# CsrA-dependent RNA accessibility landscapes prioritize new mRNA and sRNA targets and link CsrA to polyphosphate homeostasis

**DOI:** 10.64898/2026.11.18.767689

**Authors:** Ryan Buchser, Kathleen Taylor, Alexandra J. Lukasiewicz, Abigail N. Leistra, Mia K. Mihailovic, Trevor R. Simmons, José Alberto Cepeda, Lydia M. Contreras

## Abstract

The global RNA-binding protein CsrA (Carbon Storage Regulatory Protein A) regulates diverse physiological processes in *Gammaproteobacteria*, but the full set of RNAs through which it actuates and how they intersect with small RNA (sRNA) regulation and stress physiology remain incompletely defined. We asked how CsrA shapes RNA accessibility across 431 untranslated mRNA regions and 87 sRNAs in *Escherichia coli.* Using a previously published RNA accessibility profiling method (INTERFACE), we compared wild type and CsrA mutant (*csrA::kan*) strains. Permissive screening for accessibility peaks at GGA/AGA motifs nominated 87 5’UTRs, 32 3’UTRs and 28 sRNAs as candidate responsive RNAs but did not pass a permutation null (p=0.43), however it can be used to guide selection of candidates for validation. Eleven 5’UTRs had no prior experimental evidence, and all 3’UTRs and most sRNAs were nominated manually because of limited probe numbers in these targets. In reporter assays, six of nine RNAs tested showed CsrA-dependent expression *(mdtE, ribB, patA, metC, ppk and proS*) including three candidate activated-targets, although only two of five reporters from the primary candidate list responded. CsrA bound the sRNAs SsrA and RalA in vitro, and reporter assays suggested CsrA-dependent modulation of previously identified ArrS-*ribB* and GcvB-*crp* interactions. Mutating sequence elements in the *ppk* 5’ UTR altered CsrA-dependent expression, and loss of CsrA-activity was associated with reduced polyphosphate and fewer Hfq-containing HP-bodies during nitrogen starvation. Together, these findings expand the CsrA regulon and its overlap with sRNA networks, and support accessibility profiling to prioritize RNA-binding protein targets for follow-up characterization.

## Introduction

Post-transcriptional regulation is a key strategy for bacteria to rapidly respond and adapt to extracellular stress^1,2^. The Carbon Storage Regulatory (Csr) network is a major global post-transcriptional network broadly conserved across *Gammaproteobacteria* and distributed more sparsely across other classes of bacteria^3^. The Csr network is especially important to study due to the broad range of functions it regulates, including virulence and biofilm formation^4^. More recently, the role of CsrA has expanded from the study of single mRNA targets (i.e. *glgC*^5^) to its role as a regulator of broader stress response networks^6–8^, and even as a facilitator of gene regulation through biomolecular condensation^9^. CsrA regulates its RNA targets through direct binding interactions, primarily via recognition of specific structural (i.e. RNA hairpins) and sequence motifs (GGA/AGA)^10^. In *Escherichia coli* (*E. coli*), the CsrA protein is regulated by two ‘sponge’ small RNAs, CsrB and CsrC, which function by sequestering CsrA through multiple high-affinity binding sites^11^. Recent findings have identified additional small RNAs (Spot 42, McaS, FnrS, SgrS) that can also sequester CsrA^12^, albeit with lower affinity than CsrB/C. The half-lives of the CsrB/C small RNAs are regulated by the CsrD protein, which facilitates their processing in the degradosome complex depending upon the glucose levels available to the cell^13–15^. Regulation at every level of this network is tightly controlled by multiple environmental signals (mediated via sensing proteins such as BarA/UvrY^16^ and E2A^13^), as well as feedback loops that lead to a highly tuned and self-regulating system^7,16,17^. Collectively, previous literature has firmly established CsrA as a global post-transcriptional regulator affecting diverse bacterial phenotypes.

Understanding the mechanisms by which RNA-binding proteins, including CsrA, affect the function of their target RNAs remains a key challenge in the field^18,19^. Although numerous CsrA-binding RNAs have been identified^6,8,20–22^, important gaps remain concerning the coupling of identification and regulatory outcomes. Existing approaches such as motif searches^21^, CLIP-based methods^6,8,22^, WT vs knockout RNA-sequencing^6,8^, and translational reporter assays^8,20^ often identify binding events or regulatory outputs independently, making it difficult to distinguish functionally relevant interactions from incidental or non-regulatory binding. In addition, these approaches may overlook regulatory interactions that occur outside well-characterized canonical binding architectures. Most characterized CsrA interactions involve repression of translation through binding near the ribosome binding site^4^. However, a smaller subset of transcripts have been found to be activated by CsrA through diverse and less understood mechanisms^23–25^, making their identification particularly challenging using sequence-based approaches alone. Emerging evidence suggests that CsrA also interacts with bacterial sRNA-mediated regulatory networks^6,12,26^, yet the breadth of these interactions remains poorly understood. Deeper characterization of how CsrA regulates its RNA targets at multiple scales is necessary to understand the breadth of CsrA regulation in the cell.

RNA accessibility profiling provides an integrated readout of RNA-protein interactions and local RNA base pairing probabilities. We have previously demonstrated that interrogating RNA accessibility at the sequence level provides unique insight into RNA-protein interactions in bacteria at high throughput^27^, as well as in low throughput for the CsrA ‘sponge’ RNA CsrB^28^. We therefore hypothesized that accessibility profiling could be applied to identify RNA regions responsive to CsrA activity. We reasoned that these regions could nominate candidate CsrA-responsive RNAs independently of sequence-based binding site predictions. We use accessibility profiling here as a prioritization tool rather than as a predictor of regulatory outcome. Specifically, we sought to identify CsrA-dependent accessibility signatures in a high-throughput manner in *E. coli* by employing the INTERFACE method^27^ (**in** vivo **t**ranscriptional **e**longation analyzed by **R**NA-seq for **f**unctional **a**ccessibility **c**haracterization in a single **e**xperiment^27^) previously developed in the Contreras Lab. This method assesses relative levels of RNA conformational accessibility via use of RNA-based probes that capture defined 10-20nt regions along an RNA of interest. Previously, this method was successful in identifying new targets of the global RNA-binding protein Hfq^27^. As opposed to alternative methods (e.g. CLIP-seq^6^), this approach is ‘protein-agnostic’ – it measures RNA states without modifying or altering any proteins of interest beyond reducing their activity genetically. This method addresses some of the limitations discussed above by characterizing the landscape of accessibility across each RNA, as opposed to focusing on binding sites or translational outcomes, while introducing its own caveats (see Limitations). INTERFACE leverages *in vivo* structural rearrangement of antisense RNA (asRNA)-triggered toehold switches, the natural *tnaC* stalling mechanism, and rho-dependent termination to generate a signal of RNA accessibility that can be measured via RNA-sequencing (**Figure 1A**). When the asRNA probe binds to its antisense target, the ribosome binding site is opened, allowing for translation of the *tnaC* leader peptide, which stalls the ribosome^29^ (**Figure 1A**). Ribosome stalling blocks the rho-dependent termination site, allowing for elongation of the ‘elongation reporter’ sequence (**Figure 1A**). Thus, upon asRNA binding to an accessible region, the asRNA probe elongates, and the accessibility is measured via RNA-sequencing: longer transcripts correspond to more accessible probe targeting regions.

**Figure 1.**
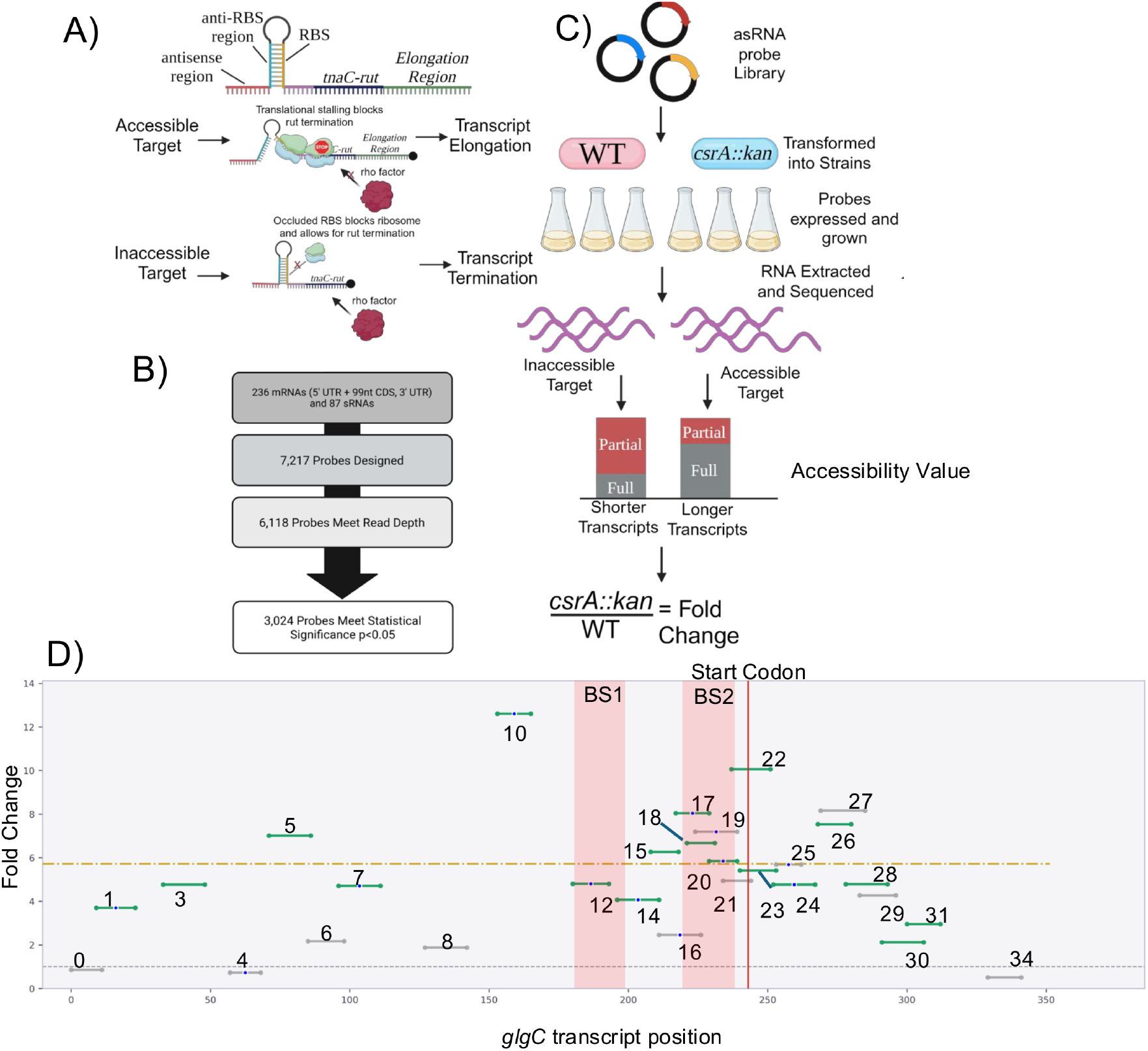
Overview of the high-throughput in vivo RNA accessibility experiment. **(A)** the INTERFACE system uses a toehold switch-based antisense RNA probe (asRNA) to detect RNA accessibility. Using a coupled transcriptional-translational reporter, the level of ribosomal occupancy is directly correlated to the length of the relevant RNA transcript. **(B)** The INTERFACE experiment was performed by transforming a library of asRNA probes into wild-type and *csrA::kan* strains (wild-type vs. reduced CsrA activity), followed by plasmid overexpression, RNA extraction and sequencing**. (C)** Probe accessibility was determined by comparing the length of each probe in both strains, thus giving an output of RNA accessibility at each region**. (D)** Accessibility map of the *glgC* 5’ UTR: Each horizontal bar in the map of the *glgC* transcript represents a single asRNA probe. The position on the x-axis signifies the location within the transcript, and the position on the y-axis represents the magnitude of the fold change between the *csrA::kan* and WT strains (fold change = *csrA::kan*/WT). Probes containing a blue dot are those that are antisense to a region containing an AGA or GGA motif, signifying a potential CsrA binding site. The color of each bar represents the statistical significance of the probe between the two strains, with green bars being statistically significant, and gray bars not significant. The vertical red bar signifies the start codon. The horizontal yellow bar indicates the mean accessibility change of the transcript. Transparent red regions represent characterized CsrA-*glgC* binding sites. Probe labels are numbered along the transcript.

In this work, we ask how CsrA activity shapes the accessibility of its candidate RNA targets in *E. coli*, and we use these accessibility landscapes to prioritize mRNAs and sRNAs for follow-up characterization. We profiled 431 UTRs and 87 sRNAs in wild type and a partial loss-of-function CsrA mutant strain (*csrA::kan*) that is characterized to retain only 10% of its activity (see Methods). The resulting data nominated candidate mRNA and sRNA targets, candidate CsrA-activated 5’UTRs and candidate CsrA-dependent sRNA-mRNA interactions. We tested a subset of these with reporter and in vitro binding assays. Finally, we examined CsrA regulation of the polyphosphate kinase (*ppk)* gene, a global stress regulator, which is essential for the production of polyphosphate (PolyP). We show that CsrA-dependent expression of a *ppk* reporter depends on specific sequence elements in the 5’UTR and coding sequence, and that loss of CsrA activity is associated with reduced polyP and fewer polyP-dependent Hfq-containing condensates ( “HP-bodies”), which are hypothesized to contribute to RNA homeostasis under stress^30^. Together, these findings broaden the CsrA regulon and its intersection with sRNA networks and illustrate the use of accessibility profiling as a prioritization approach of candidate RNA targets within a large regulatory RNA-protein network.

## Results

### Accessibility profiling of CsrA-dependent RNA landscapes recovers known CsrA-associated regions

To determine whether RNA accessibility profiles could identify CsrA binding regions, we designed a library of 7,168 asRNA probes, which cover 431 UTRs (306 5’ and 125 3’) from 236 mRNAs (including 99nt of coding sequence (CDS) for the 5’ UTRs), as well as 87 small RNAs **(Figure 1B)** to test in WT and CsrA mutant strains. These probes covered 88% and 85% of the nucleotides within the probed mRNAs and sRNAs on average, respectively. This collection of library targets includes positive control RNAs that have been shown to be regulated by CsrA (e.g. *glgC*^5^), putative targets (i.e. *ppk*^8^), and negative control targets (i.e. *fecA*^31^). Because targets were chosen largely from published CsrA datasets (see Methods), the library is enriched for RNAs with prior evidence of CsrA association, which we consider when comparing our candidates to earlier studies.

The INTERFACE experiment (**Figure 1C**) was performed in *E. coli* strains where CsrA is present (WT) and in a well-characterized reduced function mutant strain^24,25^ (*csrA::kan*) (see Methods). Evaluating the resulting RNA-seq data, we calculated an accessibility value (ratio of long reads to short reads) for each RNA probe within each strain and compared *csrA::kan* to WT to calculate a fold change in accessibility (**Figure 1C**). Of the 7,168 probes, 6,118 met our read-depth cutoff of 20 reads per probe in all replicates, and 3,024 showed a significant accessibility difference between strains (**Figure 1B**). To control for the possibility that the partial deletion of CsrA would affect transcription elongation (which would bias accessibility profiling by this method), we designed a randomized probe as control (without complementarity to the *E. coli* transcriptome) and observed a mean fold change above one that did not reach statistical significance (*p* = 0.20, n=3; **Figure S1**), although variance among replicates was large. Because most probes with significant changes were more accessible in *csrA::kan* (**Figure S2**), a global shift in reporter output in the mutant cannot be excluded. We then calculated how many probes would still show an increase in accessibility after subtracting the fold change from the scramble probe and found that 1,999 probes still showed a fold change greater than 1. To ensure that true targets were selected from any potential noise, all downstream analysis was normalized within each RNA that was probed, to select for regions that showed the highest fold changes within a given RNA (see Methods).

To validate our technique against previously published and characterized CsrA-binding controls, we first analyzed the accessibility profiles of mRNAs that have been established to directly interact with the CsrA protein via *in vitro* Electrophoretic Mobility Shift Assays (EMSAs) and footprinting assays (**Table 1**)^32–34^. We hypothesized that CsrA binding sites would be evident in the data as high accessibility probes within an RNA, or “peaks”, that contain a GGA or AGA motif, as CsrA binding in the WT strain would lower accessibility. We hypothesize that these “peaks” in accessibility between WT and *csrA::kan* may reflect CsrA binding sites or other CsrA-dependent changes in RNA structure. **Figure 1D** shows the full probe map for the *glgC* 5’UTR, where the local regions reflecting the highest fold changes in accessibility (strongest accessibility peaks) correspond to well-characterized binding sites and contain CsrA binding motifs^5^. Similar accessibility patterns were observed across additional validated targets (e.g. *pgaA, cstA*, **Figure S3**), where the largest accessibility increases frequently occurred (6 of 8 targets: *csrA*, *glgC, hfq*, *iraD, pgaA* and *sdiA*) near known GGA/AGA-containing regulatory regions. We hypothesize that the presence of additional factors such as other RNA binding proteins or ribosomes, or large changes in expression levels may account for the fact that the *rpoE* and *flhD* RNAs did not show accessibility signatures that correspond to binding. Because GGA/AGA trinucleotides are extremely common, motif presence alone may have little specificity (i.e. the screen’s signature, a high-fold-change probe overlapping a GGA/AGA site, is not highly informative beyond chance); we therefore treat accessibility as an additional information layer for ranking candidate CsrA-responsive regions and we assess below how much enrichment it provides over motif occurrence alone. As expected for negative controls, the *fecA*, *gmk*, and *phoB* 5’UTRs did not meet any of our criteria for screening and were thus not classified as candidates (**Table 2**).

**Table 1.** A list of well characterized RNA targets of CsrA that passed the filtering criteria of having a GGA/AGA peak and atleast 8 probes.

| **Gene** |
|---|
| *csrA* |
| *glgC* |
| *hfq* |
| *iraD* |
| *pgaA* |
| *sdiA* |

**Table 2.**
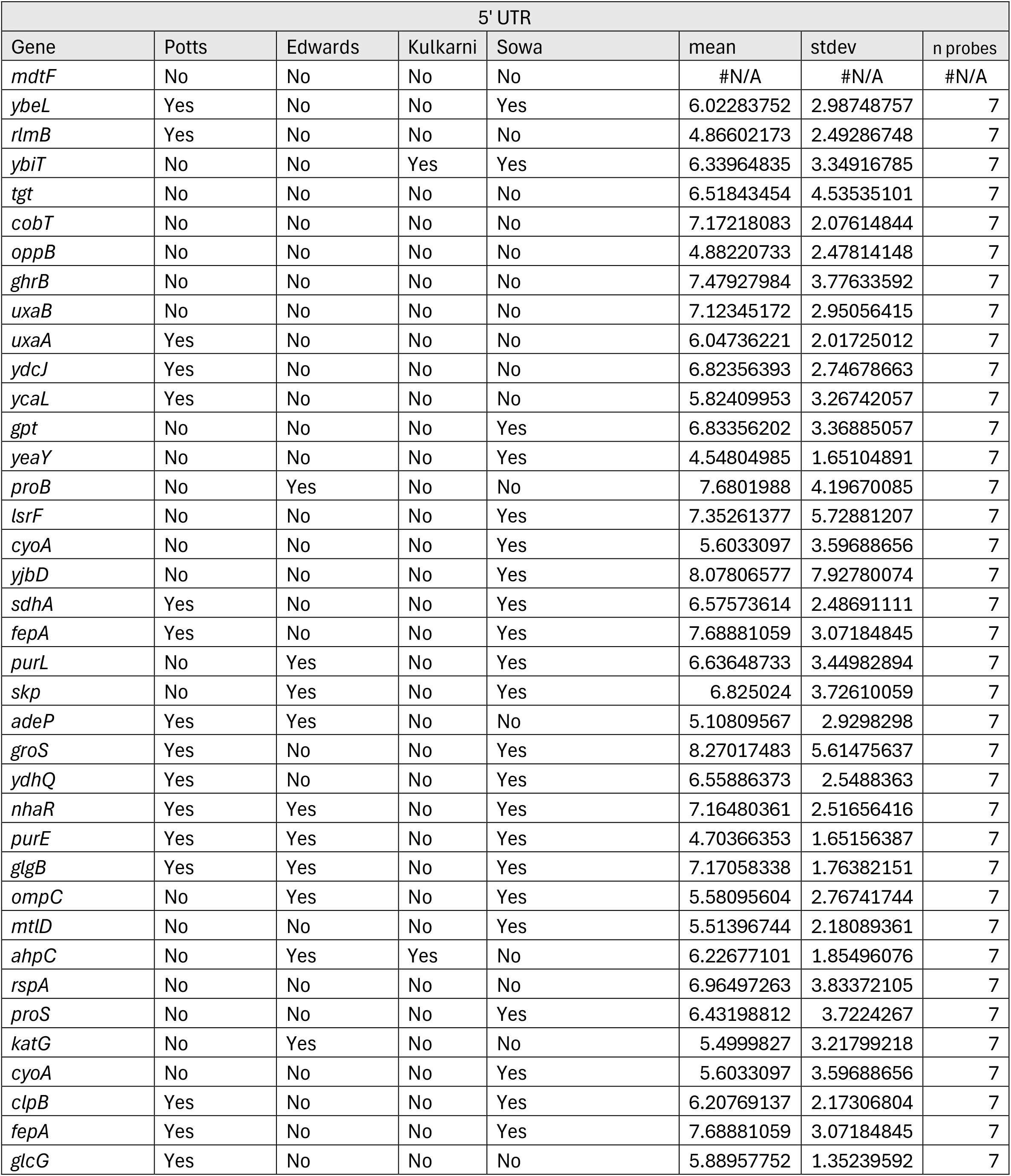
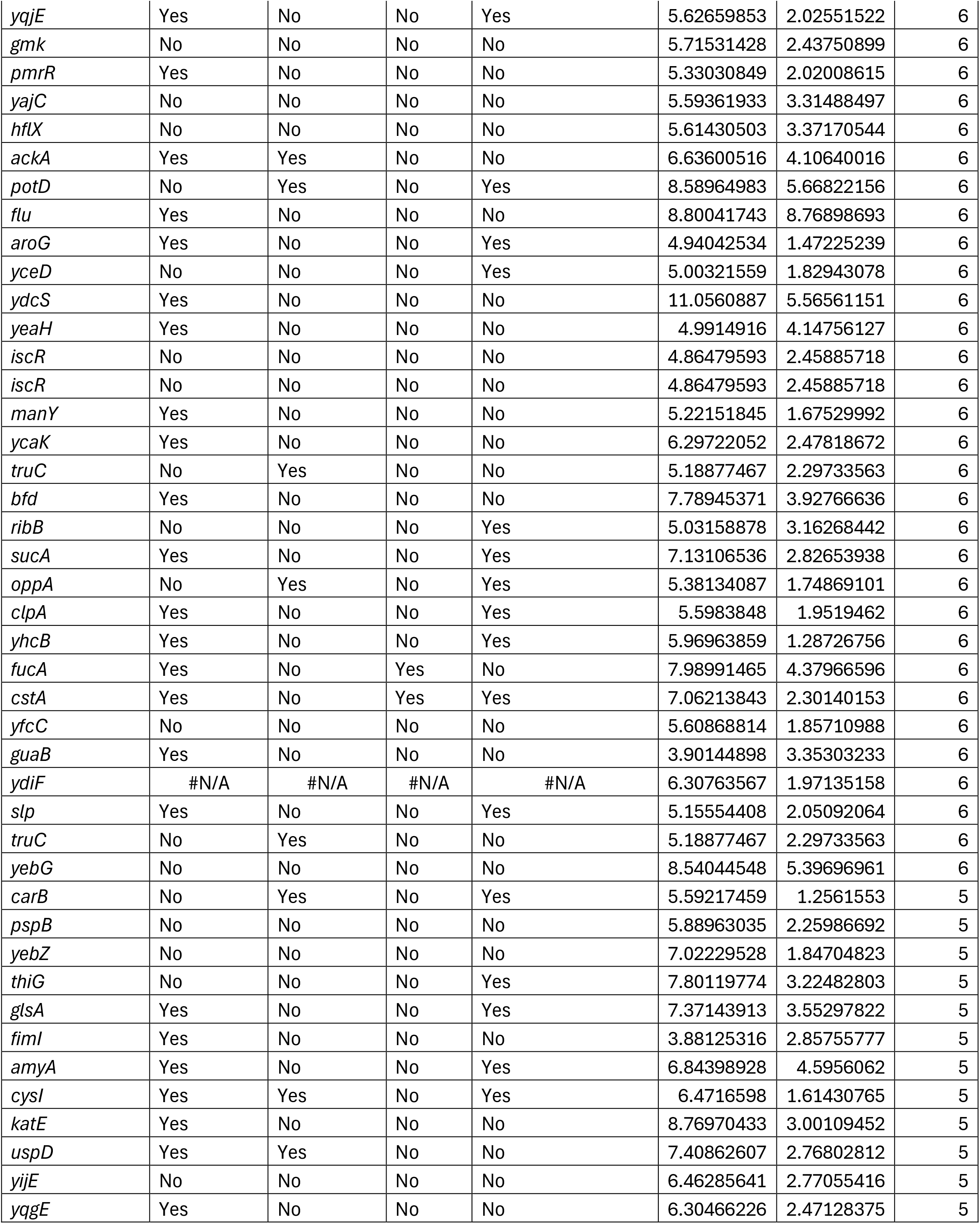
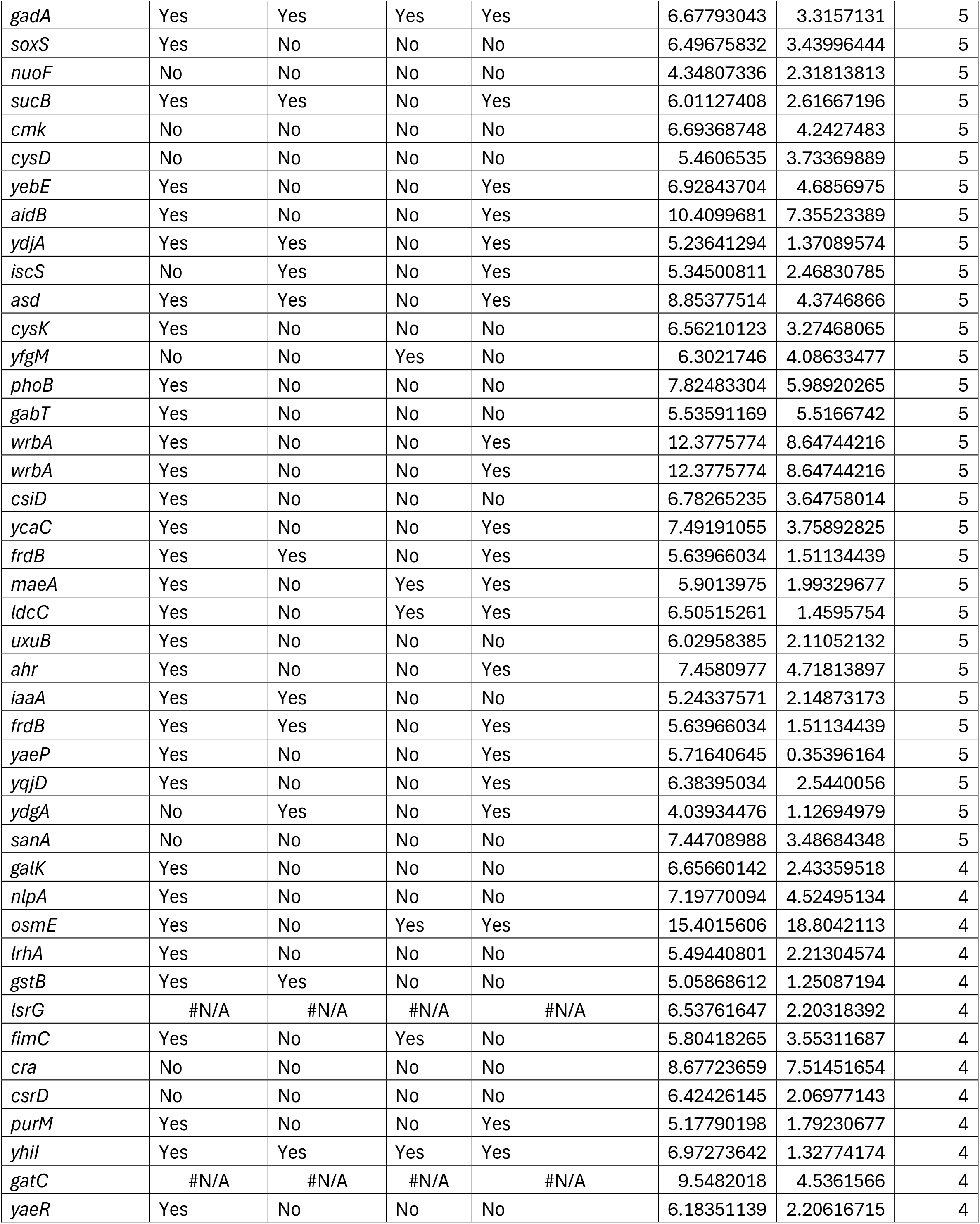
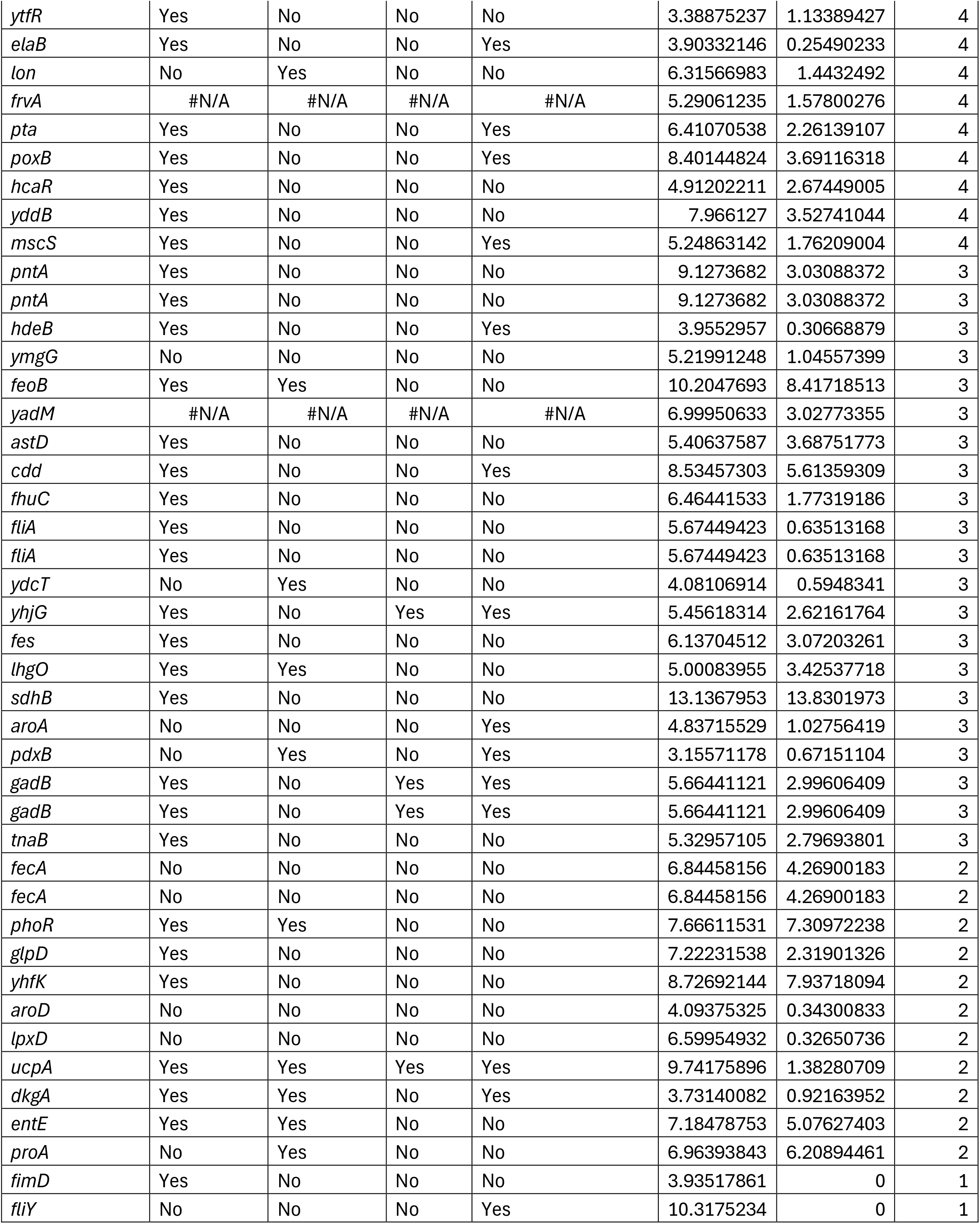
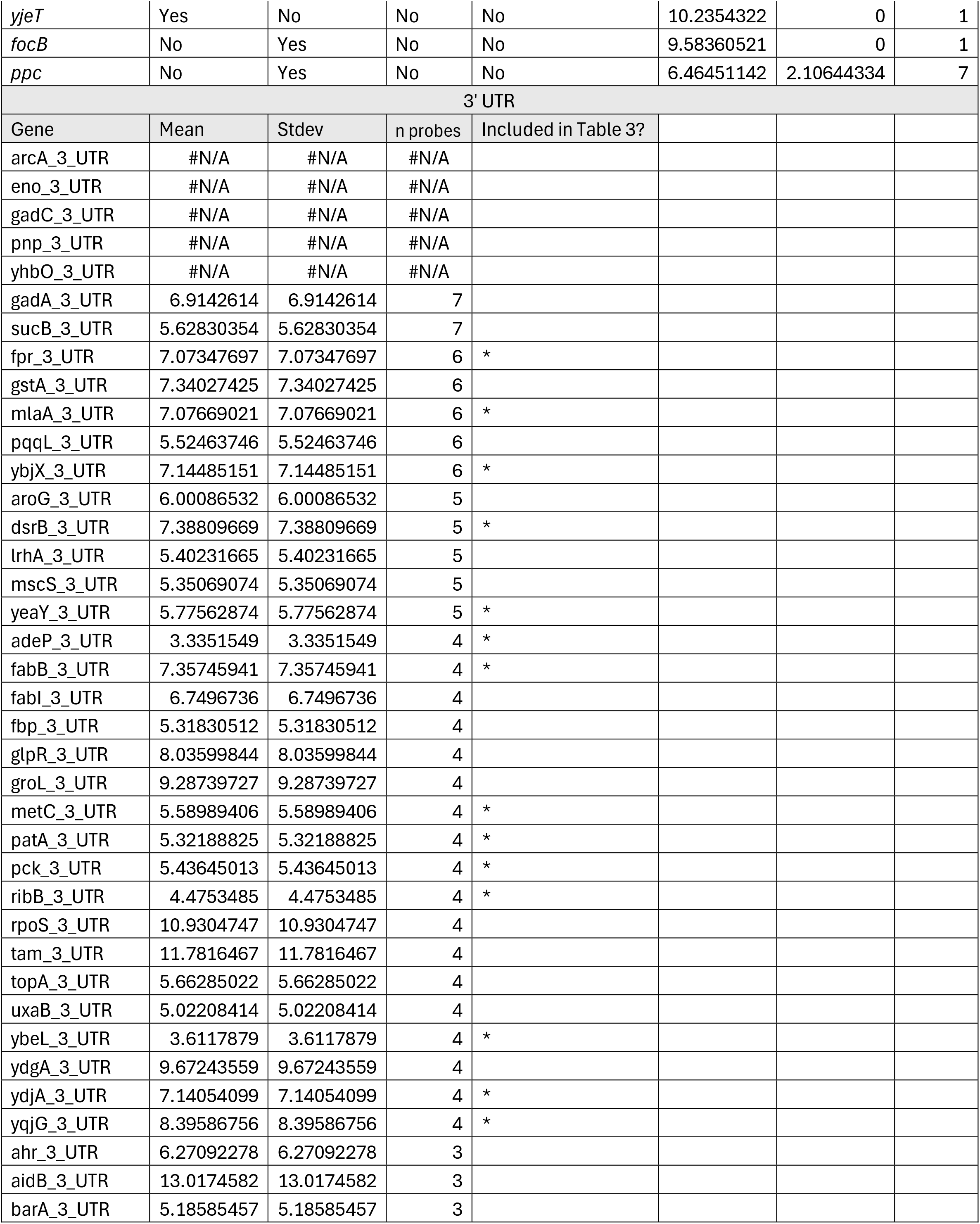
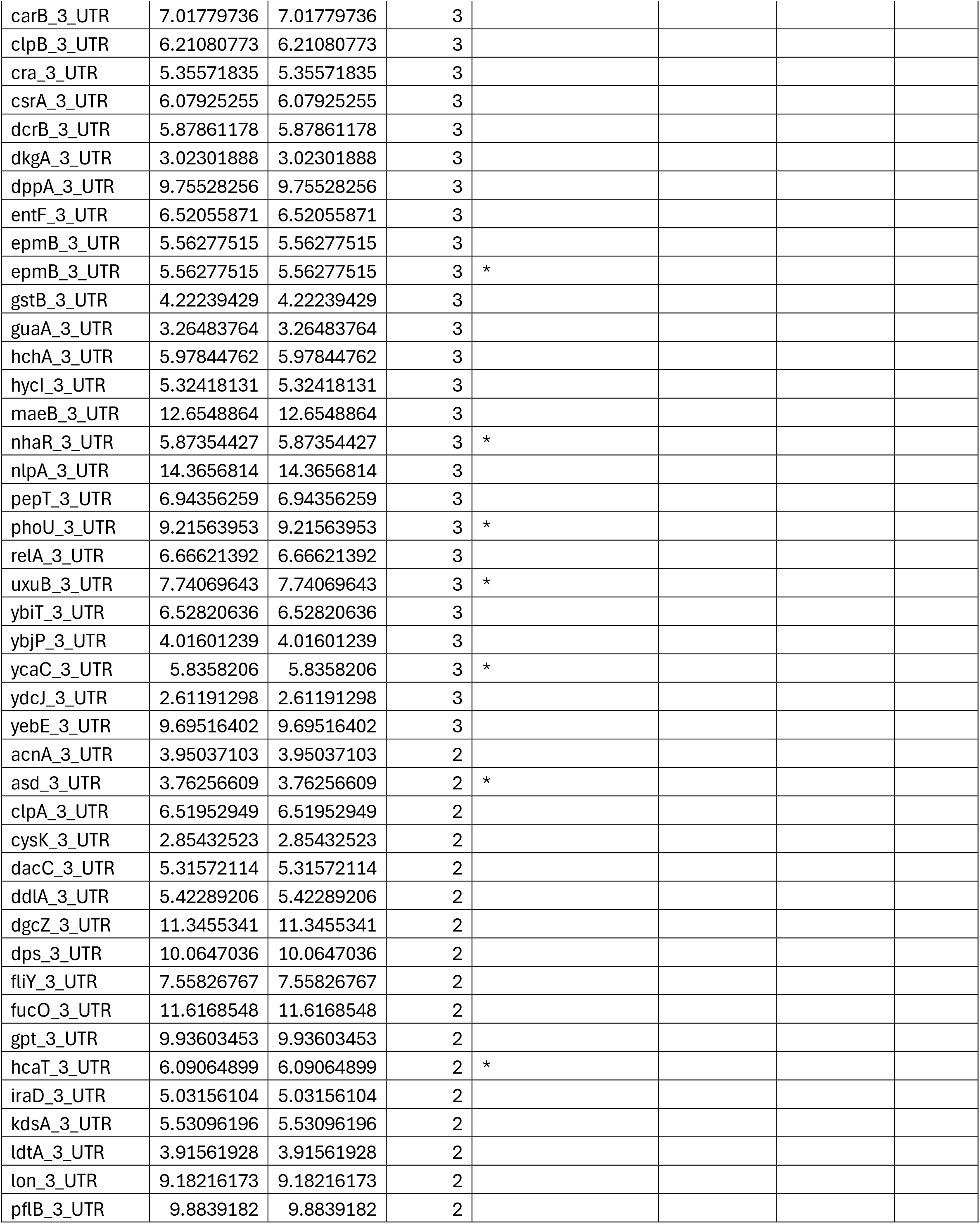
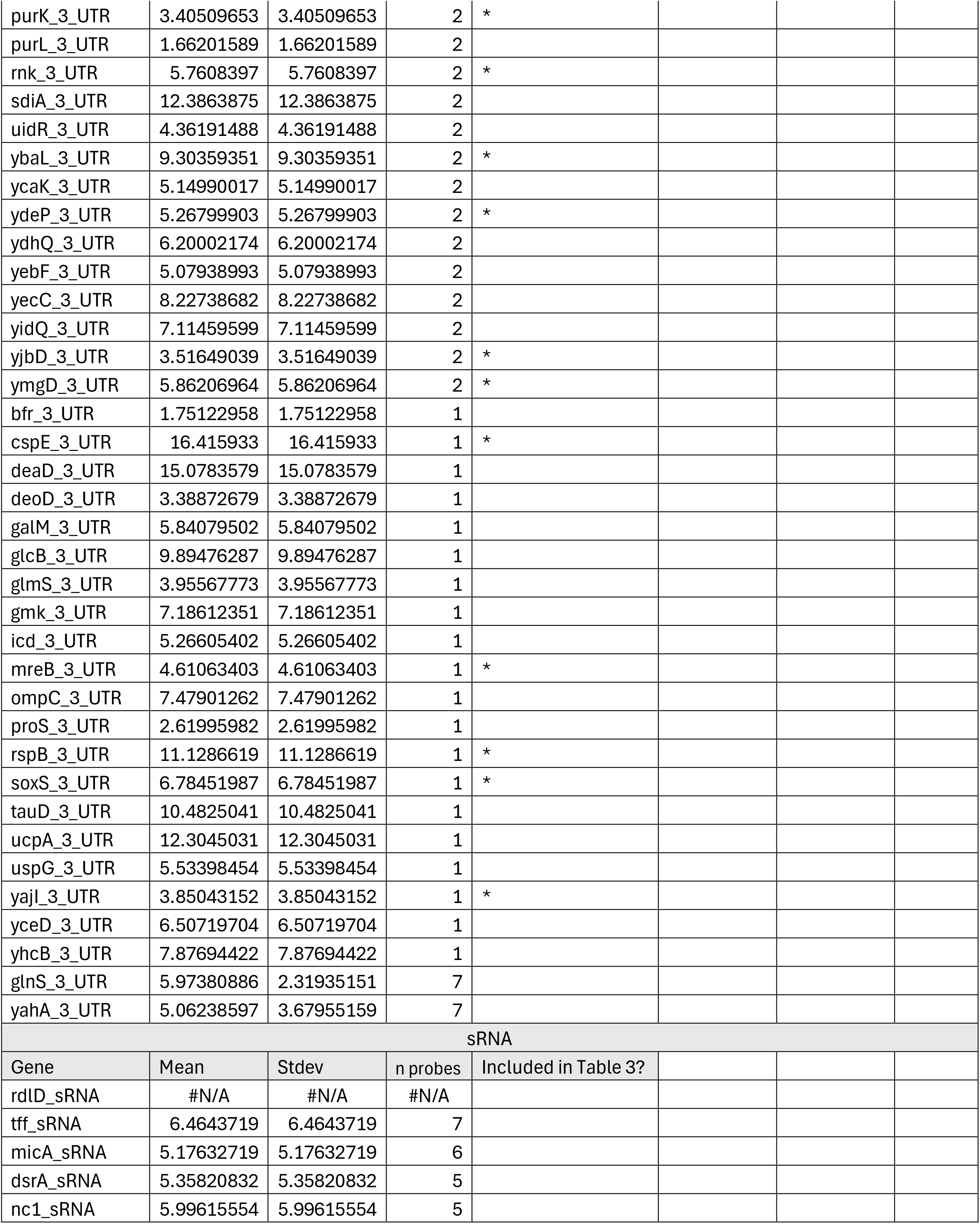
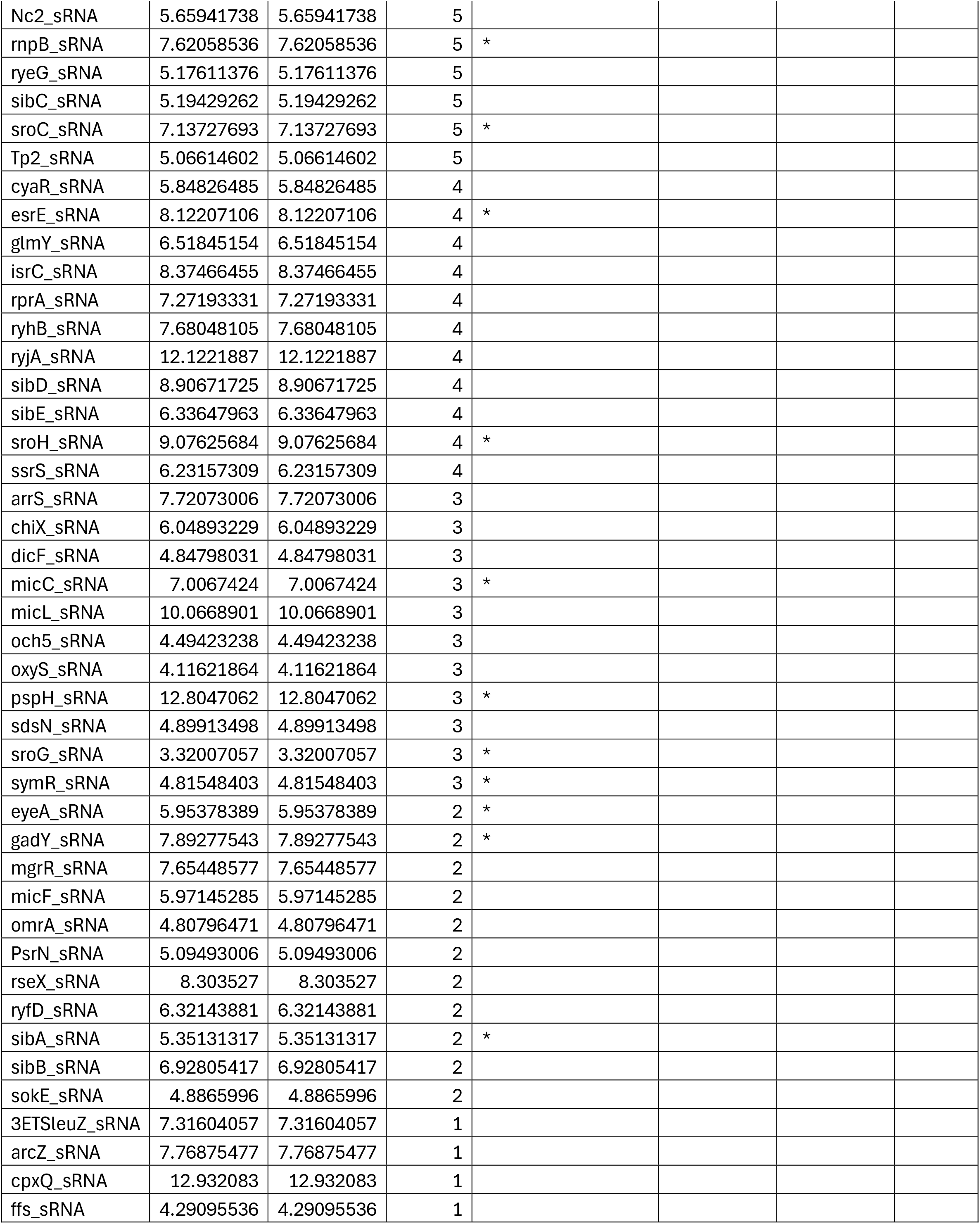
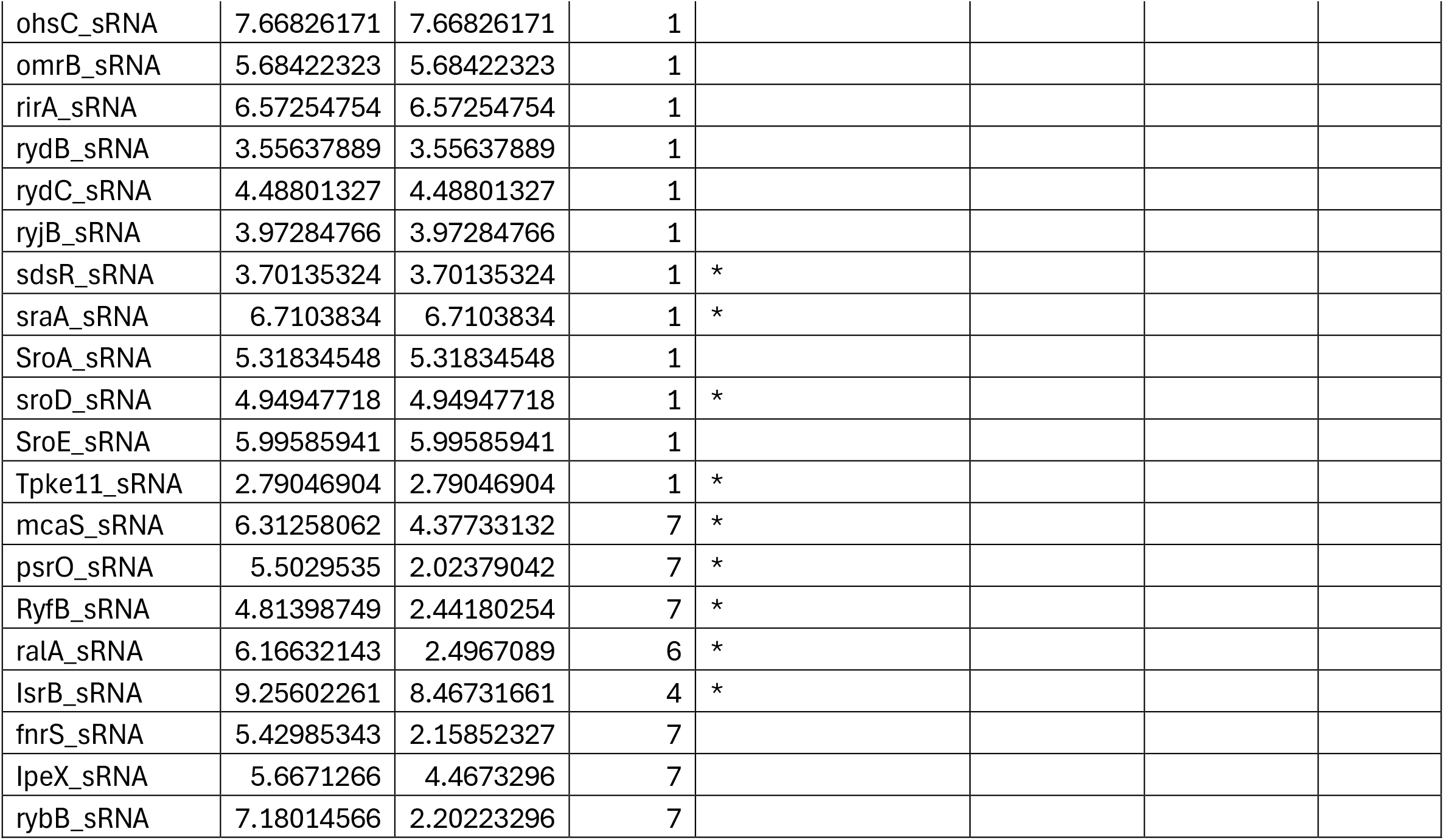
A list of all RNA targets probed in this study that did not have enough significant probes ( < 8) and thus were not considered for functional studies. Exclusions were made for those that were manually annotated and thus added to table 1 as potential targets.

|  |  |  |  | 5' UTR |  |  |  |
|---|---|---|---|---|---|---|---|
| Gene | Potts | Edwards | Kulkarni | Sowa | mean | stdev | n probes |
| *mdtF* | No | No | No | No | #N/A | #N/A | #N/A |
| *ybeL* | Yes | No | No | Yes | 6.02283752 | 2.98748757 | 7 |
| *rlmB* | Yes | No | No | No | 4.86602173 | 2.49286748 | 7 |
| *ybiT* | No | No | Yes | Yes | 6.33964835 | 3.34916785 | 7 |
| *tgt* | No | No | No | No | 6.51843454 | 4.53535101 | 7 |
| *cobT* | No | No | No | No | 7.17218083 | 2.07614844 | 7 |
| *oppB* | No | No | No | No | 4.88220733 | 2.47814148 | 7 |
| *ghrB* | No | No | No | No | 7.47927984 | 3.77633592 | 7 |
| *uxaB* | No | No | No | No | 7.12345172 | 2.95056415 | 7 |
| *uxaA* | Yes | No | No | No | 6.04736221 | 2.01725012 | 7 |
| *ydcJ* | Yes | No | No | No | 6.82356393 | 2.74678663 | 7 |
| *ycaL* | Yes | No | No | No | 5.82409953 | 3.26742057 | 7 |
| *gpt* | No | No | No | Yes | 6.83356202 | 3.36885057 | 7 |
| *yeaY* | No | No | No | Yes | 4.54804985 | 1.65104891 | 7 |
| *proB* | No | Yes | No | No | 7.6801988 | 4.19670085 | 7 |
| *lsrF* | No | No | No | Yes | 7.35261377 | 5.72881207 | 7 |
| *cyoA* | No | No | No | Yes | 5.6033097 | 3.59688656 | 7 |
| *yjbD* | No | No | No | Yes | 8.07806577 | 7.92780074 | 7 |
| *sdhA* | Yes | No | No | Yes | 6.57573614 | 2.48691111 | 7 |
| *fepA* | Yes | No | No | Yes | 7.68881059 | 3.07184845 | 7 |
| *purL* | No | Yes | No | Yes | 6.63648733 | 3.44982894 | 7 |
| *skp* | No | Yes | No | Yes | 6.825024 | 3.72610059 | 7 |
| *adeP* | Yes | Yes | No | No | 5.10809567 | 2.9298298 | 7 |
| *groS* | Yes | No | No | Yes | 8.27017483 | 5.61475637 | 7 |
| *ydhQ* | Yes | No | No | Yes | 6.55886373 | 2.5488363 | 7 |
| *nhaR* | Yes | Yes | No | Yes | 7.16480361 | 2.51656416 | 7 |
| *purE* | Yes | Yes | No | Yes | 4.70366353 | 1.65156387 | 7 |
| *glgB* | Yes | Yes | No | Yes | 7.17058338 | 1.76382151 | 7 |
| *ompC* | No | Yes | No | Yes | 5.58095604 | 2.76741744 | 7 |
| *mtlD* | No | No | No | Yes | 5.51396744 | 2.18089361 | 7 |
| *ahpC* | No | Yes | Yes | No | 6.22677101 | 1.85496076 | 7 |
| *rspA* | No | No | No | No | 6.96497263 | 3.83372105 | 7 |
| *proS* | No | No | No | Yes | 6.43198812 | 3.7224267 | 7 |
| *katG* | No | Yes | No | No | 5.4999827 | 3.21799218 | 7 |
| *cyoA* | No | No | No | Yes | 5.6033097 | 3.59688656 | 7 |
| *clpB* | Yes | No | No | Yes | 6.20769137 | 2.17306804 | 7 |
| *fepA* | Yes | No | No | Yes | 7.68881059 | 3.07184845 | 7 |
| *glcG* | Yes | No | No | No | 5.88957752 | 1.35239592 | 7 |
| *yqjE* | Yes | No | No | Yes | 5.62659853 | 2.02551522 | 6 |
| *gmk* | No | No | No | No | 5.71531428 | 2.43750899 | 6 |
| *pmrR* | Yes | No | No | No | 5.33030849 | 2.02008615 | 6 |
| *yajC* | No | No | No | No | 5.59361933 | 3.31488497 | 6 |
| *hflX* | No | No | No | No | 5.61430503 | 3.37170544 | 6 |
| *ackA* | Yes | Yes | No | No | 6.63600516 | 4.10640016 | 6 |
| *potD* | No | Yes | No | Yes | 8.58964983 | 5.66822156 | 6 |
| *flu* | Yes | No | No | No | 8.80041743 | 8.76898693 | 6 |
| *aroG* | Yes | No | No | Yes | 4.94042534 | 1.47225239 | 6 |
| *yceD* | No | No | No | Yes | 5.00321559 | 1.82943078 | 6 |
| *ydcS* | Yes | No | No | No | 11.0560887 | 5.56561151 | 6 |
| *yeaH* | Yes | No | No | No | 4.9914916 | 4.14756127 | 6 |
| *iscR* | No | No | No | No | 4.86479593 | 2.45885718 | 6 |
| *iscR* | No | No | No | No | 4.86479593 | 2.45885718 | 6 |
| *manY* | Yes | No | No | No | 5.22151845 | 1.67529992 | 6 |
| *ycaK* | Yes | No | No | No | 6.29722052 | 2.47818672 | 6 |
| *truC* | No | Yes | No | No | 5.18877467 | 2.29733563 | 6 |
| *bfd* | Yes | No | No | No | 7.78945371 | 3.92766636 | 6 |
| *ribB* | No | No | No | Yes | 5.03158878 | 3.16268442 | 6 |
| *sucA* | Yes | No | No | Yes | 7.13106536 | 2.82653938 | 6 |
| *oppA* | No | Yes | No | Yes | 5.38134087 | 1.74869101 | 6 |
| *clpA* | Yes | No | No | Yes | 5.5983848 | 1.9519462 | 6 |
| *yhcB* | Yes | No | No | Yes | 5.96963859 | 1.28726756 | 6 |
| *fucA* | Yes | No | Yes | No | 7.98991465 | 4.37966596 | 6 |
| *cstA* | Yes | No | Yes | Yes | 7.06213843 | 2.30140153 | 6 |
| *yfcC* | No | No | No | No | 5.60868814 | 1.85710988 | 6 |
| *guaB* | Yes | No | No | No | 3.90144898 | 3.35303233 | 6 |
| *ydiF* | #N/A | #N/A | #N/A | #N/A | 6.30763567 | 1.97135158 | 6 |
| *slp* | Yes | No | No | Yes | 5.15554408 | 2.05092064 | 6 |
| *truC* | No | Yes | No | No | 5.18877467 | 2.29733563 | 6 |
| *yebG* | No | No | No | No | 8.54044548 | 5.39696961 | 6 |
| *carB* | No | Yes | No | Yes | 5.59217459 | 1.2561553 | 5 |
| *pspB* | No | No | No | No | 5.88963035 | 2.25986692 | 5 |
| *yebZ* | No | No | No | No | 7.02229528 | 1.84704823 | 5 |
| *thiG* | No | No | No | Yes | 7.80119774 | 3.22482803 | 5 |
| *glsA* | Yes | No | No | Yes | 7.37143913 | 3.55297822 | 5 |
| *fimI* | Yes | No | No | No | 3.88125316 | 2.85755777 | 5 |
| *amyA* | Yes | No | No | Yes | 6.84398928 | 4.5956062 | 5 |
| *cysI* | Yes | Yes | No | Yes | 6.4716598 | 1.61430765 | 5 |
| *katE* | Yes | No | No | No | 8.76970433 | 3.00109452 | 5 |
| *uspD* | Yes | Yes | No | No | 7.40862607 | 2.76802812 | 5 |
| *yijE* | No | No | No | No | 6.46285641 | 2.77055416 | 5 |
| *yqgE* | Yes | No | No | No | 6.30466226 | 2.47128375 | 5 |
| *gadA* | Yes | Yes | Yes | Yes | 6.67793043 | 3.3157131 | 5 |
| *soxS* | Yes | No | No | No | 6.49675832 | 3.43996444 | 5 |
| *nuoF* | No | No | No | No | 4.34807336 | 2.31813813 | 5 |
| *sucB* | Yes | Yes | No | Yes | 6.01127408 | 2.61667196 | 5 |
| *cmk* | No | No | No | No | 6.69368748 | 4.2427483 | 5 |
| *cysD* | No | No | No | No | 5.4606535 | 3.73369889 | 5 |
| *yebE* | Yes | No | No | Yes | 6.92843704 | 4.6856975 | 5 |
| *aidB* | Yes | No | No | Yes | 10.4099681 | 7.35523389 | 5 |
| *ydjA* | Yes | Yes | No | Yes | 5.23641294 | 1.37089574 | 5 |
| *iscS* | No | Yes | No | Yes | 5.34500811 | 2.46830785 | 5 |
| *asd* | Yes | Yes | No | Yes | 8.85377514 | 4.3746866 | 5 |
| *cysK* | Yes | No | No | No | 6.56210123 | 3.27468065 | 5 |
| *yfgM* | No | No | Yes | No | 6.3021746 | 4.08633477 | 5 |
| *phoB* | Yes | No | No | No | 7.82483304 | 5.98920265 | 5 |
| *gabT* | Yes | No | No | No | 5.53591169 | 5.5166742 | 5 |
| *wrbA* | Yes | No | No | Yes | 12.3775774 | 8.64744216 | 5 |
| *wrbA* | Yes | No | No | Yes | 12.3775774 | 8.64744216 | 5 |
| *csiD* | Yes | No | No | No | 6.78265235 | 3.64758014 | 5 |
| *ycaC* | Yes | No | No | Yes | 7.49191055 | 3.75892825 | 5 |
| *frdB* | Yes | Yes | No | Yes | 5.63966034 | 1.51134439 | 5 |
| *maeA* | Yes | No | Yes | Yes | 5.9013975 | 1.99329677 | 5 |
| *ldcC* | Yes | No | Yes | Yes | 6.50515261 | 1.4595754 | 5 |
| *uxuB* | Yes | No | No | No | 6.02958385 | 2.11052132 | 5 |
| *ahr* | Yes | No | No | Yes | 7.4580977 | 4.71813897 | 5 |
| *iaaA* | Yes | Yes | No | No | 5.24337571 | 2.14873173 | 5 |
| *frdB* | Yes | Yes | No | Yes | 5.63966034 | 1.51134439 | 5 |
| *yaeP* | Yes | No | No | Yes | 5.71640645 | 0.35396164 | 5 |
| *yqjD* | Yes | No | No | Yes | 6.38395034 | 2.5440056 | 5 |
| *ydgA* | No | Yes | No | Yes | 4.03934476 | 1.12694979 | 5 |
| *sanA* | No | No | No | No | 7.44708988 | 3.48684348 | 5 |
| *galK* | Yes | No | No | No | 6.65660142 | 2.43359518 | 4 |
| *nlpA* | Yes | No | No | No | 7.19770094 | 4.52495134 | 4 |
| *osmE* | Yes | No | Yes | Yes | 15.4015606 | 18.8042113 | 4 |
| *lrhA* | Yes | No | No | No | 5.49440801 | 2.21304574 | 4 |
| *gstB* | Yes | Yes | No | No | 5.05868612 | 1.25087194 | 4 |
| *lsrG* | #N/A | #N/A | #N/A | #N/A | 6.53761647 | 2.20318392 | 4 |
| *fimC* | Yes | No | Yes | No | 5.80418265 | 3.55311687 | 4 |
| *cra* | No | No | No | No | 8.67723659 | 7.51451654 | 4 |
| *csrD* | No | No | No | No | 6.42426145 | 2.06977143 | 4 |
| *purM* | Yes | No | No | Yes | 5.17790198 | 1.79230677 | 4 |
| *yhiI* | Yes | Yes | Yes | Yes | 6.97273642 | 1.32774174 | 4 |
| *gatC* | #N/A | #N/A | #N/A | #N/A | 9.5482018 | 4.5361566 | 4 |
| *yaeR* | Yes | No | No | No | 6.18351139 | 2.20616715 | 4 |
| *ytfR* | Yes | No | No | No | 3.38875237 | 1.13389427 | 4 |
| *elaB* | Yes | No | No | Yes | 3.90332146 | 0.25490233 | 4 |
| *lon* | No | Yes | No | No | 6.31566983 | 1.4432492 | 4 |
| *frvA* | #N/A | #N/A | #N/A | #N/A | 5.29061235 | 1.57800276 | 4 |
| *pta* | Yes | No | No | Yes | 6.41070538 | 2.26139107 | 4 |
| *poxB* | Yes | No | No | Yes | 8.40144824 | 3.69116318 | 4 |
| *hcaR* | Yes | No | No | No | 4.91202211 | 2.67449005 | 4 |
| *yddB* | Yes | No | No | No | 7.966127 | 3.52741044 | 4 |
| *mscS* | Yes | No | No | Yes | 5.24863142 | 1.76209004 | 4 |
| *pntA* | Yes | No | No | No | 9.1273682 | 3.03088372 | 3 |
| *pntA* | Yes | No | No | No | 9.1273682 | 3.03088372 | 3 |
| *hdeB* | Yes | No | No | Yes | 3.9552957 | 0.30668879 | 3 |
| *ymgG* | No | No | No | No | 5.21991248 | 1.04557399 | 3 |
| *feoB* | Yes | Yes | No | No | 10.2047693 | 8.41718513 | 3 |
| *yadM* | #N/A | #N/A | #N/A | #N/A | 6.99950633 | 3.02773355 | 3 |
| *astD* | Yes | No | No | No | 5.40637587 | 3.68751773 | 3 |
| *cdd* | Yes | No | No | Yes | 8.53457303 | 5.61359309 | 3 |
| *fhuC* | Yes | No | No | No | 6.46441533 | 1.77319186 | 3 |
| *fliA* | Yes | No | No | No | 5.67449423 | 0.63513168 | 3 |
| *fliA* | Yes | No | No | No | 5.67449423 | 0.63513168 | 3 |
| *ydcT* | No | Yes | No | No | 4.08106914 | 0.5948341 | 3 |
| *yhjG* | Yes | No | Yes | Yes | 5.45618314 | 2.62161764 | 3 |
| *fes* | Yes | No | No | No | 6.13704512 | 3.07203261 | 3 |
| *lhgO* | Yes | Yes | No | No | 5.00083955 | 3.42537718 | 3 |
| *sdhB* | Yes | No | No | No | 13.1367953 | 13.8301973 | 3 |
| *aroA* | No | No | No | Yes | 4.83715529 | 1.02756419 | 3 |
| *pdxB* | No | Yes | No | Yes | 3.15571178 | 0.67151104 | 3 |
| *gadB* | Yes | No | Yes | Yes | 5.66441121 | 2.99606409 | 3 |
| *gadB* | Yes | No | Yes | Yes | 5.66441121 | 2.99606409 | 3 |
| *tnaB* | Yes | No | No | No | 5.32957105 | 2.79693801 | 3 |
| *fecA* | No | No | No | No | 6.84458156 | 4.26900183 | 2 |
| *fecA* | No | No | No | No | 6.84458156 | 4.26900183 | 2 |
| *phoR* | Yes | Yes | No | No | 7.66611531 | 7.30972238 | 2 |
| *glpD* | Yes | No | No | No | 7.22231538 | 2.31901326 | 2 |
| *yhfK* | Yes | No | No | No | 8.72692144 | 7.93718094 | 2 |
| *aroD* | No | No | No | No | 4.09375325 | 0.34300833 | 2 |
| *lpxD* | No | No | No | No | 6.59954932 | 0.32650736 | 2 |
| *ucpA* | Yes | Yes | Yes | Yes | 9.74175896 | 1.38280709 | 2 |
| *dkgA* | Yes | Yes | No | Yes | 3.73140082 | 0.92163952 | 2 |
| *entE* | Yes | Yes | No | No | 7.18478753 | 5.07627403 | 2 |
| *proA* | No | Yes | No | No | 6.96393843 | 6.20894461 | 2 |
| *fimD* | Yes | No | No | No | 3.93517861 | 0 | 1 |
| *fliY* | No | No | No | Yes | 10.3175234 | 0 | 1 |
| *yjeT* | Yes | No | No | No | 10.2354322 | 0 | 1 |
| *focB* | No | Yes | No | No | 9.58360521 | 0 | 1 |
| *ppc* | No | Yes | No | No | 6.46451142 | 2.10644334 | 7 |
| 3' UTR |  |  |  |  |  |  |  |
| Gene | Mean | Stdev | n probes | Included in Table 3? |  |  |  |
| arcA_3_UTR | #N/A | #N/A | #N/A |  |  |  |  |
| eno_3_UTR | #N/A | #N/A | #N/A |  |  |  |  |
| gadC_3_UTR | #N/A | #N/A | #N/A |  |  |  |  |
| pnp_3_UTR | #N/A | #N/A | #N/A |  |  |  |  |
| yhbO_3_UTR | #N/A | #N/A | #N/A |  |  |  |  |
| gadA_3_UTR | 6.9142614 | 6.9142614 | 7 |  |  |  |  |
| sucB_3_UTR | 5.62830354 | 5.62830354 | 7 |  |  |  |  |
| fpr_3_UTR | 7.07347697 | 7.07347697 | 6 | * |  |  |  |
| gstA_3_UTR | 7.34027425 | 7.34027425 | 6 |  |  |  |  |
| mlaA_3_UTR | 7.07669021 | 7.07669021 | 6 | * |  |  |  |
| pqqL_3_UTR | 5.52463746 | 5.52463746 | 6 |  |  |  |  |
| ybjX_3_UTR | 7.14485151 | 7.14485151 | 6 | * |  |  |  |
| aroG_3_UTR | 6.00086532 | 6.00086532 | 5 |  |  |  |  |
| dsrB_3_UTR | 7.38809669 | 7.38809669 | 5 | * |  |  |  |
| lrhA_3_UTR | 5.40231665 | 5.40231665 | 5 |  |  |  |  |
| mscS_3_UTR | 5.35069074 | 5.35069074 | 5 |  |  |  |  |
| yeaY_3_UTR | 5.77562874 | 5.77562874 | 5 | * |  |  |  |
| adeP_3_UTR | 3.3351549 | 3.3351549 | 4 | * |  |  |  |
| fabB_3_UTR | 7.35745941 | 7.35745941 | 4 | * |  |  |  |
| fabI_3_UTR | 6.7496736 | 6.7496736 | 4 |  |  |  |  |
| fbp_3_UTR | 5.31830512 | 5.31830512 | 4 |  |  |  |  |
| glpR_3_UTR | 8.03599844 | 8.03599844 | 4 |  |  |  |  |
| groL_3_UTR | 9.28739727 | 9.28739727 | 4 |  |  |  |  |
| metC_3_UTR | 5.58989406 | 5.58989406 | 4 | * |  |  |  |
| patA_3_UTR | 5.32188825 | 5.32188825 | 4 | * |  |  |  |
| pck_3_UTR | 5.43645013 | 5.43645013 | 4 | * |  |  |  |
| ribB_3_UTR | 4.4753485 | 4.4753485 | 4 | * |  |  |  |
| rpoS_3_UTR | 10.9304747 | 10.9304747 | 4 |  |  |  |  |
| tam_3_UTR | 11.7816467 | 11.7816467 | 4 |  |  |  |  |
| topA_3_UTR | 5.66285022 | 5.66285022 | 4 |  |  |  |  |
| uxaB_3_UTR | 5.02208414 | 5.02208414 | 4 |  |  |  |  |
| ybeL_3_UTR | 3.6117879 | 3.6117879 | 4 | * |  |  |  |
| ydgA_3_UTR | 9.67243559 | 9.67243559 | 4 |  |  |  |  |
| ydjA_3_UTR | 7.14054099 | 7.14054099 | 4 | * |  |  |  |
| yqjG_3_UTR | 8.39586756 | 8.39586756 | 4 | * |  |  |  |
| ahr_3_UTR | 6.27092278 | 6.27092278 | 3 |  |  |  |  |
| aidB_3_UTR | 13.0174582 | 13.0174582 | 3 |  |  |  |  |
| barA_3_UTR | 5.18585457 | 5.18585457 | 3 |  |  |  |  |
| carB_3_UTR | 7.01779736 | 7.01779736 | 3 |  |  |  |  |
| clpB_3_UTR | 6.21080773 | 6.21080773 | 3 |  |  |  |  |
| cra_3_UTR | 5.35571835 | 5.35571835 | 3 |  |  |  |  |
| csrA_3_UTR | 6.07925255 | 6.07925255 | 3 |  |  |  |  |
| dcrB_3_UTR | 5.87861178 | 5.87861178 | 3 |  |  |  |  |
| dkgA_3_UTR | 3.02301888 | 3.02301888 | 3 |  |  |  |  |
| dppA_3_UTR | 9.75528256 | 9.75528256 | 3 |  |  |  |  |
| entF_3_UTR | 6.52055871 | 6.52055871 | 3 |  |  |  |  |
| epmB_3_UTR | 5.56277515 | 5.56277515 | 3 |  |  |  |  |
| epmB_3_UTR | 5.56277515 | 5.56277515 | 3 | * |  |  |  |
| gstB_3_UTR | 4.22239429 | 4.22239429 | 3 |  |  |  |  |
| guaA_3_UTR | 3.26483764 | 3.26483764 | 3 |  |  |  |  |
| hchA_3_UTR | 5.97844762 | 5.97844762 | 3 |  |  |  |  |
| hycI_3_UTR | 5.32418131 | 5.32418131 | 3 |  |  |  |  |
| maeB_3_UTR | 12.6548864 | 12.6548864 | 3 |  |  |  |  |
| nhaR_3_UTR | 5.87354427 | 5.87354427 | 3 | * |  |  |  |
| nlpA_3_UTR | 14.3656814 | 14.3656814 | 3 |  |  |  |  |
| pepT_3_UTR | 6.94356259 | 6.94356259 | 3 |  |  |  |  |
| phoU_3_UTR | 9.21563953 | 9.21563953 | 3 | * |  |  |  |
| relA_3_UTR | 6.66621392 | 6.66621392 | 3 |  |  |  |  |
| uxuB_3_UTR | 7.74069643 | 7.74069643 | 3 | * |  |  |  |
| ybiT_3_UTR | 6.52820636 | 6.52820636 | 3 |  |  |  |  |
| ybjP_3_UTR | 4.01601239 | 4.01601239 | 3 |  |  |  |  |
| ycaC_3_UTR | 5.8358206 | 5.8358206 | 3 | * |  |  |  |
| ydcJ_3_UTR | 2.61191298 | 2.61191298 | 3 |  |  |  |  |
| yebE_3_UTR | 9.69516402 | 9.69516402 | 3 |  |  |  |  |
| acnA_3_UTR | 3.95037103 | 3.95037103 | 2 |  |  |  |  |
| asd_3_UTR | 3.76256609 | 3.76256609 | 2 | * |  |  |  |
| clpA_3_UTR | 6.51952949 | 6.51952949 | 2 |  |  |  |  |
| cysK_3_UTR | 2.85432523 | 2.85432523 | 2 |  |  |  |  |
| dacC_3_UTR | 5.31572114 | 5.31572114 | 2 |  |  |  |  |
| ddlA_3_UTR | 5.42289206 | 5.42289206 | 2 |  |  |  |  |
| dgcZ_3_UTR | 11.3455341 | 11.3455341 | 2 |  |  |  |  |
| dps_3_UTR | 10.0647036 | 10.0647036 | 2 |  |  |  |  |
| fliY_3_UTR | 7.55826767 | 7.55826767 | 2 |  |  |  |  |
| fucO_3_UTR | 11.6168548 | 11.6168548 | 2 |  |  |  |  |
| gpt_3_UTR | 9.93603453 | 9.93603453 | 2 |  |  |  |  |
| hcaT_3_UTR | 6.09064899 | 6.09064899 | 2 | * |  |  |  |
| iraD_3_UTR | 5.03156104 | 5.03156104 | 2 |  |  |  |  |
| kdsA_3_UTR | 5.53096196 | 5.53096196 | 2 |  |  |  |  |
| ldtA_3_UTR | 3.91561928 | 3.91561928 | 2 |  |  |  |  |
| lon_3_UTR | 9.18216173 | 9.18216173 | 2 |  |  |  |  |
| pflB_3_UTR | 9.8839182 | 9.8839182 | 2 |  |  |  |  |
| purK_3_UTR | 3.40509653 | 3.40509653 | 2 | * |  |  |  |
| purL_3_UTR | 1.66201589 | 1.66201589 | 2 |  |  |  |  |
| rnk_3_UTR | 5.7608397 | 5.7608397 | 2 | * |  |  |  |
| sdiA_3_UTR | 12.3863875 | 12.3863875 | 2 |  |  |  |  |
| uidR_3_UTR | 4.36191488 | 4.36191488 | 2 |  |  |  |  |
| ybaL_3_UTR | 9.30359351 | 9.30359351 | 2 | * |  |  |  |
| ycaK_3_UTR | 5.14990017 | 5.14990017 | 2 |  |  |  |  |
| ydeP_3_UTR | 5.26799903 | 5.26799903 | 2 | * |  |  |  |
| ydhQ_3_UTR | 6.20002174 | 6.20002174 | 2 |  |  |  |  |
| yebF_3_UTR | 5.07938993 | 5.07938993 | 2 |  |  |  |  |
| yecC_3_UTR | 8.22738682 | 8.22738682 | 2 |  |  |  |  |
| yidQ_3_UTR | 7.11459599 | 7.11459599 | 2 |  |  |  |  |
| yjbD_3_UTR | 3.51649039 | 3.51649039 | 2 | * |  |  |  |
| ymgD_3_UTR | 5.86206964 | 5.86206964 | 2 | * |  |  |  |
| bfr_3_UTR | 1.75122958 | 1.75122958 | 1 |  |  |  |  |
| cspE_3_UTR | 16.415933 | 16.415933 | 1 | * |  |  |  |
| deaD_3_UTR | 15.0783579 | 15.0783579 | 1 |  |  |  |  |
| deoD_3_UTR | 3.38872679 | 3.38872679 | 1 |  |  |  |  |
| galM_3_UTR | 5.84079502 | 5.84079502 | 1 |  |  |  |  |
| glcB_3_UTR | 9.89476287 | 9.89476287 | 1 |  |  |  |  |
| glmS_3_UTR | 3.95567773 | 3.95567773 | 1 |  |  |  |  |
| gmk_3_UTR | 7.18612351 | 7.18612351 | 1 |  |  |  |  |
| icd_3_UTR | 5.26605402 | 5.26605402 | 1 |  |  |  |  |
| mreB_3_UTR | 4.61063403 | 4.61063403 | 1 | * |  |  |  |
| ompC_3_UTR | 7.47901262 | 7.47901262 | 1 |  |  |  |  |
| proS_3_UTR | 2.61995982 | 2.61995982 | 1 |  |  |  |  |
| rspB_3_UTR | 11.1286619 | 11.1286619 | 1 | * |  |  |  |
| soxS_3_UTR | 6.78451987 | 6.78451987 | 1 | * |  |  |  |
| tauD_3_UTR | 10.4825041 | 10.4825041 | 1 |  |  |  |  |
| ucpA_3_UTR | 12.3045031 | 12.3045031 | 1 |  |  |  |  |
| uspG_3_UTR | 5.53398454 | 5.53398454 | 1 |  |  |  |  |
| yajI_3_UTR | 3.85043152 | 3.85043152 | 1 | * |  |  |  |
| yceD_3_UTR | 6.50719704 | 6.50719704 | 1 |  |  |  |  |
| yhcB_3_UTR | 7.87694422 | 7.87694422 | 1 |  |  |  |  |
| glnS_3_UTR | 5.97380886 | 2.31935151 | 7 |  |  |  |  |
| yahA_3_UTR | 5.06238597 | 3.67955159 | 7 |  |  |  |  |
| sRNA |  |  |  |  |  |  |  |
| Gene | Mean | Stdev | n probes | Included in Table 3? |  |  |  |
| rdlD_sRNA | #N/A | #N/A | #N/A |  |  |  |  |
| tff_sRNA | 6.4643719 | 6.4643719 | 7 |  |  |  |  |
| micA_sRNA | 5.17632719 | 5.17632719 | 6 |  |  |  |  |
| dsrA_sRNA | 5.35820832 | 5.35820832 | 5 |  |  |  |  |
| nc1_sRNA | 5.99615554 | 5.99615554 | 5 |  |  |  |  |
| Nc2_sRNA | 5.65941738 | 5.65941738 | 5 |  |  |  |  |
| rnpB_sRNA | 7.62058536 | 7.62058536 | 5 | * |  |  |  |
| ryeG_sRNA | 5.17611376 | 5.17611376 | 5 |  |  |  |  |
| sibC_sRNA | 5.19429262 | 5.19429262 | 5 |  |  |  |  |
| sroC_sRNA | 7.13727693 | 7.13727693 | 5 | * |  |  |  |
| Tp2_sRNA | 5.06614602 | 5.06614602 | 5 |  |  |  |  |
| cyaR_sRNA | 5.84826485 | 5.84826485 | 4 |  |  |  |  |
| esrE_sRNA | 8.12207106 | 8.12207106 | 4 | * |  |  |  |
| glmY_sRNA | 6.51845154 | 6.51845154 | 4 |  |  |  |  |
| isrC_sRNA | 8.37466455 | 8.37466455 | 4 |  |  |  |  |
| rprA_sRNA | 7.27193331 | 7.27193331 | 4 |  |  |  |  |
| ryhB_sRNA | 7.68048105 | 7.68048105 | 4 |  |  |  |  |
| ryjA_sRNA | 12.1221887 | 12.1221887 | 4 |  |  |  |  |
| sibD_sRNA | 8.90671725 | 8.90671725 | 4 |  |  |  |  |
| sibE_sRNA | 6.33647963 | 6.33647963 | 4 |  |  |  |  |
| sroH_sRNA | 9.07625684 | 9.07625684 | 4 | * |  |  |  |
| ssrS_sRNA | 6.23157309 | 6.23157309 | 4 |  |  |  |  |
| arrS_sRNA | 7.72073006 | 7.72073006 | 3 |  |  |  |  |
| chiX_sRNA | 6.04893229 | 6.04893229 | 3 |  |  |  |  |
| dicF_sRNA | 4.84798031 | 4.84798031 | 3 |  |  |  |  |
| micC_sRNA | 7.0067424 | 7.0067424 | 3 | * |  |  |  |
| micL_sRNA | 10.0668901 | 10.0668901 | 3 |  |  |  |  |
| och5_sRNA | 4.49423238 | 4.49423238 | 3 |  |  |  |  |
| oxyS_sRNA | 4.11621864 | 4.11621864 | 3 |  |  |  |  |
| pspH_sRNA | 12.8047062 | 12.8047062 | 3 | * |  |  |  |
| sdsN_sRNA | 4.89913498 | 4.89913498 | 3 |  |  |  |  |
| sroG_sRNA | 3.32007057 | 3.32007057 | 3 | * |  |  |  |
| symR_sRNA | 4.81548403 | 4.81548403 | 3 | * |  |  |  |
| eyeA_sRNA | 5.95378389 | 5.95378389 | 2 | * |  |  |  |
| gadY_sRNA | 7.89277543 | 7.89277543 | 2 | * |  |  |  |
| mgrR_sRNA | 7.65448577 | 7.65448577 | 2 |  |  |  |  |
| micF_sRNA | 5.97145285 | 5.97145285 | 2 |  |  |  |  |
| omrA_sRNA | 4.80796471 | 4.80796471 | 2 |  |  |  |  |
| PsrN_sRNA | 5.09493006 | 5.09493006 | 2 |  |  |  |  |
| rseX_sRNA | 8.303527 | 8.303527 | 2 |  |  |  |  |
| ryfD_sRNA | 6.32143881 | 6.32143881 | 2 |  |  |  |  |
| sibA_sRNA | 5.35131317 | 5.35131317 | 2 | * |  |  |  |
| sibB_sRNA | 6.92805417 | 6.92805417 | 2 |  |  |  |  |
| sokE_sRNA | 4.8865996 | 4.8865996 | 2 |  |  |  |  |
| 3ETSleuZ_sRNA | 7.31604057 | 7.31604057 | 1 |  |  |  |  |
| arcZ_sRNA | 7.76875477 | 7.76875477 | 1 |  |  |  |  |
| cpxQ_sRNA | 12.932083 | 12.932083 | 1 |  |  |  |  |
| ffs_sRNA | 4.29095536 | 4.29095536 | 1 |  |  |  |  |
| ohsC_sRNA | 7.66826171 | 7.66826171 | 1 |  |  |  |  |
| omrB_sRNA | 5.68422323 | 5.68422323 | 1 |  |  |  |  |
| rirA_sRNA | 6.57254754 | 6.57254754 | 1 |  |  |  |  |
| rydB_sRNA | 3.55637889 | 3.55637889 | 1 |  |  |  |  |
| rydC_sRNA | 4.48801327 | 4.48801327 | 1 |  |  |  |  |
| ryjB_sRNA | 3.97284766 | 3.97284766 | 1 |  |  |  |  |
| sdsR_sRNA | 3.70135324 | 3.70135324 | 1 | * |  |  |  |
| sraA_sRNA | 6.7103834 | 6.7103834 | 1 | * |  |  |  |
| SroA_sRNA | 5.31834548 | 5.31834548 | 1 |  |  |  |  |
| sroD_sRNA | 4.94947718 | 4.94947718 | 1 | * |  |  |  |
| SroE_sRNA | 5.99585941 | 5.99585941 | 1 |  |  |  |  |
| Tpke11_sRNA | 2.79046904 | 2.79046904 | 1 | * |  |  |  |
| mcaS_sRNA | 6.31258062 | 4.37733132 | 7 | * |  |  |  |
| psrO_sRNA | 5.5029535 | 2.02379042 | 7 | * |  |  |  |
| RyfB_sRNA | 4.81398749 | 2.44180254 | 7 | * |  |  |  |
| ralA_sRNA | 6.16632143 | 2.4967089 | 6 | * |  |  |  |
| IsrB_sRNA | 9.25602261 | 8.46731661 | 4 | * |  |  |  |
| fnrS_sRNA | 5.42985343 | 2.15852327 | 7 |  |  |  |  |
| IpeX_sRNA | 5.6671266 | 4.4673296 | 7 |  |  |  |  |
| rybB_sRNA | 7.18014566 | 2.20223296 | 7 |  |  |  |  |

In the positive controls (*glgC*, *flhD, rpoE, pgaA, cstA, iraD, hfq, sdiA*), the accessibility maps show position-specific features at footprinted binding sites **(Figure S3).** For instance, in *pgaA* probes that overlap previously footprinted regions^35^ near the start codon are amongst the highest-fold-change probes in the RNA (**Figure S3**, *pgaA* probes 19-21). In *glgC,* probes overlapping previously footprinted regions^5^ are above the RNA’s mean fold change, although the largest peak lies upstream (**Figure 1D**). In other controls, such as *iraD* and *hfq,* the largest peaks are elsewhere or not clearly separated from neighboring probes (**Figure S3**, *iraD* probe 20, *hfq* probe 25). We therefore quantify putative CsrA binding interactions at the site level and do not rely on visual inspection.

### A permissive motif-and-peak screen nominates candidate CsrA targets

Having shown that accessibility profiles overlap known CsrA-regulated regions, we asked whether they could prioritize previously uncharacterized candidate targets. Based upon previous analysis of similar RNA accessibility data studying the RNA binding protein Hfq^27^, we selected the top 25% of probes by fold change within each 5’ UTR, 3’ UTR, or sRNA as potential “peaks” of binding as a starting point. To confidently calculate the top 25% of probes, we first filtered for all RNAs that contained at least 8 probes that showed a significant change in accessibility between *csrA::kan* and WT. From these RNAs with the requisite number of probes, we selected probes that were within the top 25% of fold change and antisense to a GGA or AGA site. Finally, if these motif-containing probes were near to the start codon (-100 to +20 nts), the 5’ UTR was selected as a putative target (**Figure S4**). Using this filtering method, we identified 87 candidate 5’UTRs, 32 candidate 3’UTRs and 28 candidate sRNAs (**Table 3**). Because shorter RNAs cannot easily accommodate eight significant probes in library design, all the 32 3’UTRs and 20 of the 28 sRNAs were nominated by manual inspection of motif-containing, high-accessibility probes (“Manual annotation” in **Table 3**). In order to account for the odds of identifying the CsrA motif at random, w tested this screening method relative to a permutation null hypothesis in which probe sequences were randomly reassigned to within each target (20,000 permutations, probe number, fold changes and the motif count per target were fixed), the screen called 87 of the 132 (∼66%) targets with at least eight significant probes vs 85.7 expected by chance (1.02-fold enrichment; p=0.43; **Figure S5**). That is, about 98% of calls are expected without association between accessibility peaks and motif position as if the motifs were randomly distributed across the RNA sequence). This is expected because a GGA or AGA falls in roughly a third of the probes (33.7%) and the library was assembled from putative CsrA targets. Making the 25% fold-change cutoff more stringent (top 20%, 15%, 10%, 5% of probes) did not achieve any significant enrichment (**Table S1**). We did, however, observe that within our positive control targets there appeared to be regions of high AU content surrounding the GGA and AGA motifs, consistent with previous literature^36^. We therefore limited our use of the screen as a sensitive method to identify candidate regions with high CsrA dependency and use experimental validation to ultimately distinguish whether a candidate is a CsrA target.

**Table 3.**
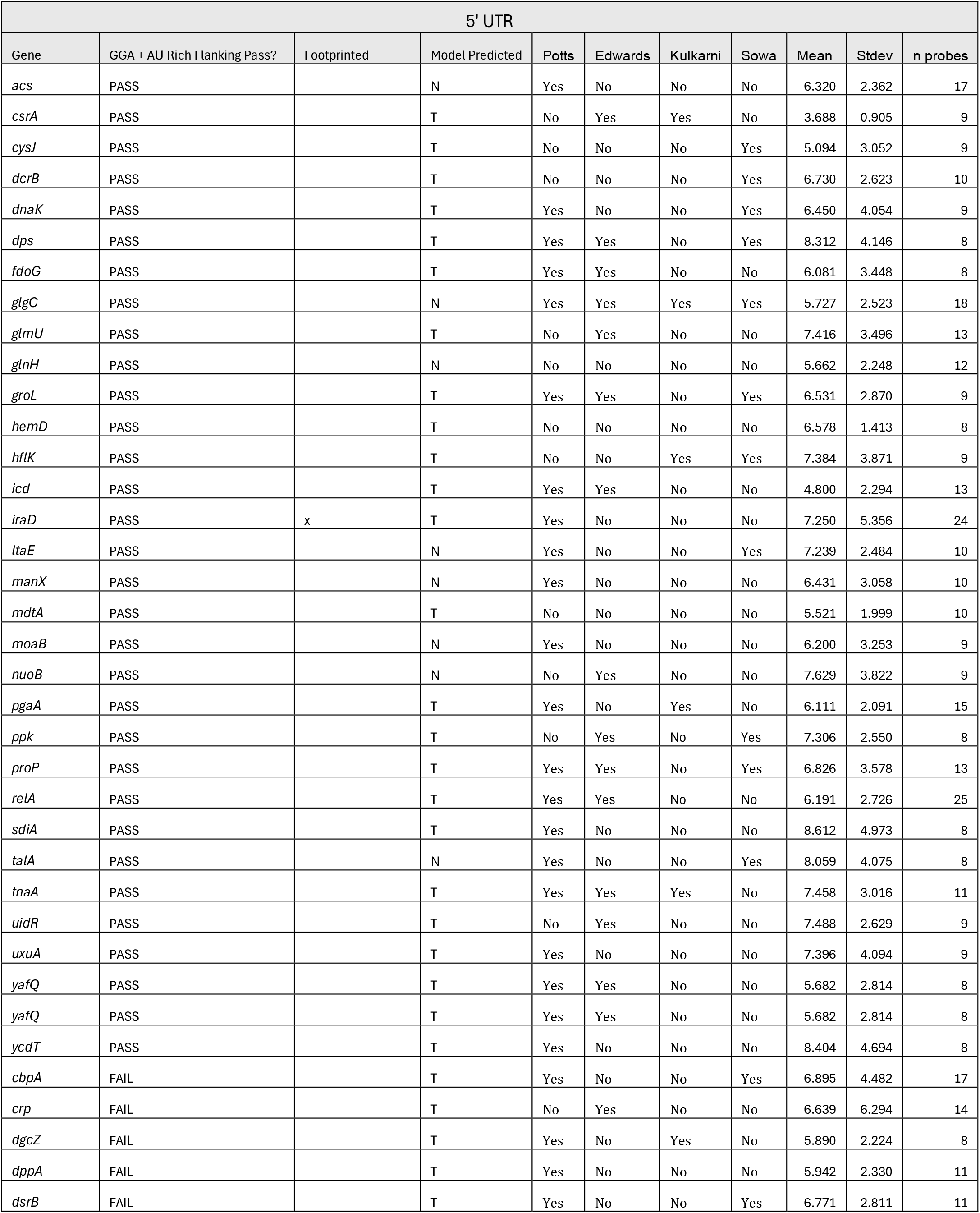
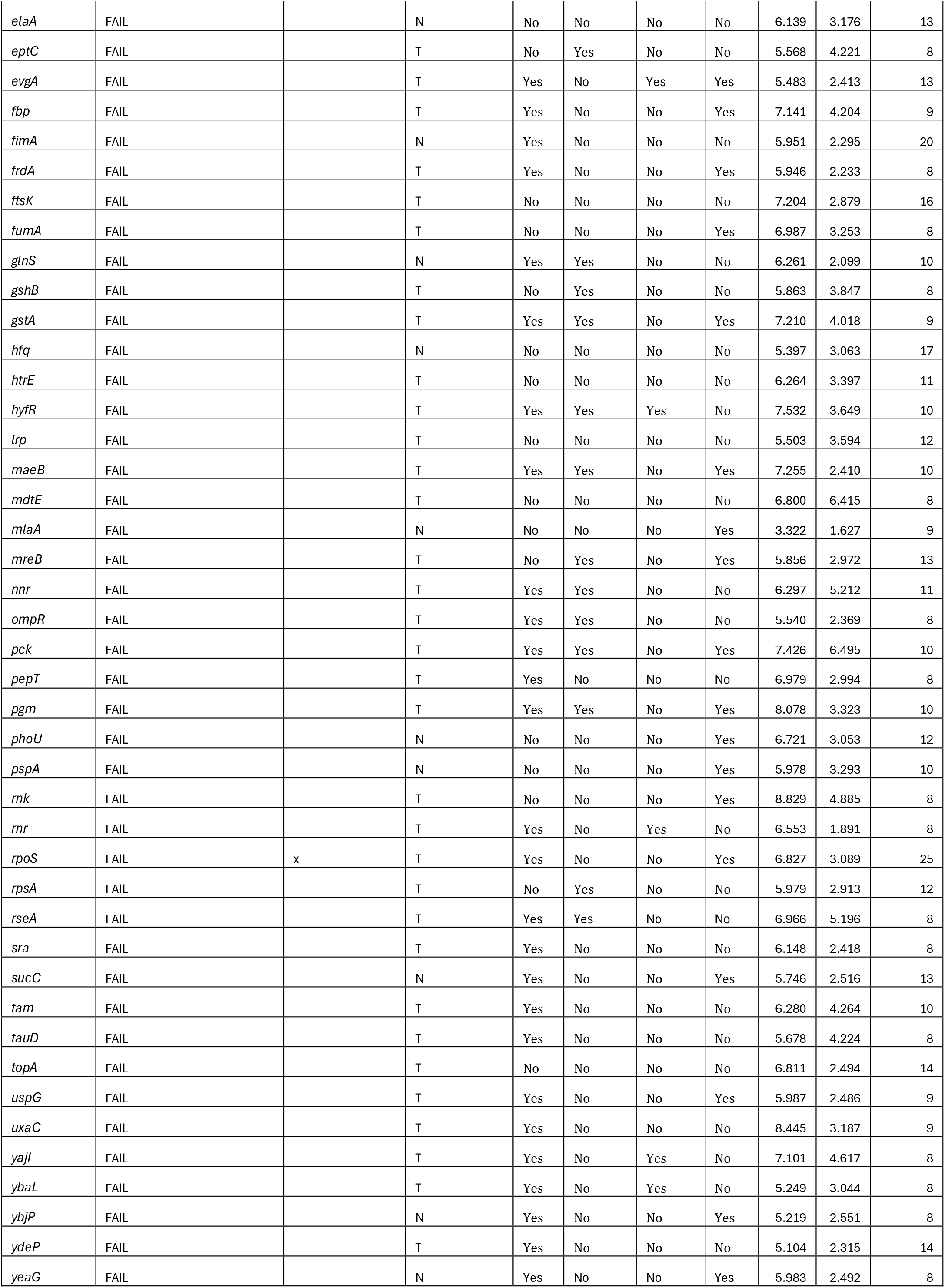
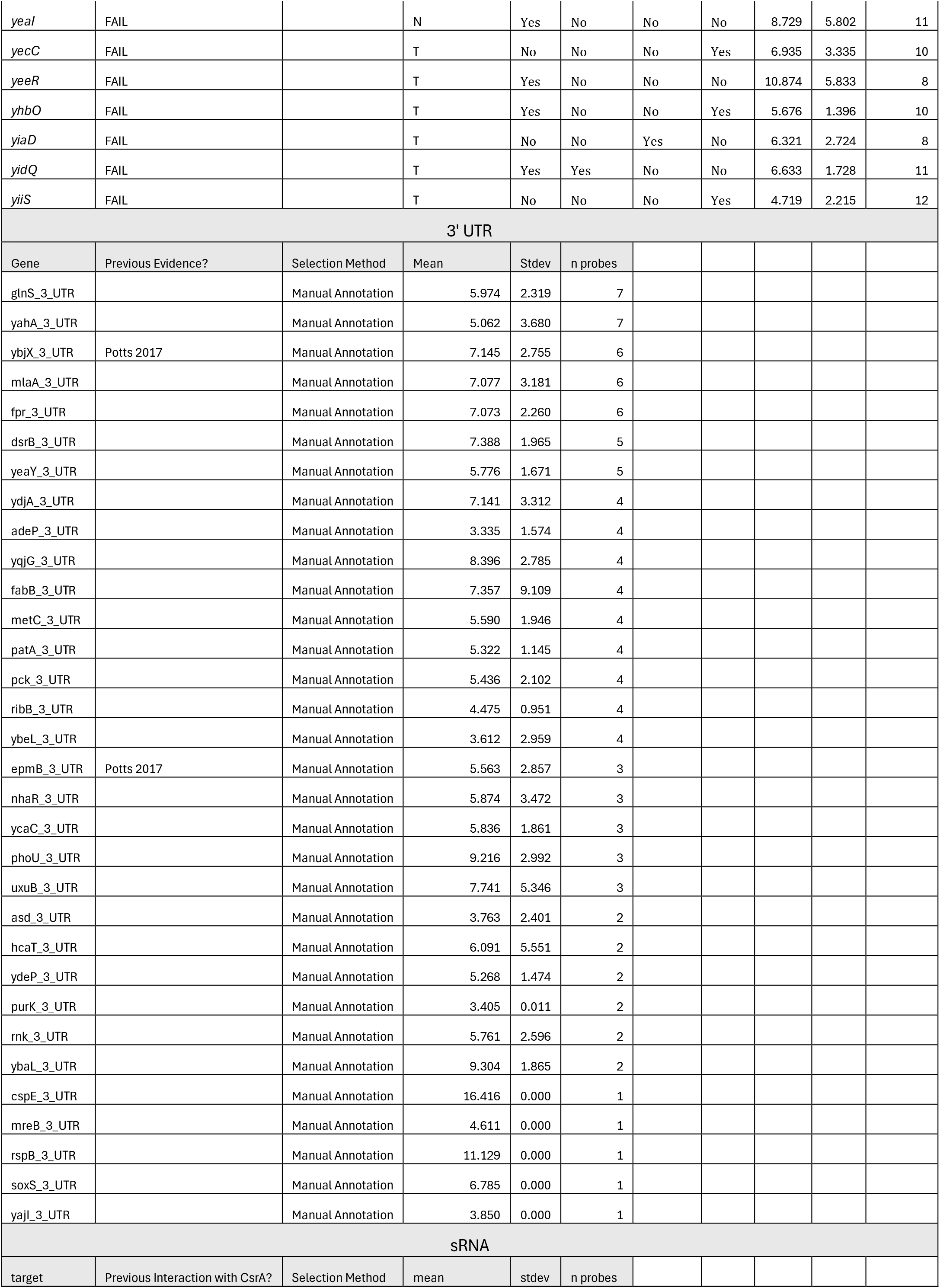
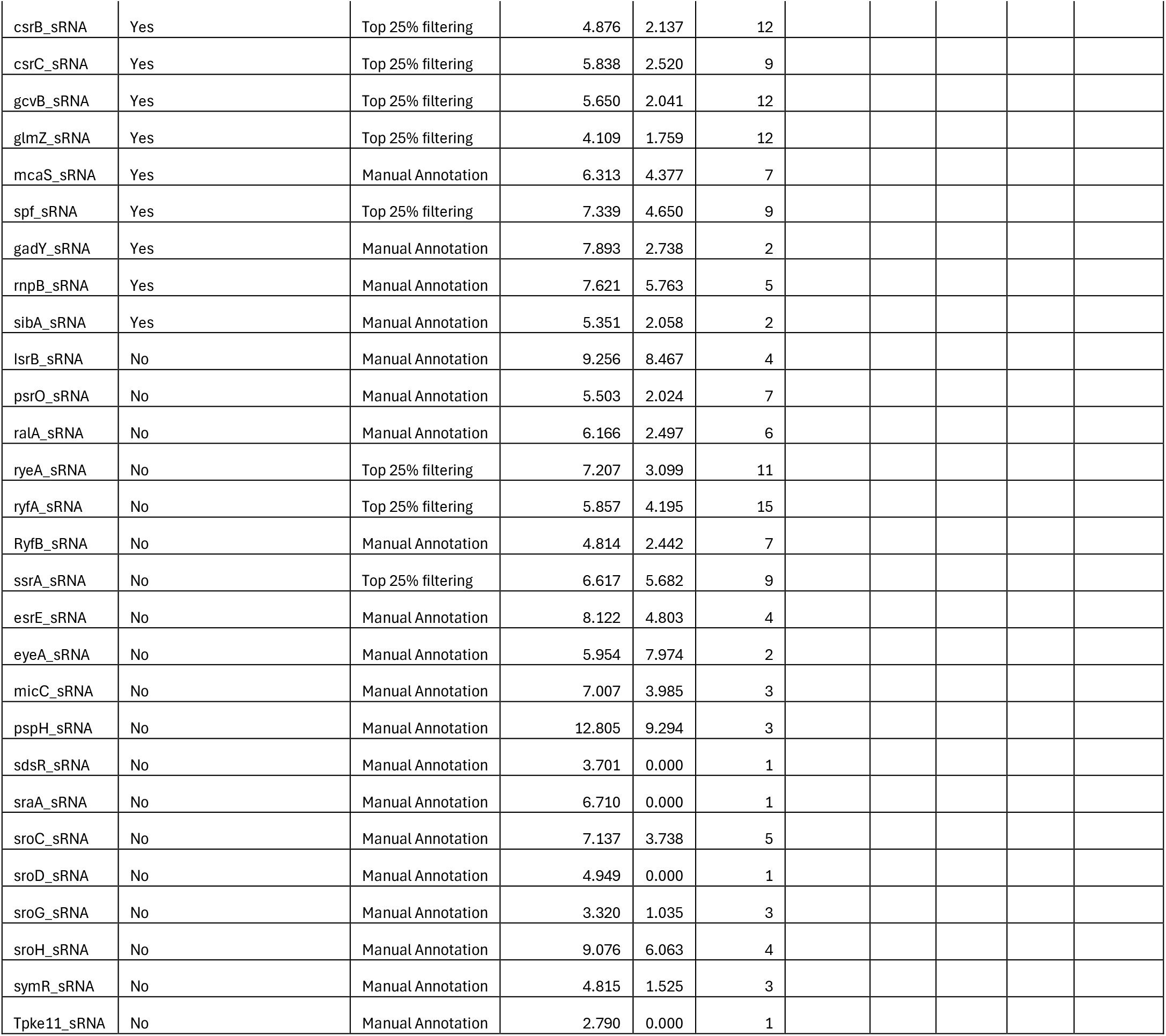
A list of all RNA targets probed in this study that met the following criteria: either (a) the RNA had atleast 8 probes with a significant change in accessibility (p< 0.05) and showed peaks that contained an AGA/GGA, representative of a CsrA binding motif, or (b) RNAs that were manually annotated as potential AGA/GGA peaks due to not meeting the 8 probe criteria because of a small number of probes

| 5' UTR |  |  |  |  |  |  |  |  |  |  |
|---|---|---|---|---|---|---|---|---|---|---|
| Gene | GGA + AU Rich Flanking Pass? | Footprinted | Model Predicted | Potts | Edwards | Kulkarni | Sowa | Mean | Stdev | n probes |
| *acs* | PASS |  | N | Yes | No | No | No | 6.320 | 2.362 | 17 |
| *csrA* | PASS |  | T | No | Yes | Yes | No | 3.688 | 0.905 | 9 |
| *cysJ* | PASS |  | T | No | No | No | Yes | 5.094 | 3.052 | 9 |
| *dcrB* | PASS |  | T | No | No | No | Yes | 6.730 | 2.623 | 10 |
| *dnaK* | PASS |  | T | Yes | No | No | Yes | 6.450 | 4.054 | 9 |
| *dps* | PASS |  | T | Yes | Yes | No | Yes | 8.312 | 4.146 | 8 |
| *fdoG* | PASS |  | T | Yes | Yes | No | No | 6.081 | 3.448 | 8 |
| *glgC* | PASS |  | N | Yes | Yes | Yes | Yes | 5.727 | 2.523 | 18 |
| *glmU* | PASS |  | T | No | Yes | No | No | 7.416 | 3.496 | 13 |
| *glnH* | PASS |  | N | No | No | No | No | 5.662 | 2.248 | 12 |
| *groL* | PASS |  | T | Yes | Yes | No | Yes | 6.531 | 2.870 | 9 |
| *hemD* | PASS |  | T | No | No | No | No | 6.578 | 1.413 | 8 |
| *hflK* | PASS |  | T | No | No | Yes | Yes | 7.384 | 3.871 | 9 |
| *icd* | PASS |  | T | Yes | Yes | No | No | 4.800 | 2.294 | 13 |
| *iraD* | PASS | x | T | Yes | No | No | No | 7.250 | 5.356 | 24 |
| *ltaE* | PASS |  | N | Yes | No | No | Yes | 7.239 | 2.484 | 10 |
| *manX* | PASS |  | N | Yes | No | No | No | 6.431 | 3.058 | 10 |
| *mdtA* | PASS |  | T | No | No | No | No | 5.521 | 1.999 | 10 |
| *moaB* | PASS |  | N | Yes | No | No | No | 6.200 | 3.253 | 9 |
| *nuoB* | PASS |  | N | No | Yes | No | No | 7.629 | 3.822 | 9 |
| *pgaA* | PASS |  | T | Yes | No | Yes | No | 6.111 | 2.091 | 15 |
| *ppk* | PASS |  | T | No | Yes | No | Yes | 7.306 | 2.550 | 8 |
| *proP* | PASS |  | T | Yes | Yes | No | Yes | 6.826 | 3.578 | 13 |
| *relA* | PASS |  | T | Yes | Yes | No | No | 6.191 | 2.726 | 25 |
| *sdiA* | PASS |  | T | Yes | No | No | No | 8.612 | 4.973 | 8 |
| *talA* | PASS |  | N | Yes | No | No | Yes | 8.059 | 4.075 | 8 |
| *tnaA* | PASS |  | T | Yes | Yes | Yes | No | 7.458 | 3.016 | 11 |
| *uidR* | PASS |  | T | No | Yes | No | No | 7.488 | 2.629 | 9 |
| *uxuA* | PASS |  | T | Yes | No | No | No | 7.396 | 4.094 | 9 |
| *yafQ* | PASS |  | T | Yes | Yes | No | No | 5.682 | 2.814 | 8 |
| *yafQ* | PASS |  | T | Yes | Yes | No | No | 5.682 | 2.814 | 8 |
| *ycdT* | PASS |  | T | Yes | No | No | No | 8.404 | 4.694 | 8 |
| *cbpA* | FAIL |  | T | Yes | No | No | Yes | 6.895 | 4.482 | 17 |
| *crp* | FAIL |  | T | No | Yes | No | No | 6.639 | 6.294 | 14 |
| *dgcZ* | FAIL |  | T | Yes | No | Yes | No | 5.890 | 2.224 | 8 |
| *dppA* | FAIL |  | T | Yes | No | No | No | 5.942 | 2.330 | 11 |
| *dsrB* | FAIL |  | T | Yes | No | No | Yes | 6.771 | 2.811 | 11 |
| *elaA* | FAIL |  | N | No | No | No | No | 6.139 | 3.176 | 13 |
| *eptC* | FAIL |  | T | No | Yes | No | No | 5.568 | 4.221 | 8 |
| *evgA* | FAIL |  | T | Yes | No | Yes | Yes | 5.483 | 2.413 | 13 |
| *fbp* | FAIL |  | T | Yes | No | No | Yes | 7.141 | 4.204 | 9 |
| *fimA* | FAIL |  | N | Yes | No | No | No | 5.951 | 2.295 | 20 |
| *frdA* | FAIL |  | T | Yes | No | No | Yes | 5.946 | 2.233 | 8 |
| *ftsK* | FAIL |  | T | No | No | No | No | 7.204 | 2.879 | 16 |
| *fumA* | FAIL |  | T | No | No | No | Yes | 6.987 | 3.253 | 8 |
| *glnS* | FAIL |  | N | Yes | Yes | No | No | 6.261 | 2.099 | 10 |
| *gshB* | FAIL |  | T | No | Yes | No | No | 5.863 | 3.847 | 8 |
| *gstA* | FAIL |  | T | Yes | Yes | No | Yes | 7.210 | 4.018 | 9 |
| *hfq* | FAIL |  | N | No | No | No | No | 5.397 | 3.063 | 17 |
| *htrE* | FAIL |  | T | No | No | No | No | 6.264 | 3.397 | 11 |
| *hyfR* | FAIL |  | T | Yes | Yes | Yes | No | 7.532 | 3.649 | 10 |
| *lrp* | FAIL |  | T | No | No | No | No | 5.503 | 3.594 | 12 |
| *maeB* | FAIL |  | T | Yes | Yes | No | Yes | 7.255 | 2.410 | 10 |
| *mdtE* | FAIL |  | T | No | No | No | No | 6.800 | 6.415 | 8 |
| *mlaA* | FAIL |  | N | No | No | No | Yes | 3.322 | 1.627 | 9 |
| *mreB* | FAIL |  | T | No | Yes | No | Yes | 5.856 | 2.972 | 13 |
| *nnr* | FAIL |  | T | Yes | Yes | No | No | 6.297 | 5.212 | 11 |
| *ompR* | FAIL |  | T | Yes | Yes | No | No | 5.540 | 2.369 | 8 |
| *pck* | FAIL |  | T | Yes | Yes | No | Yes | 7.426 | 6.495 | 10 |
| *pepT* | FAIL |  | T | Yes | No | No | No | 6.979 | 2.994 | 8 |
| *pgm* | FAIL |  | T | Yes | Yes | No | Yes | 8.078 | 3.323 | 10 |
| *phoU* | FAIL |  | N | No | No | No | Yes | 6.721 | 3.053 | 12 |
| *pspA* | FAIL |  | N | No | No | No | Yes | 5.978 | 3.293 | 10 |
| *rnk* | FAIL |  | T | No | No | No | Yes | 8.829 | 4.885 | 8 |
| *rnr* | FAIL |  | T | Yes | No | Yes | No | 6.553 | 1.891 | 8 |
| *rpoS* | FAIL | x | T | Yes | No | No | Yes | 6.827 | 3.089 | 25 |
| *rpsA* | FAIL |  | T | No | Yes | No | No | 5.979 | 2.913 | 12 |
| *rseA* | FAIL |  | T | Yes | Yes | No | No | 6.966 | 5.196 | 8 |
| *sra* | FAIL |  | T | Yes | No | No | No | 6.148 | 2.418 | 8 |
| *sucC* | FAIL |  | N | Yes | No | No | Yes | 5.746 | 2.516 | 13 |
| *tam* | FAIL |  | T | Yes | No | No | No | 6.280 | 4.264 | 10 |
| *tauD* | FAIL |  | T | Yes | No | No | No | 5.678 | 4.224 | 8 |
| *topA* | FAIL |  | T | No | No | No | No | 6.811 | 2.494 | 14 |
| *uspG* | FAIL |  | T | Yes | No | No | Yes | 5.987 | 2.486 | 9 |
| *uxaC* | FAIL |  | T | Yes | No | No | No | 8.445 | 3.187 | 9 |
| *yajI* | FAIL |  | T | Yes | No | Yes | No | 7.101 | 4.617 | 8 |
| *ybaL* | FAIL |  | T | Yes | No | Yes | No | 5.249 | 3.044 | 8 |
| *ybjP* | FAIL |  | N | Yes | No | No | Yes | 5.219 | 2.551 | 8 |
| *ydeP* | FAIL |  | T | Yes | No | No | No | 5.104 | 2.315 | 14 |
| *yeaG* | FAIL |  | N | Yes | No | No | Yes | 5.983 | 2.492 | 8 |
| *yeaI* | FAIL |  | N | Yes | No | No | No | 8.729 | 5.802 | 11 |
| *yecC* | FAIL |  | T | No | No | No | Yes | 6.935 | 3.335 | 10 |
| *yeeR* | FAIL |  | T | Yes | No | No | No | 10.874 | 5.833 | 8 |
| *yhbO* | FAIL |  | T | Yes | No | No | Yes | 5.676 | 1.396 | 10 |
| *yiaD* | FAIL |  | T | No | No | Yes | No | 6.321 | 2.724 | 8 |
| *yidQ* | FAIL |  | T | Yes | Yes | No | No | 6.633 | 1.728 | 11 |
| *yiiS* | FAIL |  | T | No | No | No | Yes | 4.719 | 2.215 | 12 |
| 3' UTR |  |  |  |  |  |  |  |  |  |  |
| Gene | Previous Evidence? | Selection Method | Mean | Stdev | n probes |  |  |  |  |  |
| glnS_3_UTR |  | Manual Annotation | 5.974 | 2.319 | 7 |  |  |  |  |  |
| yahA_3_UTR |  | Manual Annotation | 5.062 | 3.680 | 7 |  |  |  |  |  |
| ybjX_3_UTR | Potts 2017 | Manual Annotation | 7.145 | 2.755 | 6 |  |  |  |  |  |
| mlaA_3_UTR |  | Manual Annotation | 7.077 | 3.181 | 6 |  |  |  |  |  |
| fpr_3_UTR |  | Manual Annotation | 7.073 | 2.260 | 6 |  |  |  |  |  |
| dsrB_3_UTR |  | Manual Annotation | 7.388 | 1.965 | 5 |  |  |  |  |  |
| yeaY_3_UTR |  | Manual Annotation | 5.776 | 1.671 | 5 |  |  |  |  |  |
| ydjA_3_UTR |  | Manual Annotation | 7.141 | 3.312 | 4 |  |  |  |  |  |
| adeP_3_UTR |  | Manual Annotation | 3.335 | 1.574 | 4 |  |  |  |  |  |
| yqjG_3_UTR |  | Manual Annotation | 8.396 | 2.785 | 4 |  |  |  |  |  |
| fabB_3_UTR |  | Manual Annotation | 7.357 | 9.109 | 4 |  |  |  |  |  |
| metC_3_UTR |  | Manual Annotation | 5.590 | 1.946 | 4 |  |  |  |  |  |
| patA_3_UTR |  | Manual Annotation | 5.322 | 1.145 | 4 |  |  |  |  |  |
| pck_3_UTR |  | Manual Annotation | 5.436 | 2.102 | 4 |  |  |  |  |  |
| ribB_3_UTR |  | Manual Annotation | 4.475 | 0.951 | 4 |  |  |  |  |  |
| ybeL_3_UTR |  | Manual Annotation | 3.612 | 2.959 | 4 |  |  |  |  |  |
| epmB_3_UTR | Potts 2017 | Manual Annotation | 5.563 | 2.857 | 3 |  |  |  |  |  |
| nhaR_3_UTR |  | Manual Annotation | 5.874 | 3.472 | 3 |  |  |  |  |  |
| ycaC_3_UTR |  | Manual Annotation | 5.836 | 1.861 | 3 |  |  |  |  |  |
| phoU_3_UTR |  | Manual Annotation | 9.216 | 2.992 | 3 |  |  |  |  |  |
| uxuB_3_UTR |  | Manual Annotation | 7.741 | 5.346 | 3 |  |  |  |  |  |
| asd_3_UTR |  | Manual Annotation | 3.763 | 2.401 | 2 |  |  |  |  |  |
| hcaT_3_UTR |  | Manual Annotation | 6.091 | 5.551 | 2 |  |  |  |  |  |
| ydeP_3_UTR |  | Manual Annotation | 5.268 | 1.474 | 2 |  |  |  |  |  |
| purK_3_UTR |  | Manual Annotation | 3.405 | 0.011 | 2 |  |  |  |  |  |
| rnk_3_UTR |  | Manual Annotation | 5.761 | 2.596 | 2 |  |  |  |  |  |
| ybaL_3_UTR |  | Manual Annotation | 9.304 | 1.865 | 2 |  |  |  |  |  |
| cspE_3_UTR |  | Manual Annotation | 16.416 | 0.000 | 1 |  |  |  |  |  |
| mreB_3_UTR |  | Manual Annotation | 4.611 | 0.000 | 1 |  |  |  |  |  |
| rspB_3_UTR |  | Manual Annotation | 11.129 | 0.000 | 1 |  |  |  |  |  |
| soxS_3_UTR |  | Manual Annotation | 6.785 | 0.000 | 1 |  |  |  |  |  |
| yajI_3_UTR |  | Manual Annotation | 3.850 | 0.000 | 1 |  |  |  |  |  |
| sRNA |  |  |  |  |  |  |  |  |  |  |
| target | Previous Interaction with CsrA? | Selection Method | mean | stdev | n probes |  |  |  |  |  |
| csrB_sRNA | Yes | Top 25% filtering | 4.876 | 2.137 | 12 |  |  |  |  |  |
| csrC_sRNA | Yes | Top 25% filtering | 5.838 | 2.520 | 9 |  |  |  |  |  |
| gcvB_sRNA | Yes | Top 25% filtering | 5.650 | 2.041 | 12 |  |  |  |  |  |
| glmZ_sRNA | Yes | Top 25% filtering | 4.109 | 1.759 | 12 |  |  |  |  |  |
| mcaS_sRNA | Yes | Manual Annotation | 6.313 | 4.377 | 7 |  |  |  |  |  |
| spf_sRNA | Yes | Top 25% filtering | 7.339 | 4.650 | 9 |  |  |  |  |  |
| gadY_sRNA | Yes | Manual Annotation | 7.893 | 2.738 | 2 |  |  |  |  |  |
| rnpB_sRNA | Yes | Manual Annotation | 7.621 | 5.763 | 5 |  |  |  |  |  |
| sibA_sRNA | Yes | Manual Annotation | 5.351 | 2.058 | 2 |  |  |  |  |  |
| IsrB_sRNA | No | Manual Annotation | 9.256 | 8.467 | 4 |  |  |  |  |  |
| psrO_sRNA | No | Manual Annotation | 5.503 | 2.024 | 7 |  |  |  |  |  |
| ralA_sRNA | No | Manual Annotation | 6.166 | 2.497 | 6 |  |  |  |  |  |
| ryeA_sRNA | No | Top 25% filtering | 7.207 | 3.099 | 11 |  |  |  |  |  |
| ryfA_sRNA | No | Top 25% filtering | 5.857 | 4.195 | 15 |  |  |  |  |  |
| RyfB_sRNA | No | Manual Annotation | 4.814 | 2.442 | 7 |  |  |  |  |  |
| ssrA_sRNA | No | Top 25% filtering | 6.617 | 5.682 | 9 |  |  |  |  |  |
| esrE_sRNA | No | Manual Annotation | 8.122 | 4.803 | 4 |  |  |  |  |  |
| eyeA_sRNA | No | Manual Annotation | 5.954 | 7.974 | 2 |  |  |  |  |  |
| micC_sRNA | No | Manual Annotation | 7.007 | 3.985 | 3 |  |  |  |  |  |
| pspH_sRNA | No | Manual Annotation | 12.805 | 9.294 | 3 |  |  |  |  |  |
| sdsR_sRNA | No | Manual Annotation | 3.701 | 0.000 | 1 |  |  |  |  |  |
| sraA_sRNA | No | Manual Annotation | 6.710 | 0.000 | 1 |  |  |  |  |  |
| sroC_sRNA | No | Manual Annotation | 7.137 | 3.738 | 5 |  |  |  |  |  |
| sroD_sRNA | No | Manual Annotation | 4.949 | 0.000 | 1 |  |  |  |  |  |
| sroG_sRNA | No | Manual Annotation | 3.320 | 1.035 | 3 |  |  |  |  |  |
| sroH_sRNA | No | Manual Annotation | 9.076 | 6.063 | 4 |  |  |  |  |  |
| symR_sRNA | No | Manual Annotation | 4.815 | 1.525 | 3 |  |  |  |  |  |
| Tpke11_sRNA | No | Manual Annotation | 2.790 | 0.000 | 1 |  |  |  |  |  |

Given our observation that the positive control maps appeared to contain AU-rich sequence around GGA and AGA motifs in alignment with previous literature^36^, we explored whether requiring at least 60% AU content in the 6nt flanking the motif on either side improved specificity (**Table S2**). This method reduced our 87 nominated targets to a total of 34 with statistically significant but modest separation relative to the permutation null hypothesis (p = 0.029, FDR = 77%). It retained five of the eight well-established positive controls (*pgaA, csrA, iraD, glgC* and *sdiA)* but lost *hfq* and other well-characterized targets, including *mdtE* among later reporter-tested 5’UTRs and CsrB (the strongest binder of CsrA) among the sRNAs, retaining only CsrC and SsrA. Of the 34 RNAs, five overlapped previously identified CLIP-seq peaks collected under similar media conditions in a different growth phase^6^, including *glgC* and *pgaA*, while 29 did not overlap. For comparison, this same dataset identified 11 of our 87 nominated targets. We therefore keep the original screen as the primary annotation (**Table 3**), and report AU-flank regions as an additional column to help rank candidates and leave a more specific binding-context filter for future work. This analysis also confirmed that indeed the feature that best separated the verified positive controls from other targets was the number of GGA motifs in the 5’UTR. As such, the value of accessibility maps lies in showcasing where within an RNA CsrA-dependent changes occur and in which direction, in a specific cellular context. In the case of CsrA, because its recognition motif is common and the library may consist largely of true CsrA targets, we do not expect any filter based on motif and fold change to be strong.

Among the 87 candidate target 5’UTRs (**Table 3**), 77 had been previously identified experimentally by at least one of CLIP-seq^6^, synergistic combinations of various omics/genetics assays^8^, sequence motif discovery^21^, or pull-down assays of epitope-tagged CsrA^37^ (**Figure S6, Table S3**). Because the library was assembled largely from these same datasets (Methods), this overlap shows consistency with prior work but is not an independent estimate of accuracy. Specifically, we found 1 5’UTR with exclusively prior computational evidence (*yiaD*) and 66 with some prior experimental evidence, and 10 5’ UTRs with both computational and experimental evidence. Among these genes (**Table 3**), there are several genes involved in global stress responses, including the known target *hfq,* and putative targets *rpoS* and *ppk*. At least 10 5’UTRs were not reported in any of the four prior datasets (though the *hfq* RNA has been shown to be regulated in low throughput); 7 of these 10 had been predicted by a computational model from our group^38^ (**Table S3),** which provides indirect support but not independent experimental validation.

To assess whether accessibility-prioritized candidates show CsrA-dependent regulation, we tested a subset of mRNA 5’UTRs using 5’UTR-GFP translational fusion assays (**Figure 2A**). We selected six mRNA 5’UTRs with varying levels of prior evidence of CsrA interaction^6,8,21,37^ (*rpsA, uxuA, mdtE, ompR, ribB, ppk*, probe maps of these targets are displayed in **Figure S7**). Five of these (*uxuA, mdtE, ompR, rpsA, ppk)* met the criteria for inclusion in **Table 3**, whereas *ribB* had fewer than eight significant probes (7 probes; **Table 2**) and was included on the basis of prior evidence and its role in carbon metabolism; we therefore do not count it as a screen-derived candidate.

**Figure 2.**
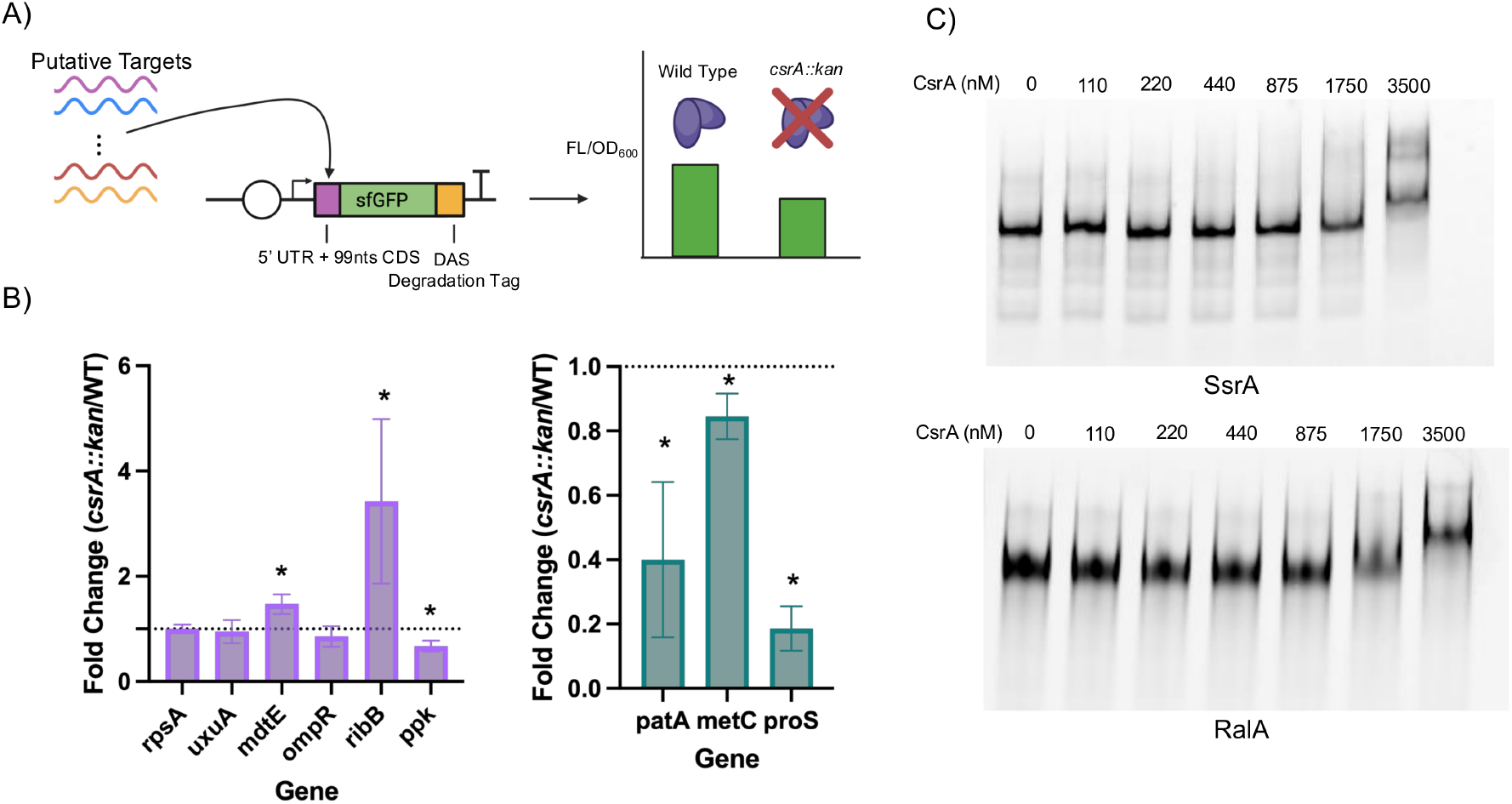
Reporter and in vitro binding assays for candidate CsrA targets. **(A)** 5’ UTR GFP fusion assays of putative CsrA targets allows for measurement of CsrA-dependent effects on the translation of 5’ UTRs. **(B)** Fluorescence data of ten candidate 5’UTRs (left: seven candidates drawn from the primary list or prior evidence; right: three candidate activated 5’UTRs), presented as the fold change of normalized GFP fluorescence between the *csrA::kan* strain and the WT strain. A * indicates a p value less than 0.05 in an unadjusted 2-tailed heteroscedastic t-test (n=3 biological replicates; error bars, SD). **(C)** Electrophoretic mobility shift assays CsrA-dependent mobility shifts of SsrA (tmRNA) and RalA small RNAs at the CsrA concentrations indicated (nM).

In these plasmid-based 5’UTR-GFP translational fusion assays (**Figure 2B**), altering the CsrA activity levels (via WT and *csrA::kan* strains) leads to changes in GFP fluorescence if these UTRs are regulated by CsrA directly or indirectly. As shown in **Figure 2B**, we observed significant (n= 3 biological replicates, *p*<0.05, two tailed independent t-test) repression by CsrA for 2 of the 6 putative genes *(mdtE, ribB*) indicating CsrA-dependent regulation of these 5’UTRs (the effect for *mdtE* was modest, approximately 1.5-fold). Three of the six (*rpsA, uxuA, and ompR)* were not significantly regulated; this may be due to condition-specific CsrA-regulatory interactions, binding events without translational consequences, or accessibility-responsive regions requiring additional regulatory factors that may be condition-dependent. Interestingly, the *ppk* reporter was lower in *csrA::kan* cells (approximately 0.65-fold), consistent with activation by CsrA, a rarely seen regulatory outcome. Collectively, three of six reporters (*mdtE, ribB and ppk*) responded, including two of the five (*mdtE* and *ppk*) drawn from the primary candidate list (**Table 3)**; *ribB* was added on prior evidence of CsrA regulation^8^ and due to its role as an essential gene in carbon metabolism^39^. Although with a low model validation rate, accessibility profiling can enrich for CsrA-responsive 5’UTRs, however a difference in the fold-change of accessibility values between wild type and *csrA::kan* does not by itself confirm direct regulation.

### Decreased accessibility in *csrA::kan* nominates candidate CsrA-activated 5’UTRs

Activation of RNA targets by CsrA is less frequently observed than repression within the literature: to date, only the following genes have been experimentally characterized as activators: *flhD*, *moaA*, *uxuB,* and *ymdA* ^23–25,31^. Considering the best characterized mechanism of CsrA activation: the *ymdA* 5’ UTR, where CsrA binding opens the ribosome binding site (RBS)^24^, we identified candidate 5’UTRs that showed decreases in accessibility in the *csrA::kan* strain near the start codon (-100 to +100 nts) as potential activators. Upon screening all 3,024 probes that showed any significant change in accessibility between WT and *csrA::kan,* we observed that only 46 of them displayed an accessibility decrease in the *csrA::kan* strain **(Figure S2).** Most significant probes instead showed increased accessibility in *csrA::kan* (**Figure S2**), which is why all regions were normalized within each target RNA. We hypothesized that these regions of accessibility change could be signatures of CsrA-dependent opening of functional regions. We identified 22 candidate 5’UTRs and compared them to published CsrA-dependent proteomics data^8^ to identify transcripts that showed potential activation by CsrA (**Table 4**). Among the 22 candidates, 40% showed protein-level activation (compared to 20% (63 of 306 genes) of the entire 5’UTR library, when compared to proteomics data^8^). Specifically, genes that showed a significant increase in protein expression from WT to *csrA::kan* or *ΔcsrBC* to WT (both indicative of an “increase” in CsrA activity) were flagged as possible activators. We found 9 such targets and flagged them as potential candidates for activation by CsrA (“Potential Activation,” **Table 4**). We selected three of the nine candidates (*patA, metC, proS*) and tested them using the same 5’ UTR-GFP assay. All three showed lower fluorescence in *csrA::kan* (fold change <1, p<0.05, two-tailed t-test; **Figure 2B**), consistent with activation by CsrA, although the effect for *metC* was modest (approximately 1.17-fold in WT over *csrA::kan*). Two other **Table 4** candidates have reporter data: *rpsA* showed no CsrA-dependent change (**Figure 2B**), and *crp,* whose reported fluorescence was higher in *csrA::kan* (**Figure 3C**), consistent with repression rather than activation. Three of the five candidates with reporter data therefore behaved as CsrA-activated. Of note, *patA* and *metC* have at least eight significant probes but no motif-containing peak (**Table 5**), and *proS* had seven significant probes (**Table 2**), so none of the three met the criteria for **Table 3**. Finally, of the three previously characterized CsrA-activated transcripts represented in our library (*flhD, moaA*, and *uxuB*; *ymdA* was not probed), only *uxuB* carried the decreased-accessibility signature (**Table 4**), so this signature identified at most a subset of CsrA-activated 5’UTRs.

**Figure 3.**
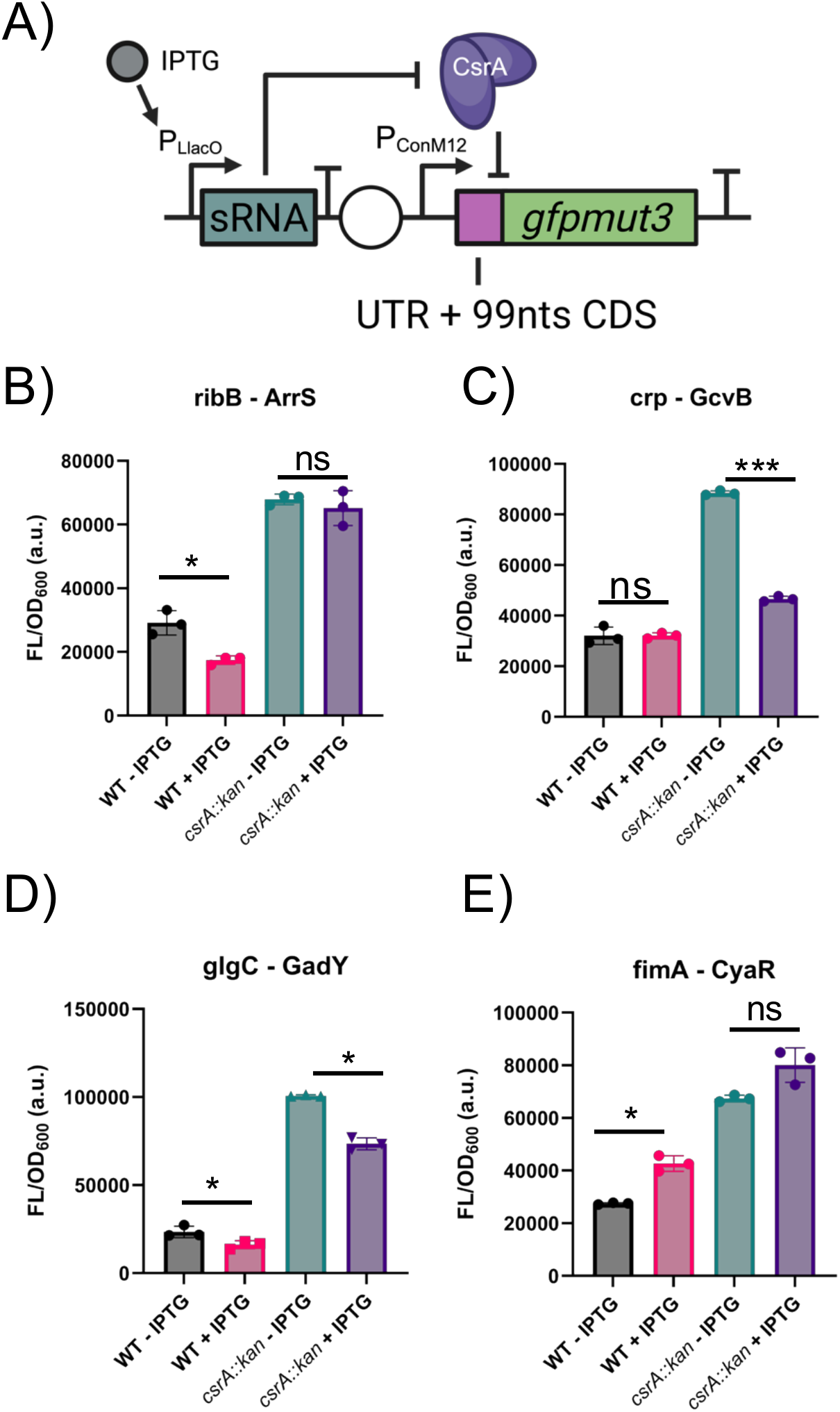
Reporter assays suggest CsrA-dependent modulation of two sRNA–mRNA pairs. **(A)** Design of experiment to test the effect of CsrA on sRNA-mRNA interactions **(B)** the *ribB* transcript is repressed by the ArrS RNA only in the presence of CsrA. **(C)** The *crp* transcript is repressed by the GcvB sRNA only in the absence of CsrA **(D)** the *glgC* transcript is repressed by the GadY sRNA, however the effect is CsrA independent **(E)** the *fimA* transcript is activated by the CyaR transcript, however the effect is CsrA independent (n = 3 biological replicates; error bars, SD; * p < 0.05, *** p < 0.001)

**Table 4.** A list of all RNA targets probed in this study that met the following criteria: 5’ UTRs that showed a region of decreased accessibility near their start codon, suggesting some level of structural rearrangement upon CsrA binding. Proteomics taken from (8).

| Gene | Identified Probe | Fold change | pval | Proteomic Changes |
|---|---|---|---|---|
| *glmU* | glmS_glmU_UTR_probe_9 | 0.567681569 | 0.017233829 | NA |
| *uxuB* | uxuA_uxuB_UTR_probe_15 | 0.571418264 | 0.02614935 | Heterogeneous |
| *adeP* | adeP_5_UTR_probe_6 | 0.623370372 | 7.86174E-06 | NA |
| *ahpF* | ahpC_ahpF_UTR_probe_8 | 0.418109099 | 0.000702315 | NA |
| *icd* | icd_5_UTR_probe_19 | 0.400092527 | 0.006969059 | NA |
| *tnaB* | tnaA_tnaB_UTR_probe_5 | 0.755021886 | 0.036620126 | NA |
| *yhfK* | crp_yhfK_UTR_probe_5 | 0.626113833 | 0.032390197 | NA |
| *yiaD* | yiaD_5_UTR_probe_15 | 0.556526588 | 0.028069235 | NA |
| *crp* | crp_5_UTR_probe_6 | 0.76160071 | 0.017631313 | Potential Activation |
| *entF* | ybdZ_entF_UTR_probe_20 | 0.387302184 | 0.014437191 | Potential Activation |
| *guaB* | guaA_guaB_UTR_probe_11 | 0.451707151 | 0.017262231 | Potential Activation |
| *metC* | metC_5_UTR_probe_19 | 0.348177737 | 0.020385313 | Potential Activation |
| *patA* | patA_5_UTR_probe_16 | 0.588851103 | 0.009827219 | Potential Activation |
| *proS* | proS_5_UTR_probe_11 | 0.692067961 | 0.005090751 | Potential Activation |
| *relA* | relA_5_UTR_probe_7 | 0.556448927 | 0.005118464 | Potential Activation |
| *rpsA* | cmk_rpsA_UTR_probe_7 | 0.57282409 | 0.014043354 | Potential Activation |
| *yjbD* | yjbD_5_UTR_probe_12 | 0.681510784 | 0.046862746 | Potential Activation |
| *cysJ* | cysJ_5_UTR_probe_16 | 0.760710746 | 0.023203384 | Repression |
| *gstA* | gstA_5_UTR_probe_16 | 0.670779756 | 0.029796169 | Repression |
| *hdeA* | hdeB_hdeA_UTR_probe_12 | 0.499679216 | 0.012530606 | Repression |
| *pck* | pck_5_UTR_probe_6 | 0.6890706 | 0.008338263 | Repression |
| *talA* | talA_5_UTR_probe_16 | 0.422245648 | 0.030649467 | Repression |

**Table 5.** A list of all RNA targets probed in this study that met the following criteria: the RNA had at least 8 probes with a significant (p <0.05) change in accessibility but were not identified to have a peak containing a GGA or AGA.

| 5' UTR |  |  |  |  |  |  |  |
|---|---|---|---|---|---|---|---|
| Gene | Potts | Edwards | Kulkarni | Sowa | Mean | Stdev | n probes |
| *ahpF* | No | No | No | No | 6.350286552 | 3.421494915 | 25 |
| *eno* | Yes | No | No | Yes | 7.761488512 | 4.466336766 | 24 |
| *arcA* | No | No | No | No | 6.015519423 | 2.877024708 | 22 |
| *kdsA* | No | Yes | No | Yes | 6.516563677 | 5.179563512 | 22 |
| *acnA* | Yes | No | No | Yes | 8.125281923 | 4.056343318 | 21 |
| *pflB* | No | Yes | No | No | 6.425746513 | 3.79647867 | 17 |
| *hchA* | No | No | No | Yes | 7.746242894 | 4.138432707 | 15 |
| *gabD* | Yes | Yes | No | Yes | 6.923086036 | 5.507468212 | 14 |
| *glpR* | NA | NA | NA | NA | 6.098307924 | 2.728026269 | 14 |
| *ytfQ* | Yes | No | No | Yes | 8.868028192 | 6.90745557 | 14 |
| *nhaA* | No | No | No | No | 7.871156825 | 4.042868665 | 12 |
| *glnH* | No | No | No | No | 5.661521323 | 2.24823144 | 12 |
| *fabI* | No | No | No | No | 7.25451022 | 4.198409575 | 12 |
| *cspE* | No | Yes | No | No | 7.415190637 | 3.159072678 | 11 |
| *tktB* | Yes | No | No | No | 7.559335903 | 4.361122669 | 11 |
| *metC* | No | No | No | No | 8.317718969 | 6.238500905 | 11 |
| *yliE* | No | Yes | No | No | 6.048481757 | 1.830488867 | 10 |
| *fabB* | Yes | No | No | No | 6.741269976 | 3.384327344 | 10 |
| *entF* | Yes | Yes | No | Yes | 4.632976512 | 2.723279998 | 10 |
| *scpA* | Yes | Yes | No | No | 10.84206819 | 16.13037283 | 10 |
| *yqjG* | Yes | No | Yes | No | 7.742443044 | 2.947472021 | 10 |
| *ltaE* | Yes | No | No | Yes | 7.238955019 | 2.484240384 | 10 |
| *barA* | No | No | No | No | 10.68699703 | 10.95947287 | 10 |
| *patA* | Yes | No | No | Yes | 4.614761778 | 2.550752589 | 9 |
| *pnp* | No | Yes | No | No | 5.695238431 | 2.553983593 | 9 |
| *hdeA* | Yes | No | No | Yes | 6.892710555 | 4.5216778 | 9 |
| *hdeA* | Yes | No | No | Yes | 6.892710555 | 4.5216778 | 9 |
| *uxaC* | Yes | No | No | No | 8.444553573 | 3.187329237 | 9 |
| *adeQ* | No | Yes | No | No | 5.217561304 | 1.433011296 | 8 |
| *rpoE* | No | Yes | No | No | 6.795885709 | 3.603119968 | 8 |
| *moaA* | Yes | Yes | No | No | 5.267185287 | 1.913252231 | 8 |
| *dctR* | NA | NA | NA | NA | 8.683190517 | 5.646035293 | 8 |
| *entC* | Yes | No | Yes | Yes | 8.149548493 | 4.712113952 | 8 |
| *hemG* | Yes | No | No | Yes | 5.655221166 | 1.942772023 | 8 |
| *dacC* | Yes | No | No | Yes | 5.081763328 | 3.161539222 | 8 |
| *deoD* | Yes | No | Yes | No | 6.272107157 | 5.738825413 | 8 |
| *fhuA* | Yes | No | No | No | 11.24862789 | 13.62715542 | 8 |
| *yihX* | Yes | No | No | Yes | 5.023676108 | 1.750768735 | 8 |
| *hipB* | No | No | No | No | 5.64516288 | 2.408695322 | 8 |
| *ugpB* | Yes | Yes | No | Yes | 5.576284219 | 2.731877979 | 8 |
| *uvrY* | Yes | No | No | No | 6.677705561 | 4.385738726 | 8 |
| *ddlA* | Yes | No | No | No | 5.695593953 | 3.199504586 | 8 |
| *flhD* | Yes | Yes | No | No | 5.466482624 | 2.486847856 | 17 |
| *rodZ* | No | No | No | No | 6.386919963 | 3.508247779 | 13 |
| 3' UTR |  |  |  |  |  |  |  |
| Gene | Mean | Stdev | n probes |  |  |  |  |
| glpD_3_UTR | 8.249082978 | 6.038111797 | 9 |  |  |  |  |
| sRNA |  |  |  |  |  |  |  |
| Gene | Mean | Stdev | n probes |  |  |  |  |
| Tpke70_sRNA | 7.256331523 | 2.601012167 | 15 |  |  |  |  |
| RybA_sRNA | 6.321409834 | 3.257199334 | 13 |  |  |  |  |
| sgrS_sRNA | 4.736306528 | 1.576290867 | 10 |  |  |  |  |
| istR_sRNA | 7.59987137 | 3.391723635 | 8 |  |  |  |  |
| psrD_sRNA | 8.713746265 | 7.735180426 | 8 |  |  |  |  |

### Accessibility profiling nominates sRNA partners of CsrA, including SsrA and RalA, which bind CsrA in vitro

Beyond mRNA targets, RNA accessibility profiling nominated a number of small RNAs as candidate CsrA partners. Among the group of 28 sRNAs identified by filtering RNA accessibility probing data (**Table 3**), we identified 9 previously characterized CsrA target sRNAs, including CsrB/C^11^, Spot 42^12,26^, and GadY^6^, as well as 19 sRNAs not previously reported to bind CsrA in the literature (**Table 3**). Only three of these 19 were called by the automated filter; the other 16 were nominated by manual annotation, 12 of the 19 because of fewer than eight significant probes and four on a single significant probe (**Table 3**). From the 19 identified putative sRNA targets, we selected two sRNA candidates that had potentially interesting phenotypic consequences: SsrA, which is a key regulator of trans-translation^40^, and RalA, a member of a toxin-antitoxin system^41^. We performed *in vitro* binding assays (EMSAs) between CsrA and each sRNA, and observed a CsrA-dependent mobility shift for both, at the highest CsrA concentrations tested (**Figure 2C**). Both of the tested sRNAs had a K_d_ < 3.5 uM CsrA, which is in line with previous literature on non-canonical CsrA targets^12^.

Overall, accessibility profiling nominated 19 previously unreported candidate sRNA partners of CsrA. Two of these targets, SsrA and RalA, bind CsrA in vitro, which suggests that the intersection between CsrA and sRNA networks may be broader than currently appreciated. Of note, whether these interactions occur in vivo remains to be tested.

### Accessibility profiles nominate sRNA-mRNA pairs whose regulation depends on CsrA

CsrA has previously been shown to impact sRNA-mRNA interactions through direct binding^12^, and we hypothesized that RNA accessibility data could identify regions of known sRNA-mRNA interactions that might be impacted by CsrA. Within each 5’ UTR found to be a putative CsrA target (**Table 3**), sRNA-mRNA recognition regions were extracted from published data containing RNA-RNA pairs bound within the Hfq protein (Hfq-CLASH^42^), which resulted in a total of 80 identified mRNA-sRNA interactions (**Table 6**). Guided by our RNA-accessibility data and these previously observed RNA-RNA interactions^42^, we selected four candidate pairs. Using GFP-fusion translational assays with IPTG inducible sRNA expression (**Figure 3A**), we observed CsrA-dependent differences in the effect of sRNA induction for two pairs, ArrS-*ribB* and GcvB-*crp*. Both pairs were previously identified by Hfq-CLASH (though not validated through reporter assays), so what is new is their apparent dependence on CsrA and not the pairs themselves. Overexpression of ArrS repressed expression of *ribB-GFP* in the presence but not in the absence of CsrA (WT strain vs. *csrA::kan* strain, **Figure 3B**), suggesting that CsrA may affect this sRNA-mRNA interaction. The accessibility data shows a decrease in the ArrS sRNA at the point of interaction between *ribB* and ArrS in the *csrA::kan* strain (Probe 2, **Figure S8**). This could suggest that CsrA alters the availability of this region of ArrS-*ribB* interactions. We observed repression of the *crp* mRNA by overexpression of the GcvB sRNA in the absence of CsrA but not in its presence (**Figure 3C**), indicating that CsrA may disrupt the GcvB-*crp* interaction. The other two tested pairs, *glgC*-GadY and *fimA*-CyaR, showed sRNA-dependent repression/activation, respectively, but that sRNA-dependent regulation did not appear to be strongly affected by CsrA (**Figure 3D,E)**. Overall, two of four tested pairs showed CsrA-dependent differences, suggesting that CsrA may shape a larger subset of sRNA-mRNA interactions relative to what has been reported thus far, and a third showed a mild effect. Although accessibility data by itself do not establish a mechanism, these data helped nominate these sRNA-mRNA pairs that show CsrA dependency.

**Table 6.** A list of the putative sRNA-mRNA pairs derived from (42) for genes identified as putative CsrA targets in Table 3.

| Gene | partner_ncRNA |
|---|---|
| *acs* | arrS |
| *acs* | cyaR |
| *acs* | gcvB |
| *acs* | omrB |
| *cbpA* | gcvB |
| *cbpA* | chiX |
| *crp* | gcvB |
| *crp* | arcZ |
| *cysJ* | cyaR |
| *dcrB* | spf |
| *dnaK* | cyaR |
| *dnaK* | arcZ |
| *dppA* | arrS |
| *dppA* | chiX |
| *dppA* | gadY |
| *dppA* | gcvB |
| *dppA* | omrB |
| *dppA* | ryeA |
| *dppA* | spf |
| *dppA* | arcZ |
| *dps* | spf |
| *fdoG* | arrS |
| *fdoG* | chiX |
| *fdoG* | csrB |
| *fdoG* | cyaR |
| *fdoG* | ffs |
| *fdoG* | gcvB |
| *fdoG* | rdlC |
| *fdoG* | ryeA |
| *fdoG* | spf |
| *fimA* | arrS |
| *fimA* | gcvB |
| *fimA* | omrA |
| *ftsK* | spf |
| *ftsK* | chiX |
| *fumA* | arcZ |
| *glgC* | gadY |
| *glmU* | glmZ |
| *hemD* | sdsR |
| *hflK* | arcZ |
| *hfq* | spf |
| *icd* | gcvB |
| *icd* | spf |
| *icd* | arcZ |
| *lrp* | cyaR |
| *lrp* | gcvB |
| *lrp* | ryeA |
| *lrp* | spf |
| *maeB* | gcvB |
| *mreB* | omrB |
| *mreB* | csrB |
| *nnr* | gcvB |
| *nuoB* | ryeA |
| *pck* | cyaR |
| *pck* | gcvB |
| *ppc* | spf |
| *proP* | cyaR |
| *ribB* | arrS |
| *rnr* | cyaR |
| *rodZ* | gcvB |
| *rodZ* | dsrA |
| *rpoS* | spf |
| *sucC* | mgrR |
| *sucC* | isrC |
| *tam* | arrS |
| *tnaA* | arcZ |
| *tnaA* | arrS |
| *tnaA* | cyaR |
| *tnaA* | gadY |
| *tnaA* | gcvB |
| *tnaA* | omrB |
| *tnaA* | rybB |
| *tnaA* | spf |
| *tnaA* | sroH |
| *tnaA* | glmZ |
| *topA* | cyaR |
| *uspG* | gcvB |
| *ybaL* | cyaR |
| *ycdT* | gcvB |
| *yiaD* | spf |

### CsrA-dependent expression of *ppk* depends on specific sequence elements in the 5’UTR and coding region

We first asked whether CsrA associates with the *ppk* transcript *in vivo*, utilizing a previously reported three-component fluorescence complementation system (TRI-FC)^31,43^ (**Figure 4A**). Together, this three-component system produces a yellow fluorescent protein (YFP) fluorescent signal if the CsrA protein and the *ppk* mRNA are in proximity due to complementation of the split YFP (**Figure 4A**). Upon deploying this assay, we observed a modest but significant increase (n = 5, FC = 1.2, p = 0.016, One-way ANOVA with Dunnett t-test) in YFP fluorescence complementation, consistent with an in vivo CsrA-*ppk* interaction, relative to the established *fecA* negative control^31^ (**Figure 4A**).

**Figure 4.**
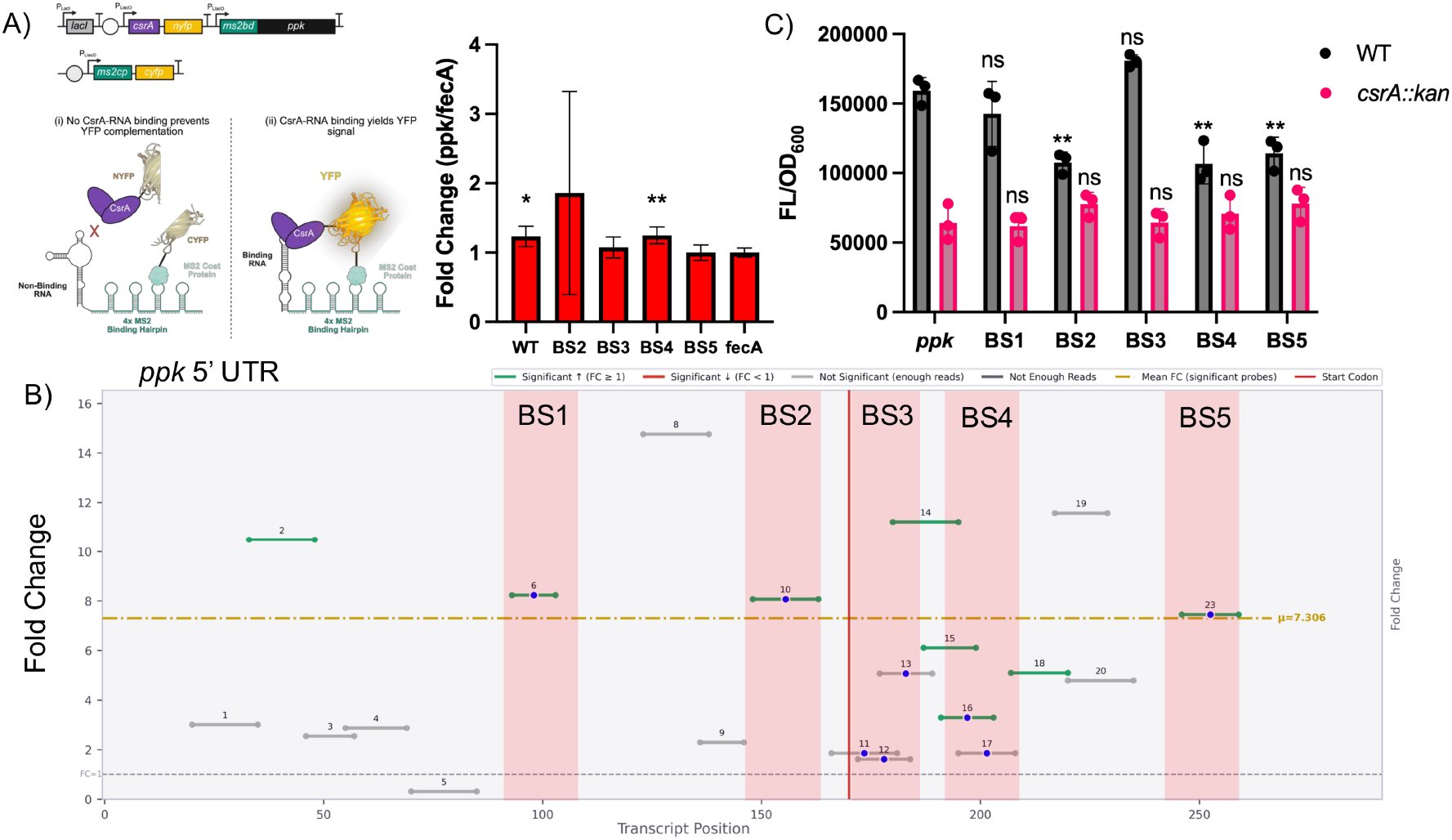
CsrA-dependent expression of *ppk* depends on specific sequence elements in its 5’UTR and coding region. **(A)** Diagram of the two-plasmid Tri-molecular fluorescence complementation (TRI-FC) system for detecting *in vivo* protein-RNA interactions. Adapted from Rojano-Nisimura & Simmons *et al* 2025, *ACS Synthetic Biology* (31), with permission. TRI-FC data (n = 5) comparing the *ppk* transcript and the BS3, BS4 and BS5 mutants to a *fecA* negative control (BS1 not tested). Multiple comparisons performed with the Dunnett test after one-way ANOVA. *p < 0.05, ** p<0.01 **(B)** Accessibility map of the *ppk* 5’ UTR: Each horizontal bar in the map of the *ppk* transcript represents a single asRNA probe. The position on the x-axis signifies the location within the transcript, and the position on the y-axis represents the magnitude of the fold change between the *csrA:kan* and WT strains. Probes containing a blue dot are those that are antisense to a region containing an AGA or GGA motif, signifying a potential CsrA binding site. The color of each bar represents the statistical significance of the probe between the two strains, with green/red bars being statistically significant, and gray bars not significant. The vertical red bar signifies the start codon. The horizontal yellow bar indicates the mean accessibility change of the transcript. Transparent red regions represent putative CsrA-*ppk* binding sites. Binding sites 1, 2, and 5 were predicted through RNA accessibility, and Binding sites 3, 4, and 5 were predicted computationally. **(C)** Fluorescence data of wild type and various putative CsrA binding site mutations of the *ppk* 5’ UTR fused to GFP. Multiple comparisons performed with the Dunnett test after one-way ANOVA. *p < 0.05, ** p<0.01. Labels above each plot represent comparison to the wild type *ppk* construct within each strain (e.g. *ppk-*sfGFP vs BS1-sfGFP in an MG1655 background). n = 3 biological replicates; error bars represent standard deviation.

To identify putative CsrA binding sites, we analyzed the RNA accessibility map of the *ppk* 5’ UTR (**Figure 4B**). Relative to the other activators identified in this work (*patA, metC, proS*, **Figure 2**), the *ppk* transcript does not show a region of significantly decreased accessibility, however it does show a pattern similar to the recently discovered CsrA-activated RNA *uxuB*^31^. Specifically, there are three peaks of accessibility, two of which are in the 5’ UTR and one is in the CDS of the mRNA (**Figure 4B**). Previous biophysical model-informed predictions of CsrA-*ppk* interactions^20^ did not identify the two putative sites found in the 5’ UTR of *ppk,* and instead found three sites (2 unique, one overlapping with our accessibility assay) all within the CDS (**Figure 4B**). This yields a total of 5 putative CsrA binding sites, 2 of which are predicted exclusively by RNA accessibility (BS1, BS2), 2 of which are predicted exclusively computationally (BS3, BS4), and 1 of which is predicted by both methods (BS5). Thus, *ppk* was nominated by prior evidence and accessibility peaks but not by the decreased-accessibility signature used for the other activated-candidates, and we therefore present it as a case study rather than a prediction.

To test whether these putative binding sites contribute to CsrA-*ppk* interaction, we generated mutant sequences that disrupted the CsrA binding motifs at each putative binding site and cloned them into the *ppk* TRI-FC construct (**Figure 4A**), based on YFP complementation^43^, to test their effect on *in vivo* interaction between CsrA and the *ppk* transcript. Mutation of Binding site 3 (BS3) or Binding Site 5 (BS5) reduced YFP complementation to levels indistinguishable from the *fecA* control (BS3: n=5, FC= 1.07, p=0.68; BS5:n=5, FC=1.00, p > 0.99; One-way ANOVA with Dunnett t-test), whereas the BS4 mutant retained signal significantly above the *fecA* control (**Figure 4A**). Binding Site 1 was not tested because the plasmid could not be constructed for unknown reasons, and mutation of Binding Site 2 abrogated binding but produced noisy results, potentially due to inconsistent binding. Because BS3 and BS5 were required for CsrA association in this assay, we asked whether the same sites were required for CsrA-dependent expression. We inserted the same CsrA binding motif mutations in the same 5’ UTR-*sfGFP* fusion as in **Figure 2A**. We observed that mutation of 3 out of 5 putative binding sites led to significant (p< 0.05) alteration in the fluorescent output of the system in the WT strain, while none showed any change in fluorescence in the *csrA::kan* strain, suggesting that the observed disruptions are CsrA-dependent effects (**Figure 4C**). Specifically, we observed that mutation of binding sites 2, 4, and 5 led to a decrease in signal in the WT strain suggesting reduced activation (**Figure 4C**). The two assays agree for BS2 and BS5; BS3 was required for CsrA association but not for expression, whereas BS4 affected expression without detectable loss of association. We do not yet have a mechanism assigned to individual sites; some sites may be responsible for CsrA binding while others are responsible for regulation, and/or it is also possible that the assays may differ in sensitivity.

Together, these data identify sequence elements in the *ppk* 5’UTR and early coding sequence, nominated by accessibility peaks and/or computational predictions, that contribute to CsrA-dependent expression; BS5, which was supported by both approaches, is also the only site supported by both reporter assays.

### Loss of CsrA activity is associated with reduced polyP and fewer HP-bodies

The *ppk* gene (encoding polyphosphate kinase) is responsible for production of polyphosphate (polyP), a global stress response molecule^44,45^, as well as a recently discovered role in the formation of biomolecular condensates under nitrogen starvation, called “HP-bodies”^30^. Polyphosphate is a globally relevant stress-responsive biomolecule, and its accumulation is induced under many stresses^44–46^, however most recent interest has focused on its role in nitrogen starved cells^30^. Based upon the potential physiological implications of a CsrA-*ppk* regulatory interaction, and our observation that the *ppk* gene showed CsrA-dependent activation (**Figure 2B**) as well as evidence for site specific CsrA-dependent reporter expression, the CsrA-*ppk* interaction was of high interest for further characterization under nitrogen stressed conditions.

We hypothesized that CsrA increases intracellular polyP levels, based on the observed activation of the *ppk* gene (responsible for polyP production) in our work (**Figure 2B)** and others^8^. To test this hypothesis, we measured accumulation of polyP under nitrogen stressed conditions (Gutnick N-media^30^) after 24 hours across three strains: MG1655 WT, MG1655 *csrA::kan,* and MG1655 *Δppk* (**Figure 5A**). In agreement with our hypothesis, we observed that the *csrA::kan* strain is deficient in the accumulation of polyP, showing no statistical difference in production relative to the Δ*ppk* strain (n = 3, FC ∼1, p-value > 0.05, Dunnett T-test) and nine fold lower polyP accumulation in *csrA::kan* relative to the WT strain (FC = 9.01, p = 0.005, Dunnett T-test) (**Figure 5A**). This result suggested that the presence of CsrA may be important for production of the stress responsive biomolecule polyP under nitrogen starvation conditions. However, *csrA::kan* cells have pleiotropic phenotypes, and polyP accumulation during nitrogen starvation is also influenced by the stringent response; notably *relA* is among the candidates flagged by our accessibility analysis (**Table 4**), and CsrA has been linked to the stringent response^37^.

**Figure 5.**
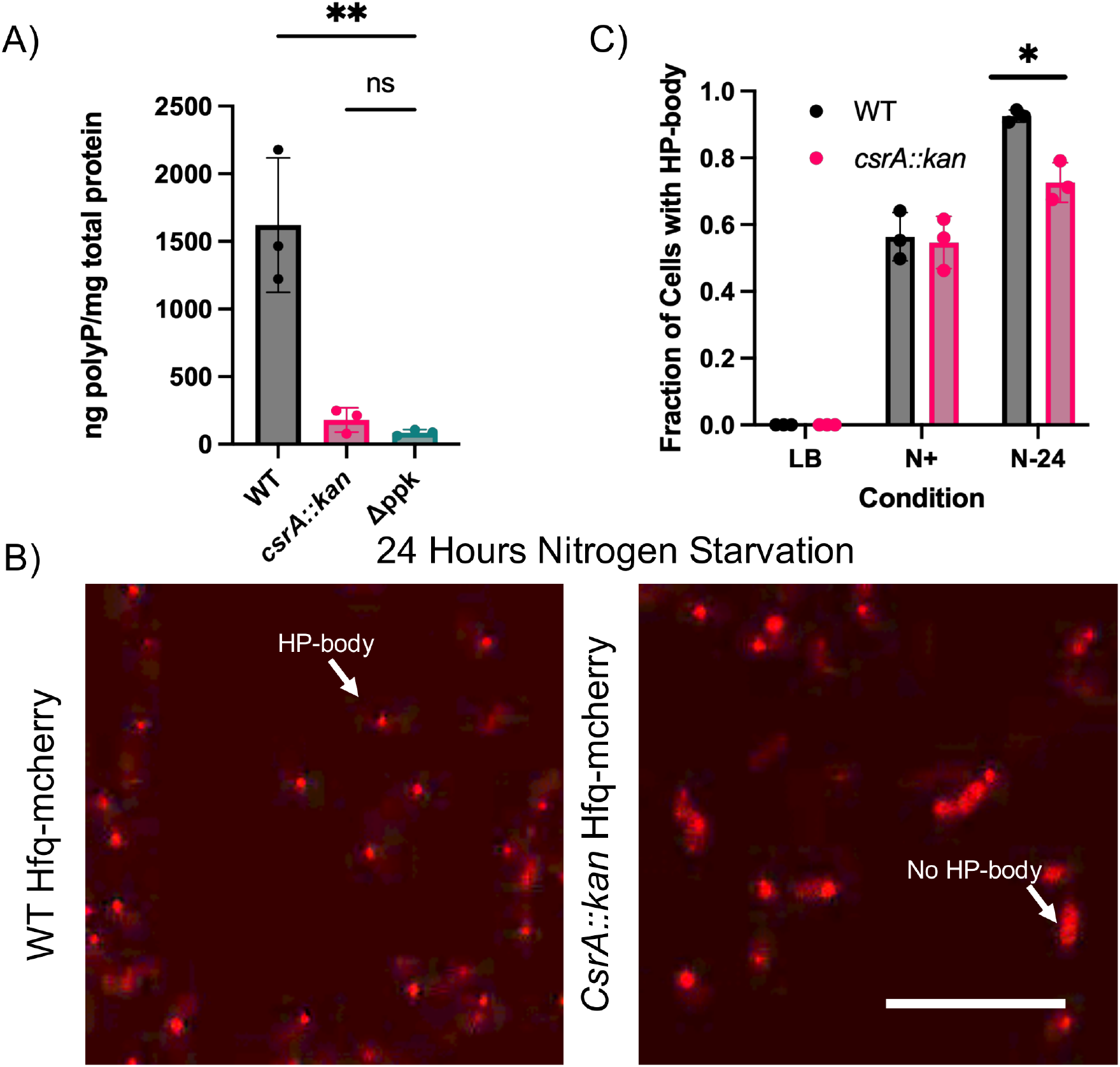
Loss of CsrA activity is associated with reduced polyP and fewer HP-bodies under nitrogen starvation**.(A)** Polyphosphate quantification on N-24 cells. PolyP calculated via mass standard curve of sodium polyphosphate and normalized to mg of total protein in cell lysate. Error bars indicate standard deviation (n= 3). Comparisons performed using the Dunnett test after one-way ANOVA, ** p<0.01; ns, not significant (**B**) Subcellular localization of Hfq-mCherry in the WT *hfq-mCherry* and *csrA::kan hfq-mCherry* strains after 24 hours of nitrogen starvation. Cells containing HP-bodies are those with distinct foci of fluorescence, whereas non HP-body containing cells show diffuse signal. Arrows and labels indicate representative cells containing HP-bodies, or cells that do not. Scale bar=10 um. **(C)** Manual counting analysis of HP-body formation in two strains (WT and *CsrA::kan*) under LB, N+ and N-conditions,, minimum 200 cells per biological replicate, n=3. Comparisons made with 2-sided independent t-test. * p<0.05. HP-Body counting was blinded to strain and condition to avoid bias in data analysis.

To ask whether the polyP difference could reflect post-transcriptional regulation of *ppk* rather than *ppk* RNA levels, we used RT-qPCR to quantify *ppk* mRNA levels in the WT and *csrA::kan* strains after 24 hours of growth in Gutnick N-media. The *ppk* mRNA levels decreased 30% in the *csrA::kan* strain (n=3, 2^-ΔΔCt = 0.707, p = 0.0438) (**Figure S9A**), which is a significant change, though not on the same order of magnitude of change as the polyP levels. Because CsrA can also affect mRNA stability^47^, this modest decrease does not distinguish between regulatory levels. We further tested whether PPK protein levels changed in the absence of csrA by constructing two additional strains (WT, *csrA::kan* background) with genomically labeled *ppk-mCherry*. Using flow cytometry, we measured the basal *ppk-mCherry* fluorescence of these strains after 24 hours of growth under nitrogen-limited media. The WT *ppk-mCherry* strain showed a modest shift in mean fluorescence relative to an unlabeled control (**Figure S9B,C; n=3** for labeled strain and n=1 for the unlabeled control), whereas the *csrA::kan ppk-mCherry* strain was indistinguishable from the control. Because the WT signal was close to background, this result is consistent with, but not conclusive for, a contribution of CsrA to *ppk* expression under nitrogen starvation.

Having observed that loss of CsrA activity is associated with reduced polyP, we were curious if CsrA also had a phenotypic impact on the newly discovered stress-responsive HP-bodies. Beyond polyP, HP-bodies also contain the global RNA binding protein Hfq. Thus, to test for CsrA-dependent alterations in HP-body formation, we used live-imaging confocal fluorescence microscopy to measure HP body formation using genomically labeled Hfq-mCherry (HMC) in WT and *csrA::kan* (**Figure 5B).** These strains, hereafter called WT HMC and *csrA::kan* HMC, were imaged under three conditions: after 4 hours of growth in LB media, after 24 hours of growth in Gutnick N+ media (nitrogen rich), and after 24 hours of nitrogen stressed growth (Gutnick N-media). HP-bodies, observed as condensed regions of high-fluorescence (**Figure 5B**) within a cell, were manually counted (≥200 cells per biological replicate, n=3) for both strains under all conditions. Because *hfq* is itself a candidate CsrA target (**Table 3**), differences in Hfq-mCherry abundance could confound HP-body scoring. In the LB media conditions, no cells showed any HP-body formation (**Figure 5C**), in agreement with the stress-free nature of these conditions. Cells began forming HP-bodies in the nitrogen rich Gutnick N+ conditions, however we observed no significant difference between the WT HMC and the *csrA::kan* HMC strains (**Figure 5C)**. In the nitrogen starvation conditions, however, we observed that the WT HMC strain had the higher fraction of HP-bodies, and the *csrA::kan* HMC strain showed a significant reduction (p = 0.02, 2-tailed independent t-test, n = 3) in the number of HP-bodies (**Figure 5C)**. Specifically, the WT HMC cells had 92.8 ± 1.9% (mean ± s.d.) of cells with HP-bodies, consistent with previously reported values^30^, whereas the *csrA::kan* HMC had 75.1 ± 6.3% (mean ± s.d.) (**Figure 5C)**. Interestingly, the fraction of *csrA::kan* HMC cells with HP-bodies aligns with literature reported values for a *Δppk* HMC strain of approximately 75%^30^. This association is consistent with a model in which CsrA affects HP-body formation through *ppk* in *csrA::kan* cells.

## Discussion

This work broadens the set of mRNAs, sRNAs and cellular processes connected to CsrA. By comparing accessibility in cells with wild-type and reduced CsrA activity, we prioritized candidate CsrA-responsive 5’UTRs, 3’UTRs and sRNAs, tested a subset with reporter assays and in vitro binding assays, and examined one candidate, *ppk,* in more depth. We view accessibility profiling as a prioritization tool to nominate RNA and regions for study that, when paired with orthogonal assays, can establish binding and regulatory outcomes. Existing methods typically identify binding events or regulatory outputs independently^6^, and accessibility profiling offers a complementary in vivo view of RNA state without modifying the protein itself.

Accessibility maps of characterized targets such as *glgC* showed the largest changes near known binding regions, which provides a benchmark for the approach (**Table 1**; **Figure 1D**). Six of nine 5’UTR reporters tested (*mdtE, ribB, proS, metC*, *patA,* and *ppk*) showed CsrA-dependent expression, and CsrA bound the sRNAs SsrA and RalA in vitro (**Figure 2**). Only two of five reporters drawn from the primary candidate list showed CsrA-dependency, however, and four of the six responsive 5’UTRs fell outside that list (**Tables 5** and **6**); the screen is therefore best understood as a sensitive (but unspecific) annotation of candidate regions that allow prioritization for experimental validation. In general, accessibility probing in the presence and absence of CsrA allows for: (i) a position-resolved view of where CsrA-dependent changes occur (as shown strongly for known targets such as *pgaA* and *glgC*), (ii) site-level leads that are functional (as in the case of accessibility nominated BS1, BS2 and BS5 for *ppk,* where BS2 was not even predicted computationally), and (iii) direction-specific signal that motifs and sequence alone cannot give (as decreased accessibility nominated 22 candidate activators with 9 with agreement to previously published proteomics^8^). From an RNA-structure perspective, increased accessibility in *csrA::kan* at most candidate regions, and decreased accessibility at the start codons of candidate activated 5’UTRs, are consistent with CsrA either occluding or remodeling local structure; this model could be tested by structure probing (for example, using SHAPE-MaP^48^) of selected 5’UTRs with and without CsrA in future studies. More broadly, this study raises the possibility that RNA accessibility profiling and enrichment may be applicable to studies of other global bacterial RNA-binding proteins.

CsrA-activated transcripts are rare, and only four have been characterized ^23–25^. Using decreased accessibility near the start codon together with published proteomics, we nominated nine candidate activated 5’UTRs. Reporter data for three of them (*patA, metC, proS*) were consistent with CsrA-dependent activation. Comparing our mechanism for CsrA activation to known activators, only *uxuB* (which both activates and de-activates in a concentration dependent manner) carried the signature of showing a significantly decreased probe, and the *ppk* 5’ UTR was activated without the observed accessibility trend. This suggests that decreased accessibility selects for a subset of CsrA-activated 5’UTRs, perhaps being regulated through similar mechanisms. The *patA* and *metC* genes play a role in putrescine catabolism and methionine biosynthesis respectively, suggesting additional metabolic connections of CsrA. Likewise, the *proS* gene is essential and encodes for a proline-tRNA ligase protein and suggests that CsrA may be involved in regulating translational machinery in the cell rather than just translational efficiency. Each of these candidates requires further work to establish direct regulation and elucidation of a mechanism, which will be of focus for future manuscripts.

Our data are consistent with a broader overlap between CsrA and sRNA networks than currently appreciated, although several caveats apply. Of the 19 previously unreported candidate sRNAs (most of which were selected manually due to a low number of probes) we tested *in vitro* binding for only two (RalA and SsrA (tmRNA)) at high CsrA concentrations. Therefore, although our results suggest an interaction, it is unknown whether CsrA binds and regulates SsrA or RalA *in vivo*, and whether that impacts the phenotype of these sRNAs (**Figure 2**). In addition, reporter assays suggest that CsrA may alter the output of the ArrS-*ribB* and GcvB-*crp* pairs, extending prior evidence that CsrA can modulate sRNA-mRNA regulation^12^; these data do not yet distinguish effects on sRNA expression, sRNA stability or target accessibility.

Not all accessibility-positive targets yielded measurable translational regulation under the conditions examined. Of six candidate targets tested (*ribB, mdtE, uxuA, ompR, ppk, rpsA),* two showed clear translational repression (*ribB, mdtE*), one was activated (*ppk*), and three showed no effect (*ompR, uxuA, rpsA*) in our assay. Only two of the five reporters drawn from the primary candidate list (*mdtE* and *ppk*) showed CsrA-dependency in our assays, whereas *ribB* was added based on prior evidence and fell just below the probe threshold (**Table 2**). These outcomes highlight an important distinction between accessibility changes and regulatory outputs. Accessibility signatures may reflect direct binding events, condition-dependent regulation, interactions with additional RNA-binding factors, or regulatory effects occurring outside the specific experimental conditions tested here. Our INTERFACE accessibility probing and the reporter assays were performed in LB media, and it is possible that regulation specific to other conditions would be missed. Indeed, previous work has identified several CsrA targets that were bound by CsrA *in vivo* but not regulated, or regulated *in vivo* but seemingly not bound^8^. For example, the *yecC* 5’ UTR shows binding without regulation in previous work^8^ and is identified as a potential binder in this work. Similarly, previous CLIP-seq data did not capture a majority of the RNAs with altered translation or translation efficiency^6^. Consequently, accessibility profiling should be viewed as a prioritization strategy for identifying biologically relevant candidate interactions rather than a direct predictor of regulatory function in every case.

The *ppk* gene, identified as a putative CsrA target by RNA accessibility screening, was selected for further characterization due to its relevance to stress and its newly discovered role as a mediator of biomolecular condensation. Using a combination of our RNA accessibility pipeline and previous computational studies^20^ we identified 5 putative CsrA binding sites on the *ppk* transcript (**Figure 4**). We tested the five sites by mutagenesis; BS5 was supported by both reporter assays, whereas the other sites were supported by one assay or neither. The computational method identified binding sites near the start codon that our accessibility data did not flag; one possible explanation is competition with the ribosome for RNA access, which we did not test. Similarly, the computational screen did not identify two binding sites in the 5’ UTR of the *ppk* transcript. This supports the notion that neither computational nor experimental techniques are exhaustive for identifying CsrA-RNA binding sites and interactions. *In vivo*, loss of CsrA activity was associated with reduced accumulation of polyP under nitrogen limited conditions and with fewer HP-bodies (**Figure 5**), but these associations were measured in a single partial mutant and do not yet show that *ppk* regulation accounts for them. Interestingly, *ppk* appeared to be primarily activated by CsrA, without a strong region of decreased accessibility in the *csrA::kan* strain. This suggests that there are other, less obvious, mechanisms of activation by CsrA than those used to identify *patA, metC,* and *proS*. We quantified polyP levels in two strains with varied activity levels of CsrA in addition to a *ppk* knockout negative control strain. We showed that the *csrA::kan* (low CsrA activity) strain accumulated polyP equivalent to the *ppk* knockout strain and far lower than the WT strain. Both polyP and CsrA have been linked to virulence in diverse bacterial systems^4,44,49,50^, so a connection between them may be of interest, although our data do not address virulence. Finally, loss of CsrA activity was associated with reduction in the formation of HP-bodies, comparable to that reported for a strain lacking *ppk* ^30^. Together, these data suggest that CsrA may connect to polyP homeostasis and HP-body condensate formation, a possibility that merits direct testing.

In this work, we identify several new putative CsrA targets, both mRNAs and sRNAs. Compared to several previous studies^6,8,21,37^, we identified nine new putative mRNA targets, however many of the mRNAs that were not classified as putative targets in this work showed significant previous evidence of CsrA regulation. RNA accessibility is fundamentally a less specific screen for CsrA binding, as it cannot account for indirect interactions, whereas methods such as CLIP-seq are driven by physical RNA-protein interactions. RNA accessibility thus may find CsrA-RNA interactions less easily observed (whether through steric or other limitations), however it may also miss other targets if the RNA is regulated or bound by several RNA binding proteins or other factors.

### Limitations

We acknowledge that there are limitations to this analysis, as we could not capture all known CsrA targets, nor were we able to confirm all potential targets that showed strong accessibility signatures. First, *csrA::kan* is a partial loss-of-function mutant (approximately 10% of wild type activity) with plieotropic effects, and our in vivo conclusions rest on comparisons with this mutant; therefore, based on this assay alone, we describe effects as CsrA-dependent rather than direct unless supported by motif mutations or binding site data. Second, we observed a global increase in probe accessibility in this *csrA::kan* mutant, including for the scrambled control **(Figures S1 and S2)** and cannot exclude contributions from indirect effects on probe expression, reporter translation or transcription termination; we address this by normalization in our data analysis (Methods), but orthogonal validation is needed. Third, the candidate-calling criteria (top 25% of probes within each RNA, antisense to GGA/AGA) have limited specificity because a GGA or AGA motif is likely to occur somewhere within the 10-20 nt windows within probes. Furthermore, the library is enriched for literature-reported CsrA targets, and 3’UTRs and most sRNAs were nominated manually because shorter RNA regions cannot carry eight significant probes.

A permutation analysis showed no enrichment of motif-containing probes among top-quartile probes beyond chance (87 called versus 85.7 expected, *p* = 0.43), so the screen has high sensitivity but low specificity, and an estimated 98% of its calls are expected by chance. It is worth noting that this null randomizes motif assignments among probes and does not account for secondary structure, probe overlap, local sequence composition or other features that could shape fold-change patterns. The library was built from putative CsrA targets and may consist largely of true targets, in which case no filter could enrich for them, and with the numbers of probes and targets available a strong filter is unlikely. A stricter screen that required AU-rich flanks around the motif separated modestly from the null (*p* = 0.029) but lost known targets (CsrB, hfq and mdtE), and only five of its 34 targets overlapped published CLIP-seq peaks; this comparison may also reflect differences in strain, growth condition and what each method measures. We therefore consider validation of prioritized targets as essential.

Fourth, accessibility probing and reporter assays were performed in LB so condition specific effects may be missed. Fifth, the associations between CsrA, polyP and HP-bodies were measured in *csrA::kan* cells and could involve routes other than *ppk,* including the stringent response and *hfq* expression as CsrA has also been linked with regulating *hfq*^51^ and *relA/dksA* in the stringent response pathway^37^. It was also infeasible to design a library of probes with efficient coverage over the entire transcriptome, due to the size of the library and the ability to capture enough read depth for all probes. While there are many advantages to this systems-level probing approach used in this study to better understand the CsrA network, it is possible that loss/inactivation of CsrA causes other effects that influence RNA accessibility patterns including transcriptional changes, altered expression of other RNA-binding proteins, altered ribosome occupancy and altered RNA expression and stability. While there is ample evidence that altering CsrA impacts steady-state RNA abundance^6,8,47^, we do not have evidence to elucidate the role of target RNA abundance on the fidelity of asRNA probing. To mitigate this concern, the fold changes for probes were internally normalized to the mean abundance of every probe on their target RNA. It was not feasible to perform plasmid-based CsrA re-complementation for assays subjecting the *csrA::kan* strain to nitrogen stress, due to the cells’ inability to grow to high densities under antibiotic stress. To mitigate any potential effects, the genome of the *csrA::kan* strain was sequenced, and no obvious compromising mutations were found. Despite these limitations, we believe the data support the use of accessibility profiling to prioritize CsrA-responsive RNA candidates for follow up and that this could be a useful approach to dissect large datasets to uncover the regulome of global RNA-regulatory proteins like CsrA.

## Conclusion

Together, our findings broaden the known landscape of CsrA-responsive RNAs and highlight connections between CsrA, sRNA regulatory networks and polyphosphate homeostasis. By comparing RNA accessibility under wild-type and reduced CsrA activity, we prioritized candidate mRNA, 3’UTR and sRNA targets, nominated candidate CsrA-activated 5’UTRs, and found CsrA-dependent expression for six of nine reporters tested. Accessibility profiling is therefore a useful way to prioritize RNAs for study, with a validation rate that we report transparently. Future work should test direct binding at prioritized sites by mutagenesis and in vivo crosslinking, determine whether SsrA and RalA interact with CsrA in vivo, and establish whether reduced polyP and HP-body formation in CsrA-deficient cells reflect reduced PPK expression. Lastly, the effects of how cellular fluctuations of CsrA concentration affect RNA accessibility still need further exploration.

## Supporting information

Raw Microscopy Images

## Acknowledgements

This work was supported by the Genome Sequencing and Analysis Facility (GSAF) at UT Austin, Center for Biomedical Research Support, and the Microscopy and Flow Cytometry Core at UT Austin (RRID:SCR_021756). Computational analysis in this work was supported by the Texas Advanced Computer Center (TACC) at the University of Texas at Austin. URL: http://www.tacc.utexas.edu. We would like to thank Matthew Burroughs for his thoughtful feedback, and the rest of the Contreras Lab for their helpful input. Some images were made using Biorender and can be accessed at the following links: Fig. 1 (BioRender.com/7qqxjlz) Data were visualized using GraphPad Prism.

## Materials and Methods

### Bacterial Strains and Growth Conditions

All strains are available in **Table S4.** *E. coli* strains used in this study are based upon K-12 MG1655. All cloning was done in *E. coli* DH5α. The INTERFACE experiment was performed in strains WT (MG1655 ΔtnaA ΔaraA) and *csrA::kan* (MG1655 *csrA::kan* ΔtnaA ΔaraA), where CsrA is partially knocked out and functionally ∼10% as effective as WT. Deletions of *tnaA* and *araA* were included to ensure maximum probe elongation and probe expression, respectively, in accordance with optimal conditions for the RNA-seq compatible INTERFACE accessibility probing method used^52–54^. Cell growth in rich media was performed in LB media grown at 37C with orbital shaking at 200rpm.

### Strain construction

All genome editing was performed using an established CRISPR-cas9 protocol^55^. gRNAs were designed using the gRNA tool by atum.bio (https://www.atum.bio/eCommerce/cas9/input).

### Plasmid Construction

The INTERFACE library plasmids were constructed by Genscript. Plasmids to test UTRs and sRNA-UTR interactions were constructed using Gibson assembly. Oligonucleotides and gBlocks were purchased from integrated DNA technologies (idtdna.com) and Twist Biosciences (twistbioscience.com). These constructs are available in **Table S5**. Gibson assembly primers were designed according to previous protocols^55^. Plasmids were verified via Sanger sequencing (Eton Biosciences, etonbio.com) and Plasmidsaurus (plasmidsaurus.com). Vanillic acid inducible promoters were sourced from the Marionette collection^56,57^.

### Probe Library Design

An automated pipeline for designing the INTERFACE antisense RNA (asRNA) probes targeting mRNA 5’ and 3’ UTR regions was implemented in Python 3.7, leveraging Biopython modules for sequence handling and NCBI BLAST command-line tool integration. For each nucleotide across the library of UTRs, base pairing probabilities were first calculated using the pairs command in NUPACK 3.2, which generates position-specific pairing probabilities, *p(i_n_ · j_n_),* for each nucleotide in the input sequence^58^. Positions with no reported value were treated as unbound and assigned a probability of 0. To minimize targeting of structurally dynamic regions, probes were positioned within windows of low structural variability by computing the standard deviation of unbound probability (1-P) at each position as described in previous work^59^ using the Python statistics module (N-1 degrees of freedom), then centering each probe on the nucleotide position with the lowest variance. The probe-design and analysis code is available at github.com/ryanbuchser/INTERFACE_Analysis.

Genome specificity of each candidate probe was assessed using the NCBI BLAST+ command-line tool (v2.6.0)^60^, which was selected over global alignment approaches for its statistical e-value framework. A local BLAST database was constructed from the *Escherichia coli* K-12 MG1655 reference genome (accession: U00096.3) using makeblastdb, and queries were executed programmatically via the Biopython NcbiblastnCommandLine^61^ wrapper with the following parameters: task set to blastn-short, output format 6 (tab-delimited), e-value cutoff of 50, and a maximum of 10 reported alignments per query (max_hsps=10). Rather than applying a conventional e-value threshold, probe specificity was evaluated by the magnitude of separation between the top two reported alignment scores, calculated as the 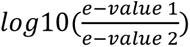. Probes were designated as passing if this magnitude shift exceeded 0.5.

Targets were selected from multiple literature sources, including translational (5’UTR-GFP fusion) reporter assays^8,20^, RNAs suspected from high throughput datasets^6,20,37,47^, as well as RNAs predicted via sequence searches^21,62^. Consequently, the library is enriched for RNAs with prior evidence of CsrA association. The mRNA targets were also biased for 5’UTRs, as most direct regulatory CsrA-mRNA interactions reported involve the 5’UTR^4,6,8^, with a few more recent exceptions involving the 3’UTR^6^. To supplement the UTR-tiling probe library, additional probes were designed to directly target regions of known or predicted protein-RNA and RNA-RNA interaction. Using previously predicted CsrA binding site coordinates^20^, 504 site-specific probes were generated extending outward from each central binding region. These probes ensure direct coverage of protein-RNA interaction sites and may also facilitate antisense regulation through a toehold-switch mechanism, in which hybridization near a stem loop can expose a sequestered ribosome binding site (RBS).

### Plasmid Library Preparation

1.5 ug of purified library was transformed into competent cells of the strains of interest and recovered in 1000uL of LB Media supplemented with 0.4% glucose. The transformants were allowed to recover at 37C with shaking for 90 minutes. After 90 minutes, 2x 10uL samples were removed from the recovery and plated on LB Agar + Carbenicillin at 37C overnight. The resultant colonies were counted to ensure that there were at least 10x the library size in number of colonies (∼71,000) per strain when accounting for the dilution factor of 600x. The remaining 5980uL of recovery was added to 44 mL of LB + Carbenicillin and grown overnight at 37C with orbital shaking. The next day, the OD_600nm_ was measured to determine seeding density, and the cells were mixed 1:1 with 30% (w/v) glycerol and frozen at -80C to be used in the INTERFACE experiment.

### INTERFACE Experiment

The frozen stocks containing the INTERFACE library for each strain were thawed on ice and diluted in three 50 mL flasks with LB media and carbenicillin to a starting OD_600nm_ of 0.05. After each sample was grown to an OD_600nm_ of 0.2-0.3, they were induced with 0.2% arabinose to induce probe expression and allowed to grow for 5 hours to ensure maximum probe expression. Cells were pelleted and frozen at -80C to prepare for RNA extraction.

### RNA Extraction

RNA was extracted following previous methods^7^ with alterations. Briefly, cell pellets were thawed on ice before being resuspended in lysozyme buffer (2mg/mL, Thermo Fisher Scientific) and incubated at room temperature for 10 minutes. DNA/RNA Shield (Zymo Research) was added to each lysate before being spun down at 21,300xg. The supernatant from this spin was used in the Quick DNA/RNA Miniprep Plus Kit (Zymo Research).

### Small RNA Library Preparation

Small RNA libraries were prepared at the University of Texas Genomic Sequencing and Analysis Facility using the NEBNext Small RNA Library Prep Set (New England Biolabs, Cat. No. E7330). For each sample, 5 µg of total RNA was used as input. Library construction followed the manufacturer’s protocol with the following modifications: the 3’ adapter ligation step was performed with an 18-hour incubation at 16 °C without a heated lid, and library amplification was performed using 12 PCR cycles, with a 70 °C, 1-minute extension for each cycle.

Following amplification, libraries were purified using AMPure XP beads (Beckman Coulter) at a 2.4X bead-to-sample ratio. Library size distribution and quality were assessed using an Agilent 2100 Bioanalyzer with the High Sensitivity DNA Kit (Agilent Technologies, Cat. No. 5067-4626). Libraries were pooled equimolarly and processed for size selection on a BluePippin system (Sage Science) using a 2% agarose gel cassette, with a target size range of 200–500 bp. Prior to sequencing, pooled libraries were quantified using the KAPA Library Quantification Kit for Illumina platforms (Roche, Cat. No. KK4602) to determine appropriate loading concentrations.

Sequencing was performed on an Illumina NextSeq2000 platform using paired-end, 150-bp reads (PE150).

### RNA-seq data analysis and calculation of relative accessibility

RNA reads were processed using resources available through the Texas Advanced Computing Center (TACC) using previously developed pipelines^27^. Raw reads from the INTERFACE sequencing were first aligned to a created reference genome containing all the probes using burrows wheeler alignment^63^. The resulting sam file was then converted into a bam file using samtools^64^ before finally being converted into a bed file using bedtools^65^. The resulting bed files were processed using previously published python code^27^ that maps the number of reads at each end position for each unique INTERFACE probe. RNA accessibility was calculated based upon the ratio of reads that extended beyond the 3’ end of the *rut* site (long reads) relative to the number of reads which did not (short reads). This length ratio (probes longer than the 3’ end of the *rut* site relative to probe shorter) was computed for all probes across replicates allowing for calculation of a mean and standard standard deviation length ratio per probe per strain.

### INTERFACE Data Analysis

Probes which reached adequate sequencing depth (20+ total reads in all replicates) were compared between the WT and *csrA::kan* strains for statistical significance of accessibility changes. A minimum sequencing depth of 20 reads was selected to reduce noise associated with low-count probes and to enable reliable estimation of accessibility ratios across biological replicates. An adjusted p-value cutoff of 0.05 was used to determine statistical significance using a Benjamini-Hochberg multiple t-test correction^66^. INTERFACE data were collected three independent cultures per strain started from a single library transformation. A table of each RNA probe (including its sequence, location on the target RNA, whether it had sufficient RNA-seq data, fold change, and adjusted pvalue) is included as **Table S6**; per-RNA coverage statistics are included in **Table S7**. All RNA probing maps are available in **Figure S10.**

### Filtering for Potential Binding Sites using INTERFACE Data

First, we calculated the mean accessibility fold change for all probes that showed a significant change in accessibility (*p* <0.05, two-tailed unpaired t-test, Benjamini-Hochberg correction) between WT and *csrA::kan,* yielding a total of 3,024 probes. These probes covered all targets in the library with at least one probe, excluding 1 5’UTR, 5 3’ UTRs, and 1 sRNA (**Table S8**). All target RNAs that contain at least 8 probes with a significant change in fold change of accessibility between the two strains were moved to downstream analysis. A minimum of eight significant probes was required to ensure that accessibility signatures reflected consistent regional behavior rather than isolated probe-level fluctuations. To identify the most prominent accessibility features within each RNA, probes were split into quartiles of fold change (bottom 25%, middle 50%, top 25%). The top 25% of probes by accessibility change between the two strains were selected for further analysis as ‘high fold change’ or ‘peak’ probes. From these high fold change probes, we screened for those that paired to a region containing a GGA or AGA motif, suggesting a possible CsrA binding site. To identify potential functional 5’ UTR targets we further screened for probes that bound to their targets within -100 nt to +20 nt of the start codons, as CsrA often interacts with its targets in this region. Additionally, RNAs containing two GGA/AGA motif probes that were both above the average accessibility for the given RNA were flagged as potential CsrA targets and added to **Table 3**. This approach reduced the influence of low-amplitude accessibility fluctuations and enriched for regions corresponding to established CsrA binding sites. To test whether the screen’s signature, a high-fold-change probe overlapping a GGA/AGA site, is informative beyond chance (i.e. specificity), we compared the number of targets it called with the number called when motif-carrying probes were randomly reassigned to positions within each target. Specifically, we performed 20,000 permutations in which each target’s number of probes, fold changes and positions were held fixed and probes were randomly reassigned to positions within the target (preserving the motif count per target). The expected number of calls is the mean over permutations, enrichment is the observed number divided by the expected number, the *p*-value is the fraction of permutations with at least as many calls as observed, and the permutation-based false-discovery estimate is the expected number divided by the observed number (**Figure S5**). We repeated the analysis for the top 25%, 20%, 15%, 10% and 5% of probes, for GGA and AGA separately (**Table S1**), and for GGA/AGA with at least 50% or 60% AU in the 6 nt flanking the motif (**Table S9**), and recorded recovery of the verified positive controls (*csrA, flhD, glgC, hfq, iraD, pgaA, rpoE and sdiA*). CLIP-seq comparison used the peaks reported by Potts^6^. As a sequence-only baseline, we counted GGA/AGA motifs in each 5’UTR and compared verified positive controls with other targets.

RNAs that met the probing criteria and contained GGA/AGA peaks or were manually annotated to have GGA/AGA peaks are included in **Table 3** (87 5’UTR, 32 3’UTR and 28 sRNAs). RNAs that met the probing criteria but did not have any GGA/AGA peaks are included in **Table 5** (44 5’UTR, 1 3’UTR and 5 sRNAs), and RNAs that did not meet the adequate probing criteria are included in **Table 2**, (170 5’UTR,117 3’UTR and 68 sRNAs). Several sRNAs and 3’ UTRs exhibited a low number of probes, and thus peaks were manually selected if they showed high accessibility and presence of an AGA/GGA motif. Those RNAs are included in **Table 3** and noted as “manually annotated.”

### 5’UTR-GFP Assays

3 colonies of each strain (WT, *csrA::kan*) containing the plasmid of interest were grown overnight in 5 mL of LB + their respective antibiotic at 37C with 200 rpm shaking. The next day, the cultures were reinoculated into fresh 5 mL LB + antibiotic and grown overnight at 37C with 200 rpm shaking. After 18 hours, the cells were diluted 1:4 in 1x phosphate buffered saline and fluorescence was measured on a Bio-Tek Cytation3 plate reader using 488/525 ex/em.

### CsrA Purification

CsrA was purified using previously published methods with modification^12^. Briefly, His6-SUMO-CsrA was expressed from a pET-28a(+) vector in BL21 DE3 *E. coli*. 5mL saturated overnight cultures were diluted 1:100 in 500mL of LB + kanamycin. Protein expression was induced with 1mM IPTG after growth to OD_600_ of 0.6 at 37C with 200rpm shaking. After overnight growth at room temperature (25C), the culture was harvested via centrifugation. The pellet was resuspended in 1X PBS before being frozen at -80C. Pellets were resuspended in lysis buffer (50 mM NaH2PO4, 300 mM NaCl, 10 mM Imidazole, 5 mM MgCl2 and 10% glycerol, adjusted to pH 8.0 with NaOH) before being sonicated for 15 minutes (5s on, 5s off) at 40% power (Q125 sonicator). Soluble lysates were collected via centrifugation, and purification was performed on Qiagen Ni-NTA agarose columns. Columns were washed with 20mM, 50mM and 75mM imidazole before being eluted with 250mM imidazole. The resulting eluate was dialyzed overnight at 4C into storage buffer (10 mM Tris-HCl, 100 mM KCl, 10 mM MgCl2, 25% glycerol, pH 7.0) before being concentrated down and used for EMSAs.

### Electrophoretic Mobility Shift (EMSA) Assays

Electrophoretic mobility shift assays were performed in line with previously described literature^12^ with the following modification: RNAs were generated using *in vitro* transcription (MEGAScript) and subsequently Cy5 fluorescently labeled using Mirus Bio LabelIT, and yeast total RNA was omitted. Primers used to generate T7-ssrA and T7-ralA are in **Table S5.** Protein was serially diluted 1:1 for each concentration. Gels were run at 170V for 2 hours in 0.5X TBE before being imaged on the Typhoon RGB with automatic voltage adjustment.

### CsrA-dependent sRNA regulation assays

3 colonies of each strain containing the plasmid of interest (5’UTR-GFPmut3 + pLac-sRNA) were grown overnight in LB + antibiotic with 200rpm shaking. These cultures were inoculated into 30 mL of LB + antibiotic in flasks and grown for 2 hours at 37C with 200 rpm shaking. After 2 hours, expression of the sRNA target was induced via addition of 1 mM IPTG. Fluorescence was measured using a Bio-Tek plate reader using 484/536 ex/em after 4 hours.

### Measuring *in vivo* binding with Tri-Fluorescence Complementation

Fluorescence assays were performed as described in previous work^31^ with small modification. Both plasmids (MS2BD-*ppk*/CsrA-NYFP + MS2CYFP) were transformed into MG1655 *ΔcsrBC*. Five colonies for each test condition were picked and grown overnight in 5mL LB media at 37C with 200 rpm shaking. These overnight cultures were inoculated (1:100) in fresh LB media with antibiotics and grown for 22 hours at 37C with 200rpm shaking. YFP fluorescence was measured on the BioTek Cytation 3 plate reader with an excitation/emission of 500nm/530nm rather than through flow cytometry.

### Extraction and Quantification of PolyP

Cultures of each strain were grown overnight in 5mL LB media, followed by a 1:100 dilution into 5mL Gutnick N+ media, and then grown to their respective time points in a 1:100 dilution of 5mL Gutnick N-media. The 5mL cultures of N-media were pelleted and resuspended in 250 uL GITC lysis buffer (4M Guanidinium isothiocyanate, 50mM Tris-HCl, pH 7)^67^ and lysed at 95C for 5 minutes before the polyP was column purified according to previous methods^68^. Purified polyP was mixed 1:1 with 30uM JC-D7 in HEPES pH 8.0 buffer, and allowed to sit for 5 minutes before being measured via plate reader (BioTek Cytation 3) with an excitation emission of 410/510nm. TE buffer + JC-D7 was used as a blank control. polyP concentration was compared to a standard curve of sodium polyphosphate (Thermo Fisher Cat. No. 390932500).

### Growth of Bacterial Samples for RT-qPCR

Three colonies of each CsrA variant strain (MG1655 WT, MG1655 *csrA::kan*) were first grown overnight in 5mL LB media until saturation. Next, these were grown overnight in 5mL Gutnick N+ media (33.8 mM KH2PO4, 77.5 mM K2HPO4, 5.74 mM K2SO4, 0.41 mM MgSO4, 10mM NH4Cl, 0.4% Glucose, M9 trace elements) until saturation. Each N+ overnight was then inoculated into 5mL of fresh Gutnick N-media (33.8 mM KH2PO4, 77.5 mM K2HPO4, 5.74 mM K2SO4, 0.41 mM MgSO4, 3mM NH4Cl, 0.4% Glucose, M9 trace elements) and grown to the appropriate time point. These samples were pelleted at 4,000 rpm, supernatant was decanted, and the pellets were frozen at -80C. These samples were stored for RNA extraction and the resulting RNA was used for RT-qPCR.

### Reverse-Transcription Quantitative PCR

Relevant RNA samples were thawed on ice before being subjected to RT-qPCR using the Luna one-step RT-qPCR kit (New England Biolabs). Primer efficiencies were calculated by performing 10-fold dilutions of RNA samples and calculating the slope of the C_T_ vs. log_10_ dilution. The efficiencies were then used for all following RT-qPCR experiments. Relative quantification was performed by calculating the C_T_ difference between the gene of interest and the 16s rRNA housekeeping gene for each replicate and then calculating mean and standard deviation^69^.

### Flow Cytometry

Samples containing genomically labelled *ppk* were grown overnight in 5mL of LB media at 37C with orbital shaking at 200 rpm, followed by overnight to saturation at the same conditions in 5mL of Gutnick N+, and finally 5mL of Gutnick N-media to the desired time point. Samples were then diluted 5uL to 1mL of phosphate buffered saline (GrowCells) and mCherry fluorescence was quantified on the Fortessa flow cytometer (Becton Dickenson) using the PE-mCherry setting (561 excitation laser and 610/20 bandpass filter).

### Fluorescence Microscopy

For nitrogen stress experiments, individual *E. coli* colonies containing genomic fluorescently labeled protein (WT HMC, csrA::kan HMC **Table S4**) were grown overnight in 5mL LB media at 37C with 200rpm shaking. The next day, these cells were inoculated and grown overnight in 5 mL Gutnick Media N+ (33.8 mM KH2PO4, 77.5 mM K2HPO4, 5.74 mM K2SO4, 0.41 mM MgSO4, 0.4% (w/v) glucose, 10 mM NH4Cl and M9 trace elements)^30^. The third day, the cells were inoculated 1:100 in Gutnick Media N-(33.8 mM KH2PO4, 77.5 mM K2HPO4, 5.74 mM K2SO4, 0.41 mM MgSO4, 0.4% (w/v) glucose, 3 mM NH4Cl and M9 trace elements), and grown for 24 hours. Afterwards, 1 mL of sample was spun down at 8,000 rcf for 1 min. Supernatant was removed and cells were resuspended in 400 uL of their respective media (Gutnick N+, Gutnick N-, LB). Corresponding (N-media, N+ media, LB) agarose covers were made to immobilize the cells^70^. In total, 10 uL of sample was pipetted in 2 uL fractions on each microscopy chamber well (Cellvis 8-chambered coverglass system C8-1.5H-N, California). An agarose coverslip was added to the surface in order to keep the cells at the bottom of the slide. N+ media controls used 1:100 dilution into Gutnick N+ media (10 mM NH4Cl) and LB controls were inoculated from LB overnights and imaged after 4 hours of growth. Fluorescence microscopy was performed on the Nikon AXR-NSPARC microscope, at the UT Austin microscopy and flow cytometry facility (RRID:SCR_021756), using a 60x objective. The 561 nm laser was used for excitation of mcherry-Hfq. Due to live cell imaging, all samples were imaged within 90 minutes and samples were imaged in randomized order to ensure limited and random granule changes over time. Granules were then manually counted for >200 cells per biological replicate. The images in **Figure 5B** are converted tiff files with the LUTs altered to have a maximum of 2000 to more easily see the results.

### Statistical Testing

Statistical significance of accessibility change between the two strains was calculated using two-tailed heteroscedastic t-tests between the *csrA::kan* and WT strains, and p-values were corrected for multiple t-tests using the Benjamini-Hochberg correction^66^. For testing between strains in the 5’ UTR assays, an unadjusted heteroscedastic t-test was used for comparisons. For computation of the permutation null, p value was calculated using previously published statistical methods for permutational analysis and Monte-Carlo simulation^71^, where p = (B+1)/(N+1), B is the number of permutations that reach or exceed the observed number of hits, and N is the number of permutations tested.

## Author Contributions

This study was designed by R.B, A.J.L., M.K.M, A.N.L, T.S and L.M.C. Probe library design was performed by A.J.L, A.N.L, M.M, and T.S. All microscopy was performed by K.T., and all other experiments were performed by R.B. and J.A.C. The manuscript was written by R.B. and edited by all authors. All authors have approved the final version of the manuscript.

## Generative AI Use

Generative AI was used to assist with the preparation of data analysis code. All generated code was tested using controls and manual analysis to validate its accuracy. Generative AI was never used for generation of *de novo* text, figures, or references.

