## Supplementary figures and images for "CsrA-dependent RNA accessibility landscapes prioritize new mRNA and sRNA targets and link CsrA to polyphosphate homeostasis"

### CSRAKAN_LB_REP1_IMAGE1_RGB_MCHERRY.tif

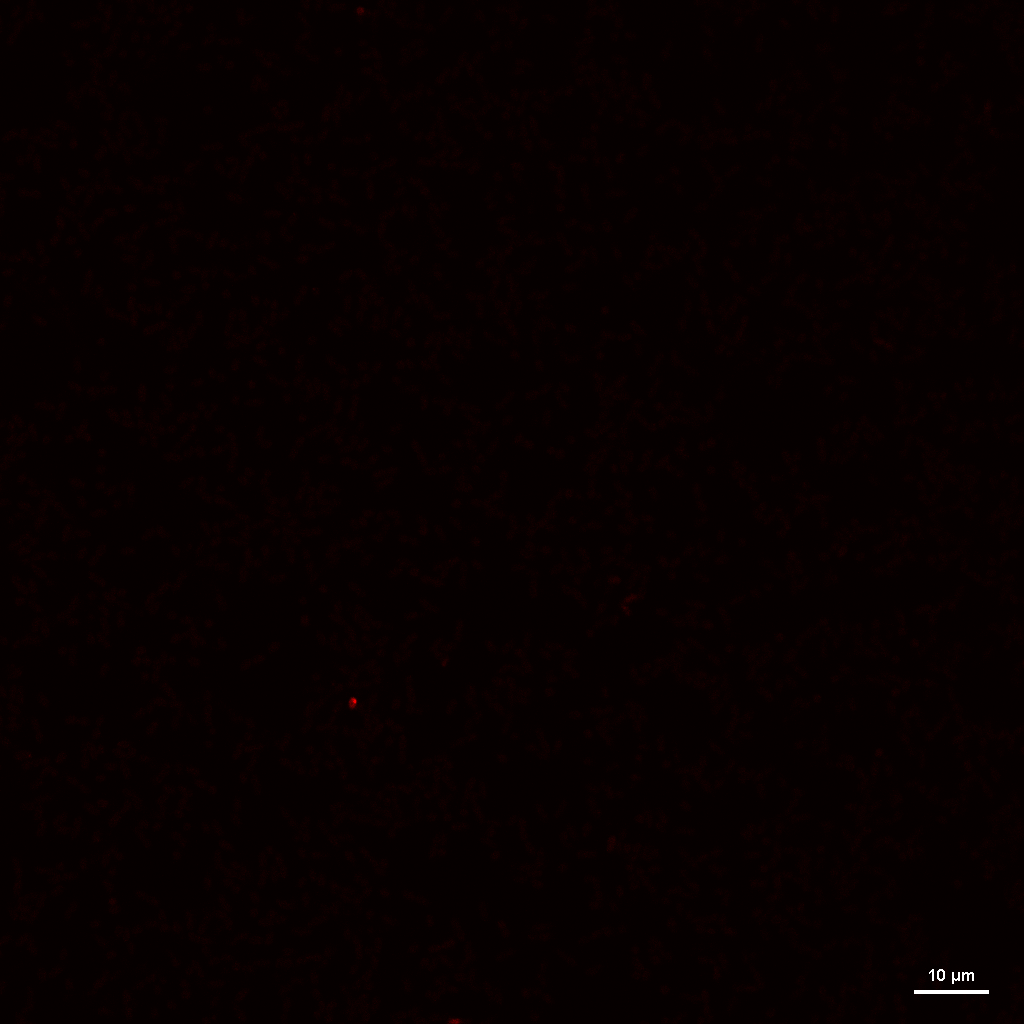

### CSRAKAN_LB_REP1_IMAGE2_RGB_MCHERRY.tif

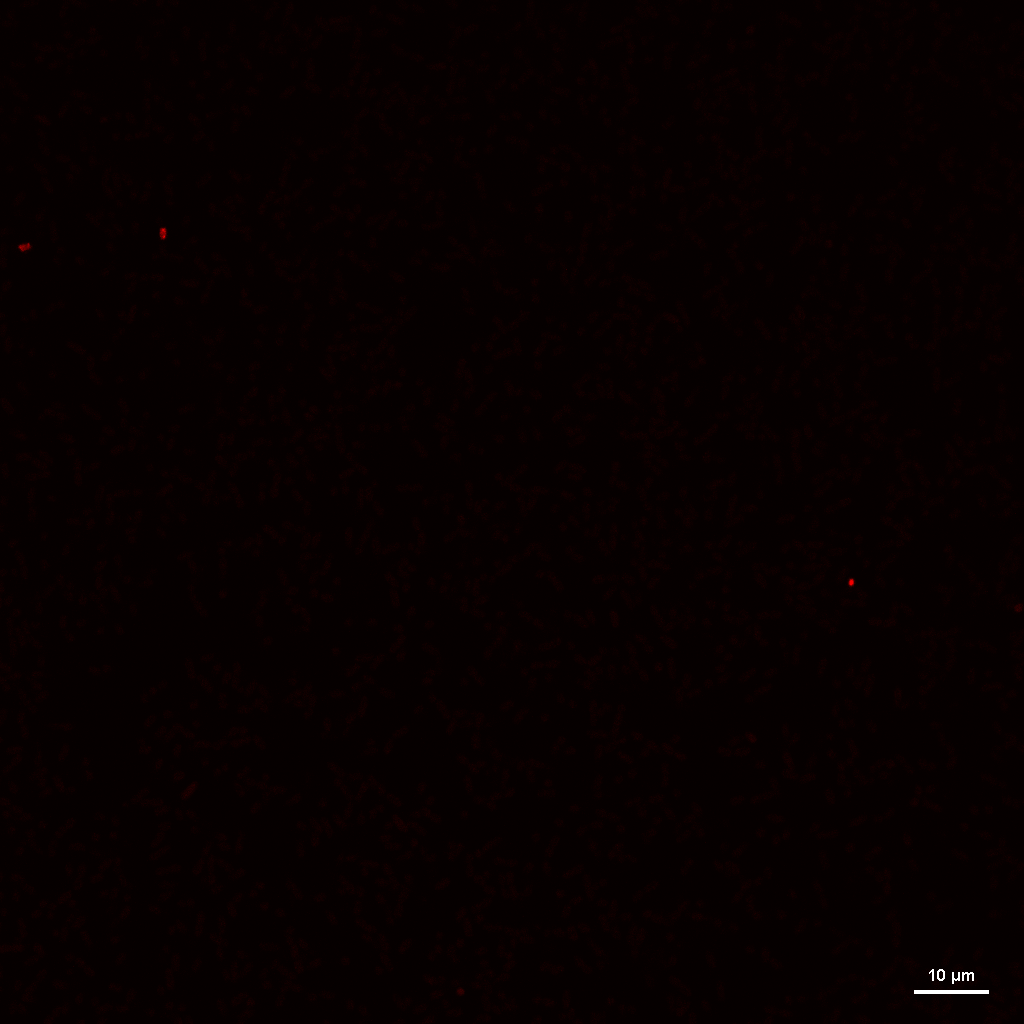

### CSRAKAN_LB_REP2_IMAGE1_RGB_MCHERRY.tif

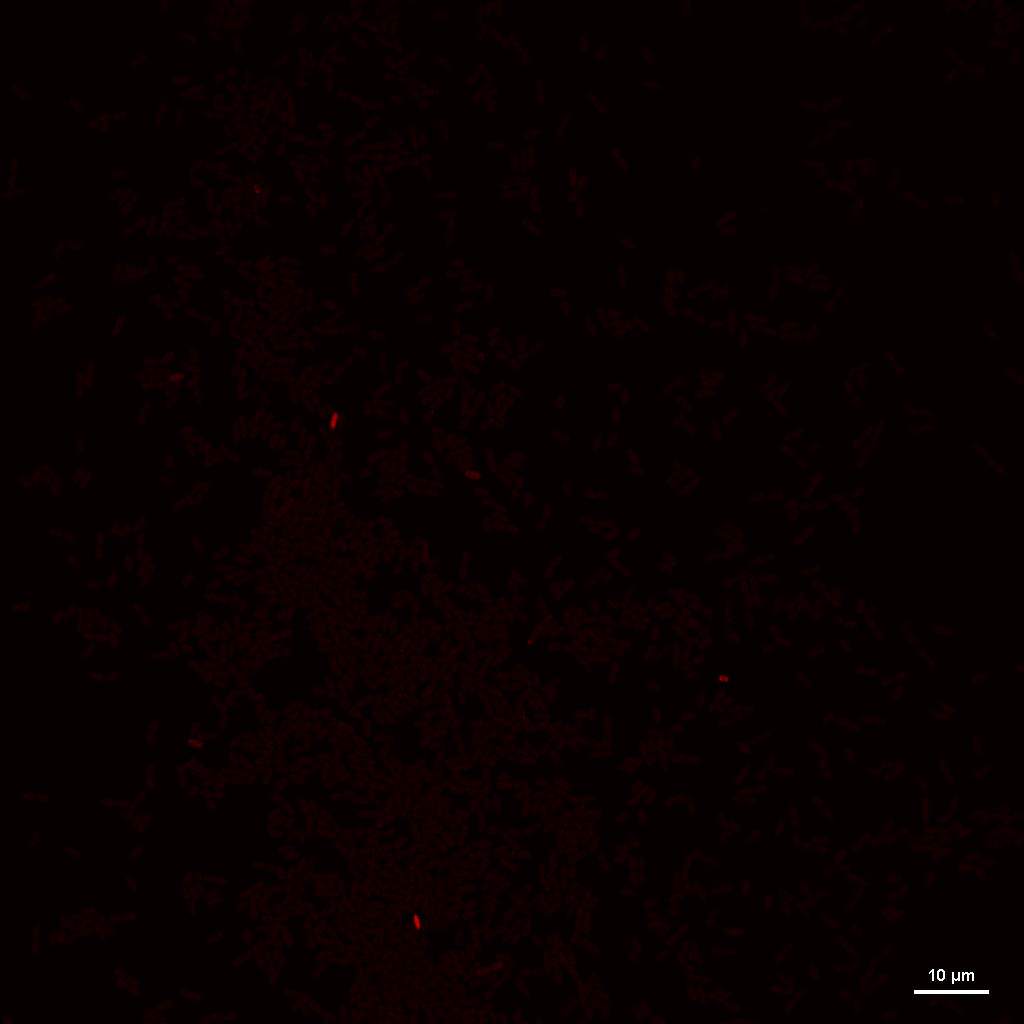

### CSRAKAN_LB_REP2_IMAGE3_RGB_MCHERRY.tif

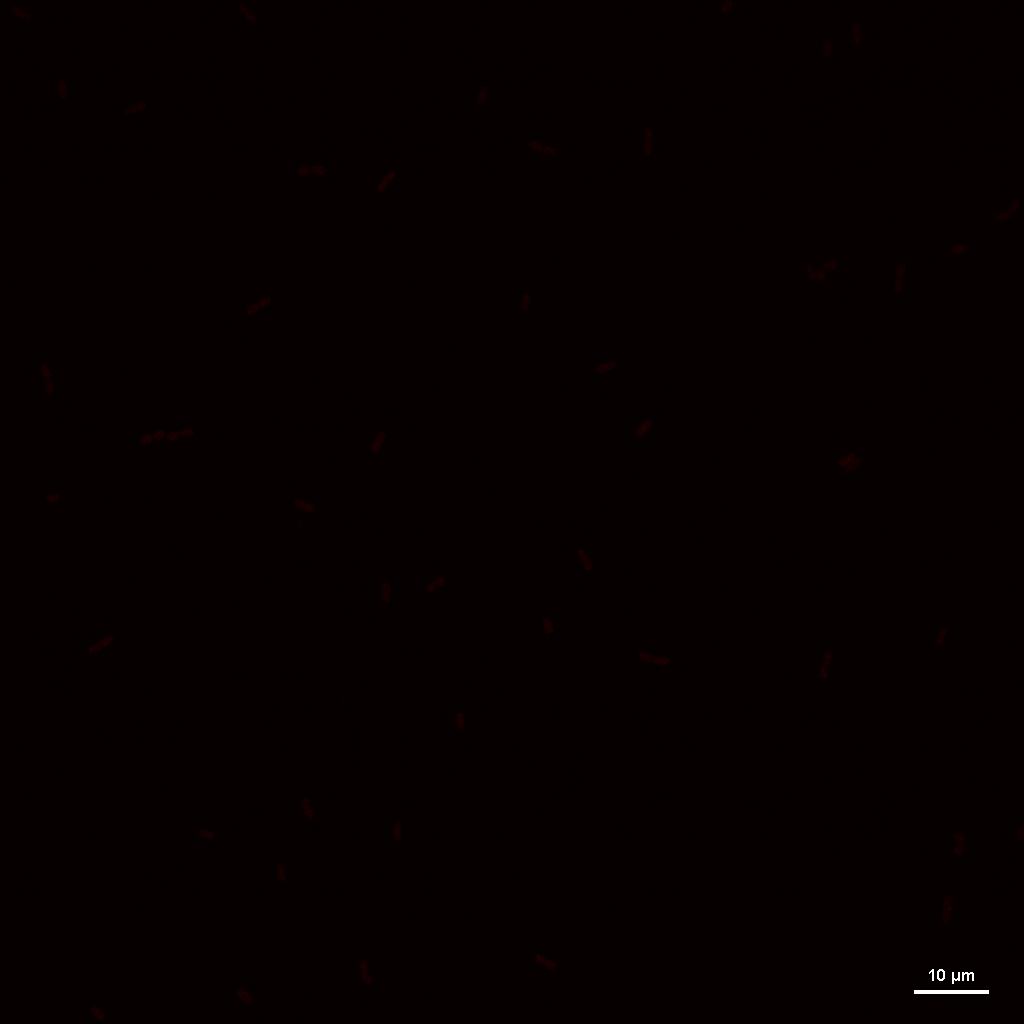

### CSRAKAN_LB_REP2_IMAGE5_RGB_MCHERRY.tif

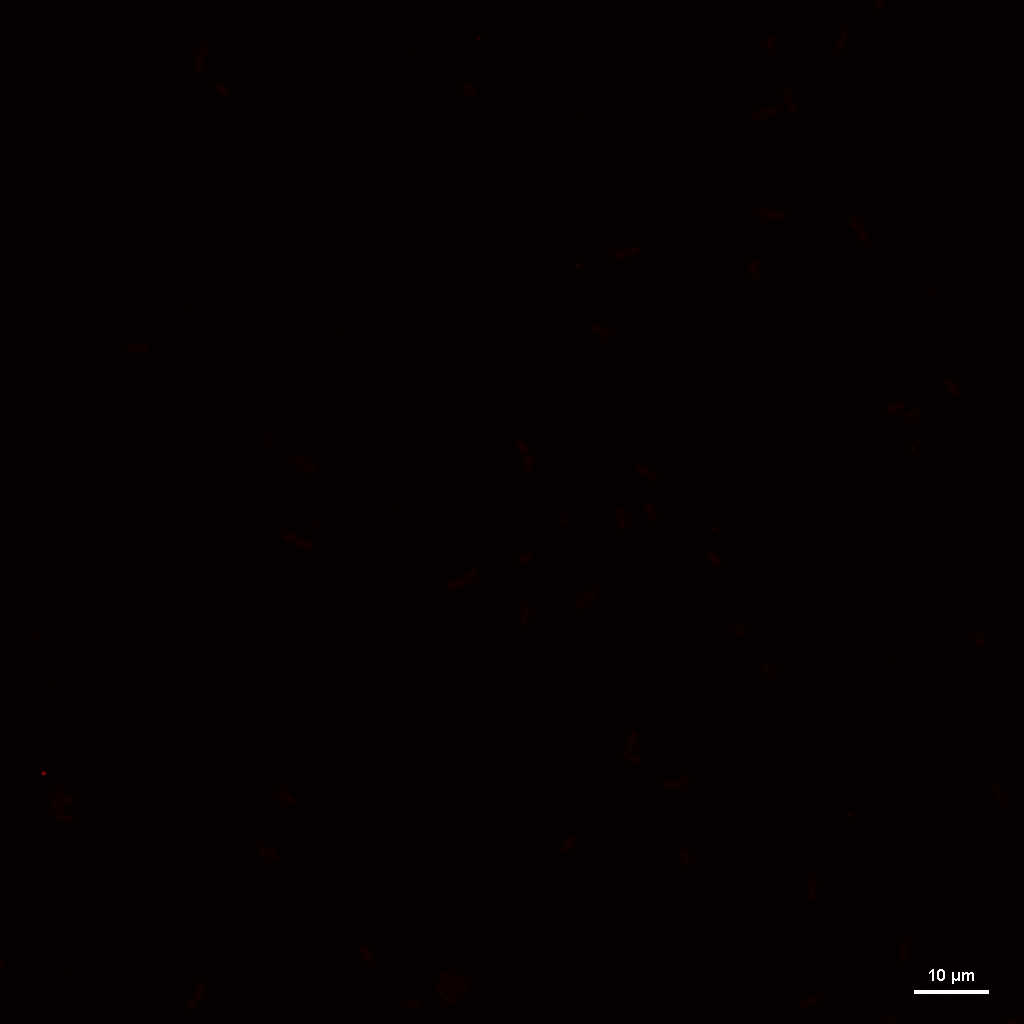

### CSRAKAN_LB_REP3_IMAGE1_RGB_MCHERRY.tif

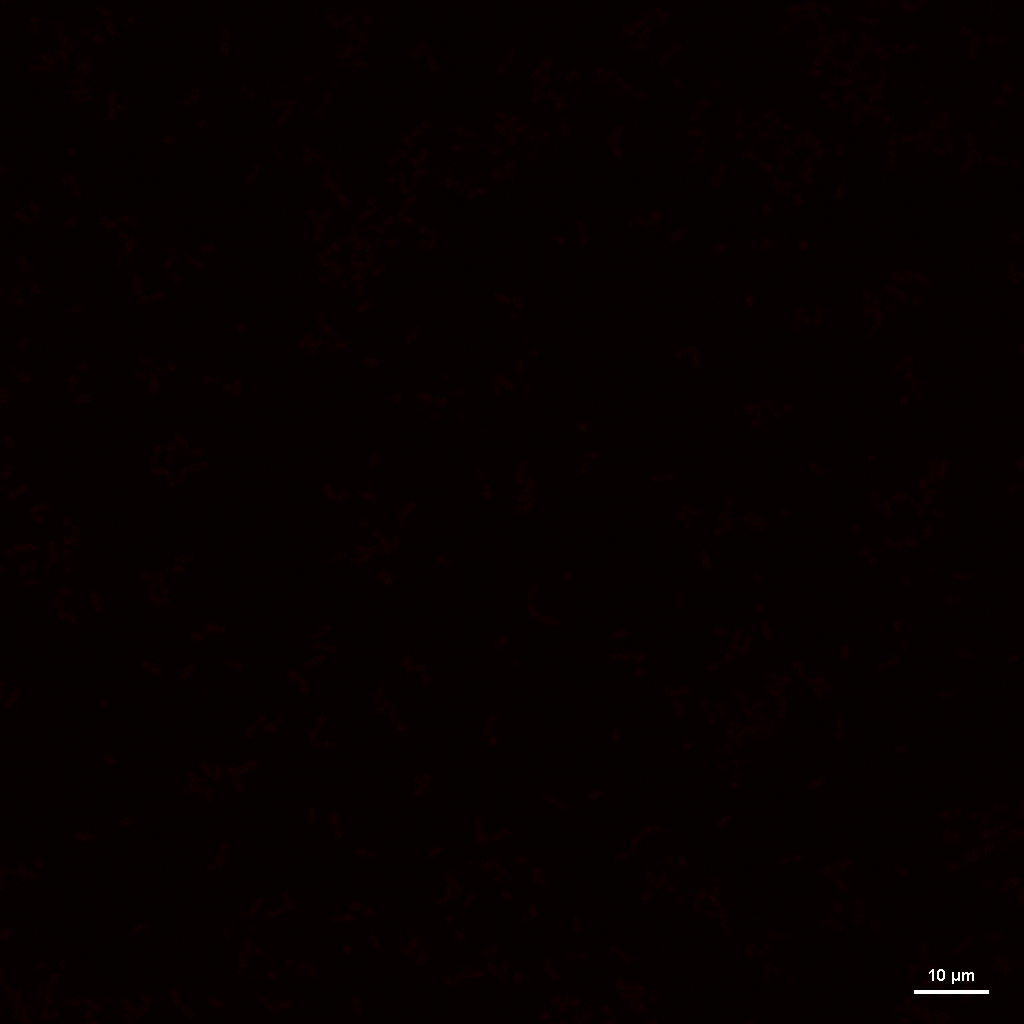

### CSRAKAN_LB_REP3_IMAGE2_RGB_MCHERRY.tif

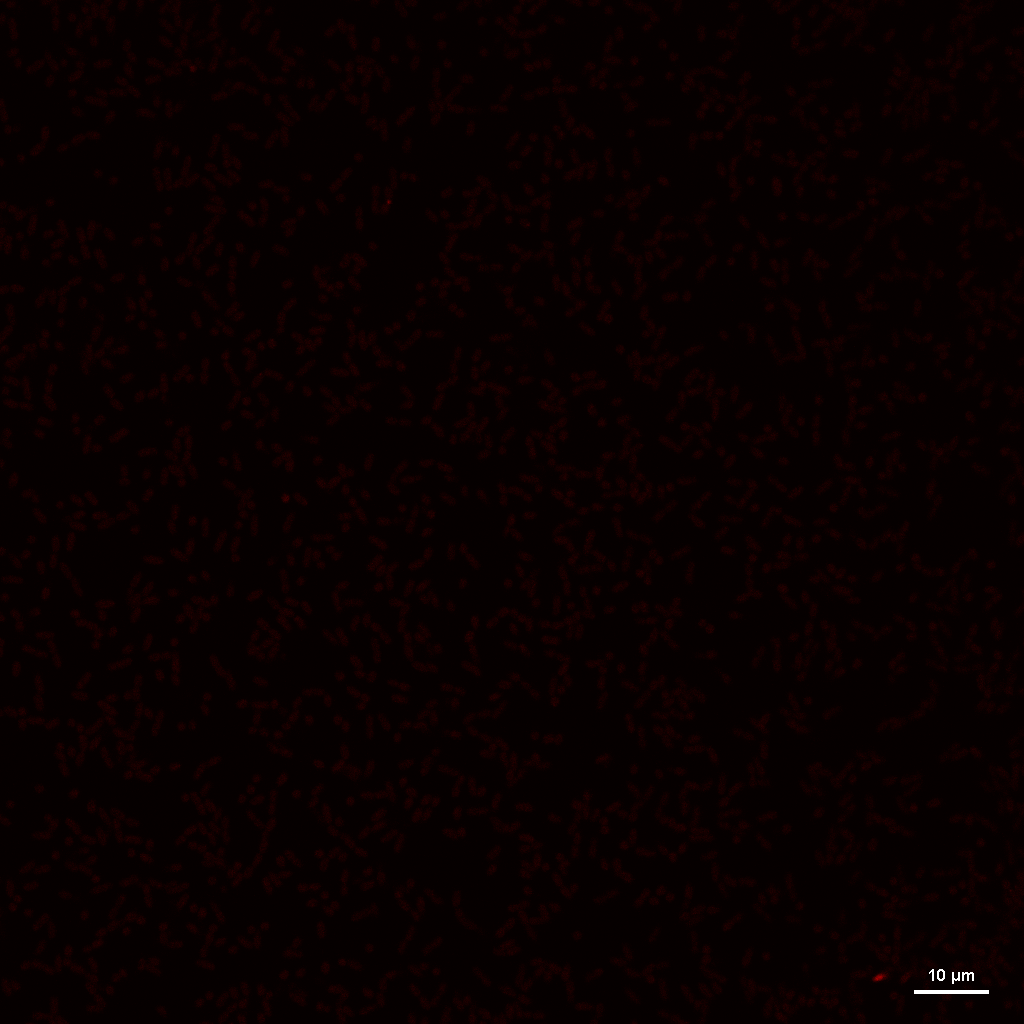

### CSRAKAN_N+24_REP1_IMAGE1_RGB_MCHERRY.tif

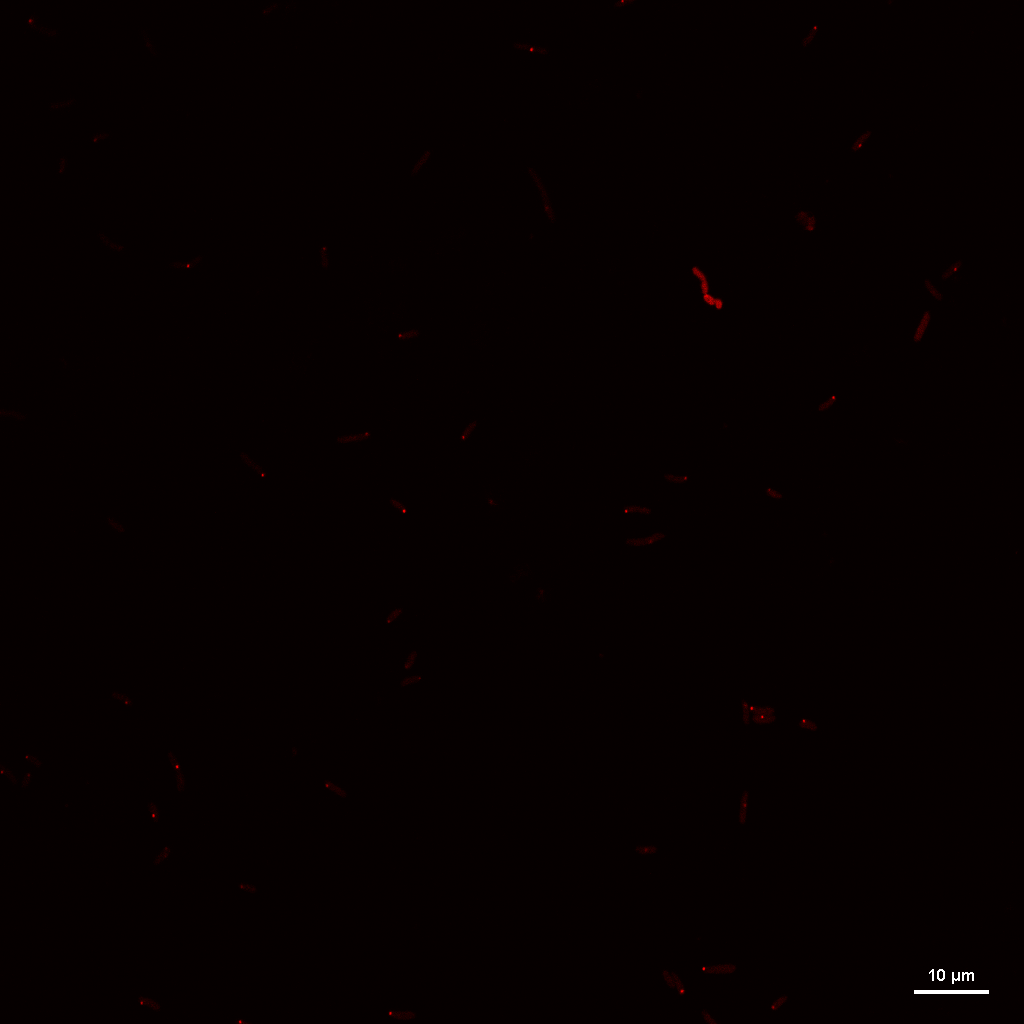

### CSRAKAN_N+24_REP1_IMAGE2_RGB_MCHERRY.tif

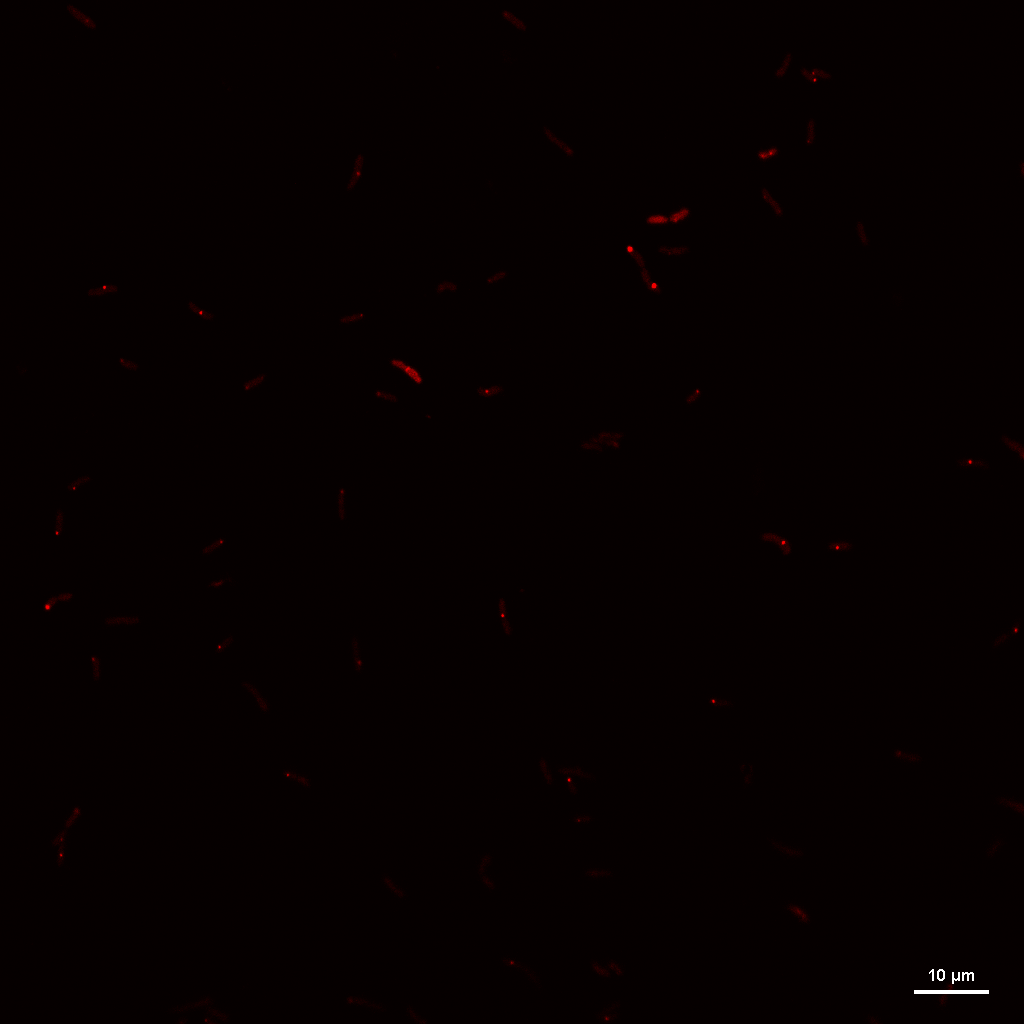

### CSRAKAN_N+24_REP1_IMAGE3_RGB_MCHERRY.tif

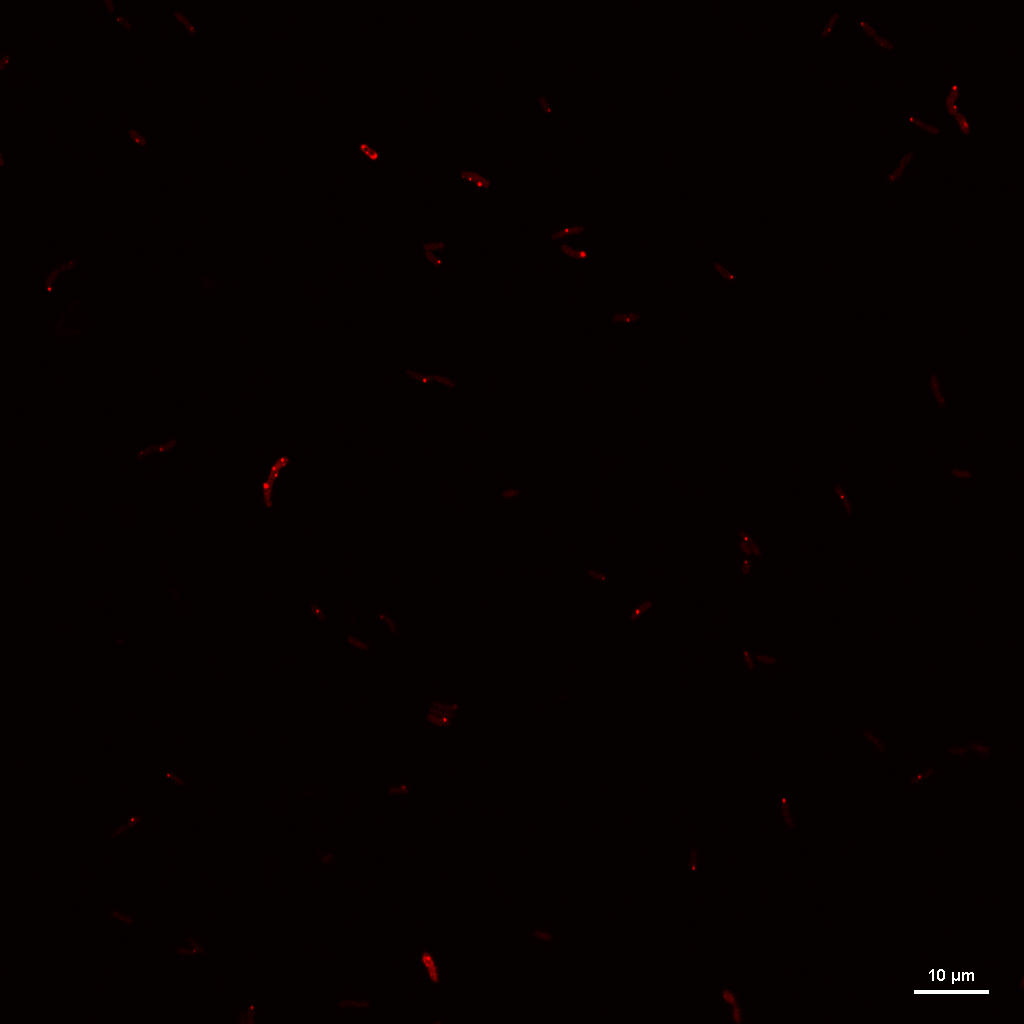

### CSRAKAN_N+24_REP1_IMAGE4_RGB_MCHERRY.tif

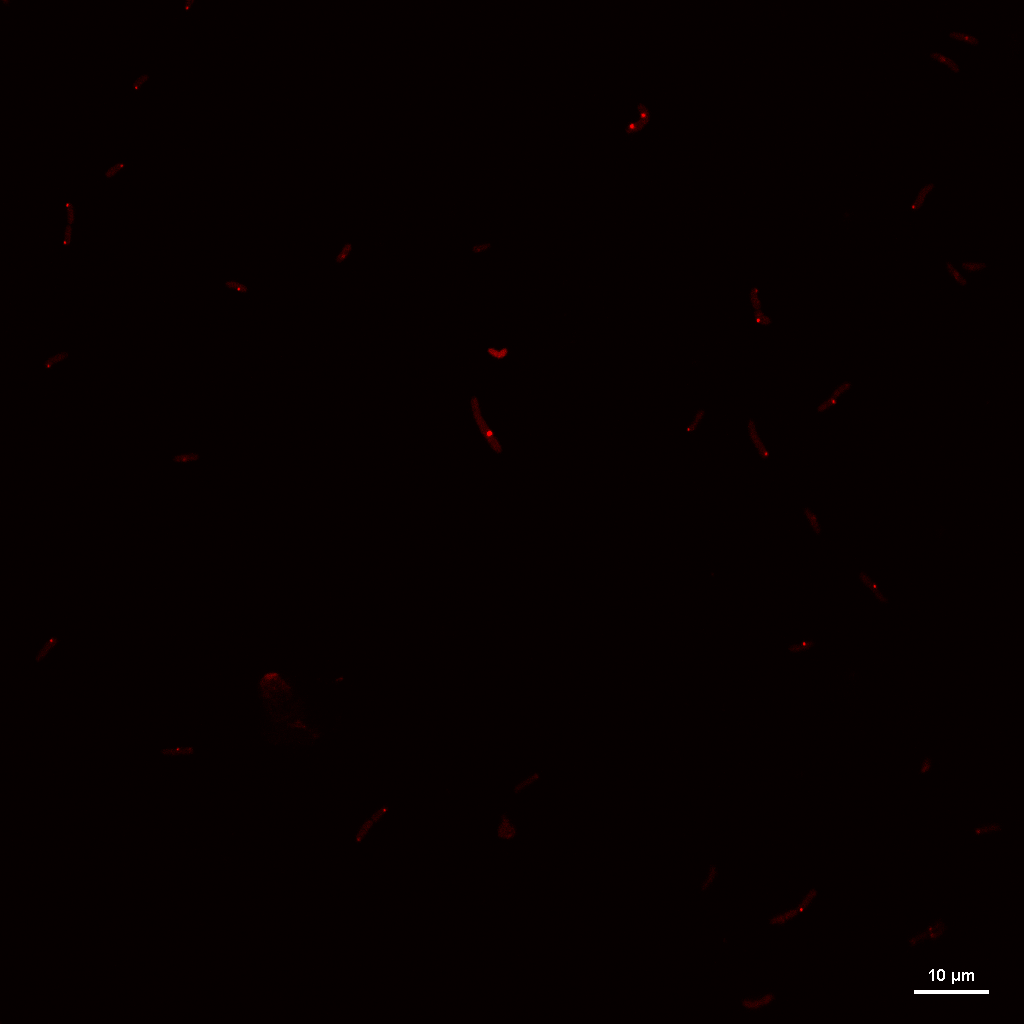

### CSRAKAN_N+24_REP1_IMAGE5_RGB_MCHERRY.tif

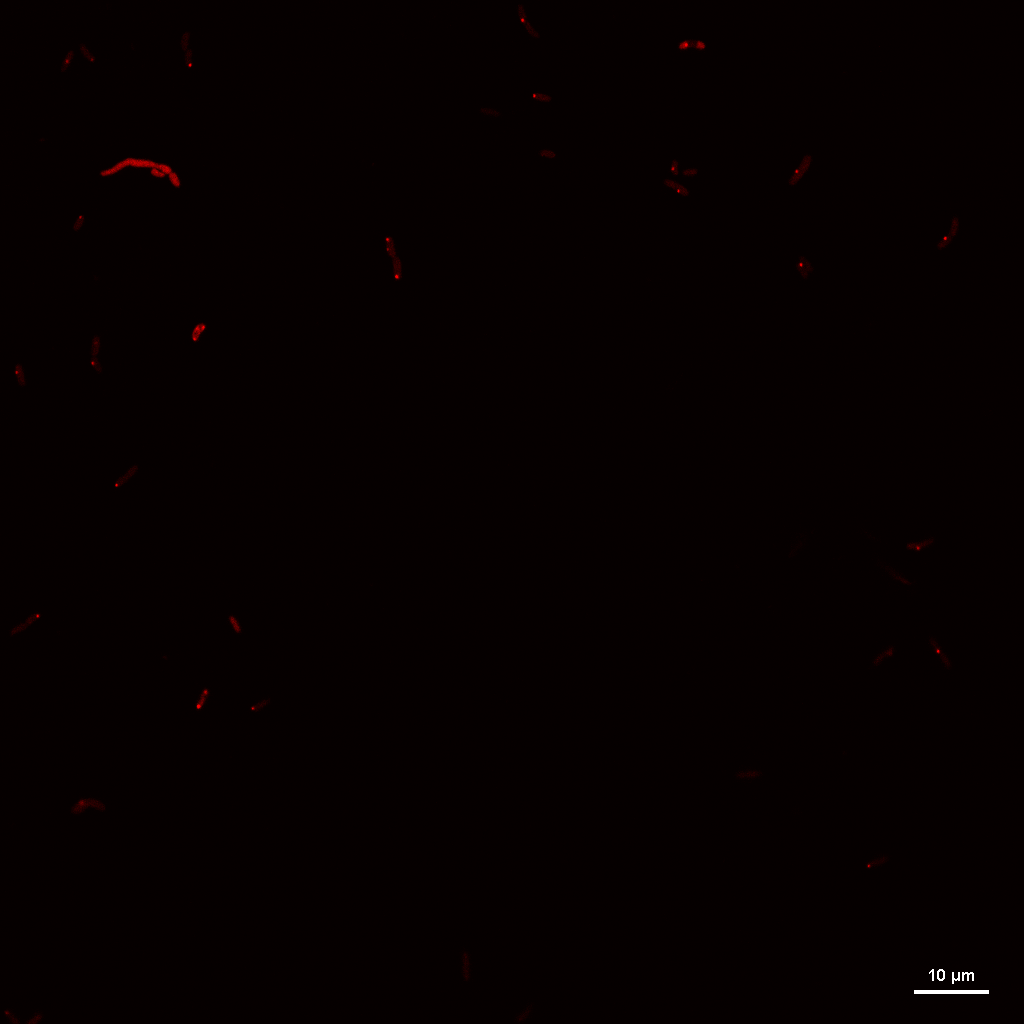

### CSRAKAN_N+24_REP2_IMAGE1_RGB_MCHERRY.tif

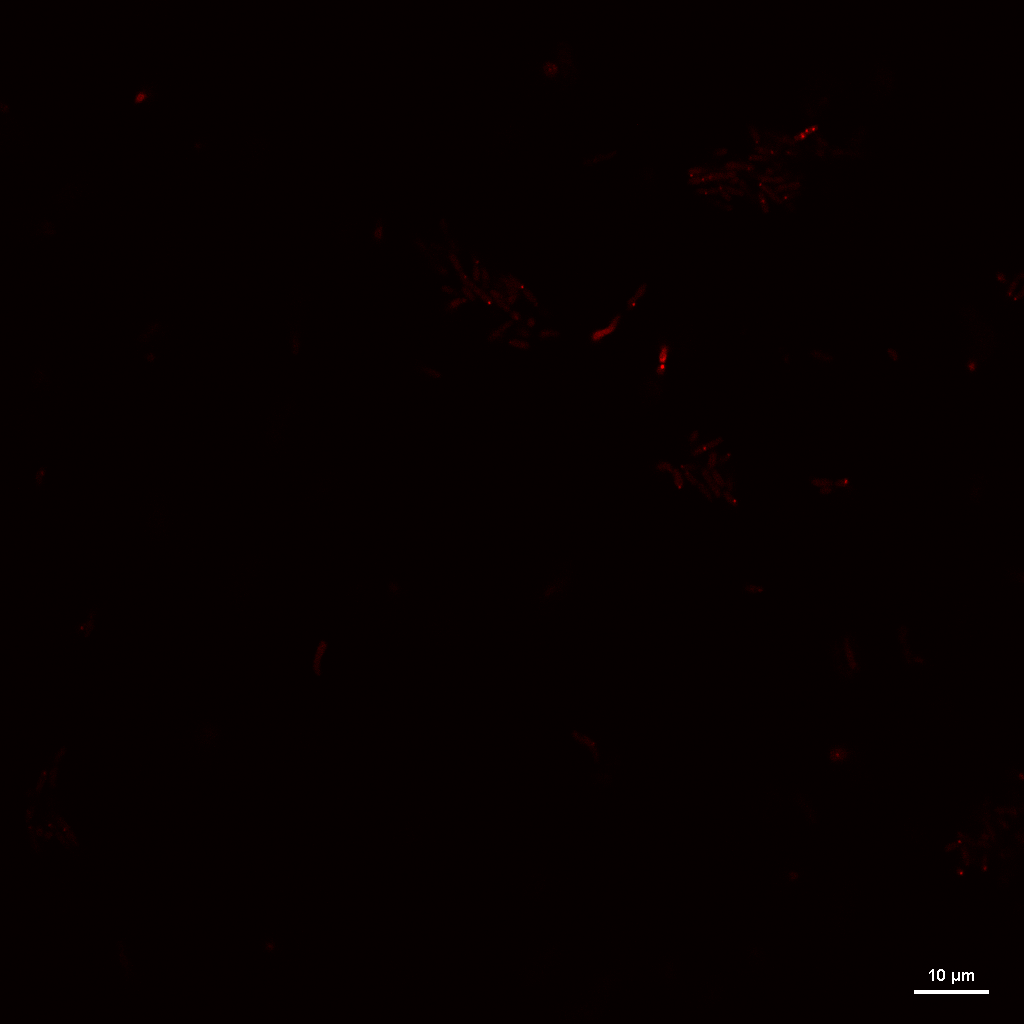

### CSRAKAN_N+24_REP2_IMAGE2_RGB_MCHERRY.tif

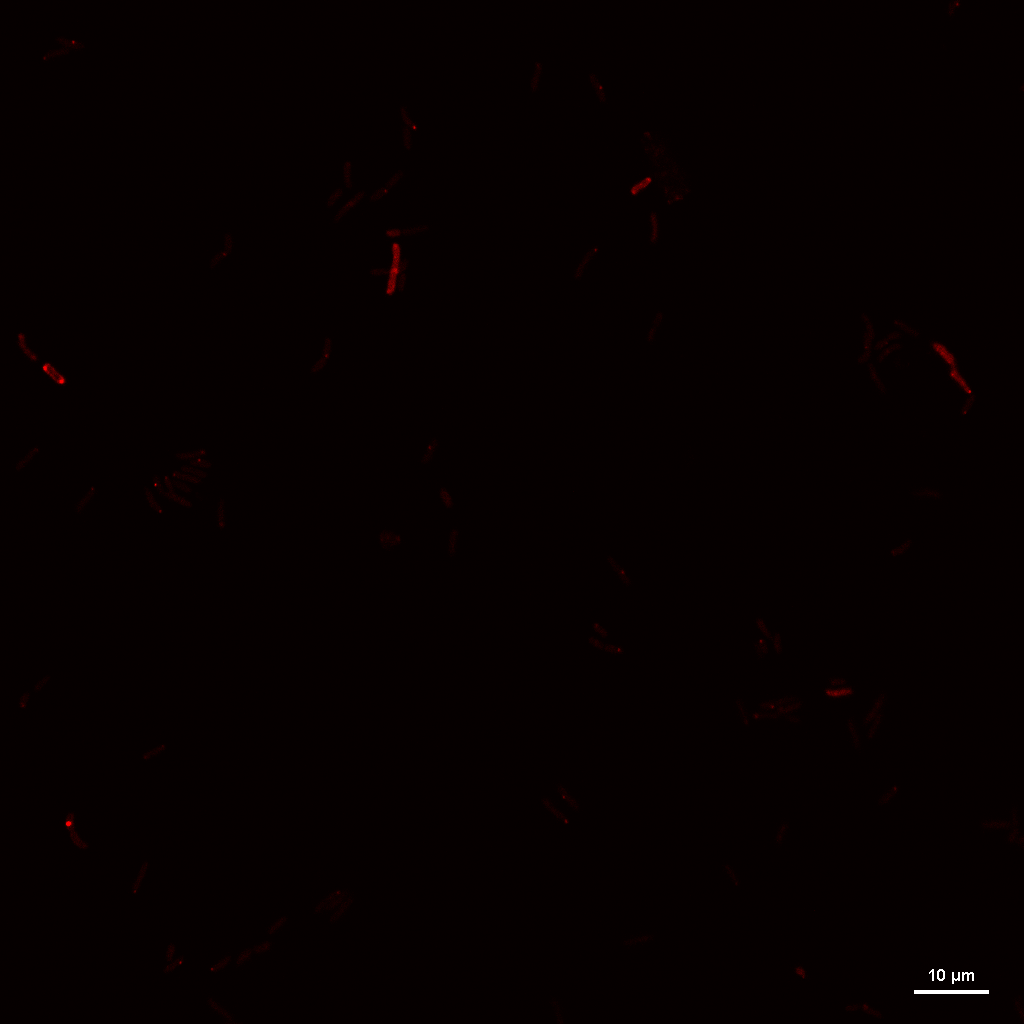

### CSRAKAN_N+24_REP2_IMAGE3_RGB_MCHERRY.tif

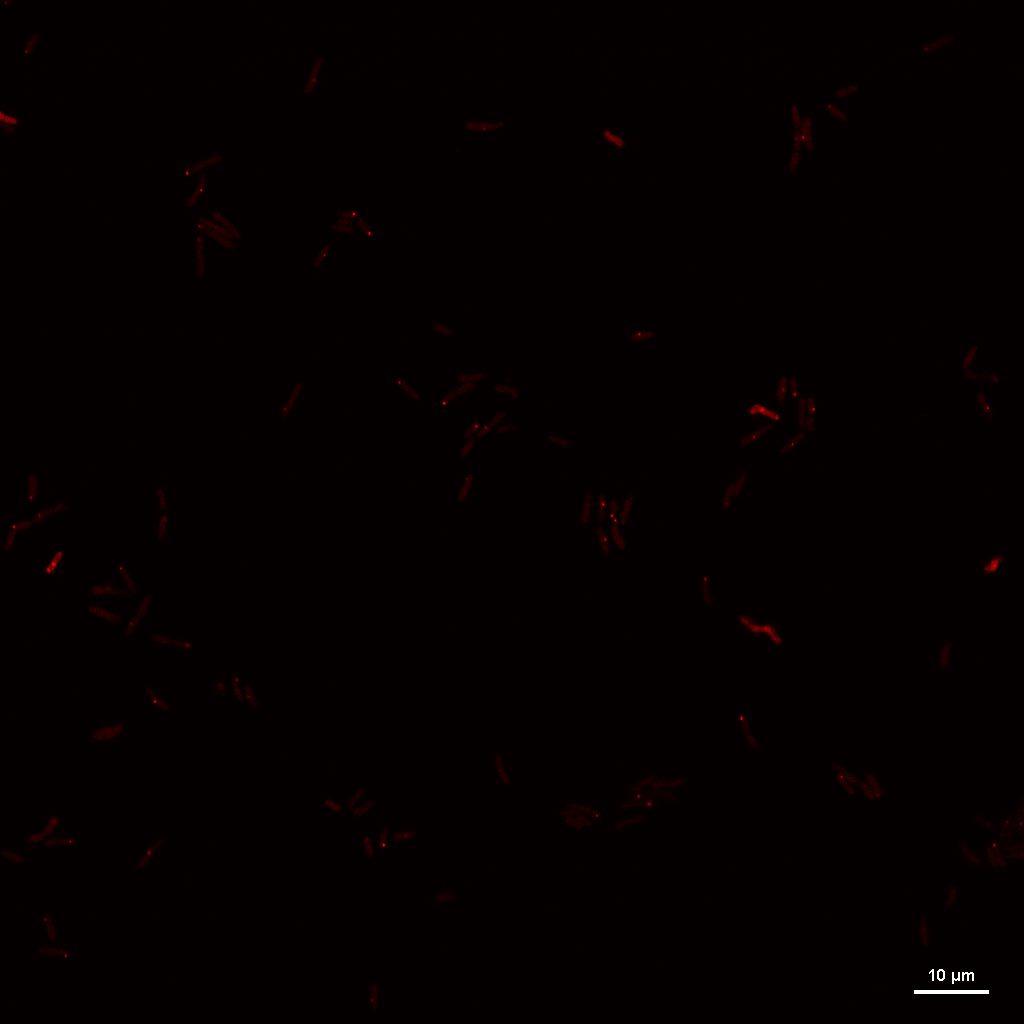

### CSRAKAN_N+24_REP3_IMAGE4_RGB_MCHERRY.tif

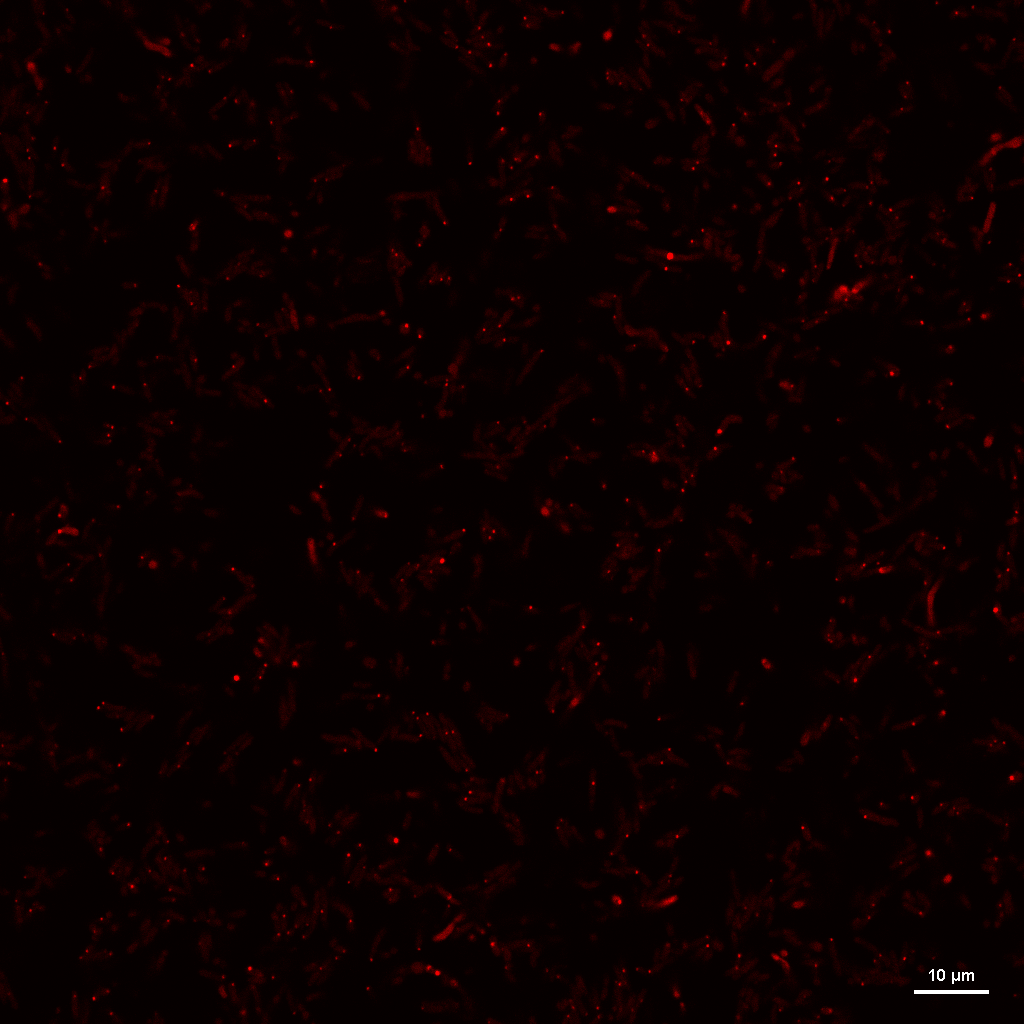

### CSRAKAN_N-_REP1_IMAGE1_RGB_MCHERRY.tif

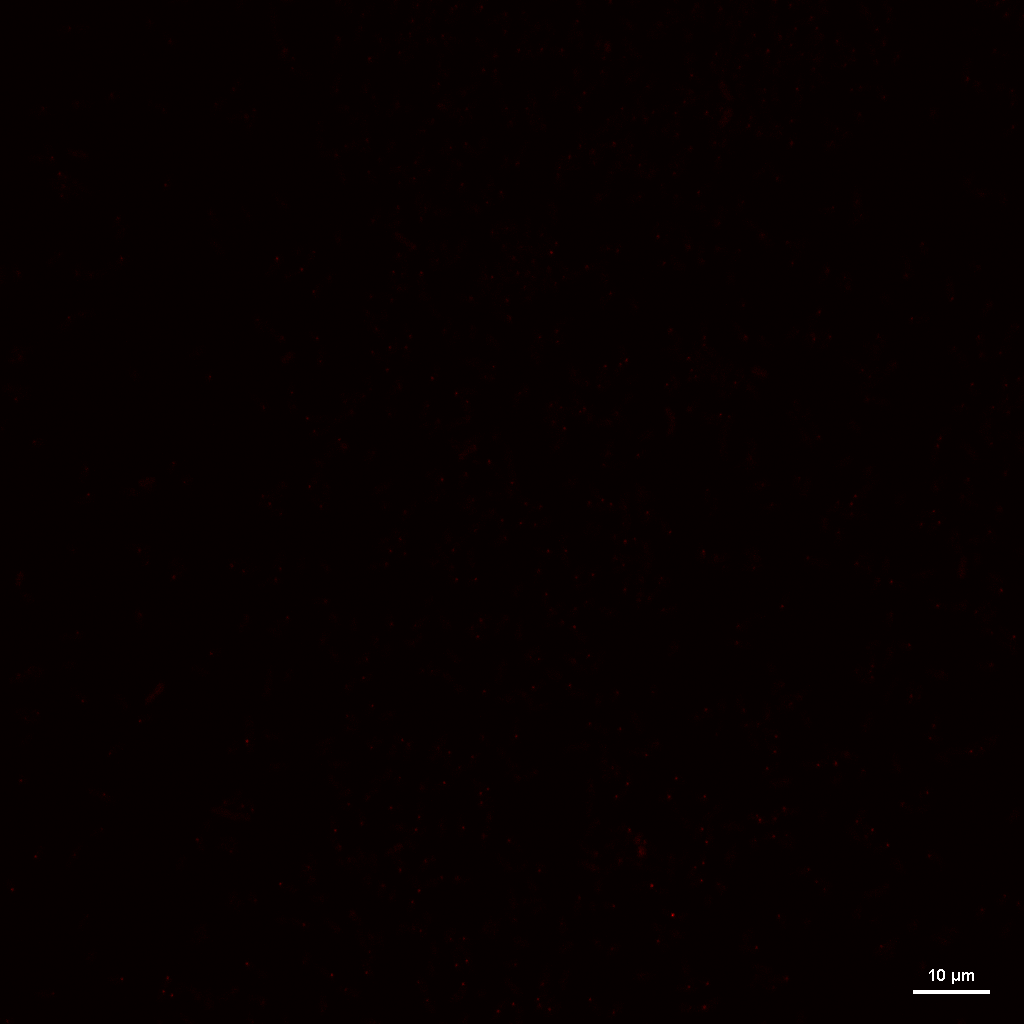

### CSRAKAN_N-_REP1_IMAGE2_RGB_MCHERRY.tif

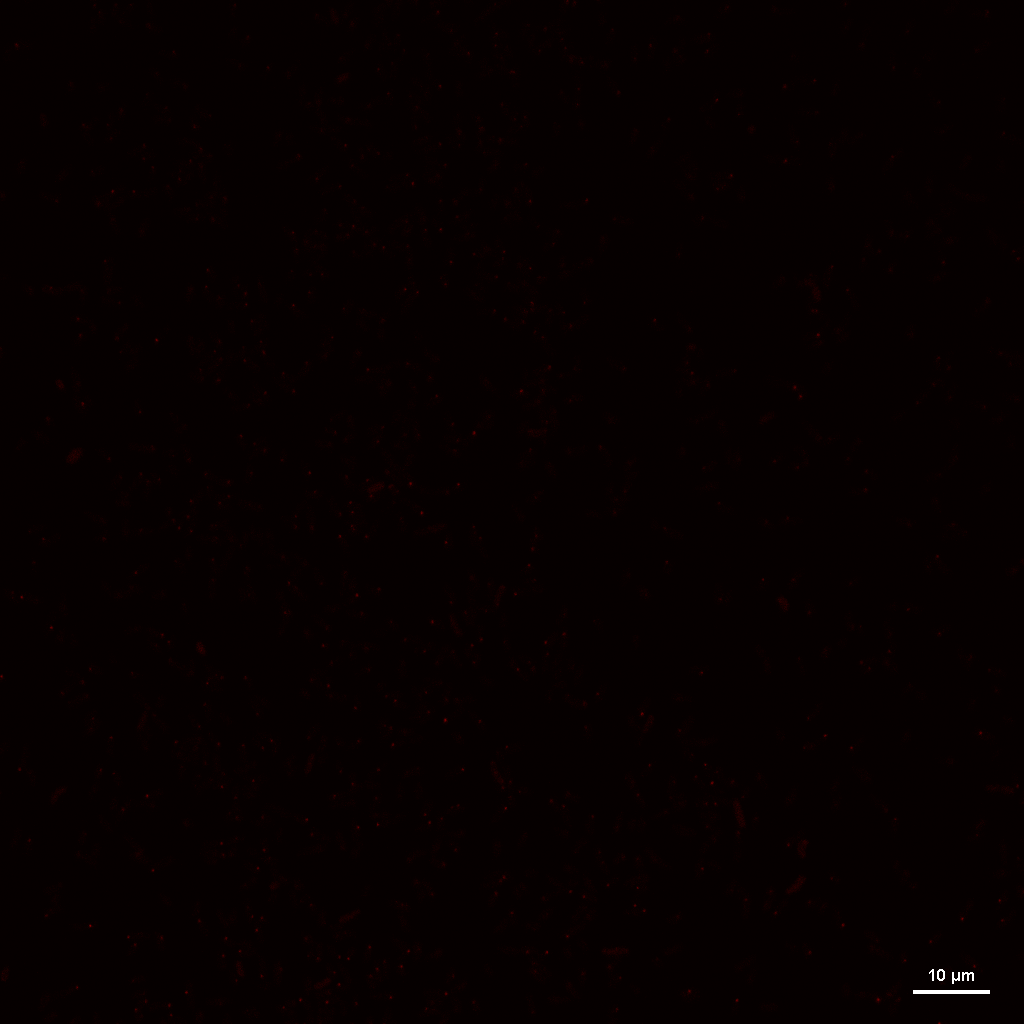

### CSRAKAN_N-_REP2_IMAGE1_RGB_MCHERRY.tif

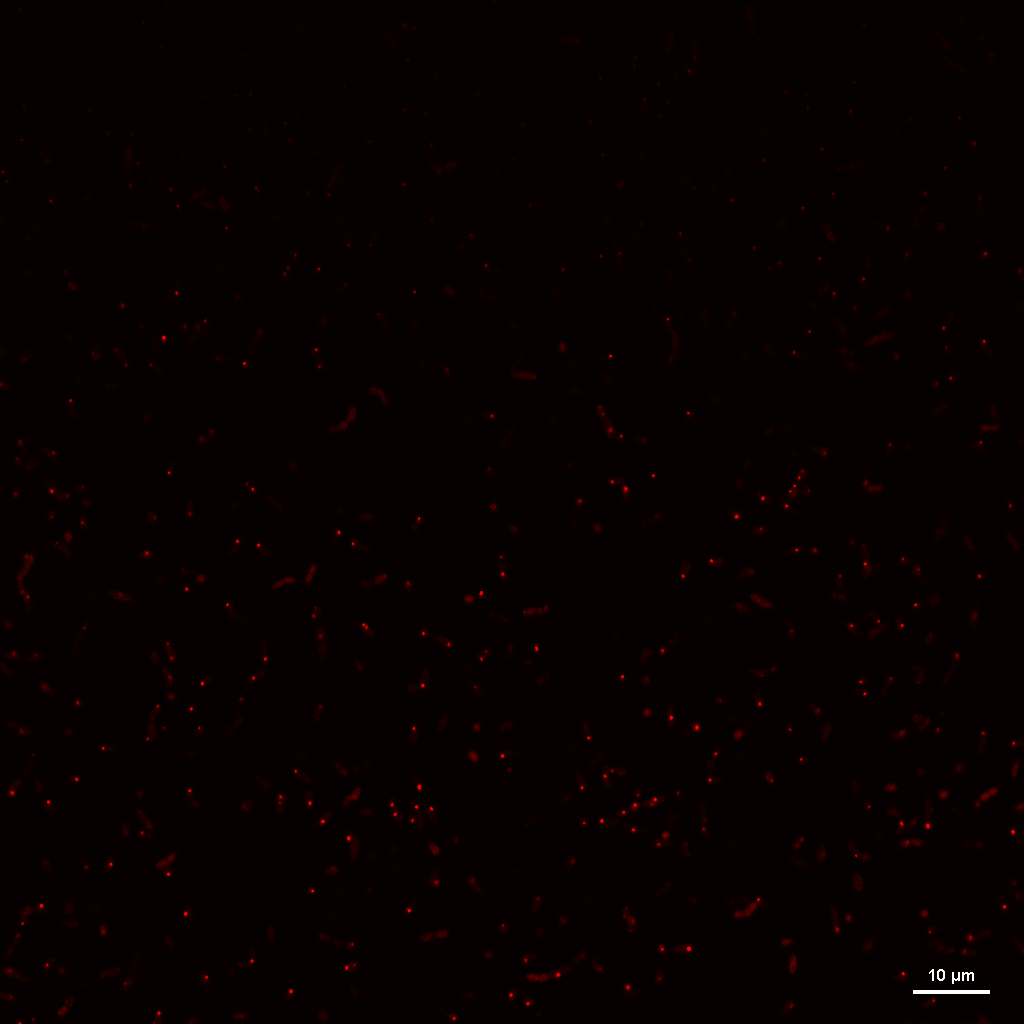

### CSRAKAN_N-_REP3_IMAGE1_RGB_MCHERRY.tif

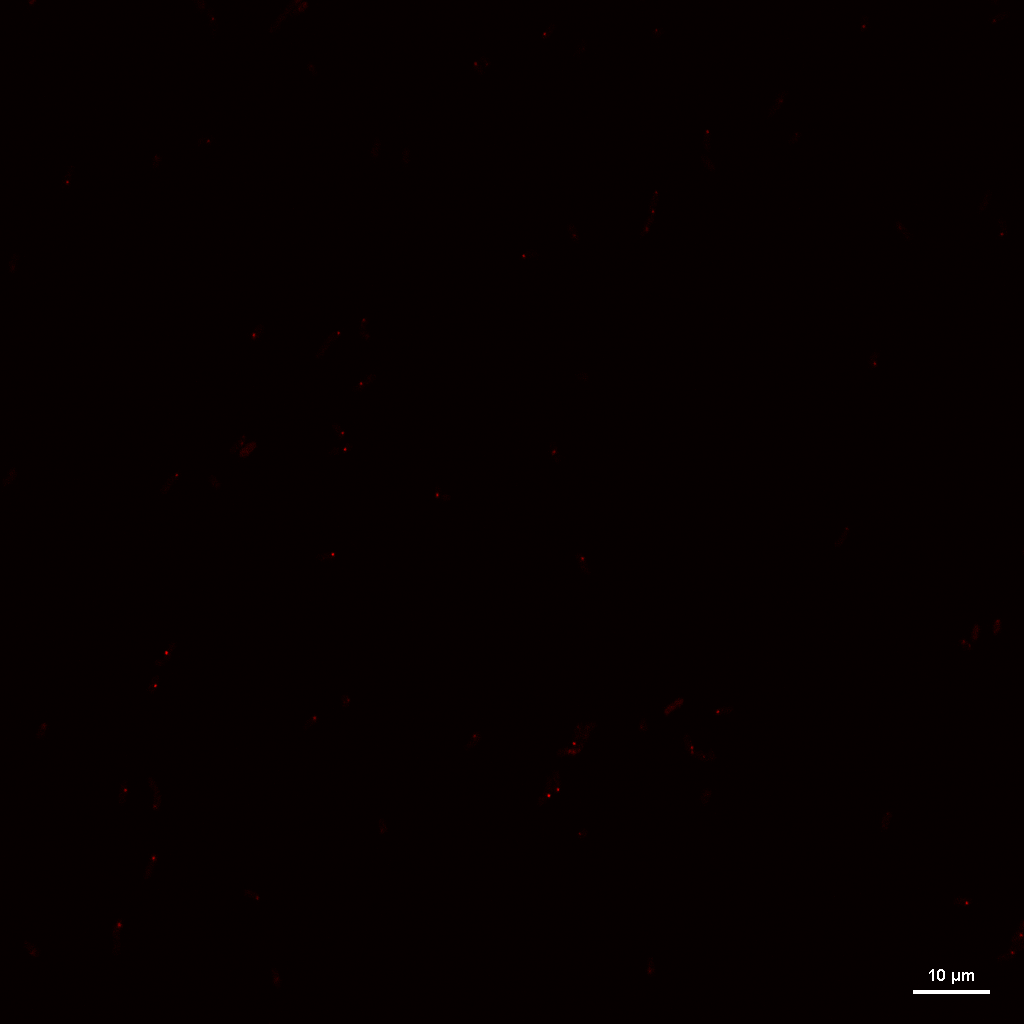

### CSRAKAN_N-_REP3_IMAGE2_RGB_MCHERRY.tif

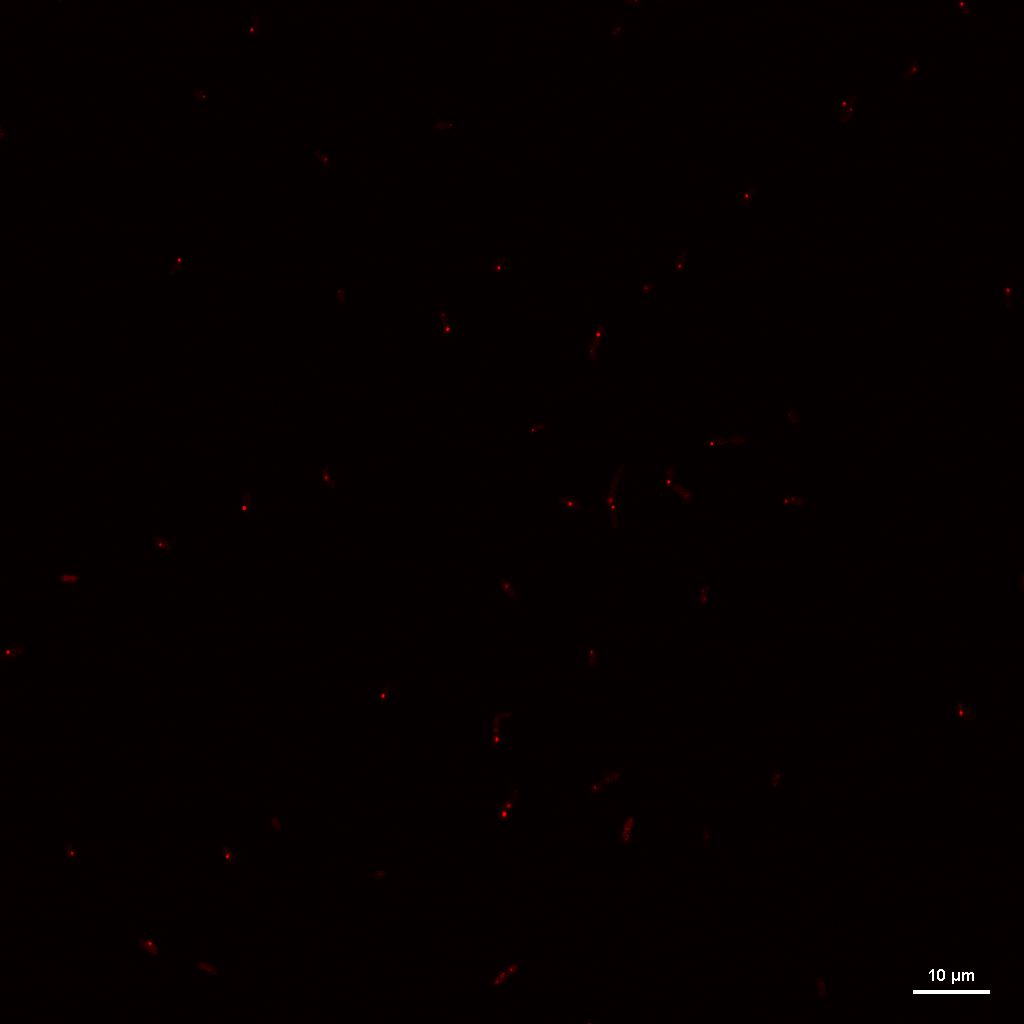

### CSRAKAN_N-_REP3_IMAGE5_RGB_MCHERRY.tif

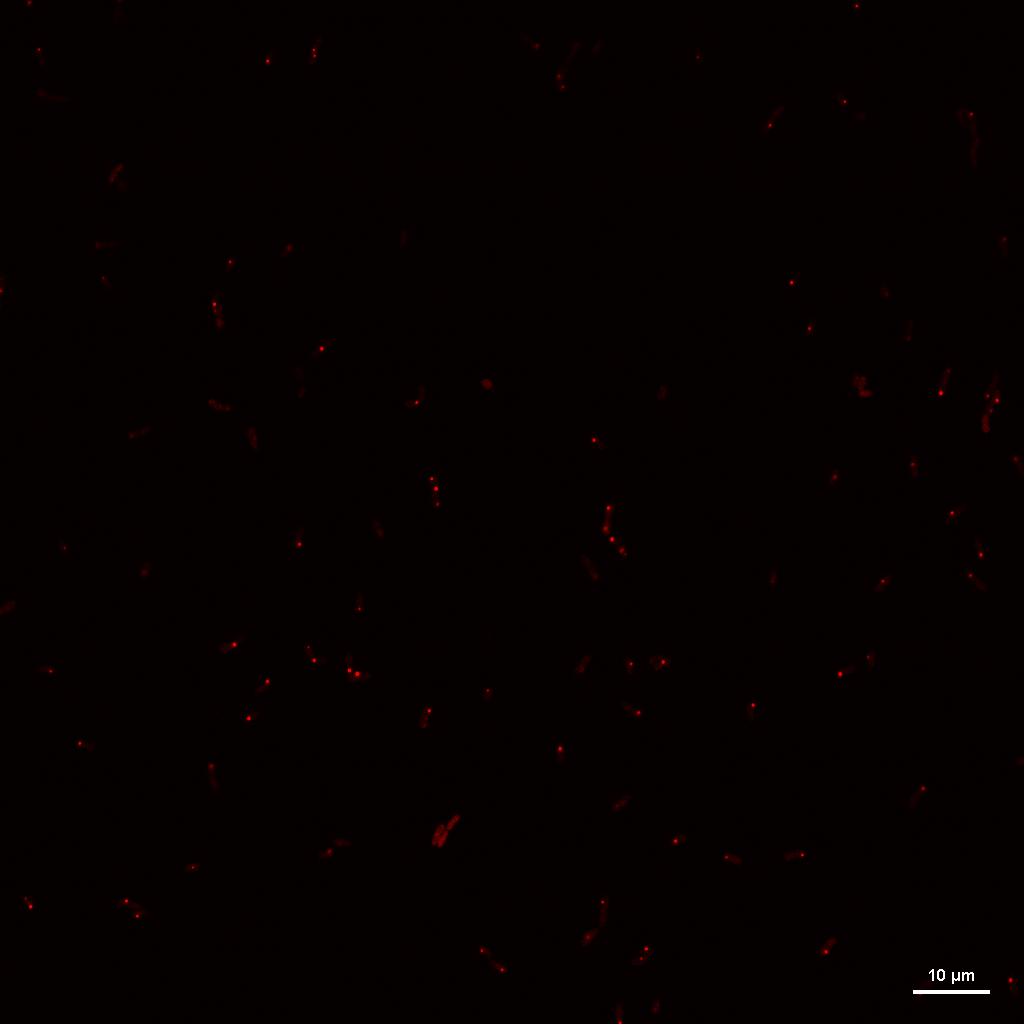

### HMC_LB_REP1_IMAGE1_RGB_MCHERRY.tif

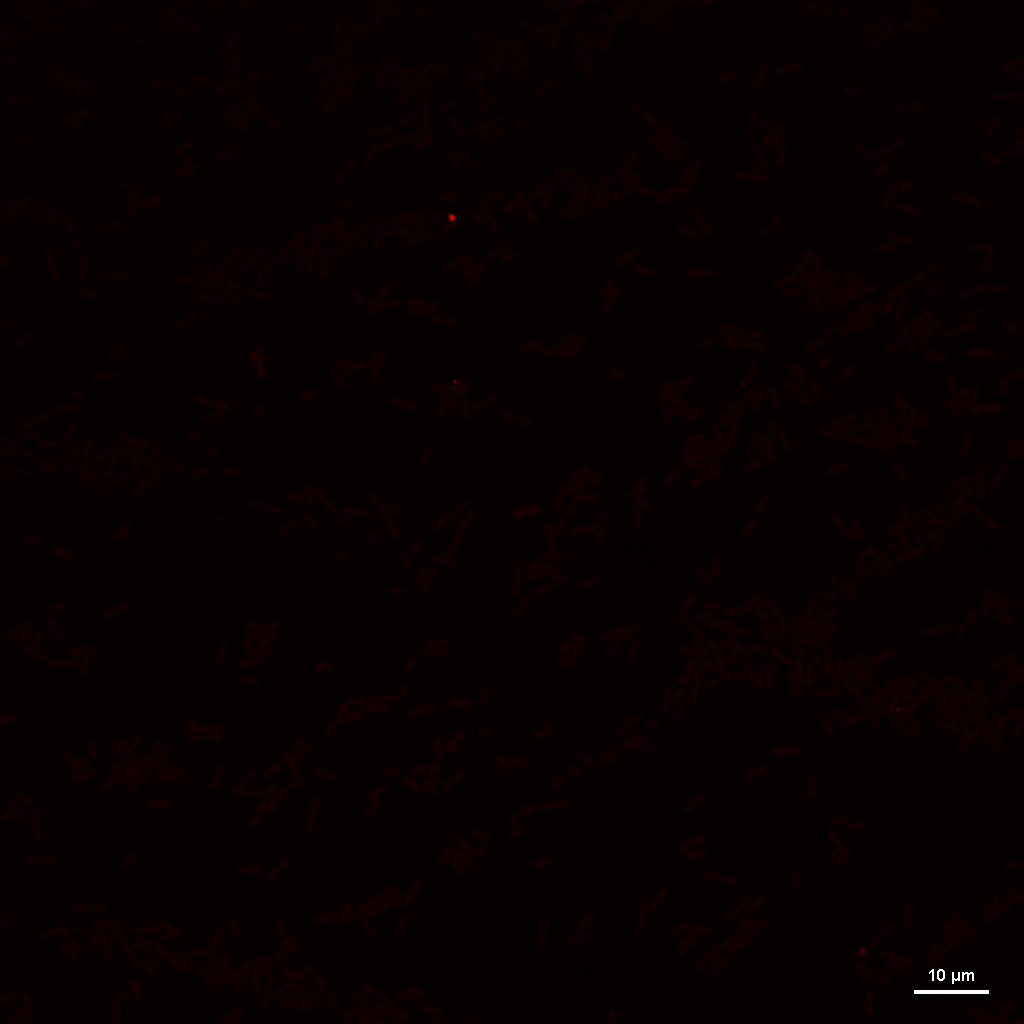

### HMC_LB_REP1_IMAGE2_RGB_MCHERRY.tif

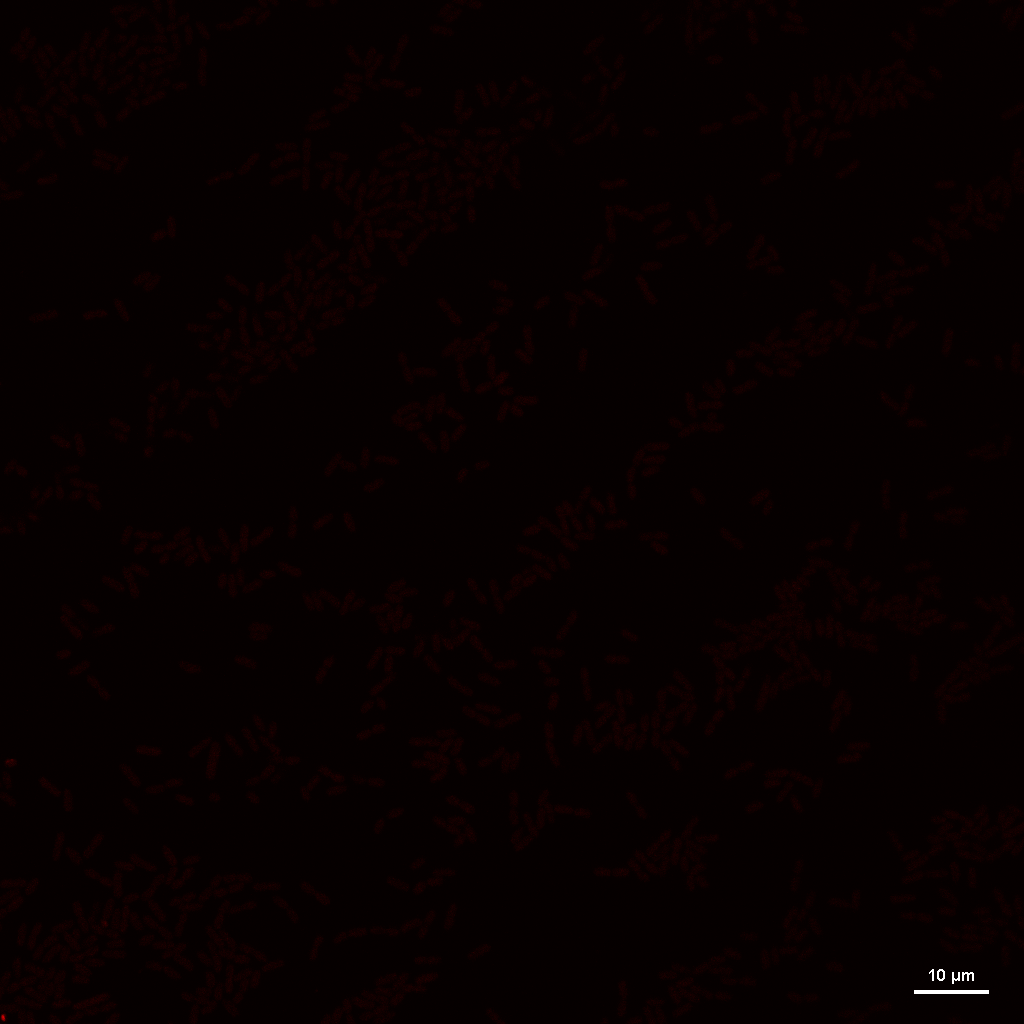

### HMC_LB_REP1_IMAGE3_RGB_MCHERRY.tif

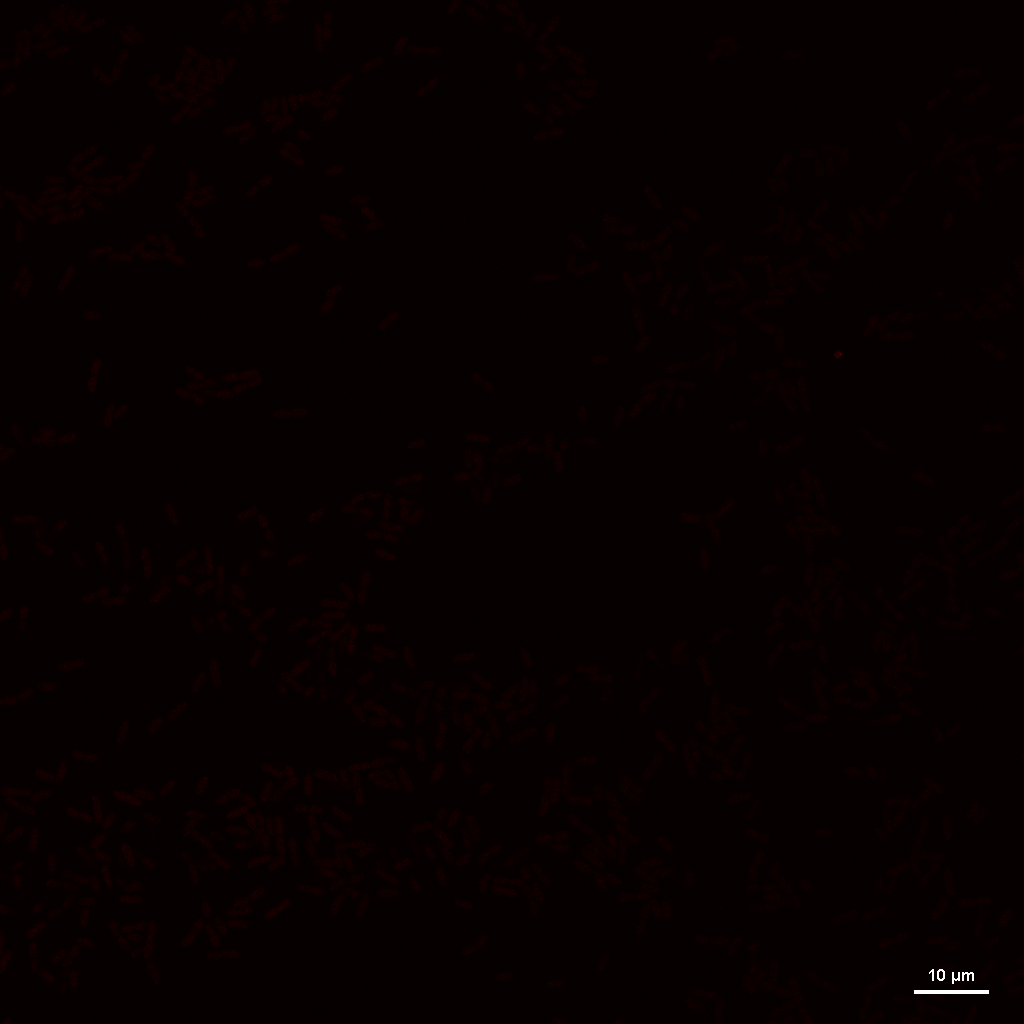

### HMC_LB_REP2_IMAGE1_RGB_MCHERRY.tif

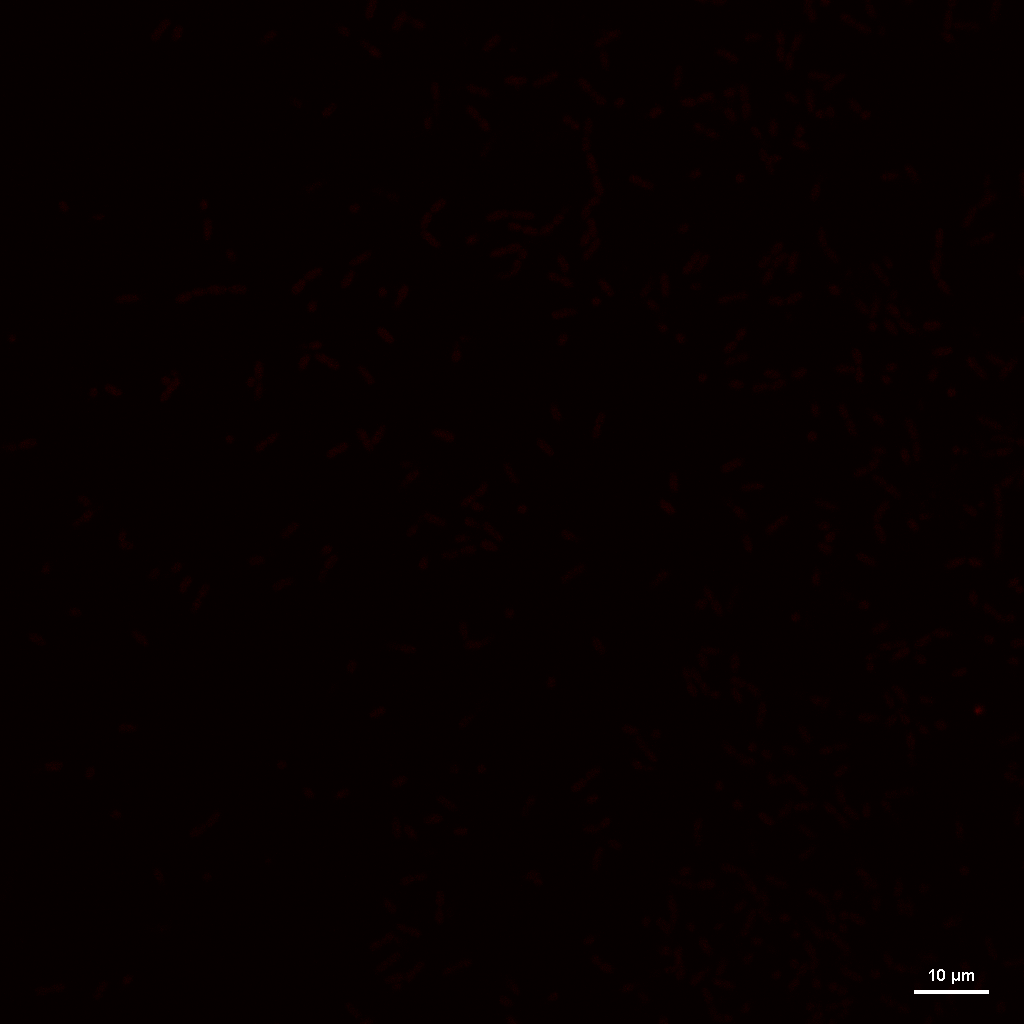

### HMC_LB_REP2_IMAGE2_RGB_MCHERRY.tif

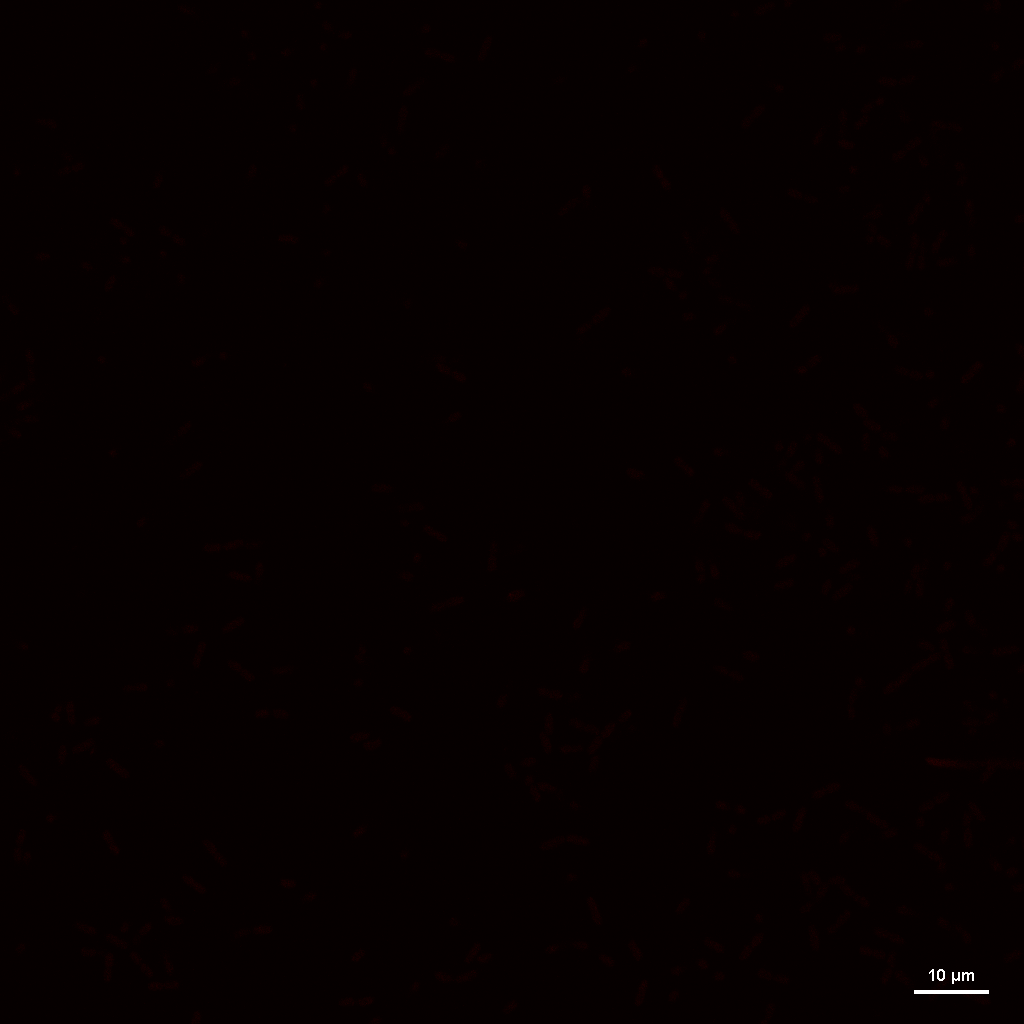

### HMC_LB_REP3_IMAGE3_RGB_MCHERRY.tif

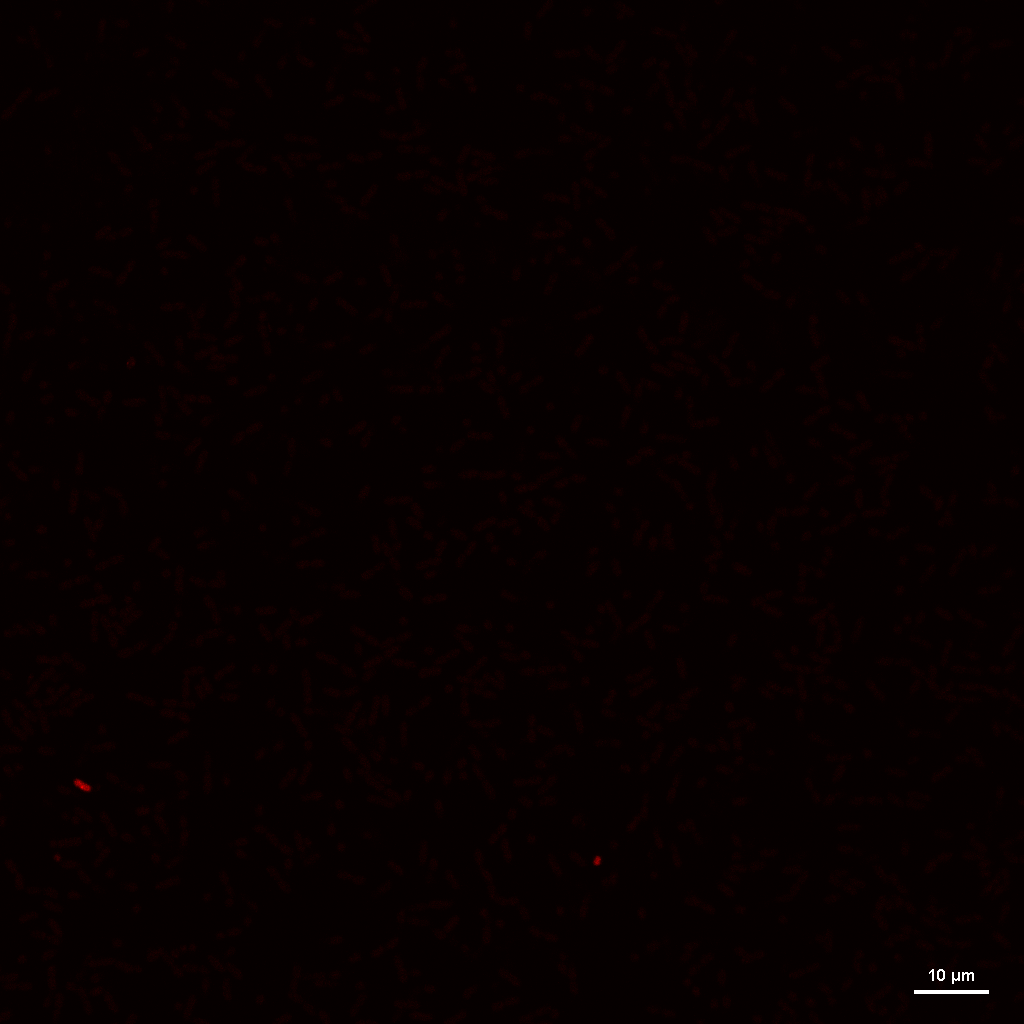

### HMC_LB_REP3_IMAGE4_RGB_MCHERRY.tif

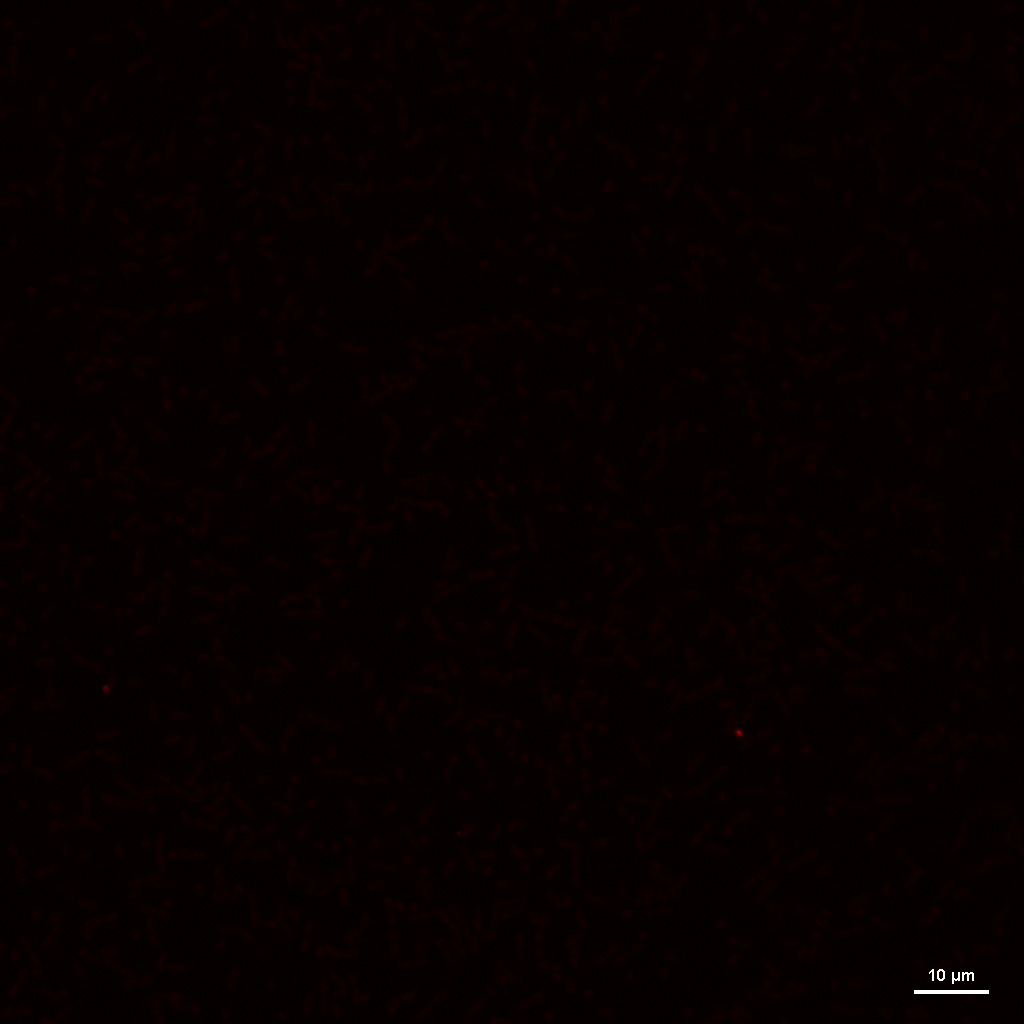

### HMC_N+24_REP1_IMAGE6_RGB_MCHERRY.tif

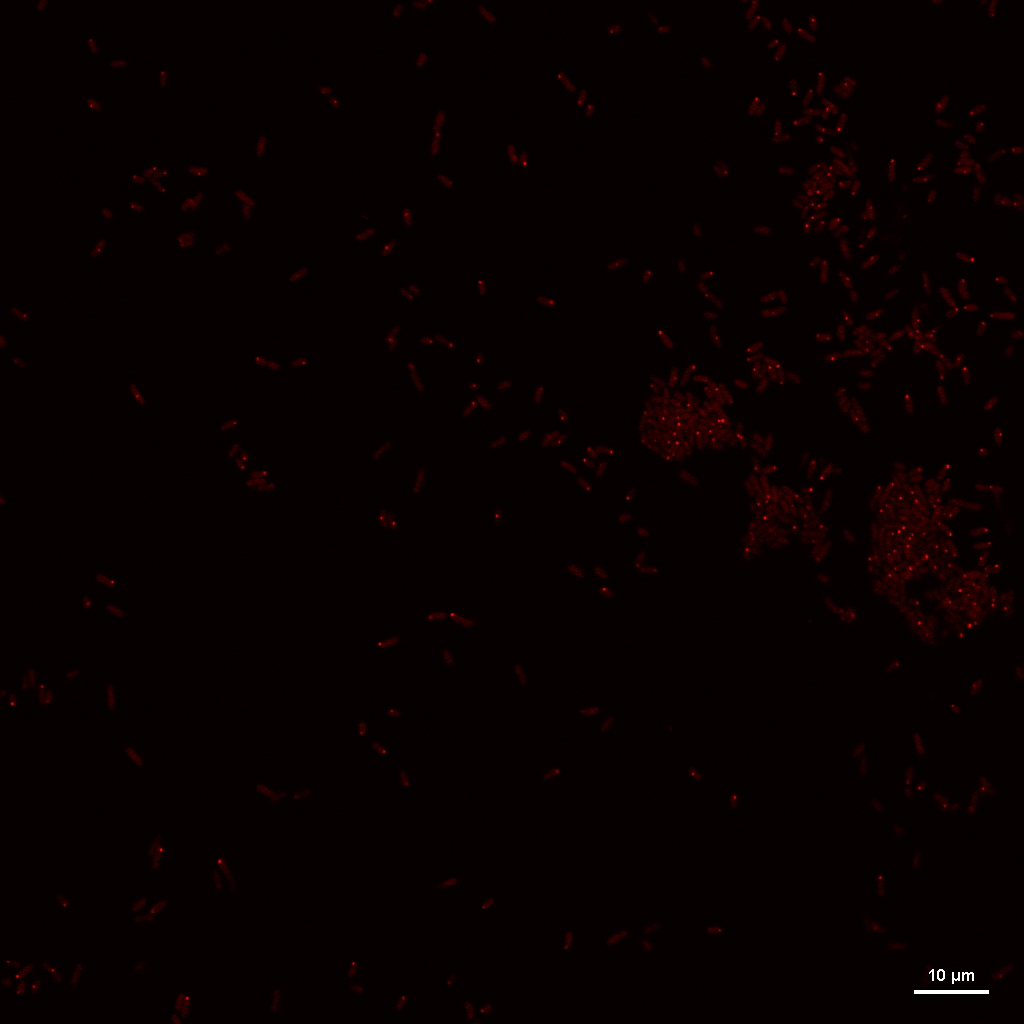

### HMC_N+24_REP2_IMAGE2_RGB_MCHERRY.tif

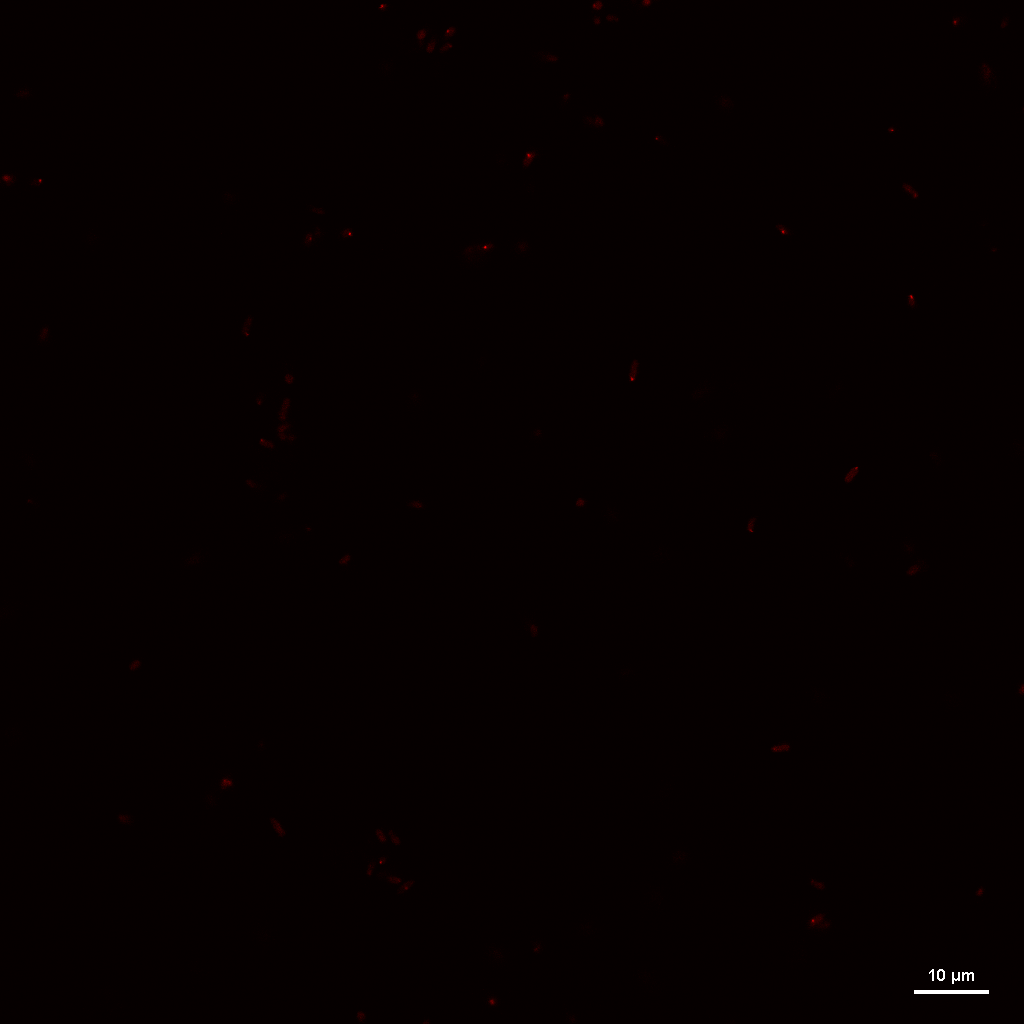

### HMC_N+24_REP2_IMAGE3_RGB_MCHERRY.tif

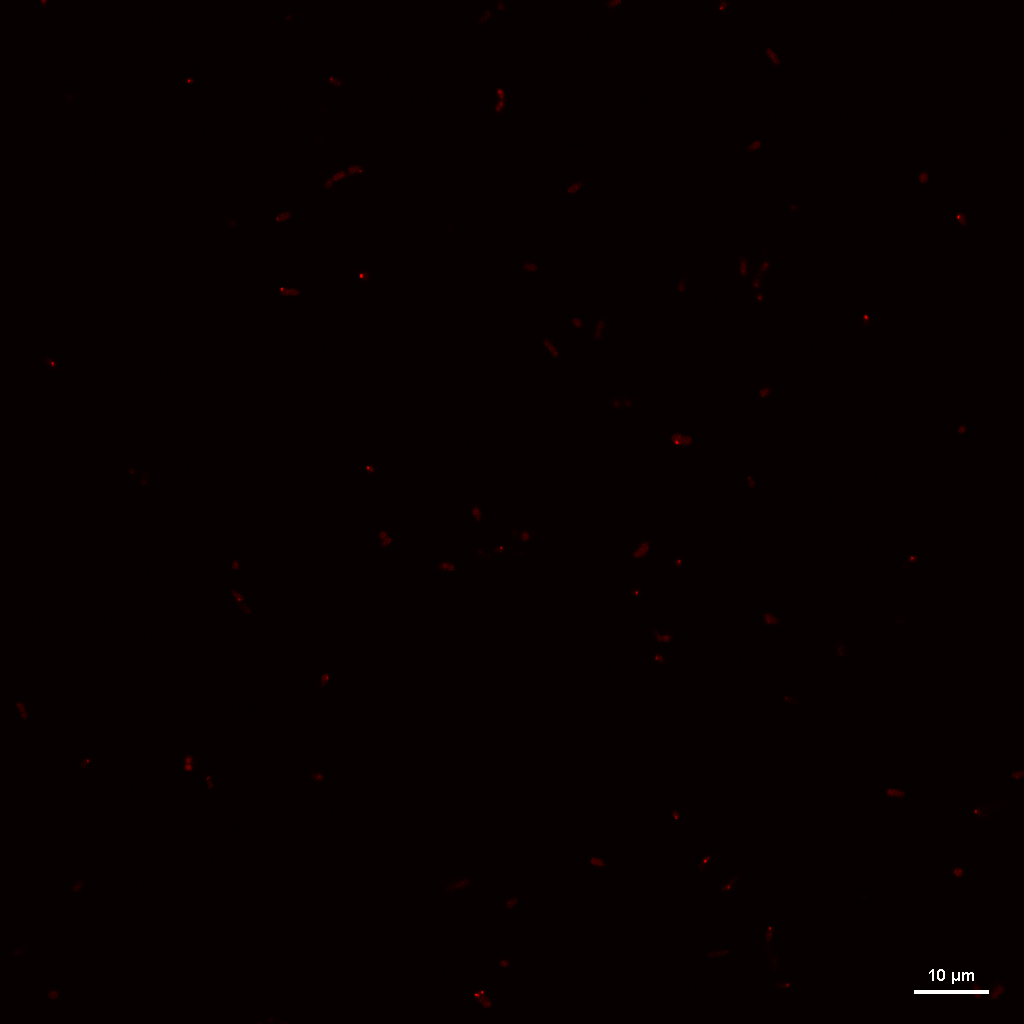

### HMC_N+24_REP2_IMAGE4_RGB_MCHERRY.tif

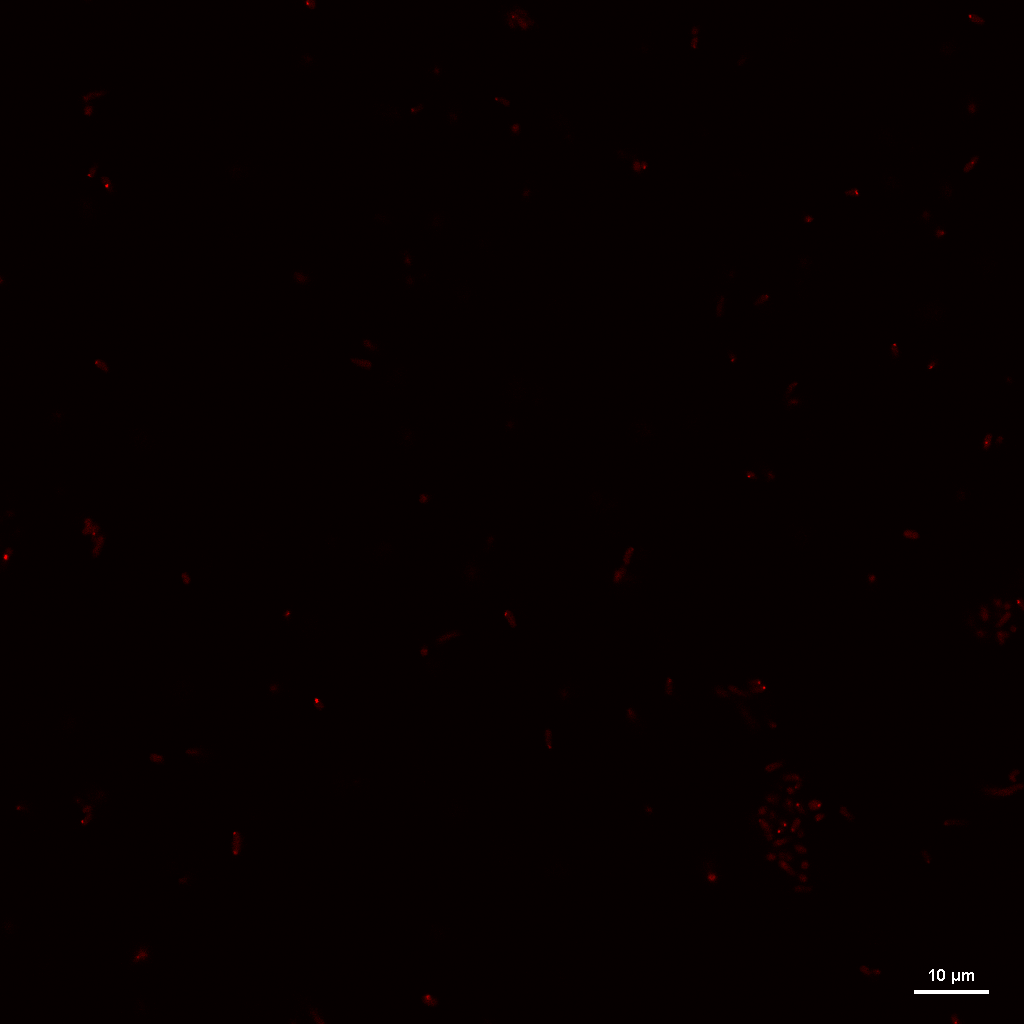

### HMC_N+24_REP2_IMAGE5_RGB_MCHERRY.tif

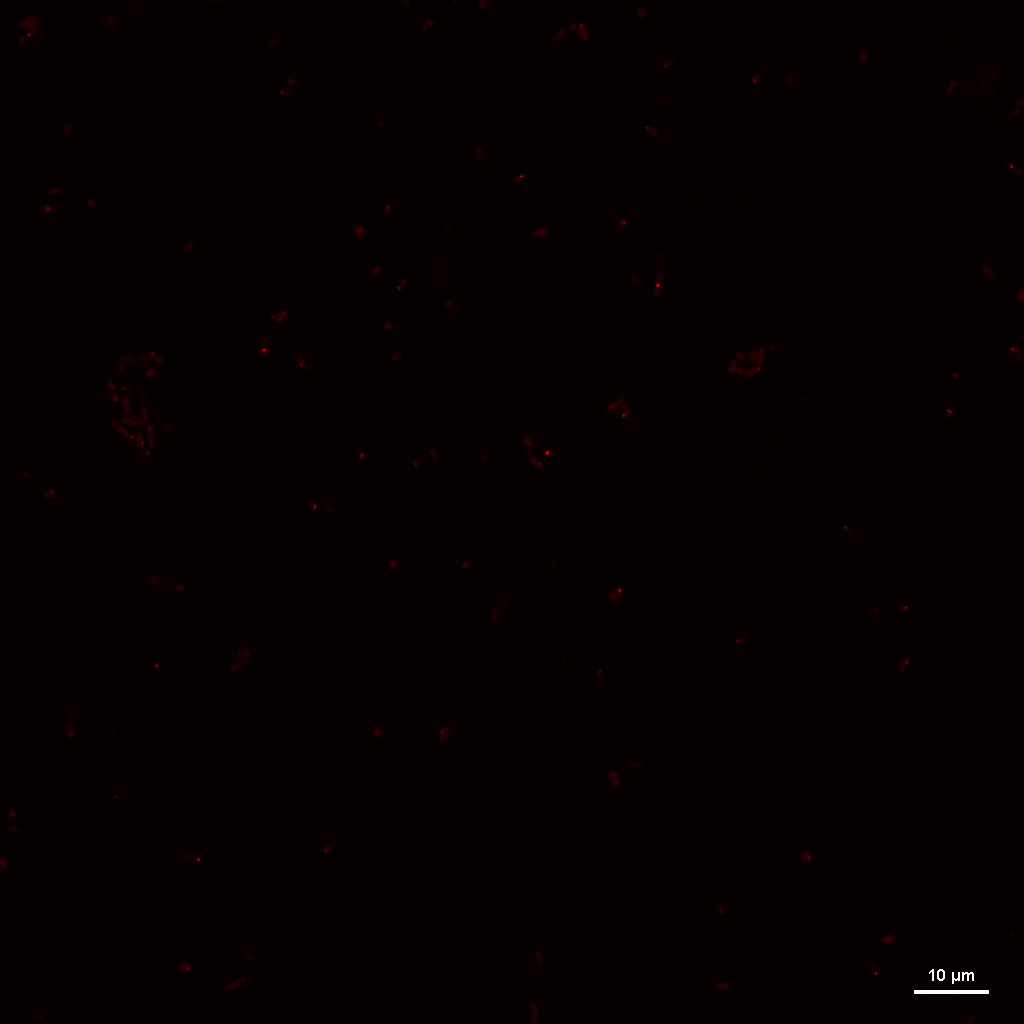

### HMC_N+24_REP3_IMAGE1_RGB_MCHERRY.tif

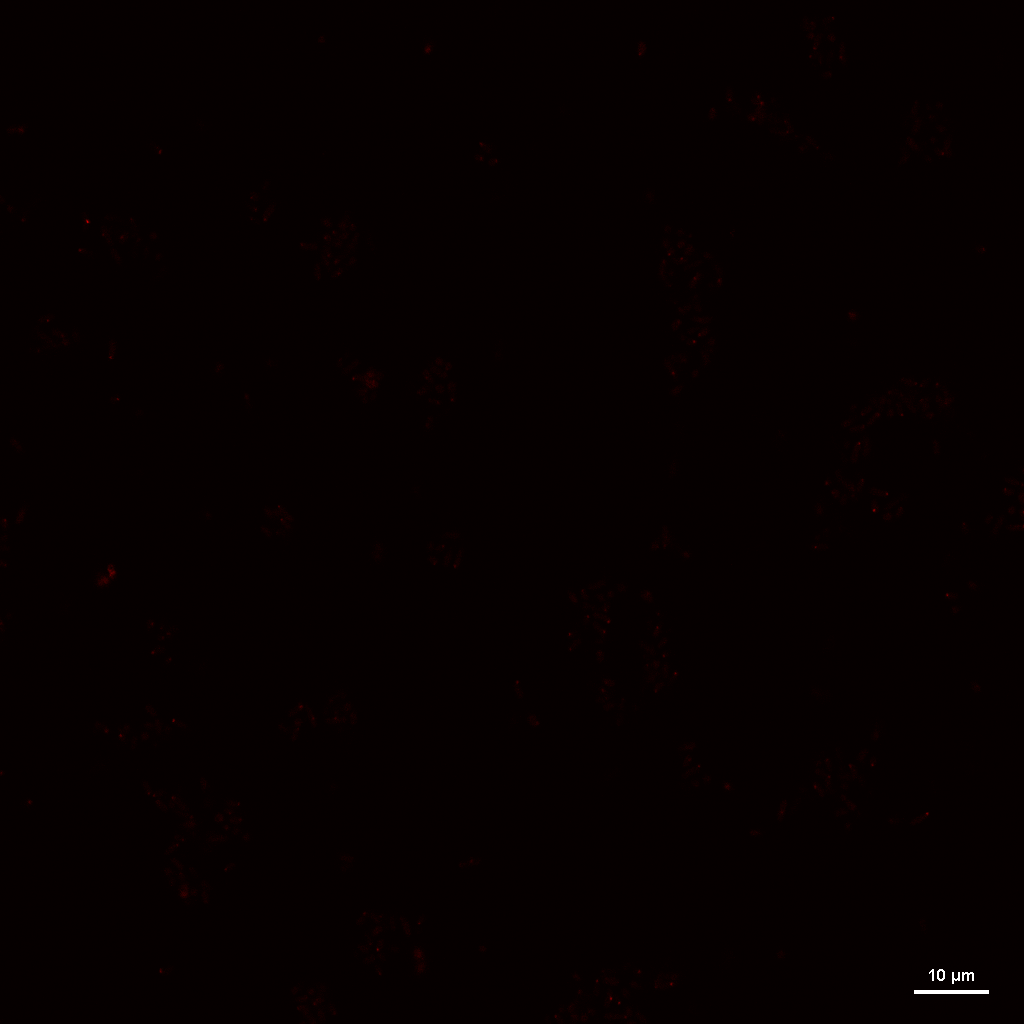

### HMC_N-_REP1_IMAGE1_RGB_MCHERRY.tif

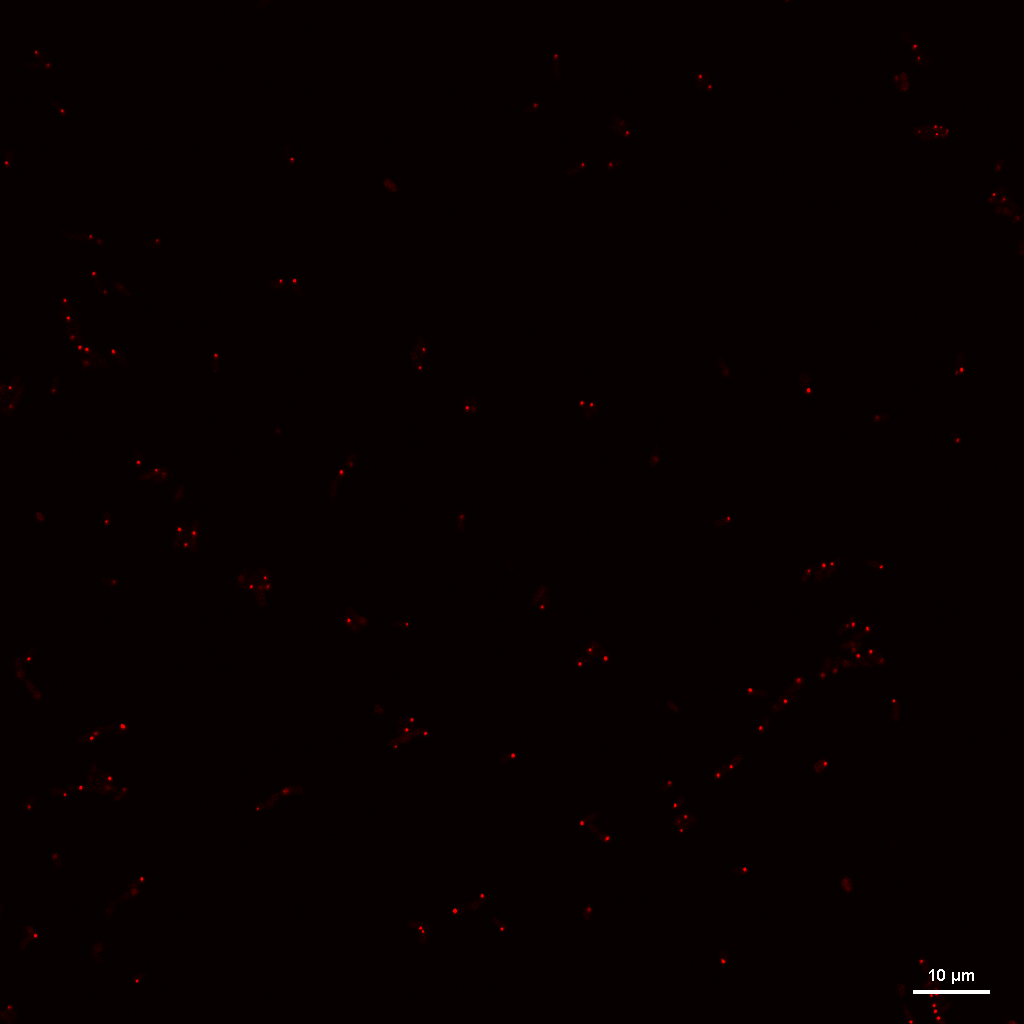

### HMC_N-_REP1_IMAGE2_RGB_MCHERRY.tif

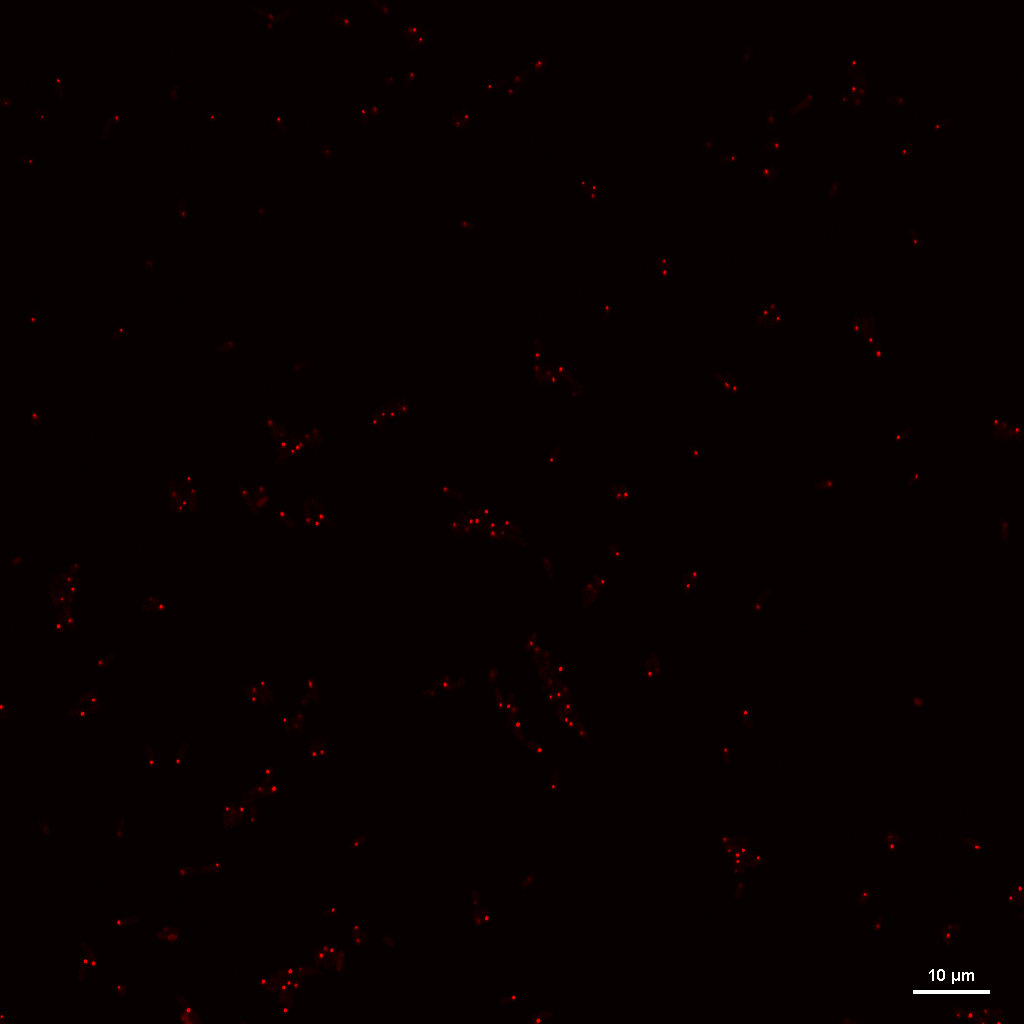

### HMC_N-_REP1_IMAGE3_RGB_MCHERRY.tif

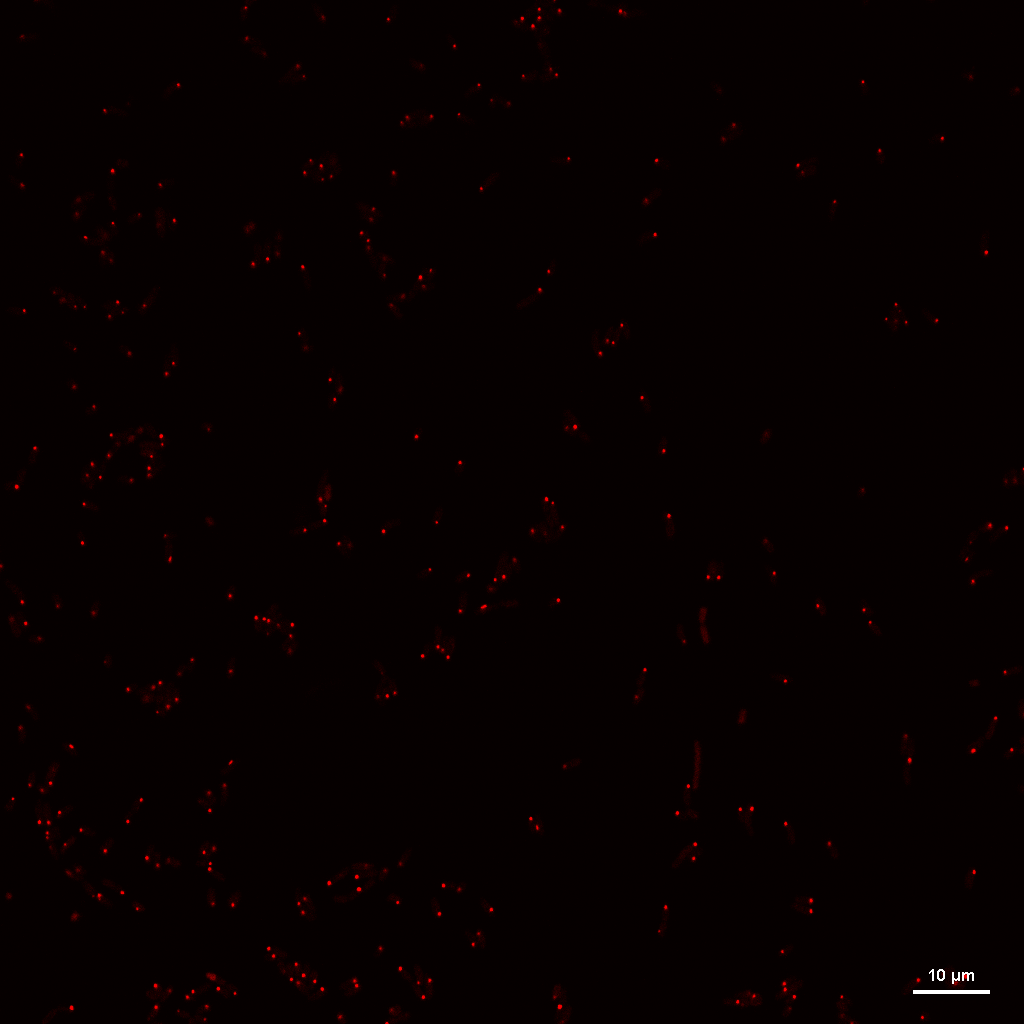

### HMC_N-_REP1_IMAGE4_RGB_MCHERRY.tif

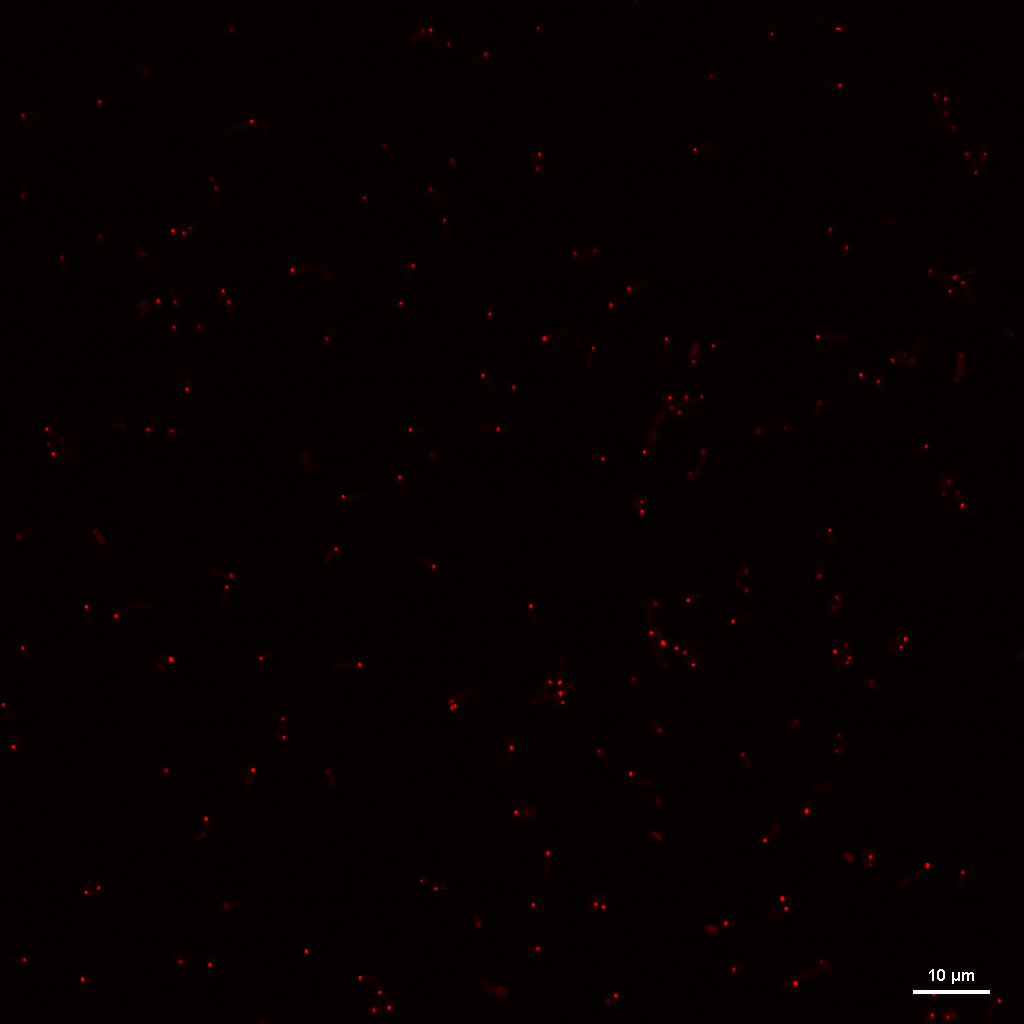

### HMC_N-_REP2_IMAGE1_RGB_MCHERRY.tif

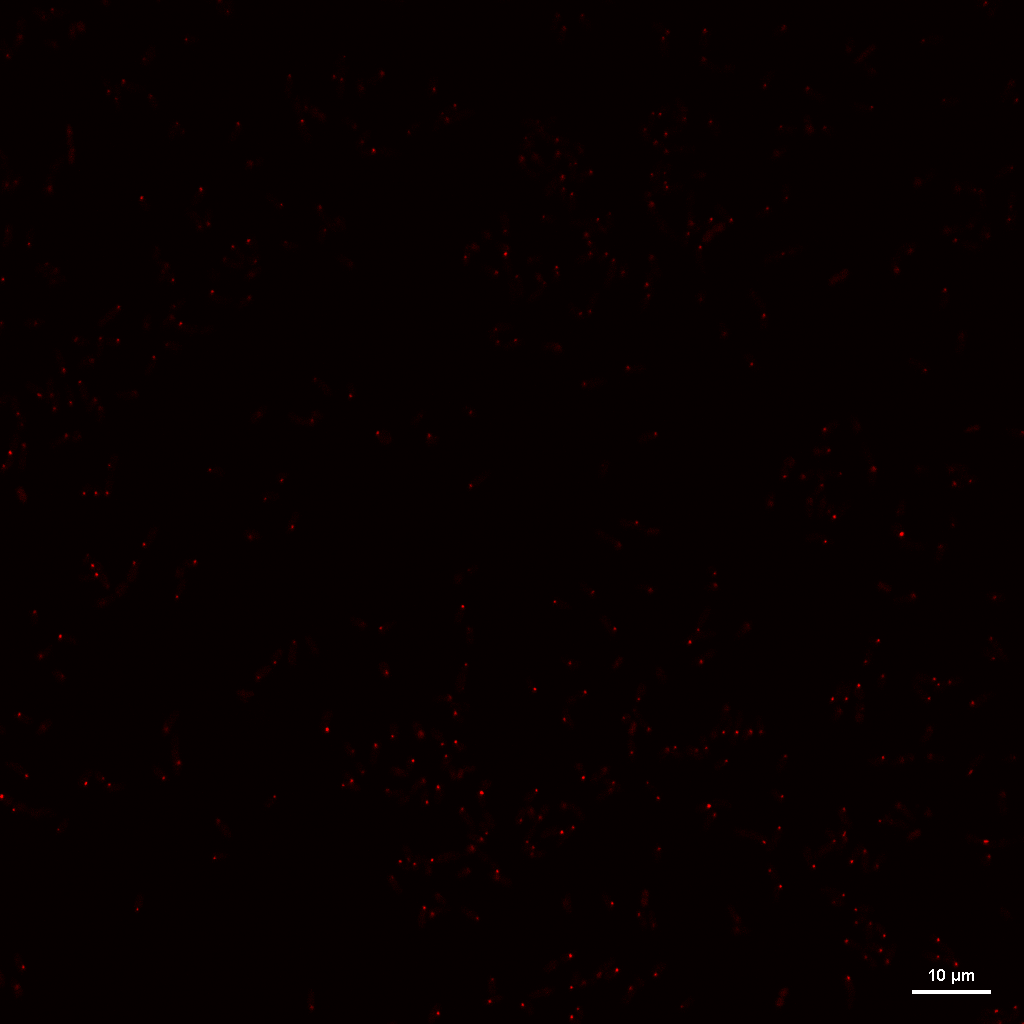

### HMC_N-_REP2_IMAGE2_RGB_MCHERRY.tif

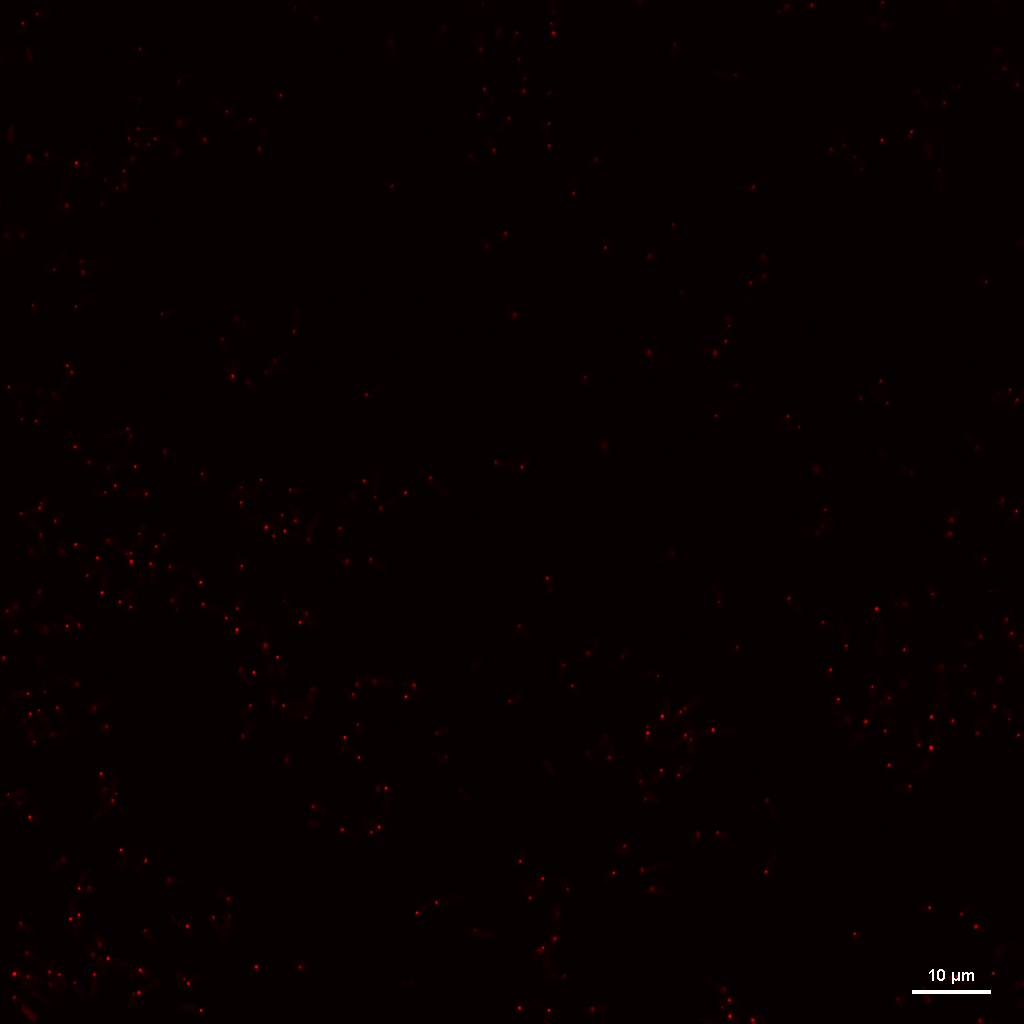

### HMC_N-_REP2_IMAGE3_RGB_MCHERRY.tif

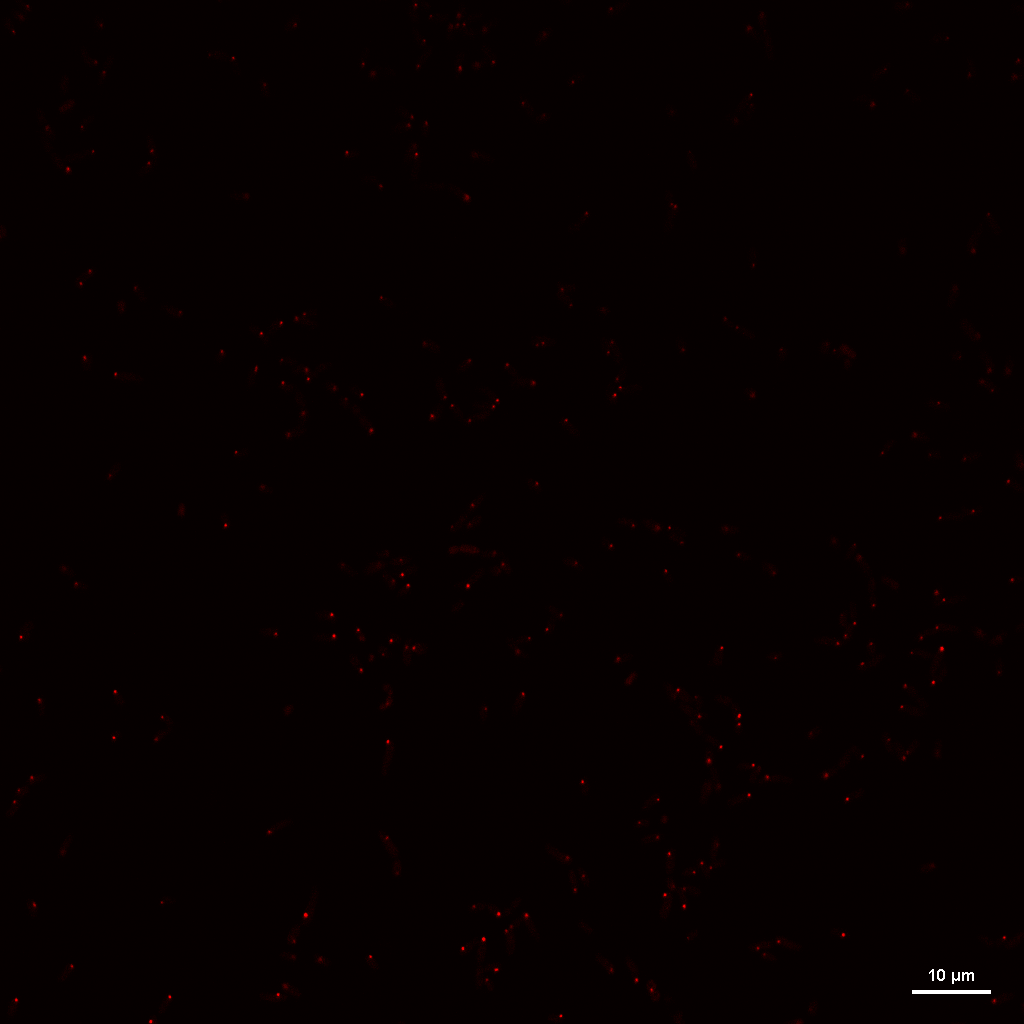

### HMC_N-_REP2_IMAGE4_RGB_MCHERRY.tif

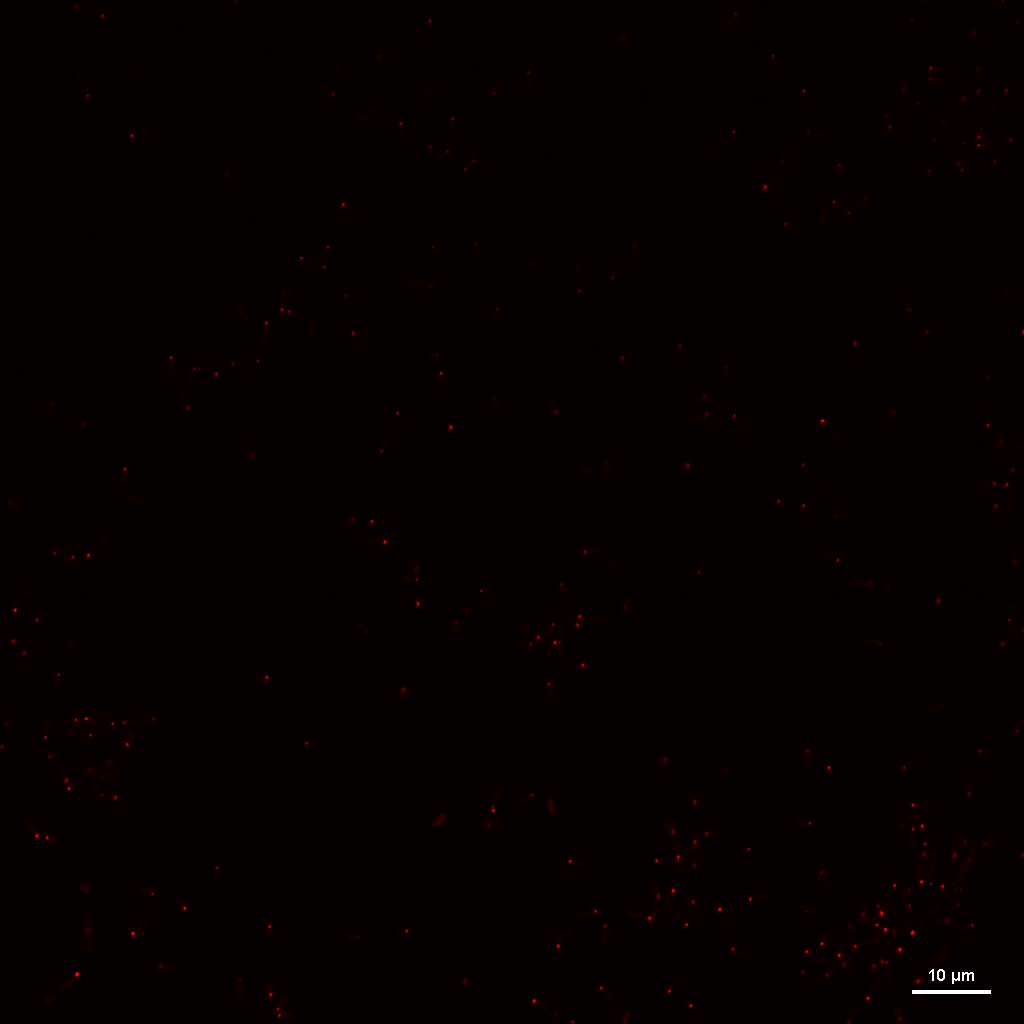

### HMC_N-_REP3_IMAGE1_RGB_MCHERRY.tif

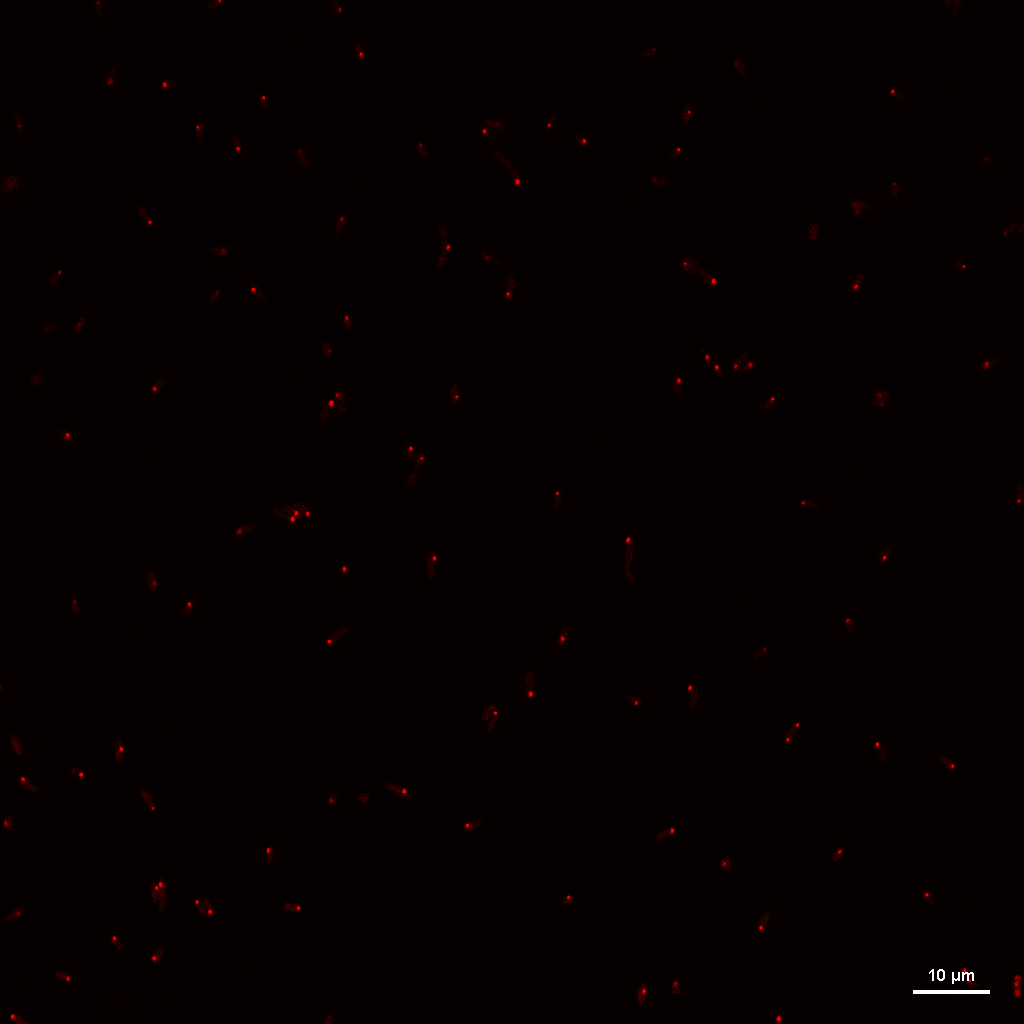

### HMC_N-_REP3_IMAGE2_RGB_MCHERRY.tif

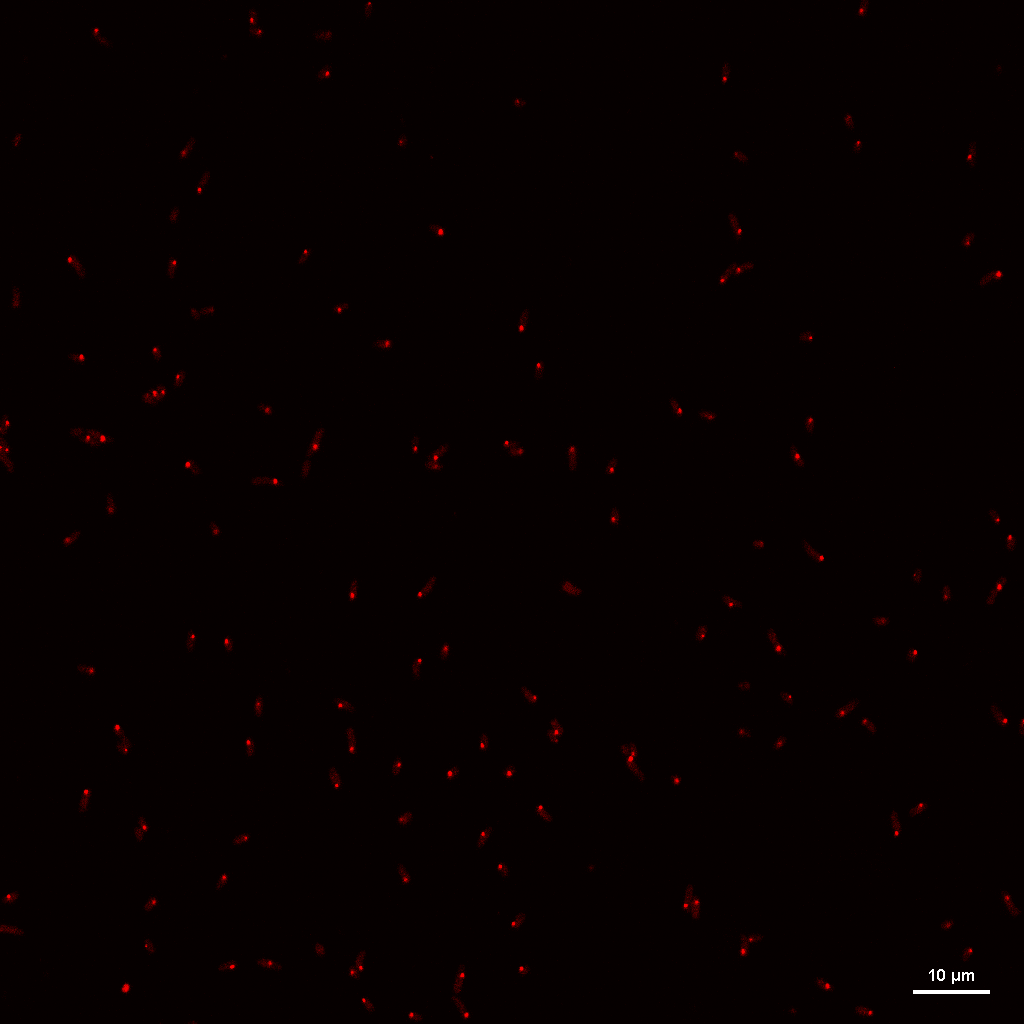
