## Supplementary Figures for "CsrA-dependent RNA accessibility landscapes prioritize new mRNA and sRNA targets and link CsrA to polyphosphate homeostasis"

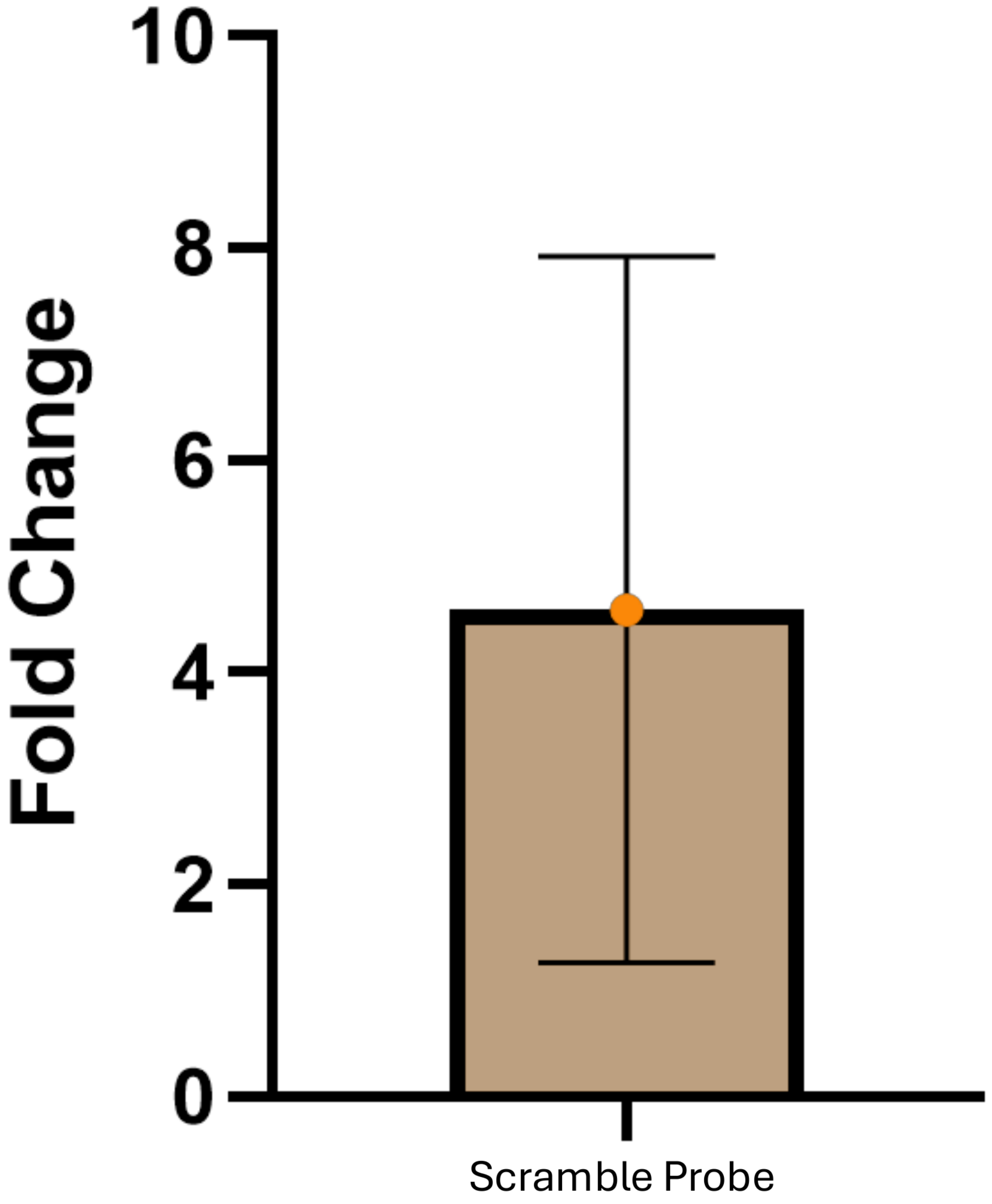

Fold Change
10
8
6
4
2
0
Scramble Probe

***Figure S1.*** Fold change of the scramble probe. Assaying the fold change between the three biological replicates for *csrA::kan* and WT strains. Error bars represent standard deviation. The mean fold change is above one ($p=0.20$ for the difference between strains, $n=3$); see Results for how a possible global shift is handled by our normalization analysis.

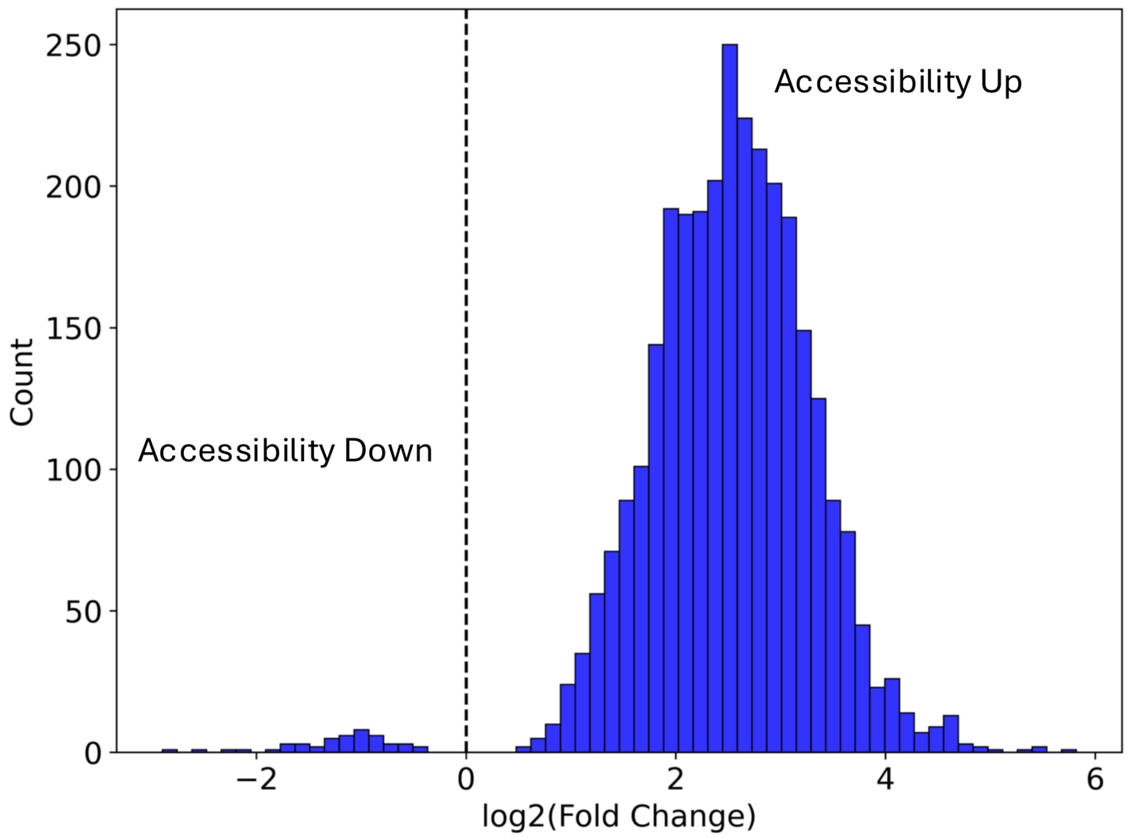

Accessibility Up
Accessibility Down
Count
log2(Fold Change)
250
200
150
100
50
0
−2
0
2
4
6

***Figure S2.*** Distribution of all probe fold changes. The histogram represents the distribution of the accessibility fold change between *csrA::kan* and WT for every probe that had a significant change in accessibility. The x-axis is the log2(fold change) and the y-axis is the count of probes. The vertical line differentiates between “accessibility down” where the probe is more accessible in the WT strain, and “accessibility up” where the strain is more accessible in the *csrA::kan* strain. Most significant probes are more accessible in *csrA::kan* (median log2 fold change = 2.53; fold changes in downstream analyses are normalized with RNA targets to avoid any global shifts (see Results).

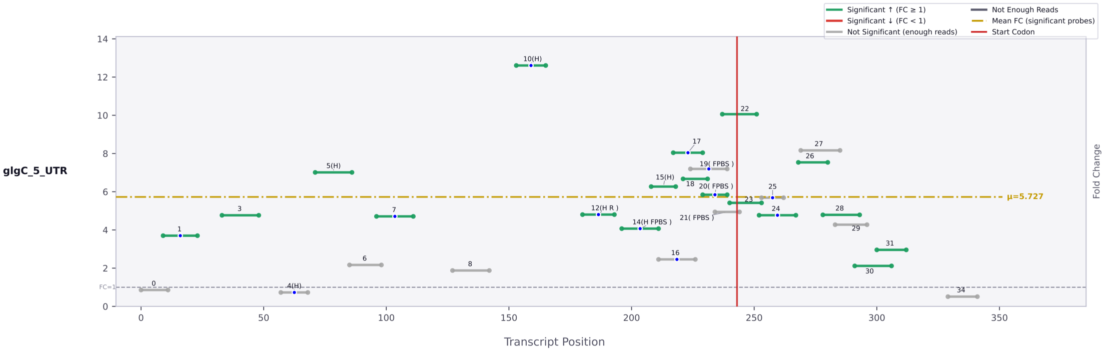
glgC_5_UTR
Significant ↑ (FC ≥ 1)
Significant ↓ (FC < 1)
Not Significant (enough reads)
Not Enough Reads
Mean FC (significant probes)
Start Codon
μ=5.727
FC=1
Fold Change
Transcript Position

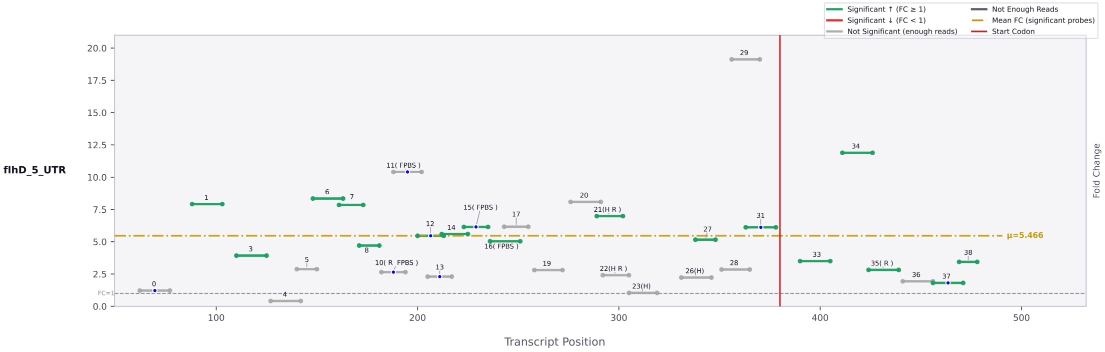
flhD_5_UTR
Significant ↑ (FC ≥ 1)
Significant ↓ (FC < 1)
Not Significant (enough reads)
Not Enough Reads
Mean FC (significant probes)
Start Codon
μ=5.466
FC=1
Fold Change
Transcript Position

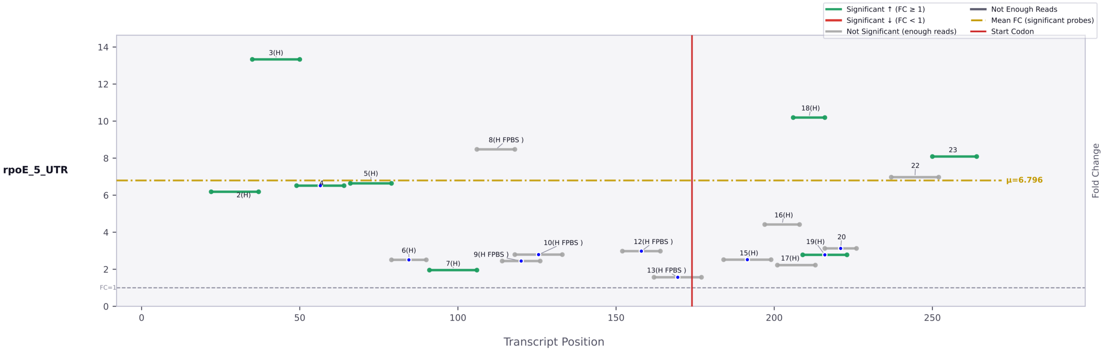
rpoE_5_UTR
Significant ↑ (FC ≥ 1)
Significant ↓ (FC < 1)
Not Significant (enough reads)
Not Enough Reads
Mean FC (significant probes)
Start Codon
μ=6.796
FC=1
Fold Change
Transcript Position

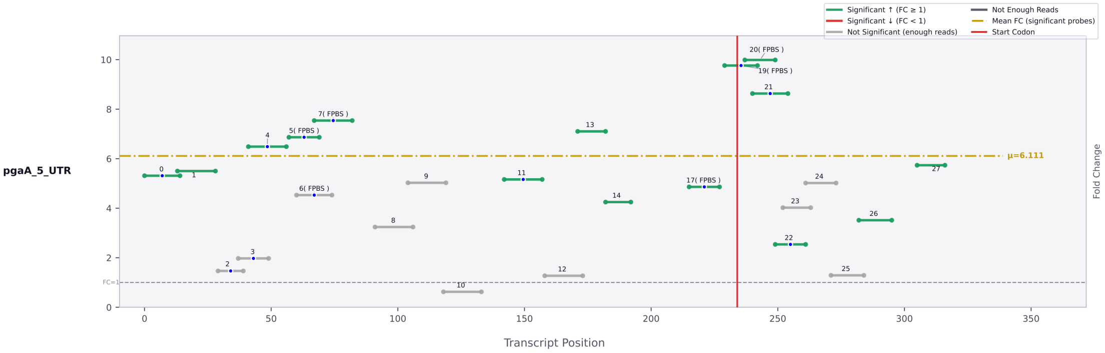
pgaA_5_UTR
Significant ↑ (FC ≥ 1)
Significant ↓ (FC < 1)
Not Significant (enough reads)
Not Enough Reads
Mean FC (significant probes)
Start Codon
μ=6.111
FC=1
Fold Change
Transcript Position

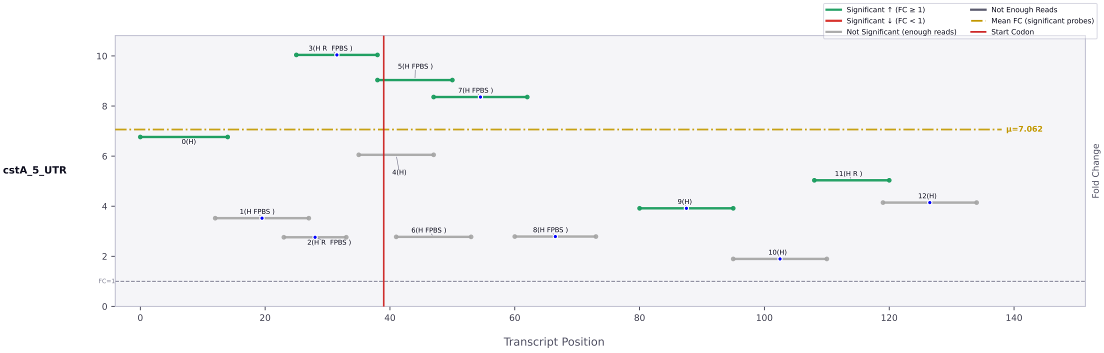
cstA_5_UTR
Significant ↑ (FC ≥ 1)
Significant ↓ (FC < 1)
Not Significant (enough reads)
Not Enough Reads
Mean FC (significant probes)
Start Codon
μ=7.062
FC=1
Transcript Position
Fold Change

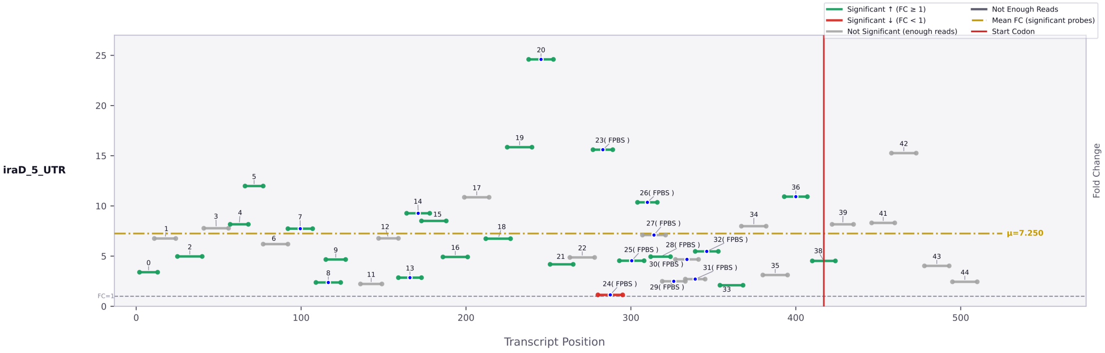
iraD_5_UTR
Significant ↑ (FC ≥ 1)
Significant ↓ (FC < 1)
Not Significant (enough reads)
Not Enough Reads
Mean FC (significant probes)
Start Codon
μ=7.250
FC=1
Transcript Position
Fold Change

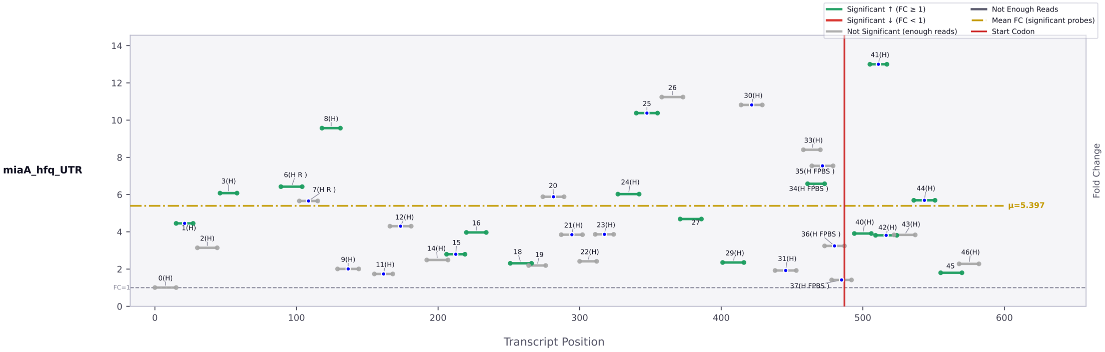
miaA_hfq_UTR
Significant ↑ (FC ≥ 1)
Significant ↓ (FC < 1)
Not Significant (enough reads)
Not Enough Reads
Mean FC (significant probes)
Start Codon
μ=5.397
FC=1
Transcript Position
Fold Change

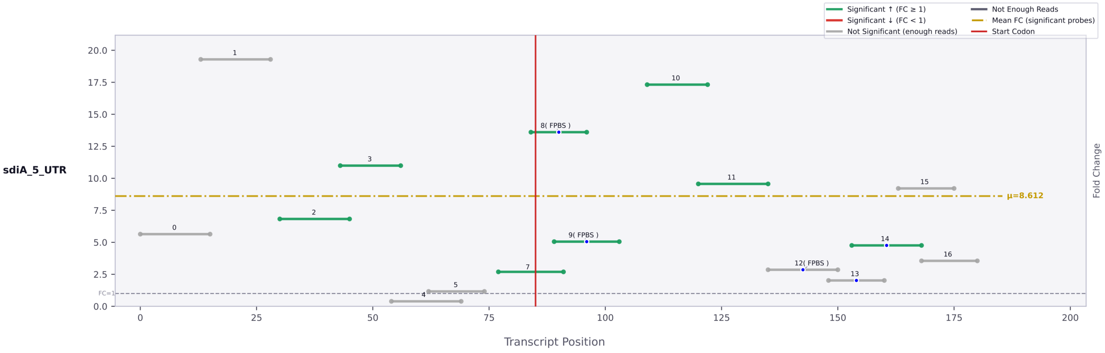
sdiA_5_UTR
Significant ↑ (FC ≥ 1)
Significant ↓ (FC < 1)
Not Significant (enough reads)
Not Enough Reads
Mean FC (significant probes)
Start Codon
μ=8.612
FC=1
Transcript Position
Fold Change

***Figure S3.*** Probe maps of positive control targets. Accessibility map of known CsrA target 5' UTRs: Each horizontal bar in the map of the transcript represents a single asRNA probe. The position on the x-axis signifies the location within the transcript, and the position on the y-axis represents the magnitude of the fold change between the *csrA:kan* and WT strains. Probes containing a blue dot are those that are antisense to a region containing an AGA or GGA motif, signifying a potential CsrA binding site. The color of each bar represents the statistical significance of the probe between the two strains, with green/red bars being statistically significant, and gray bars not significant. The vertical red bar signifies the start codon. The horizontal yellow bar indicates the mean accessibility change of the transcript.

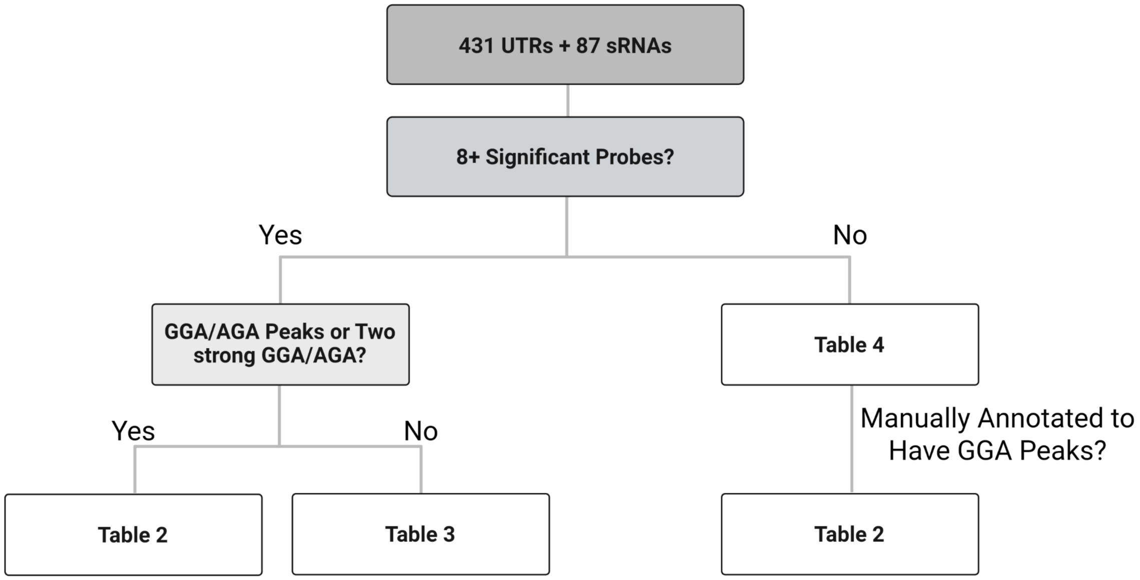
431 UTRs + 87 sRNAs
8+ Significant Probes?
Yes
No
GGA/AGA Peaks or Two strong GGA/AGA?
Table 4
Yes
No
Manually Annotated to Have GGA Peaks?
Table 2
Table 3
Table 2

***Figure S4.*** Filtering Flow Chart. The 431 UTRs and 87 sRNAs studied in this work were first filtered by whether they contained 8+ significant probes. This threshold allows for reduction of noise and identification of high peaks. RNAs containing 8+ significant probes were then filtered by whether they contained a probe in the top 25% of the given RNA that also was antisense to an AGA/GGA motif, suggesting that they may bind CsrA. Those targets were moved to Table 2. RNAs with adequate reads but no peaks were moved to Table 5. Table 6 contains all RNAs that did not meet the 8+ probe threshold. Some RNAs that lacked a large number of probes in the library were manually annotated to have GGA/AGA peaks and were moved into Table 2. All 32 3'UTRs and 20 of the 28 sRNAs in Table 2 were manually annotated' 90 5'UTRs and 8 sRNAs were called by automated filter.

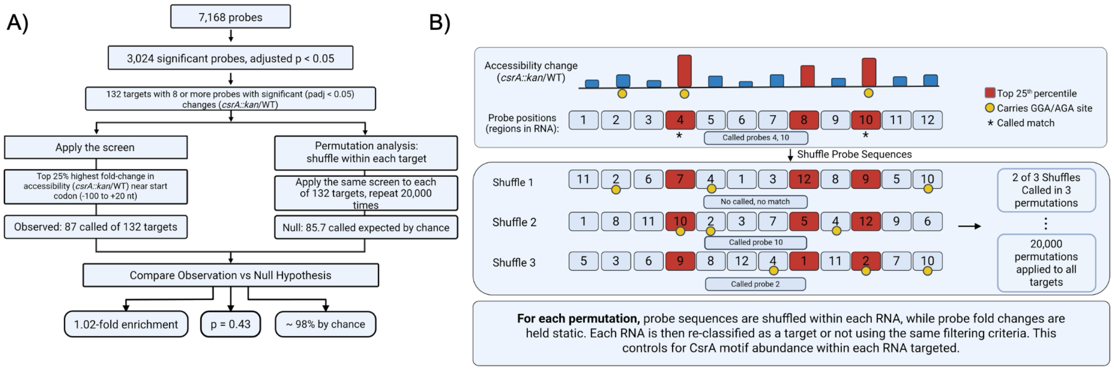

A)
7,168 probes
3,024 significant probes, adjusted p < 0.05
132 targets with 8 or more probes with significant (padj < 0.05) changes (csrA::kan/WT)
Apply the screen
Top 25% highest fold-change in accessibility (csrA::kan/WT) near start codon (-100 to +20 nt)
Observed: 87 called of 132 targets
Permutation analysis: shuffle within each target
Apply the same screen to each of 132 targets, repeat 20,000 times
Null: 85.7 called expected by chance
Compare Observation vs Null Hypothesis
1.02-fold enrichment
p = 0.43
~ 98% by chance
B)
Accessibility change (csrA::kan/WT)
Probe positions (regions in RNA):
Top 25th percentile
Carries GGA/AGA site
* Called match
Called probes 4, 10
Shuffle Probe Sequences
Shuffle 1
No called, no match
Shuffle 2
Called probe 10
Shuffle 3
Called probe 2
2 of 3 Shuffles Called in 3 permutations
20,000 permutations applied to all targets
For each permutation, probe sequences are shuffled within each RNA, while probe fold changes are held static. Each RNA is then re-classified as a target or not using the same filtering criteria. This controls for CsrA motif abundance within each RNA targeted.

***Figure S5.*** Permutational analysis of filtering methodology. (A) The flow of how permutational analysis was performed. At a macro scale, all targets were screened using filtering described in **Figure S3.** Then, for all targets permutations were performed 20,000 times and a mean number of discovered targets was calculated. This was converted into an enrichment and a p-value in order to determine the likelihood of enrichment over random noise. (B) Permutational analysis was performed by fixing the position and fold changes of each probe and shuffling their locations. For each shuffle, it was determined whether the combination of probe sequence, location, and fold change met the criteria to be considered a target. The higher the number of unique permutations that successfully passed the screen, the more likely that the effects observed were due to random chance.

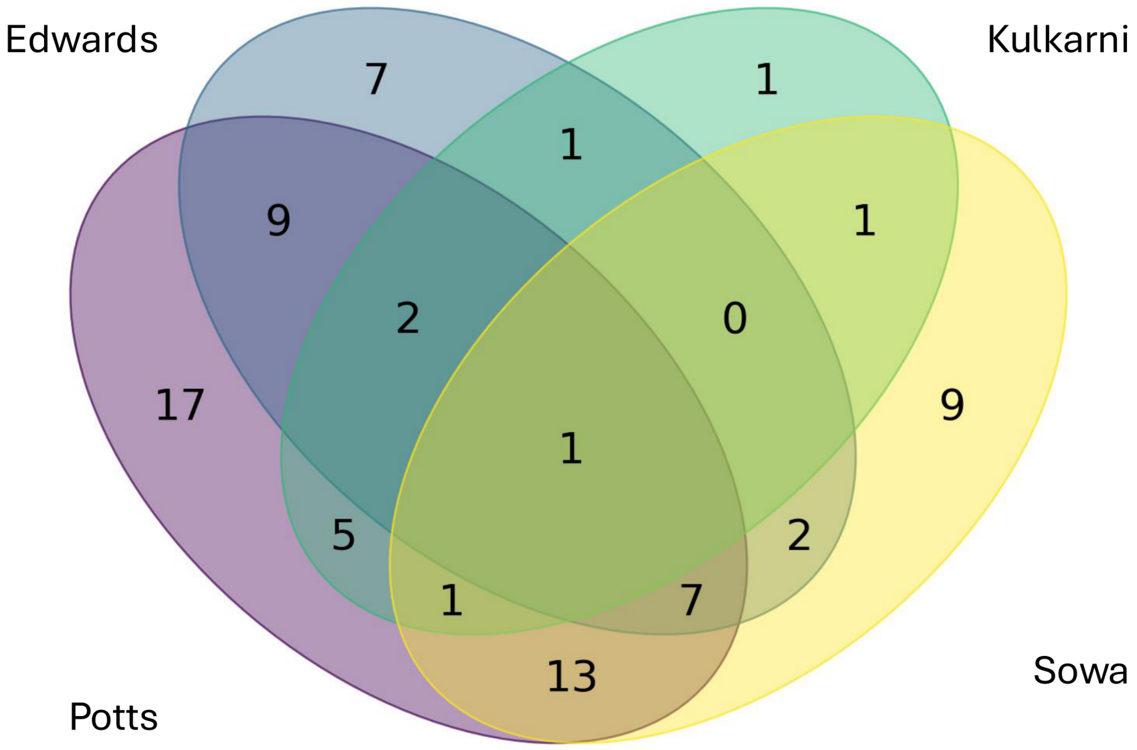
Edwards
Kulkarni
7
1
1
9
1
2
0
17
1
9
5
2
1
7
13
Sowa
Potts

***Figure S6.*** Comparing identified targets to published literature. The overlap of putative CsrA targets with previously published high throughput or computational datasets. Numbers in the Venn diagram sum to 79 of 90 candidate 5'UTRs with prior evidence.

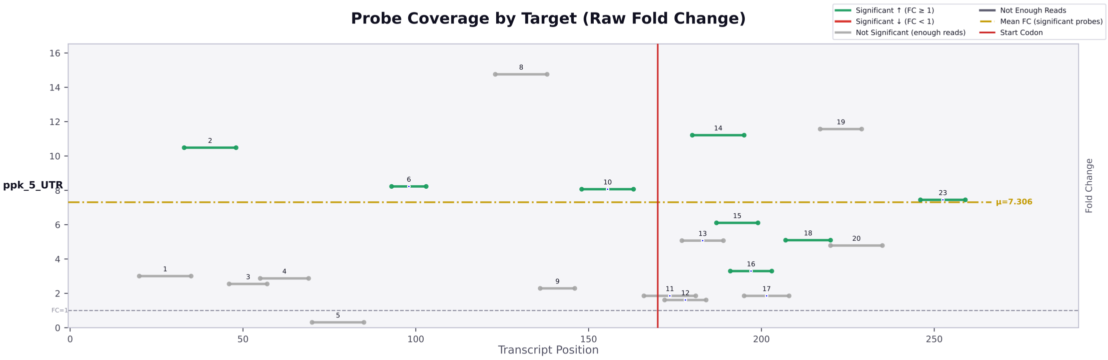

Probe Coverage by Target (Raw Fold Change)
Significant ↑ (FC ≥ 1)
Significant ↓ (FC < 1)
Not Significant (enough reads)
Not Enough Reads
Mean FC (significant probes)
Start Codon
ppk_5_UTR
μ=7.306
FC=1
Transcript Position
Fold Change


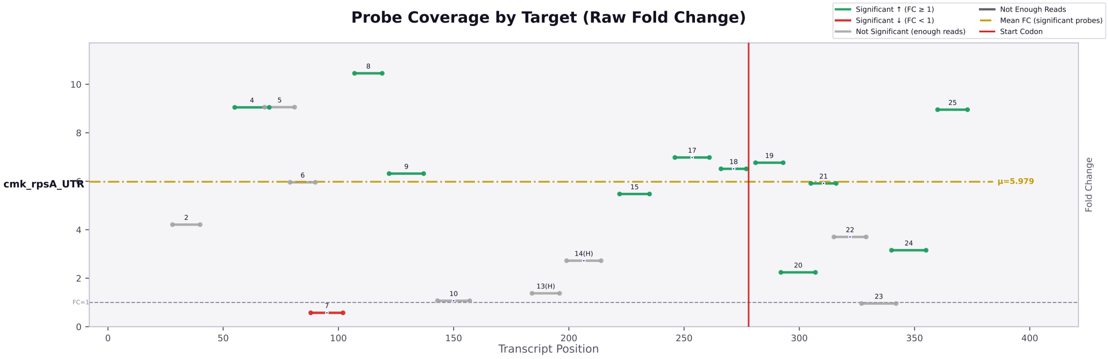

Probe Coverage by Target (Raw Fold Change)
Significant ↑ (FC ≥ 1)
Significant ↓ (FC < 1)
Not Significant (enough reads)
Not Enough Reads
Mean FC (significant probes)
Start Codon
cmk_rpsA_UTR
μ=5.979
FC=1
Transcript Position
Fold Change


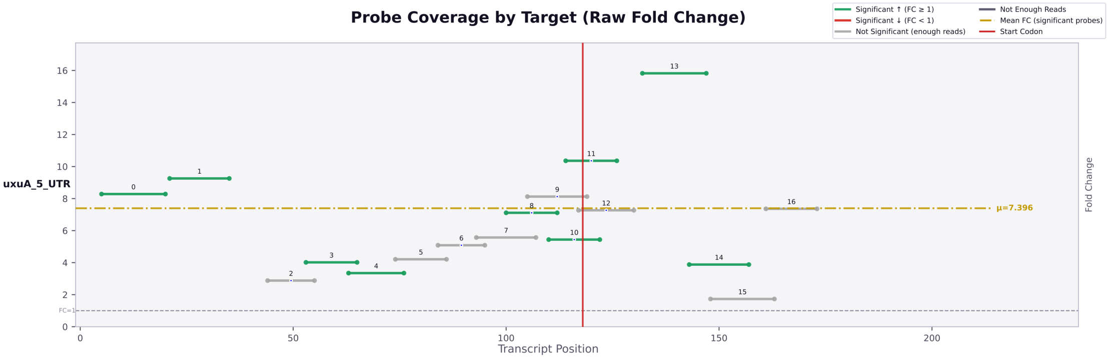

Probe Coverage by Target (Raw Fold Change)
Significant ↑ (FC ≥ 1)
Significant ↓ (FC < 1)
Not Significant (enough reads)
Not Enough Reads
Mean FC (significant probes)
Start Codon
uxuA_5_UTR
μ=7.396
FC=1
Transcript Position
Fold Change


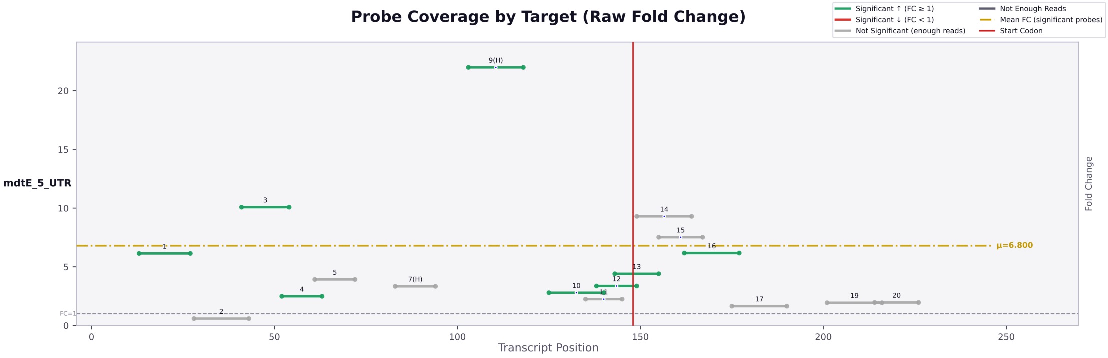

Probe Coverage by Target (Raw Fold Change)
Significant ↑ (FC ≥ 1)
Significant ↓ (FC < 1)
Not Significant (enough reads)
Not Enough Reads
Mean FC (significant probes)
Start Codon
mdtE_5_UTR
μ=6.800
FC=1
Transcript Position
Fold Change

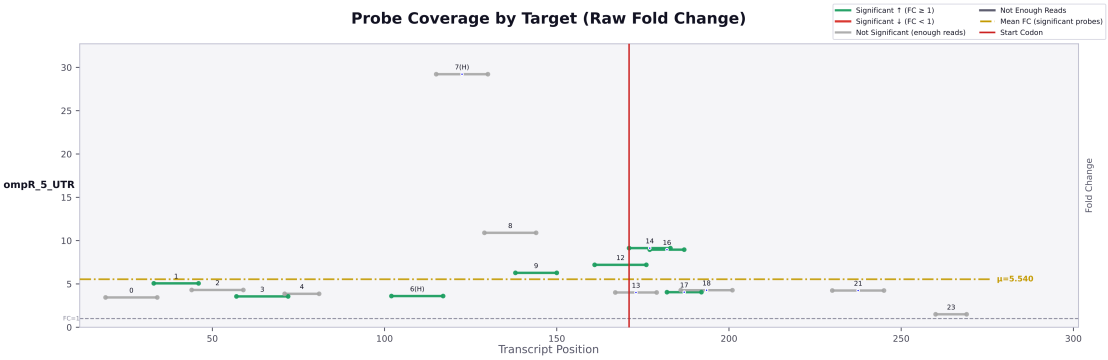
Probe Coverage by Target (Raw Fold Change)
Significant ↑ (FC ≥ 1)
Significant ↓ (FC < 1)
Not Significant (enough reads)
Not Enough Reads
Mean FC (significant probes)
Start Codon
ompR_5_UTR
Fold Change
Transcript Position
μ=5.540
FC=1

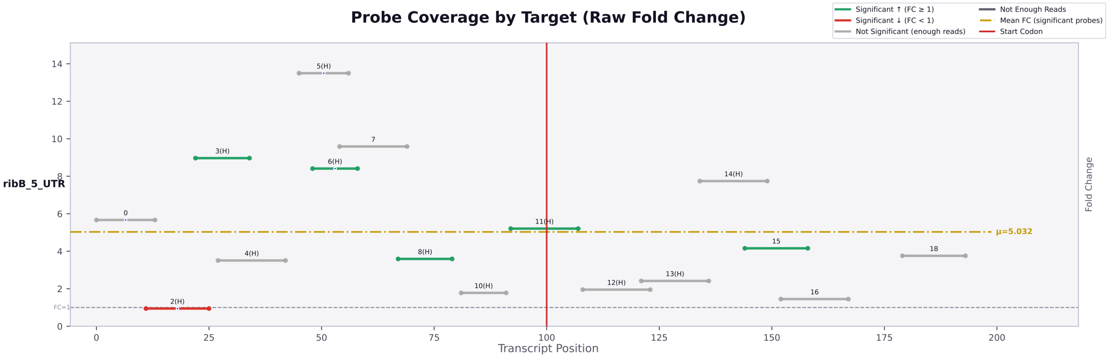
Probe Coverage by Target (Raw Fold Change)
Significant ↑ (FC ≥ 1)
Significant ↓ (FC < 1)
Not Significant (enough reads)
Not Enough Reads
Mean FC (significant probes)
Start Codon
ribB_5_UTR
Fold Change
Transcript Position
μ=5.032
FC=1

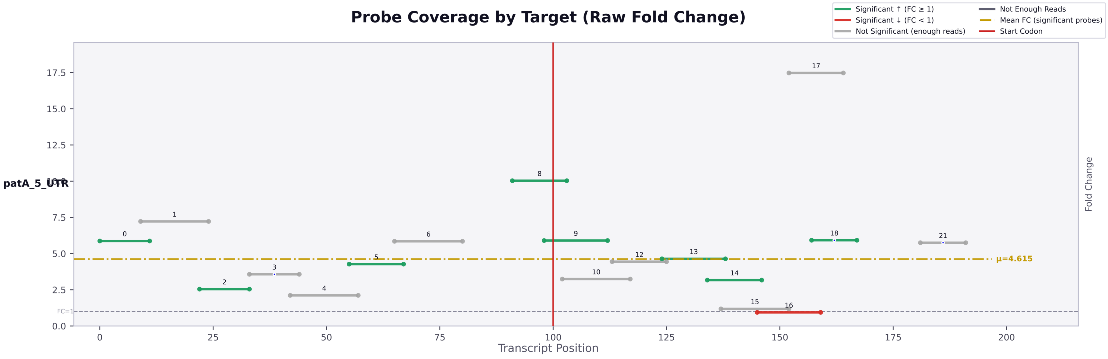
Probe Coverage by Target (Raw Fold Change)
Significant ↑ (FC ≥ 1)
Significant ↓ (FC < 1)
Not Significant (enough reads)
Not Enough Reads
Mean FC (significant probes)
Start Codon
patA_5_UTR
Fold Change
Transcript Position
μ=4.615
FC=1

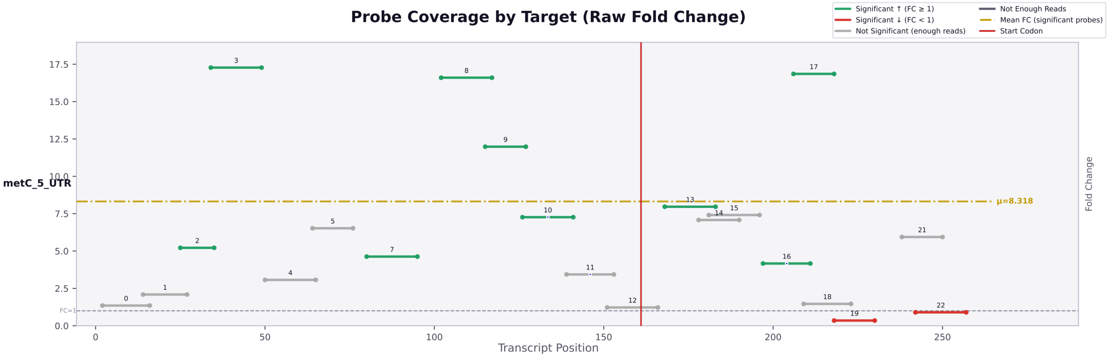
Probe Coverage by Target (Raw Fold Change)
Significant ↑ (FC ≥ 1)
Significant ↓ (FC < 1)
Not Significant (enough reads)
Not Enough Reads
Mean FC (significant probes)
Start Codon
metC_5_UTR
Fold Change
Transcript Position
μ=8.318
FC=1

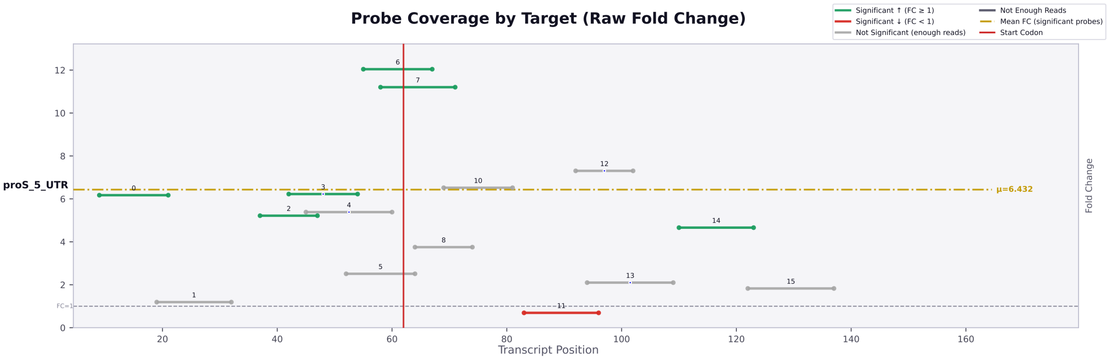

Probe Coverage by Target (Raw Fold Change)
Significant ↑ (FC ≥ 1)
Significant ↓ (FC < 1)
Not Significant (enough reads)
Not Enough Reads
Mean FC (significant probes)
Start Codon
proS_5_UTR
μ=6.432
FC=1
Transcript Position
Fold Change

***Figure S7.*** Accessibility maps of putative CsrA target 5' UTRs tested in Figure 2B. Each horizontal bar in the map of the transcript represents a single asRNA probe. The position on the x-axis signifies the location within the transcript, and the position on the y-axis represents the magnitude of the fold change between the *csrA:kan* and WT strains. Probes containing a blue dot are those that are antisense to a region containing an AGA or GGA motif, signifying a potential CsrA binding site. The color of each bar represents the statistical significance of the probe between the two strains, with green/red bars being statistically significant, and gray bars not significant. The vertical red bar signifies the start codon. The horizontal yellow bar indicates the mean accessibility change of the transcript.

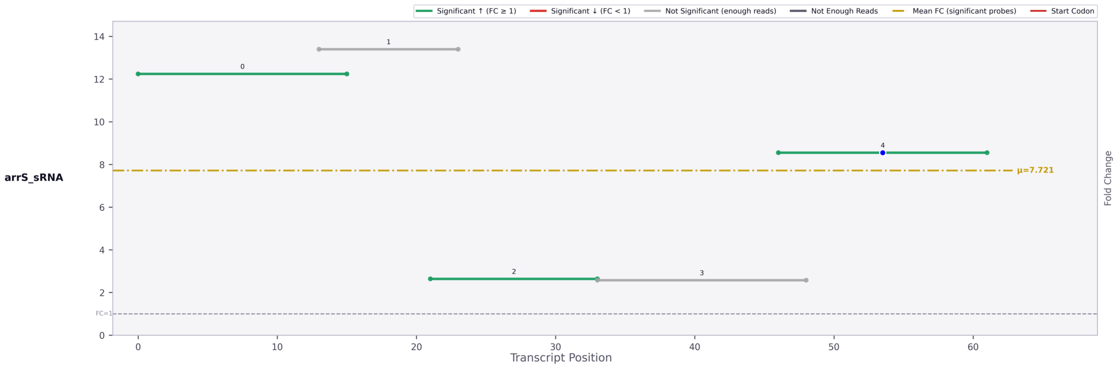

Significant ↑ (FC ≥ 1)
Significant ↓ (FC < 1)
Not Significant (enough reads)
Not Enough Reads
Mean FC (significant probes)
Start Codon
arrS_sRNA
0
1
2
3
4
μ=7.721
FC=1
Fold Change
Transcript Position

***Figure S8.*** Accessibility map of the ArrS sRNA**.** (Probe 2 marks the ribB-interacting region) tested in Figure 3**:** Each horizontal bar in the map of the transcript represents a single asRNA probe. The position on the x-axis signifies the location within the transcript, and the position on the y-axis represents the magnitude of the fold change between the *csrA::kan* and WT strains. Probes containing a blue dot are those that are antisense to a region containing an AGA or GGA motif, signifying a potential CsrA binding site. The color of each bar represents the statistical significance of the probe between the two strains, with green/red bars being statistically significant, and gray bars not significant. The vertical red bar signifies the start codon. The horizontal yellow bar indicates the mean accessibility change of the transcript.

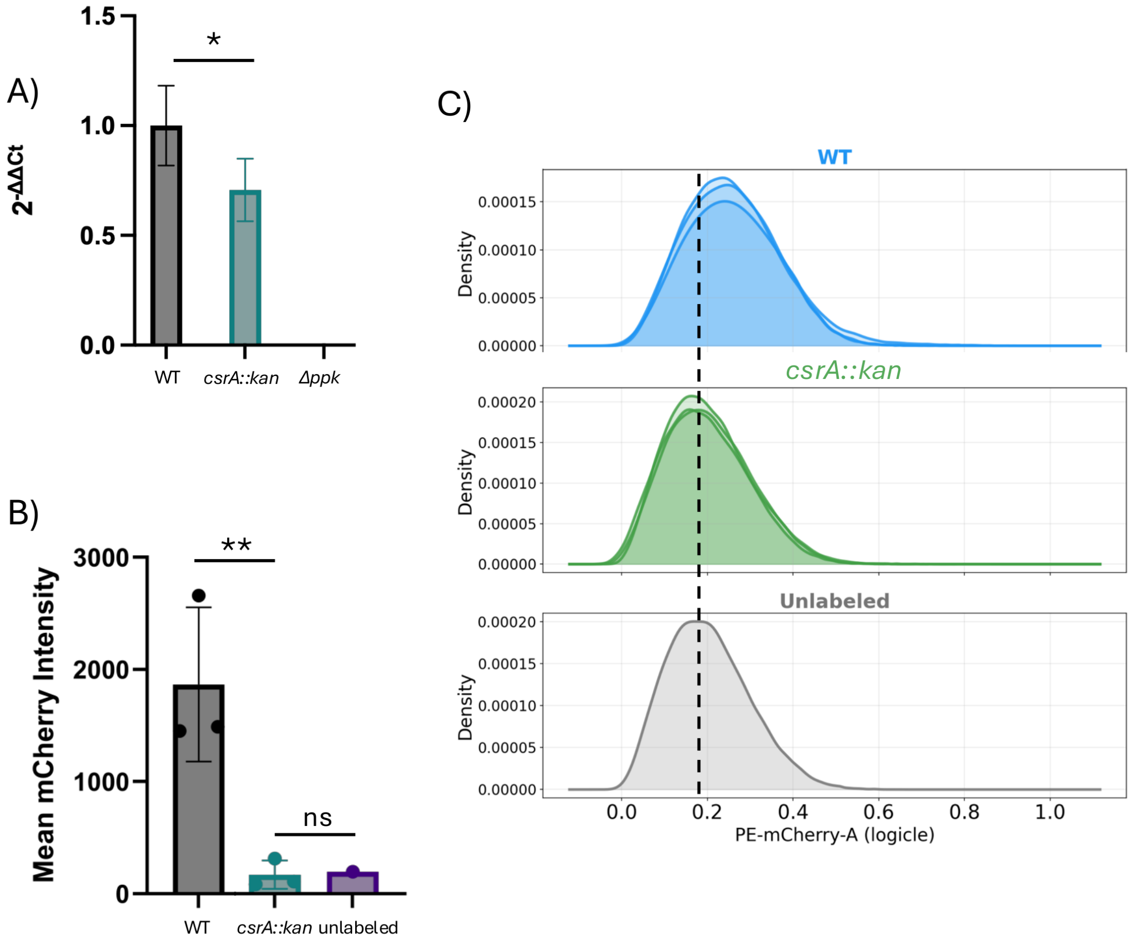

A)
1.5
1.0
0.5
0.0
2-ΔΔCt
*
WT
csrA::kan
Δppk
B)
3000
2000
1000
0
Mean mCherry Intensity
**
ns
WT
csrA::kan
unlabeled
C)
WT
csrA::kan
Unlabeled
Density
0.00000
0.00005
0.00010
0.00015
0.00020
0.0
0.2
0.4
0.6
0.8
1.0
PE-mCherry-A (logicle)

***Figure S9.*** Impacts of CsrA on *ppk* RNA and protein levels *in vivo*. **(A)** RT-qPCR on 24 hour nitrogen starved cells, quantifying the amount of *ppk* mRNA WT and *csrA::kan* strains (n=3) with varying levels of CsrA activity and a *ppk* knockout control (value too small to visualize) **(B)** Comparison of mean values of mCherry intensity in genomically labeled *ppk*-mCherry strains. Error bars indicate SD **(C)** Flow cytometry was performed on three strains with mCherry labeled *ppk,* and an unlabeled MG1655 control. Data normalized using logicle transformation. (n=3 independent biological replicates for WT, BC, CK, n=1 replicate for unlabeled cells). WT = WT MG1655, BC = MG1655 *Δ*csrBC, and CK = MG1655 *csrA::kan.* . All Comparisons made using Dunnett multiple t test following one-way ANOVA. *p <0.05, ** p< 0.01, *** p<0.001, **** p<0.0001

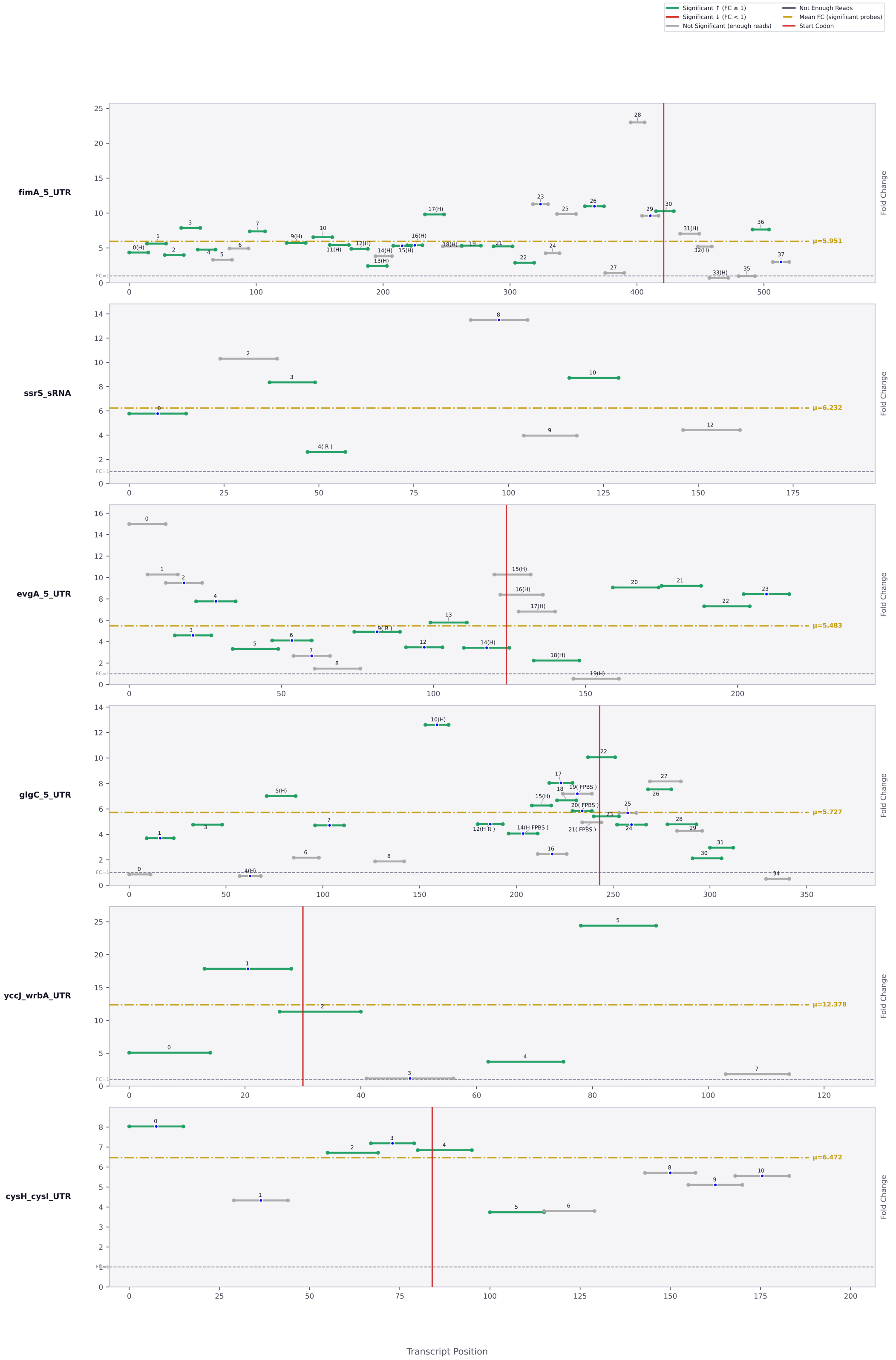

Significant ↑ (FC ≥ 1)
Significant ↓ (FC < 1)
Not Significant (enough reads)
Not Enough Reads
Mean FC (significant probes)
Start Codon
fimA_5_UTR
μ=5.951
ssrS_sRNA
μ=6.232
evgA_5_UTR
μ=5.483
glgC_5_UTR
μ=5.727
yccJ_wrbA_UTR
μ=12.378
cysH_cysI_UTR
μ=6.472
Fold Change
FC=1
Transcript Position

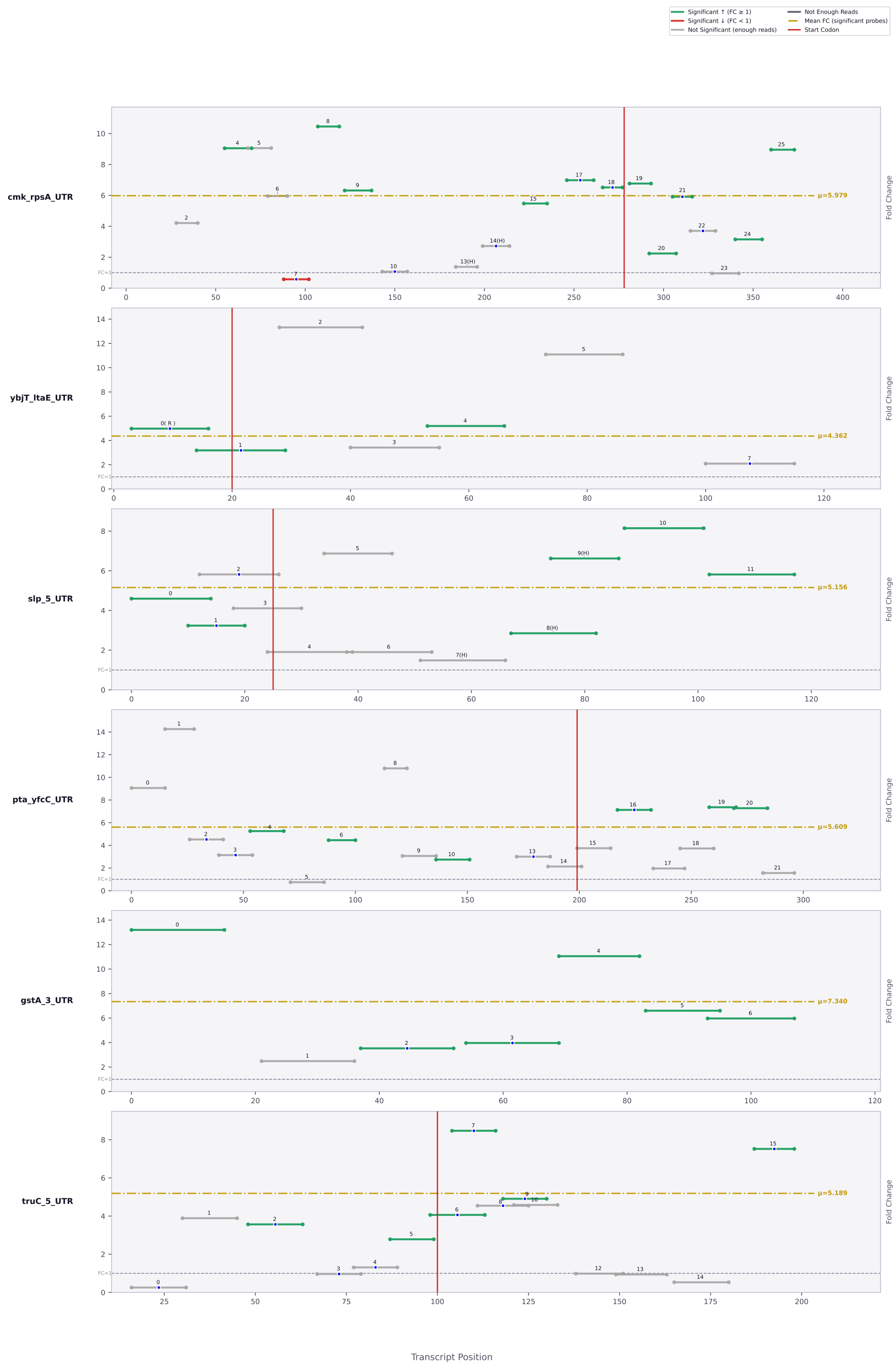

Significant ↑ (FC ≥ 1)
Significant ↓ (FC < 1)
Not Significant (enough reads)
Not Enough Reads
Mean FC (significant probes)
Start Codon
cmk_rpsA_UTR
μ=5.979
ybjT_ltaE_UTR
μ=4.362
slp_5_UTR
μ=5.156
pta_yfcC_UTR
μ=5.609
gstA_3_UTR
μ=7.340
truC_5_UTR
μ=5.189
Fold Change
Transcript Position

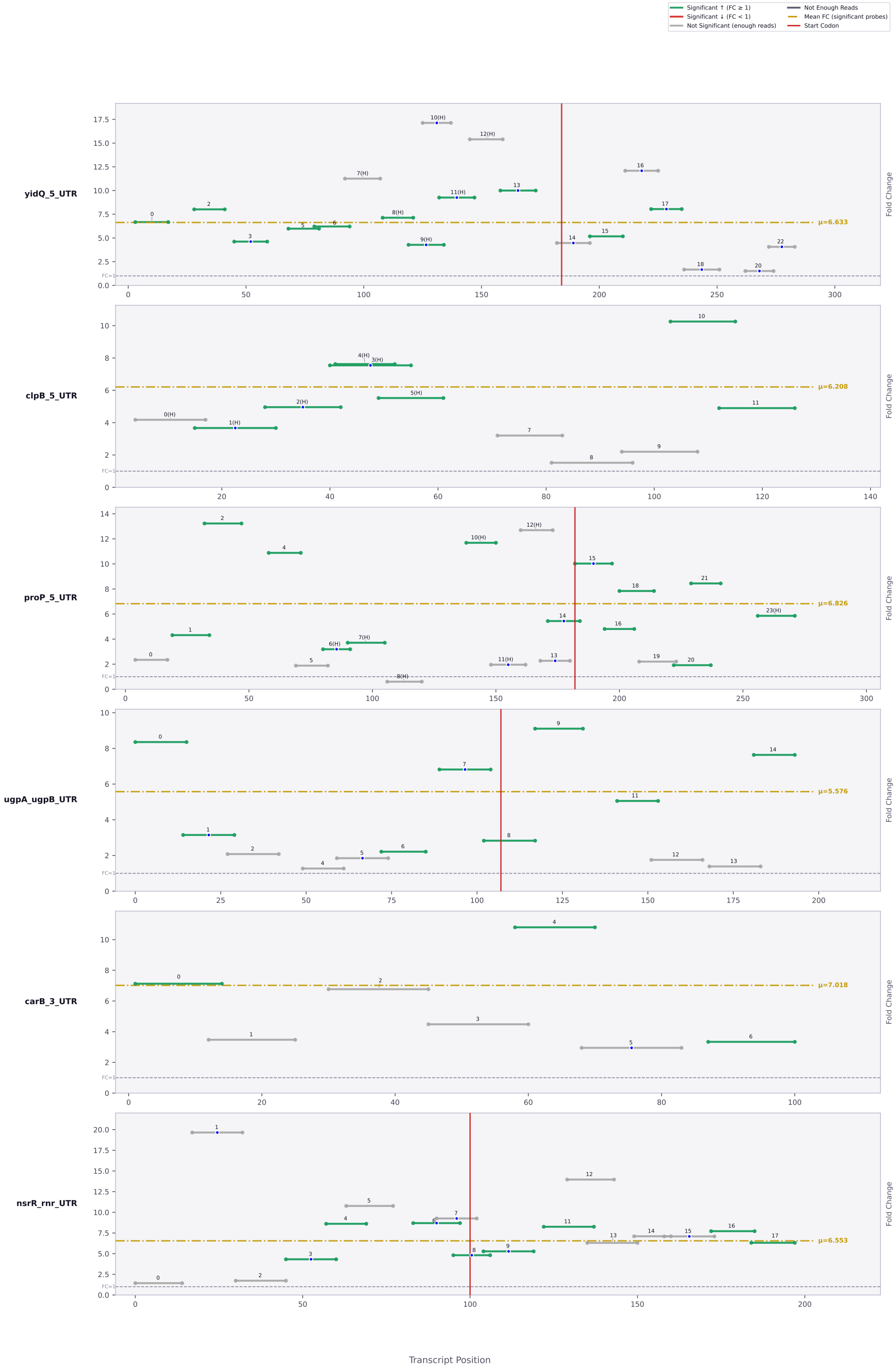
Significant ↑ (FC ≥ 1)
Significant ↓ (FC < 1)
Not Significant (enough reads)
Not Enough Reads
Mean FC (significant probes)
Start Codon
yidQ_5_UTR
μ=6.633
clpB_5_UTR
μ=6.208
proP_5_UTR
μ=6.826
ugpA_ugpB_UTR
μ=5.576
carB_3_UTR
μ=7.018
nsrR_rnr_UTR
μ=6.553
Fold Change
FC=1
Transcript Position

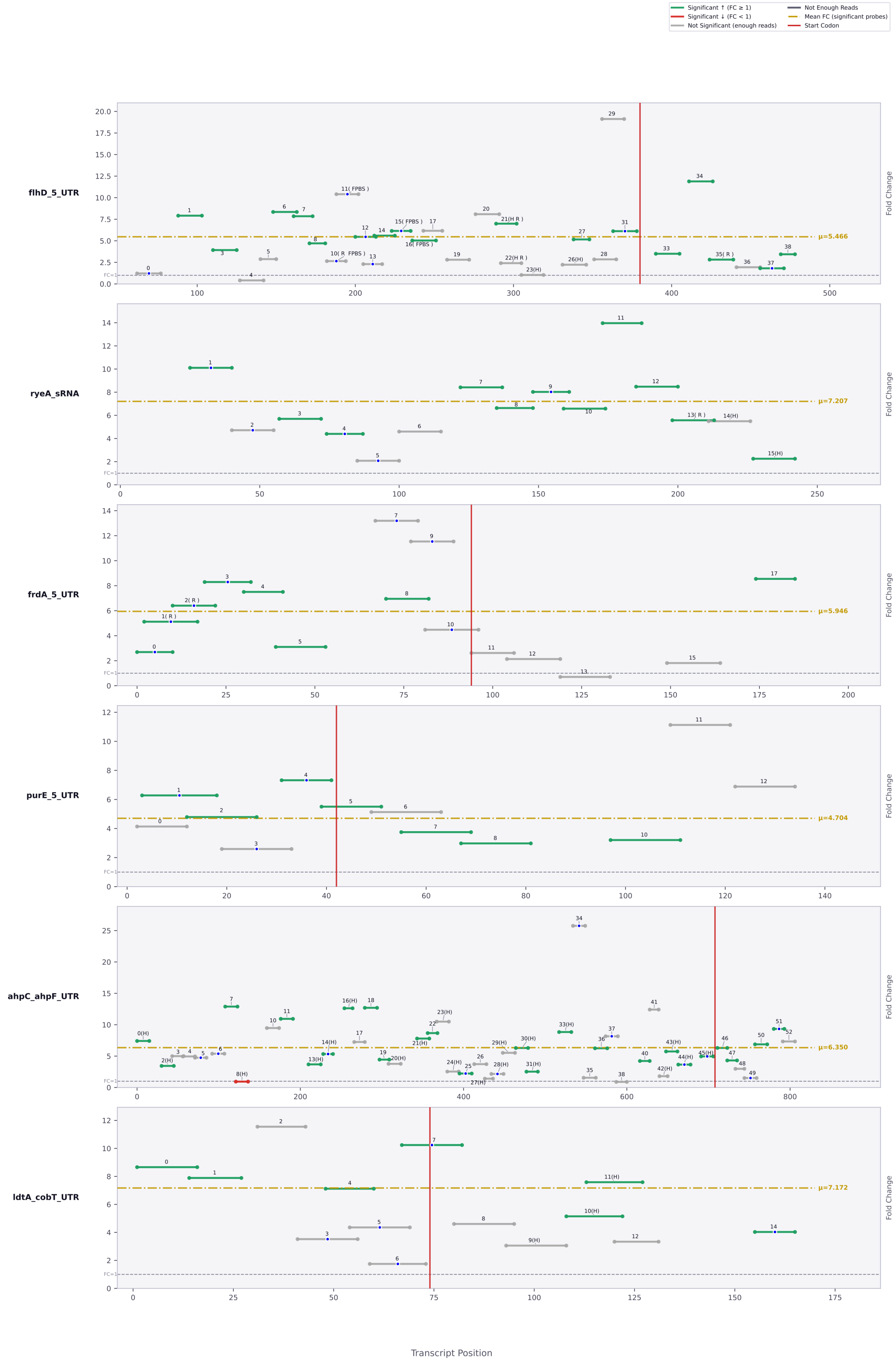

Significant ↑ (FC ≥ 1)
Significant ↓ (FC < 1)
Not Significant (enough reads)
Not Enough Reads
Mean FC (significant probes)
Start Codon
flhD_5_UTR
μ=5.466
ryeA_sRNA
μ=7.207
frdA_5_UTR
μ=5.946
purE_5_UTR
μ=4.704
ahpC_ahpF_UTR
μ=6.350
ldtA_cobT_UTR
μ=7.172
Fold Change
FC=1
Transcript Position

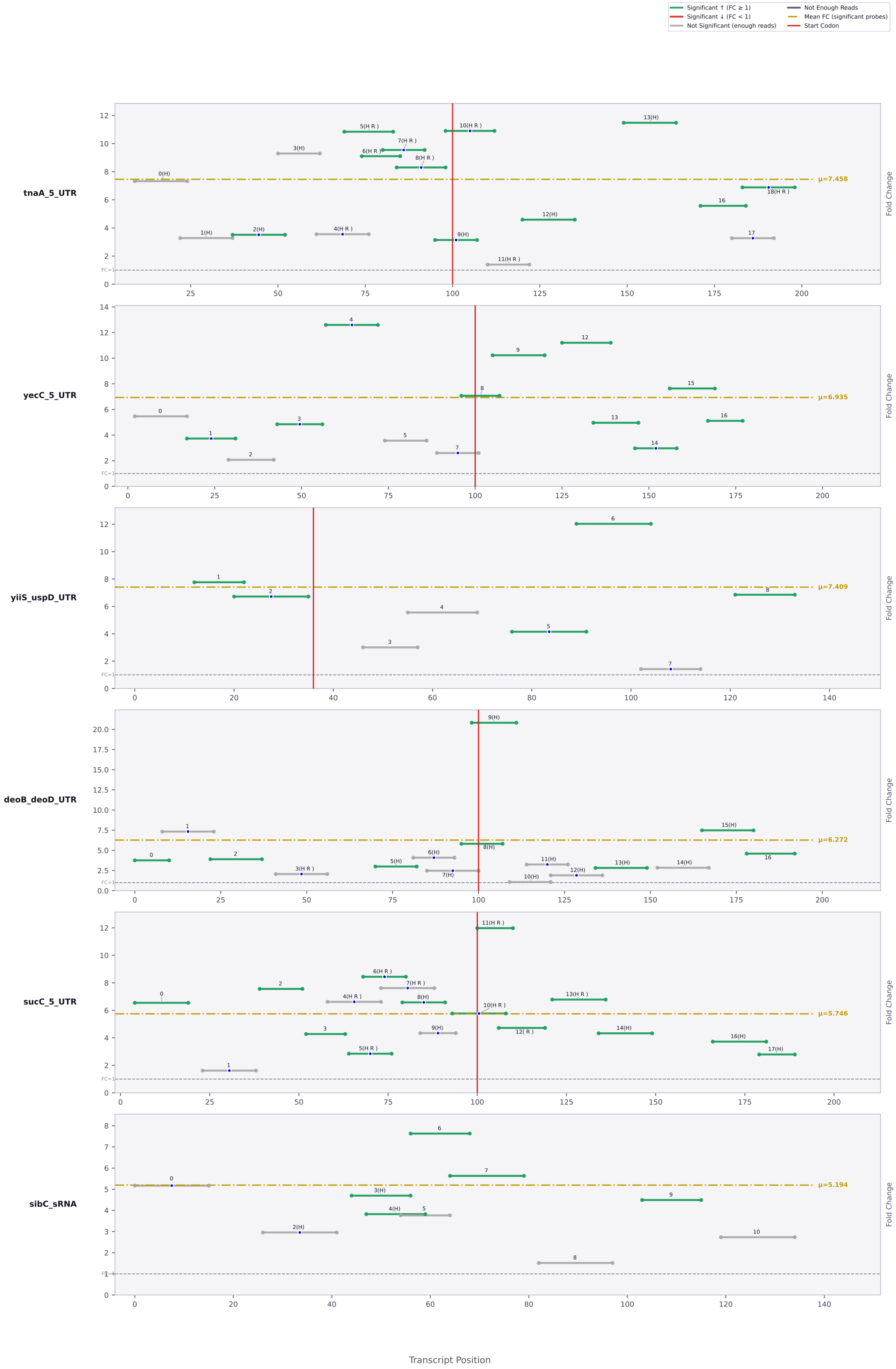

Significant ↑ (FC ≥ 1)
Significant ↓ (FC < 1)
Not Significant (enough reads)
Not Enough Reads
Mean FC (significant probes)
Start Codon
tnaA_5_UTR
μ=7.458
yecC_5_UTR
μ=6.935
yiiS_uspD_UTR
μ=7.409
deoB_deoD_UTR
μ=6.272
sucC_5_UTR
μ=5.746
sibC_sRNA
μ=5.194
Fold Change
FC=1
Transcript Position

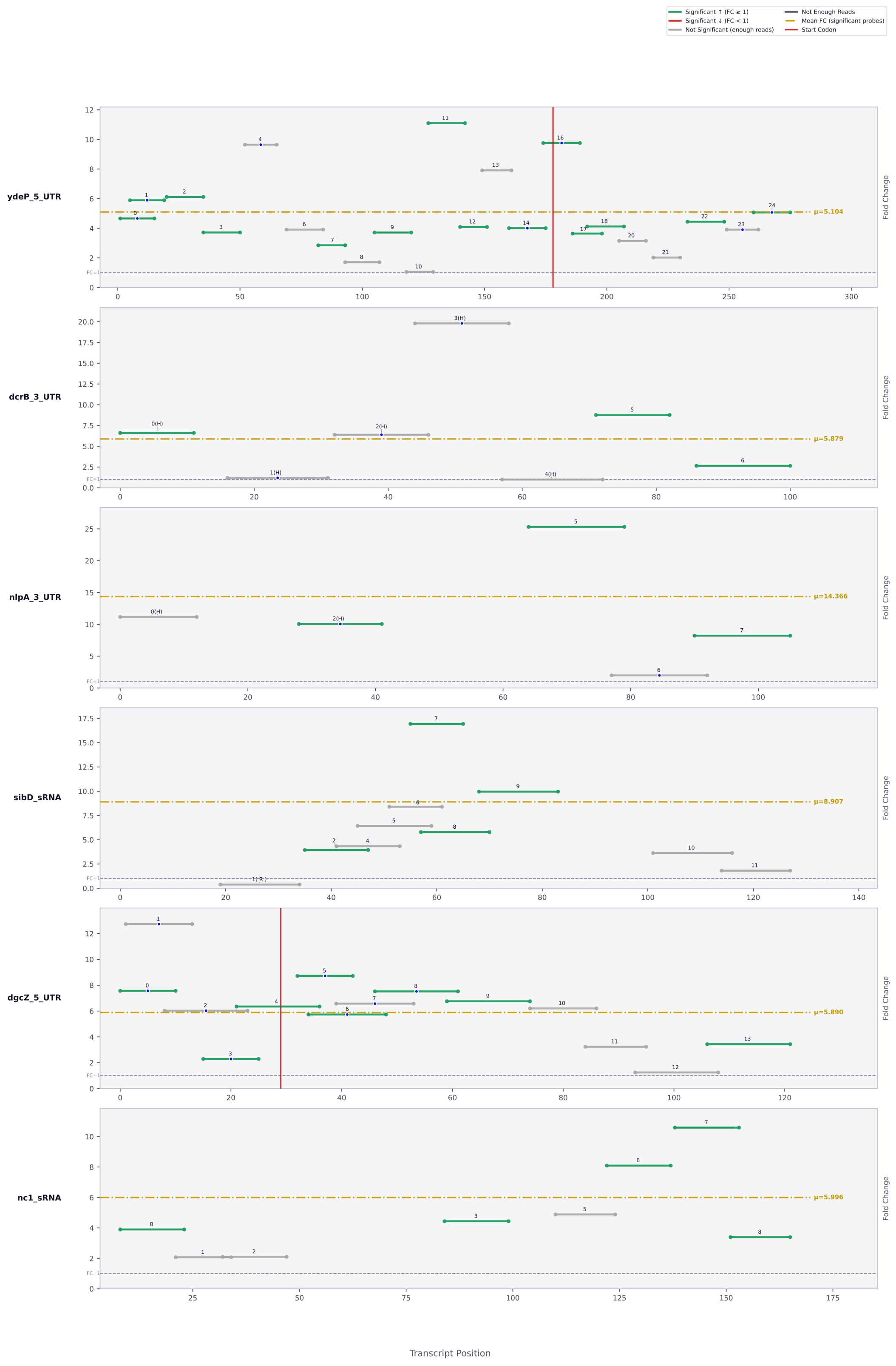

Significant ↑ (FC ≥ 1)
Significant ↓ (FC < 1)
Not Significant (enough reads)
Not Enough Reads
Mean FC (significant probes)
Start Codon
ydeP_5_UTR
μ=5.104
dcrB_3_UTR
μ=5.879
nlpA_3_UTR
μ=14.366
sibD_sRNA
μ=8.907
dgcZ_5_UTR
μ=5.890
nc1_sRNA
μ=5.996
Fold Change
FC=1
Transcript Position

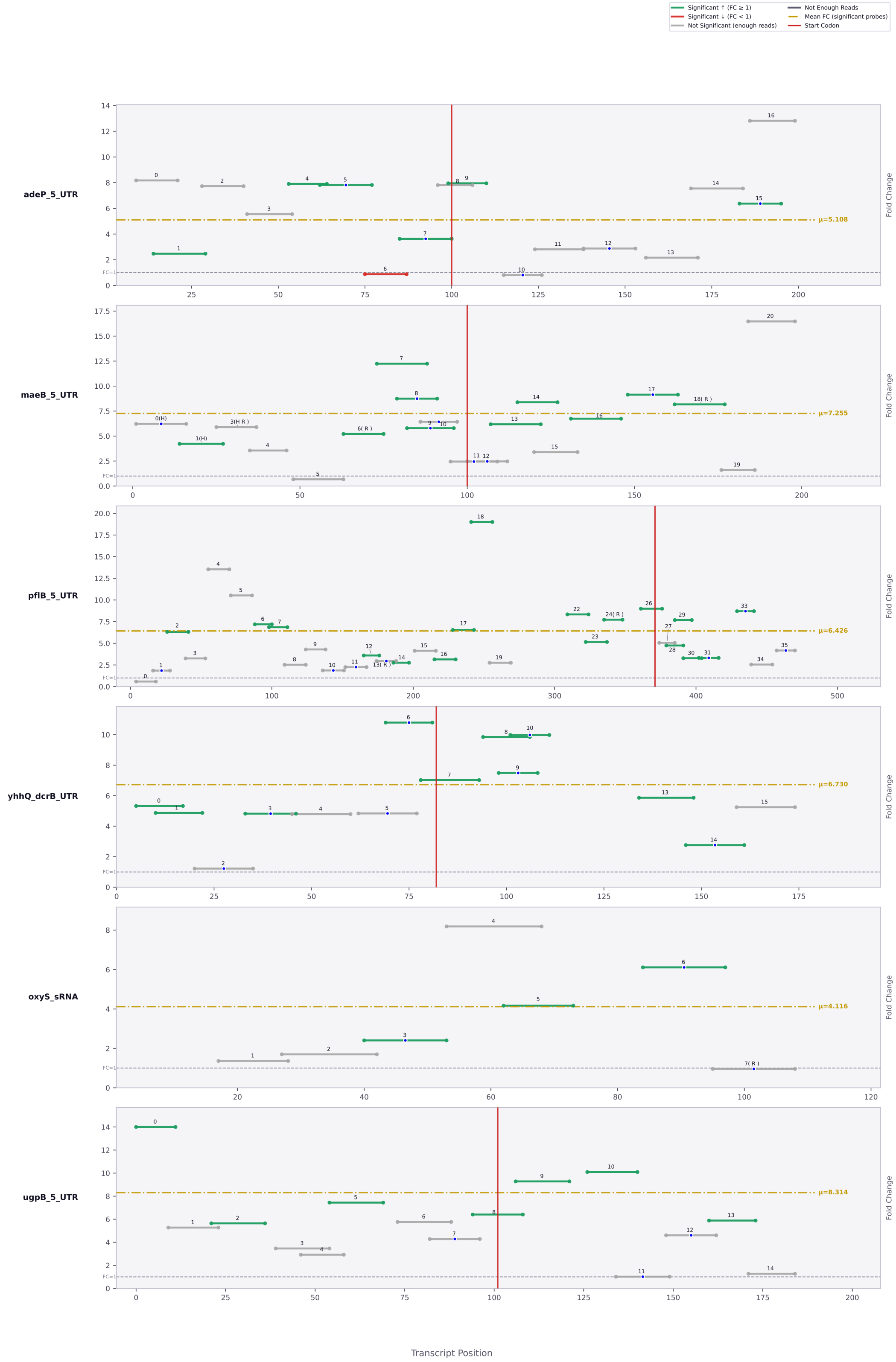

Significant ↑ (FC ≥ 1)
Significant ↓ (FC < 1)
Not Significant (enough reads)
Not Enough Reads
Mean FC (significant probes)
Start Codon
adeP_5_UTR
μ=5.108
maeB_5_UTR
μ=7.255
pflB_5_UTR
μ=6.426
yhhQ_dcrB_UTR
μ=6.730
oxyS_sRNA
μ=4.116
ugpB_5_UTR
μ=8.314
Fold Change
Transcript Position

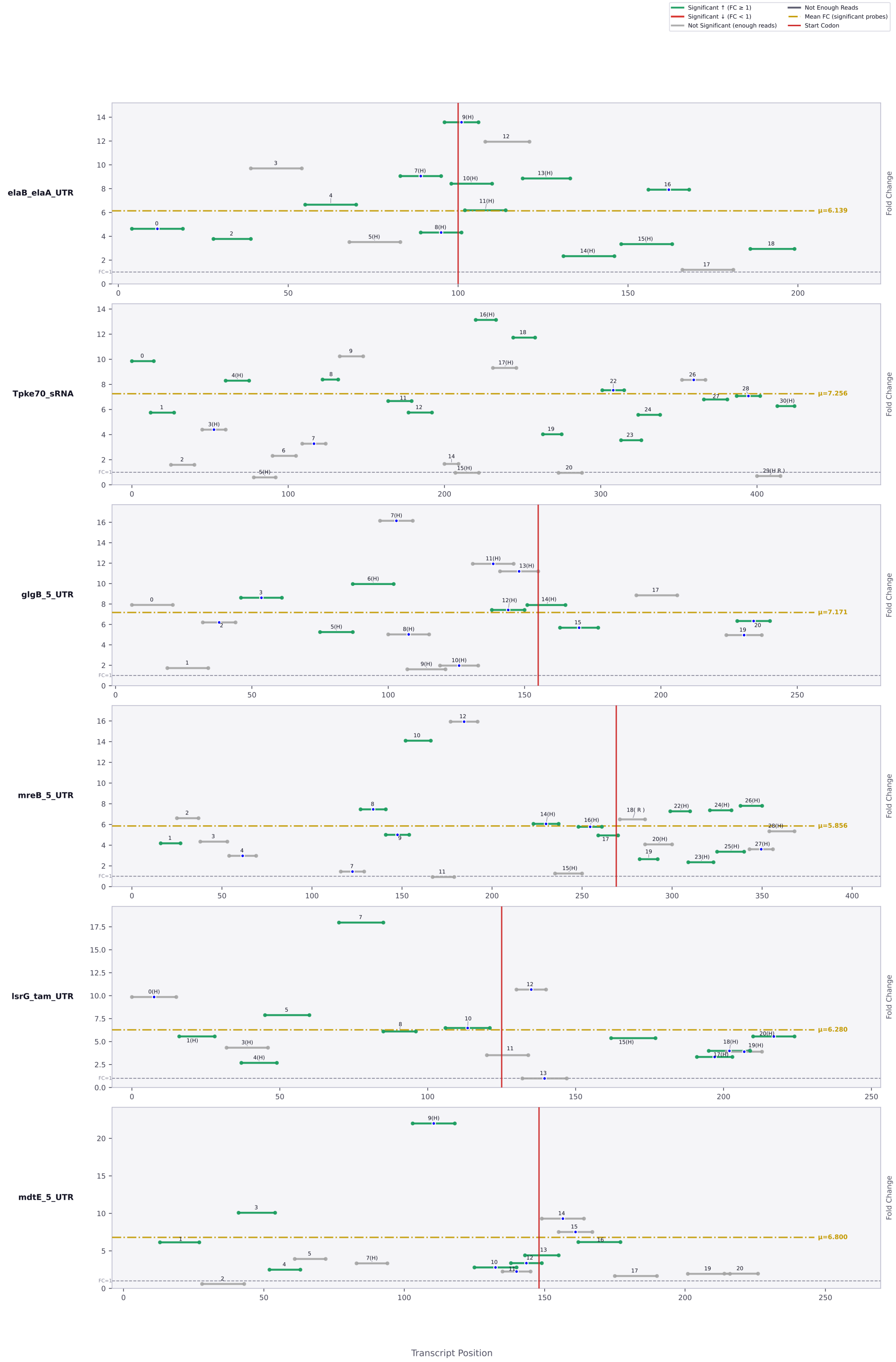
Significant ↑ (FC ≥ 1)
Significant ↓ (FC < 1)
Not Significant (enough reads)
Not Enough Reads
Mean FC (significant probes)
Start Codon
elaB_elaA_UTR
μ=6.139
Tpke70_sRNA
μ=7.256
glgB_5_UTR
μ=7.171
mreB_5_UTR
μ=5.856
lsrG_tam_UTR
μ=6.280
mdtE_5_UTR
μ=6.800
Fold Change
FC=1
Transcript Position

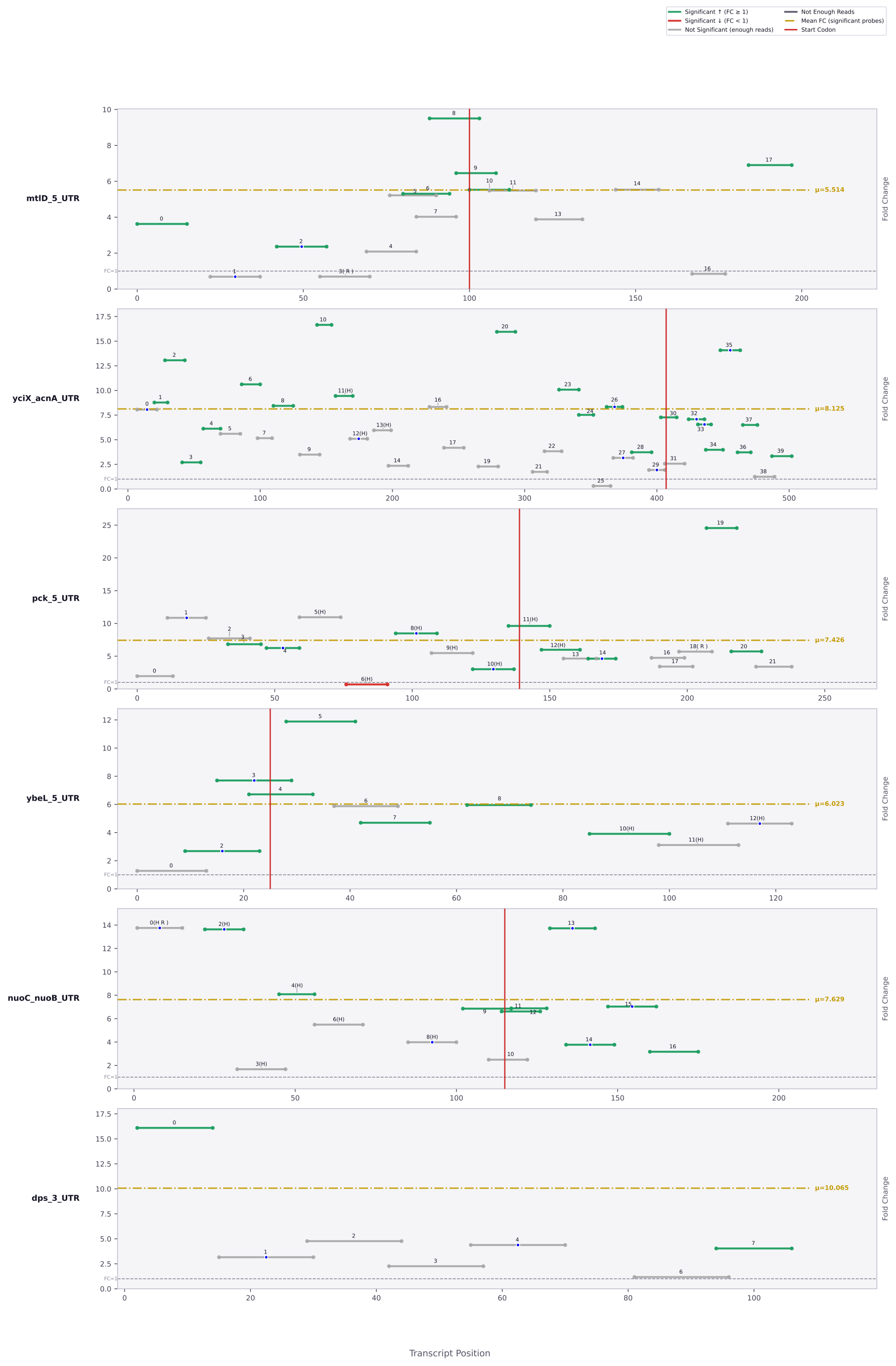

Significant ↑ (FC ≥ 1)
Significant ↓ (FC < 1)
Not Significant (enough reads)
Not Enough Reads
Mean FC (significant probes)
Start Codon
mtlD_5_UTR
μ=5.514
yciX_acnA_UTR
μ=8.125
pck_5_UTR
μ=7.426
ybeL_5_UTR
μ=6.023
nuoC_nuoB_UTR
μ=7.629
dps_3_UTR
μ=10.065
Fold Change
FC=1
Transcript Position

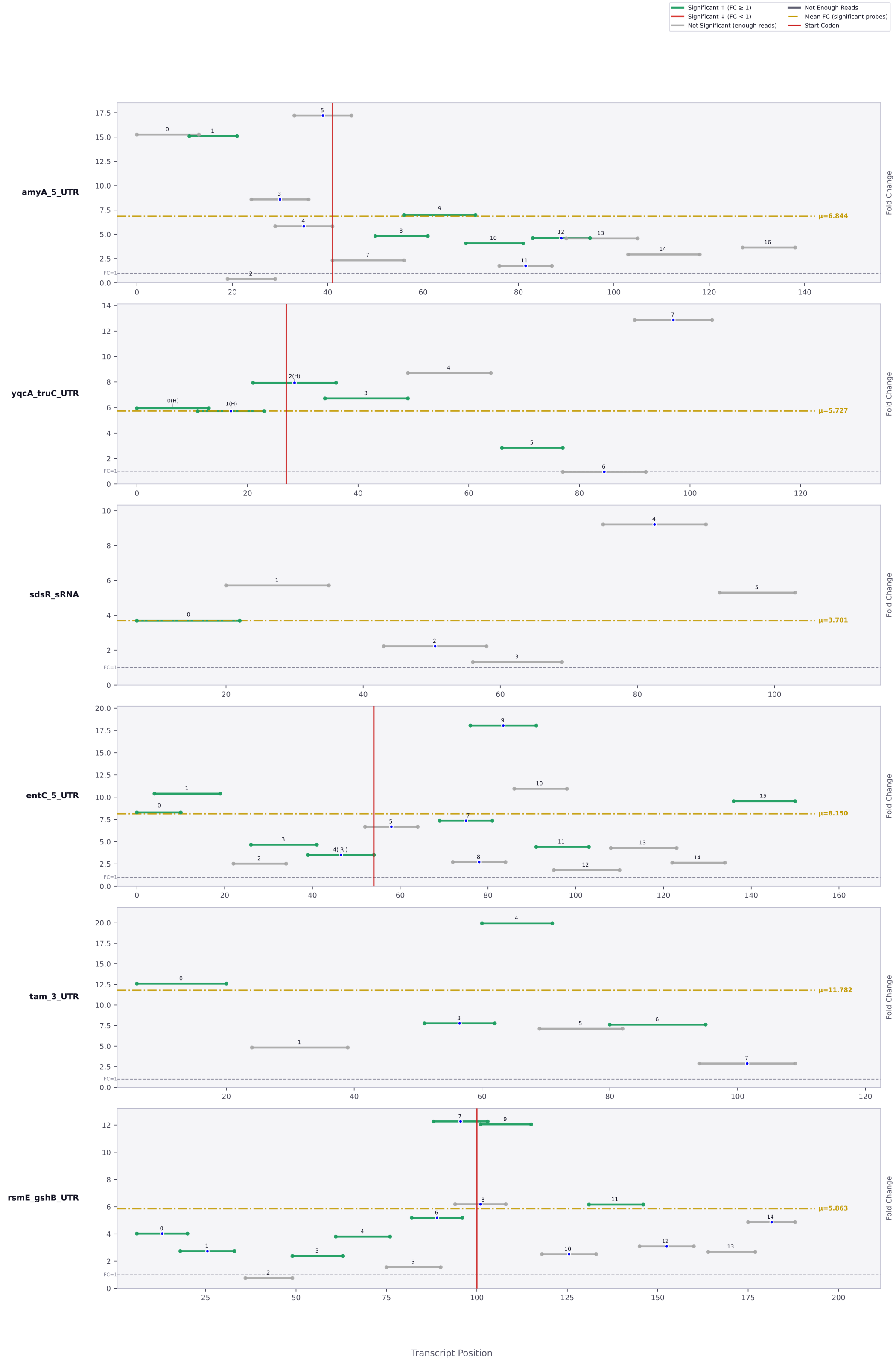

Significant ↑ (FC ≥ 1)
Significant ↓ (FC < 1)
Not Significant (enough reads)
Not Enough Reads
Mean FC (significant probes)
Start Codon
amyA_5_UTR
μ=6.844
yqcA_truC_UTR
μ=5.727
sdsR_sRNA
μ=3.701
entC_5_UTR
μ=8.150
tam_3_UTR
μ=11.782
rsmE_gshB_UTR
μ=5.863
Fold Change
FC=1
Transcript Position

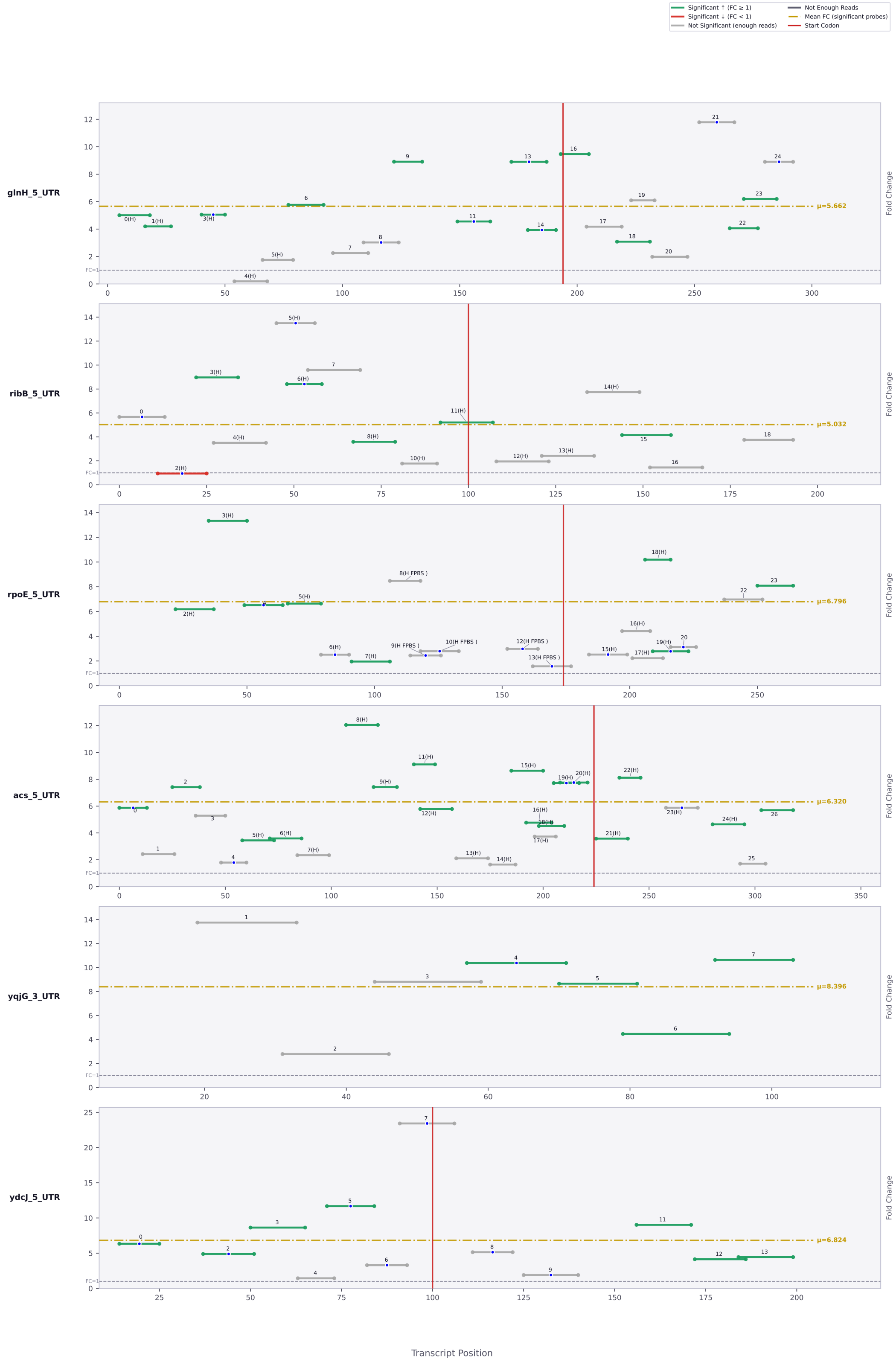

Significant ↑ (FC ≥ 1)
Significant ↓ (FC < 1)
Not Significant (enough reads)
Not Enough Reads
Mean FC (significant probes)
Start Codon
glnH_5_UTR
μ=5.662
ribB_5_UTR
μ=5.032
rpoE_5_UTR
μ=6.796
acs_5_UTR
μ=6.320
yqjG_3_UTR
μ=8.396
ydcJ_5_UTR
μ=6.824
Fold Change
FC=1
Transcript Position

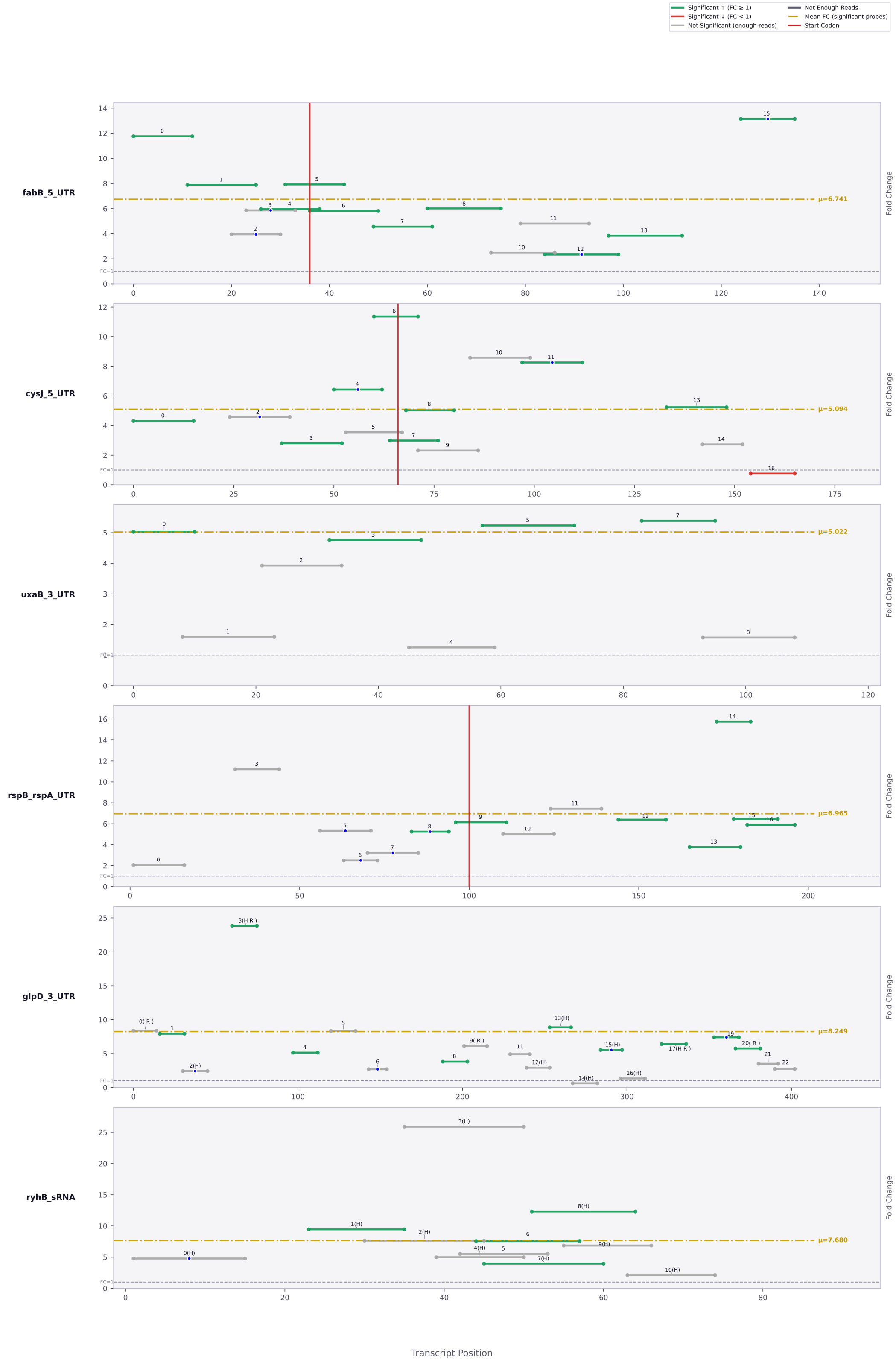

Significant ↑ (FC ≥ 1)
Significant ↓ (FC < 1)
Not Significant (enough reads)
Not Enough Reads
Mean FC (significant probes)
Start Codon
fabB_5_UTR
μ=6.741
cysJ_5_UTR
μ=5.094
uxaB_3_UTR
μ=5.022
rspB_rspA_UTR
μ=6.965
glpD_3_UTR
μ=8.249
ryhB_sRNA
μ=7.680
Fold Change
FC=1
Transcript Position

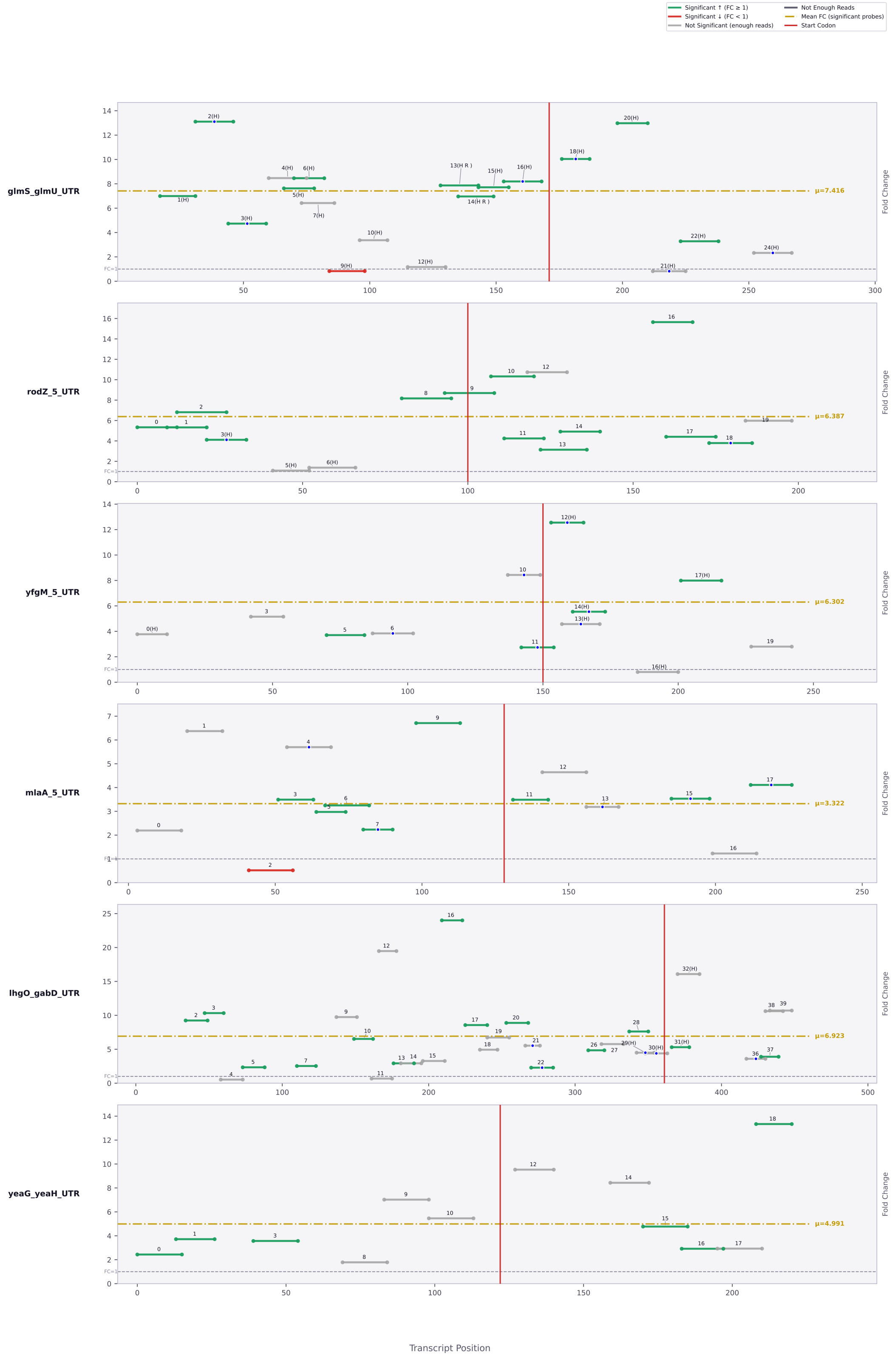

Significant ↑ (FC ≥ 1)
Significant ↓ (FC < 1)
Not Significant (enough reads)
Not Enough Reads
Mean FC (significant probes)
Start Codon
glmS_glmU_UTR
μ=7.416
rodZ_5_UTR
μ=6.387
yfgM_5_UTR
μ=6.302
mlaA_5_UTR
μ=3.322
lhgO_gabD_UTR
μ=6.923
yeaG_yeaH_UTR
μ=4.991
Fold Change
Transcript Position

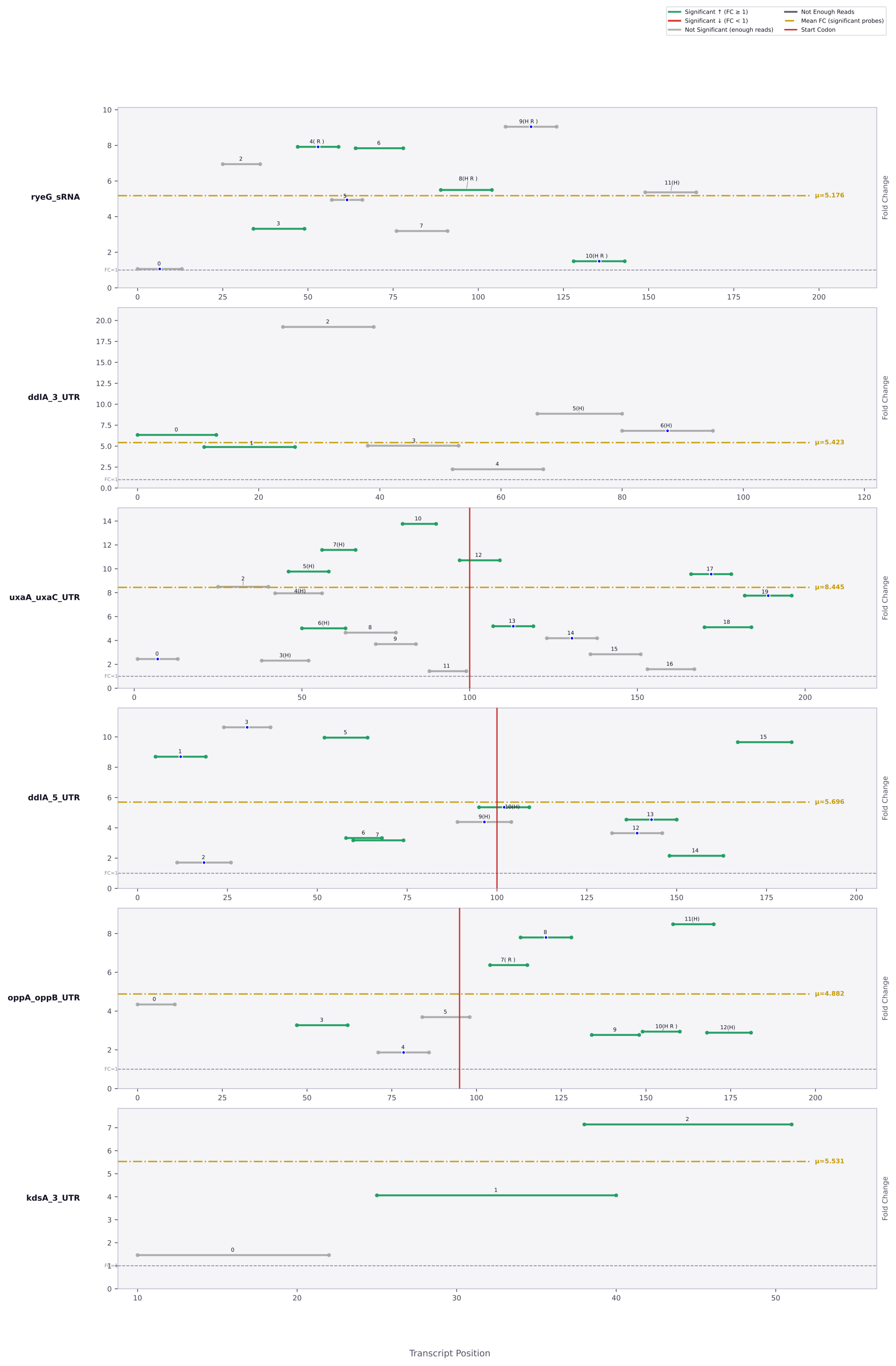

Significant ↑ (FC ≥ 1)
Significant ↓ (FC < 1)
Not Significant (enough reads)
Not Enough Reads
Mean FC (significant probes)
Start Codon
ryeG_sRNA
μ=5.176
ddlA_3_UTR
μ=5.423
uxaA_uxaC_UTR
μ=8.445
ddlA_5_UTR
μ=5.696
oppA_oppB_UTR
μ=4.882
kdsA_3_UTR
μ=5.531
Fold Change
Transcript Position

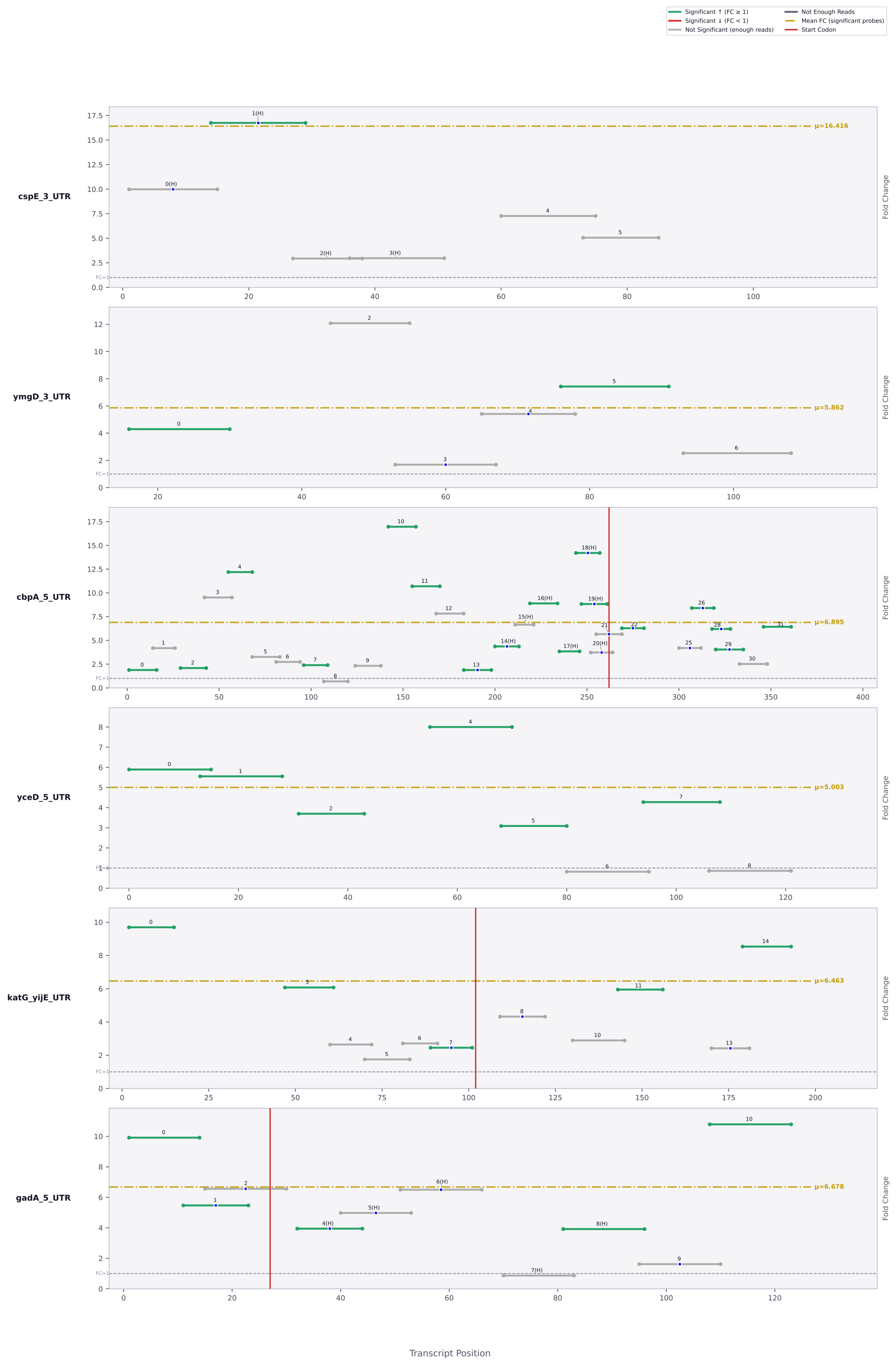

Significant ↑ (FC ≥ 1)
Significant ↓ (FC < 1)
Not Significant (enough reads)
Not Enough Reads
Mean FC (significant probes)
Start Codon
cspE_3_UTR
μ=16.416
ymgD_3_UTR
μ=5.862
cbpA_5_UTR
μ=6.895
yceD_5_UTR
μ=5.003
katG_yijE_UTR
μ=6.463
gadA_5_UTR
μ=6.678
FC=1
Fold Change
Transcript Position

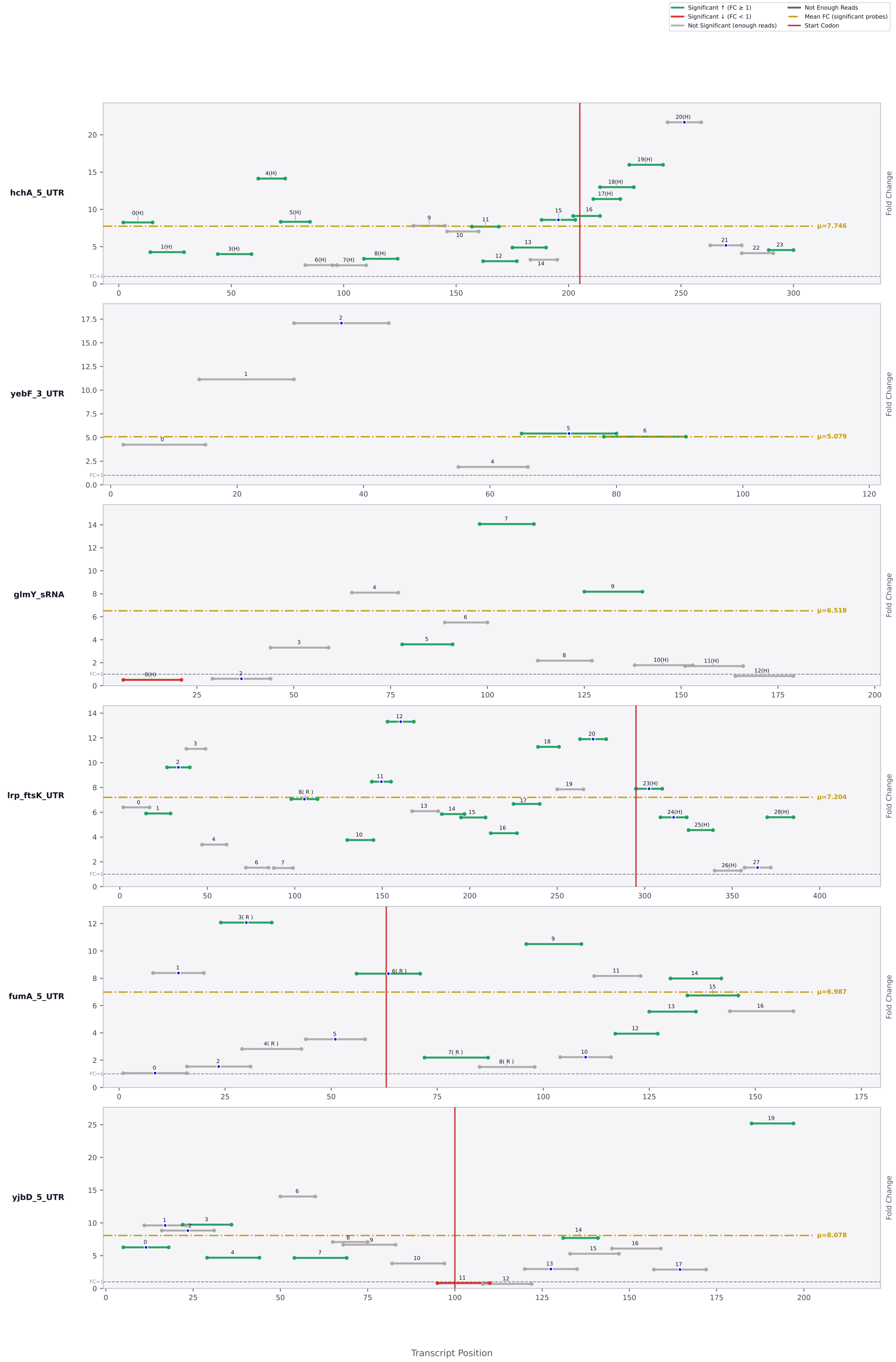

Significant ↑ (FC ≥ 1)
Significant ↓ (FC < 1)
Not Significant (enough reads)
Not Enough Reads
Mean FC (significant probes)
Start Codon
hchA_5_UTR
μ=7.746
yebF_3_UTR
μ=5.079
glmY_sRNA
μ=6.518
lrp_ftsK_UTR
μ=7.204
fumA_5_UTR
μ=6.987
yjbD_5_UTR
μ=8.078
Fold Change
FC=1
Transcript Position

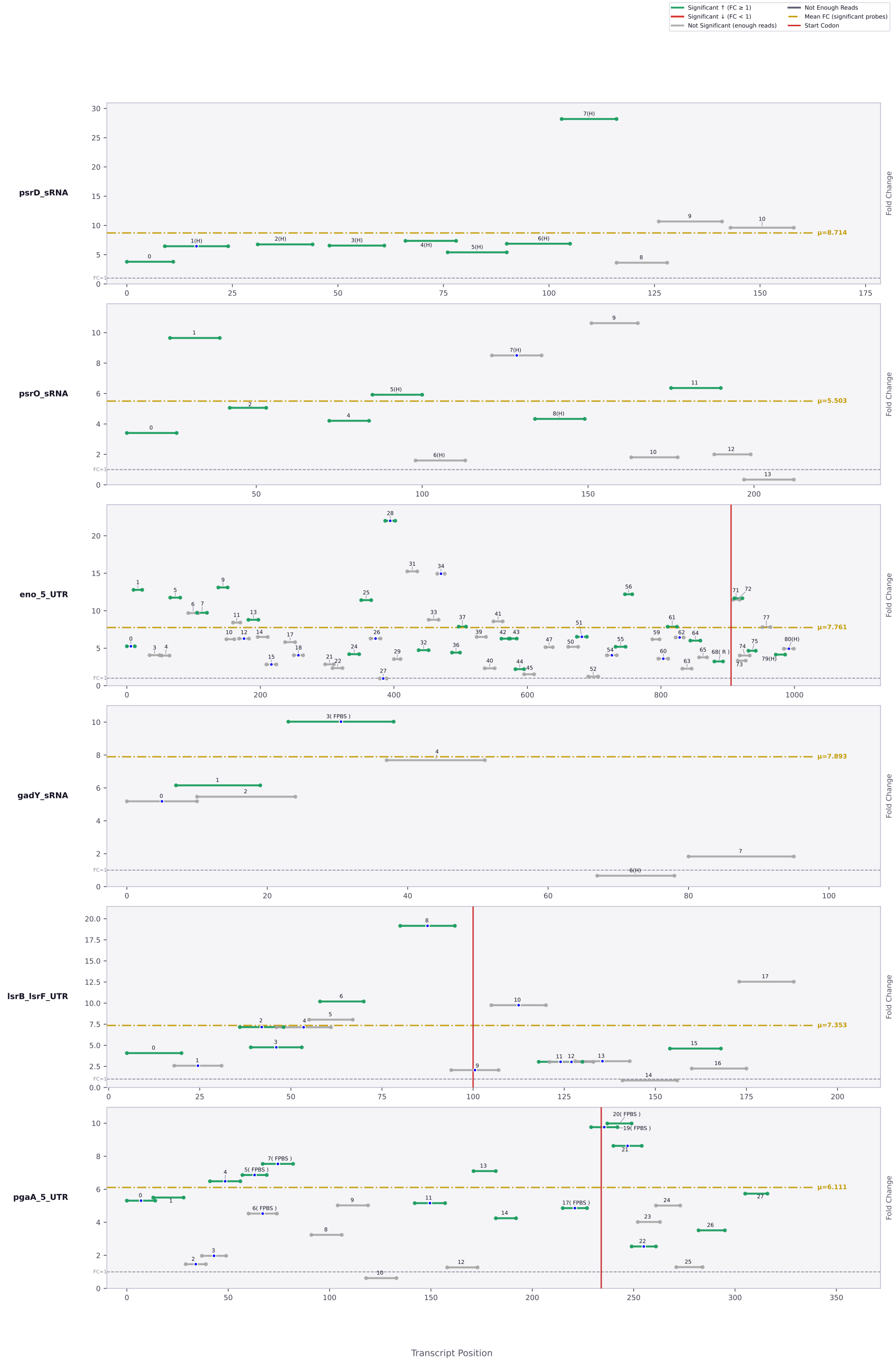

Significant ↑ (FC ≥ 1)
Significant ↓ (FC < 1)
Not Significant (enough reads)
Not Enough Reads
Mean FC (significant probes)
Start Codon
psrD_sRNA
μ=8.714
psrO_sRNA
μ=5.503
eno_5_UTR
μ=7.761
gadY_sRNA
μ=7.893
lsrB_lsrF_UTR
μ=7.353
pgaA_5_UTR
μ=6.111
Fold Change
FC=1
Transcript Position

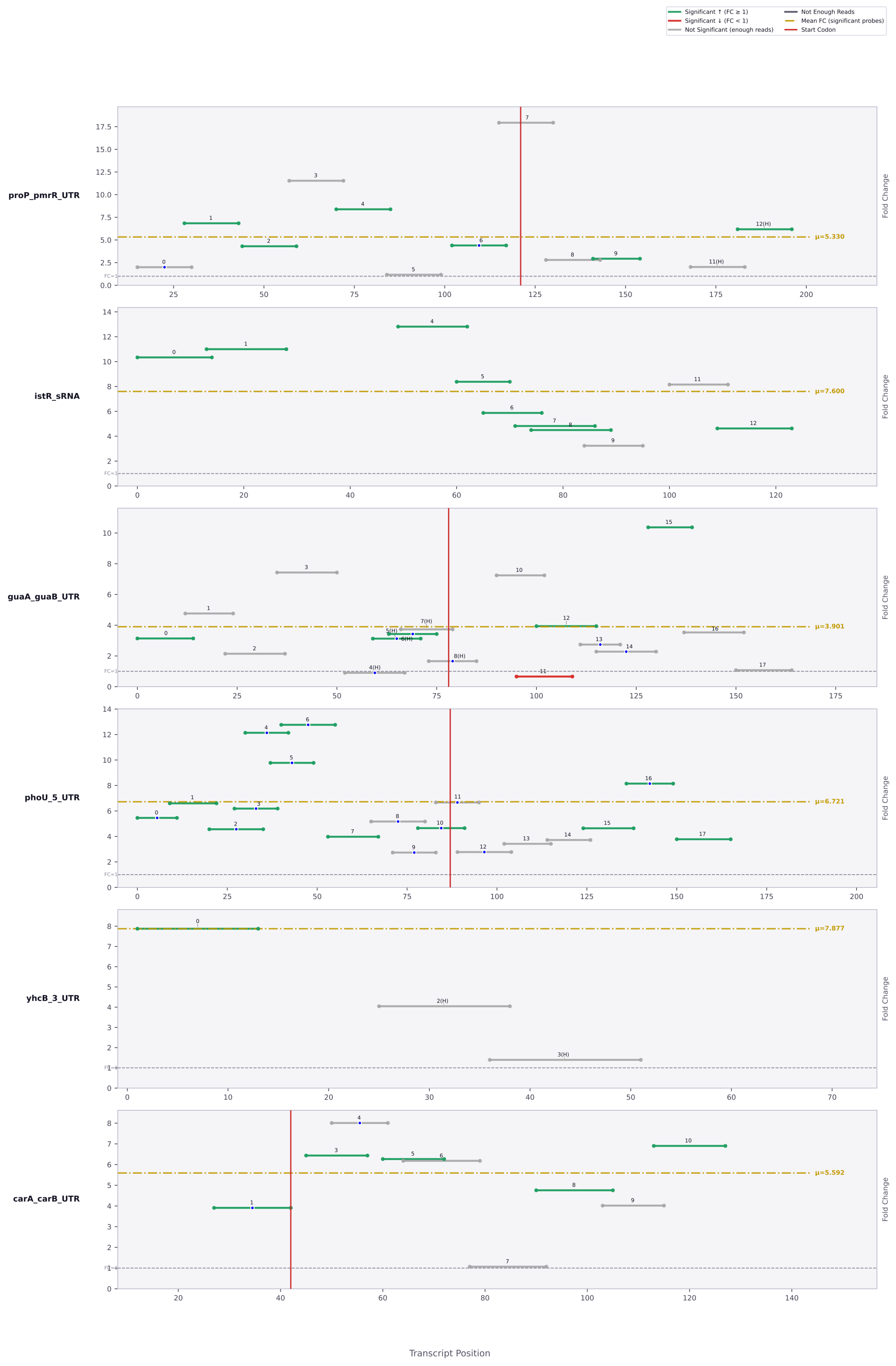

Significant ↑ (FC ≥ 1)
Significant ↓ (FC < 1)
Not Significant (enough reads)
Not Enough Reads
Mean FC (significant probes)
Start Codon
proP_pmrR_UTR
μ=5.330
istR_sRNA
μ=7.600
guaA_guaB_UTR
μ=3.901
phoU_5_UTR
μ=6.721
yhcB_3_UTR
μ=7.877
carA_carB_UTR
μ=5.592
FC=1
Fold Change
Transcript Position

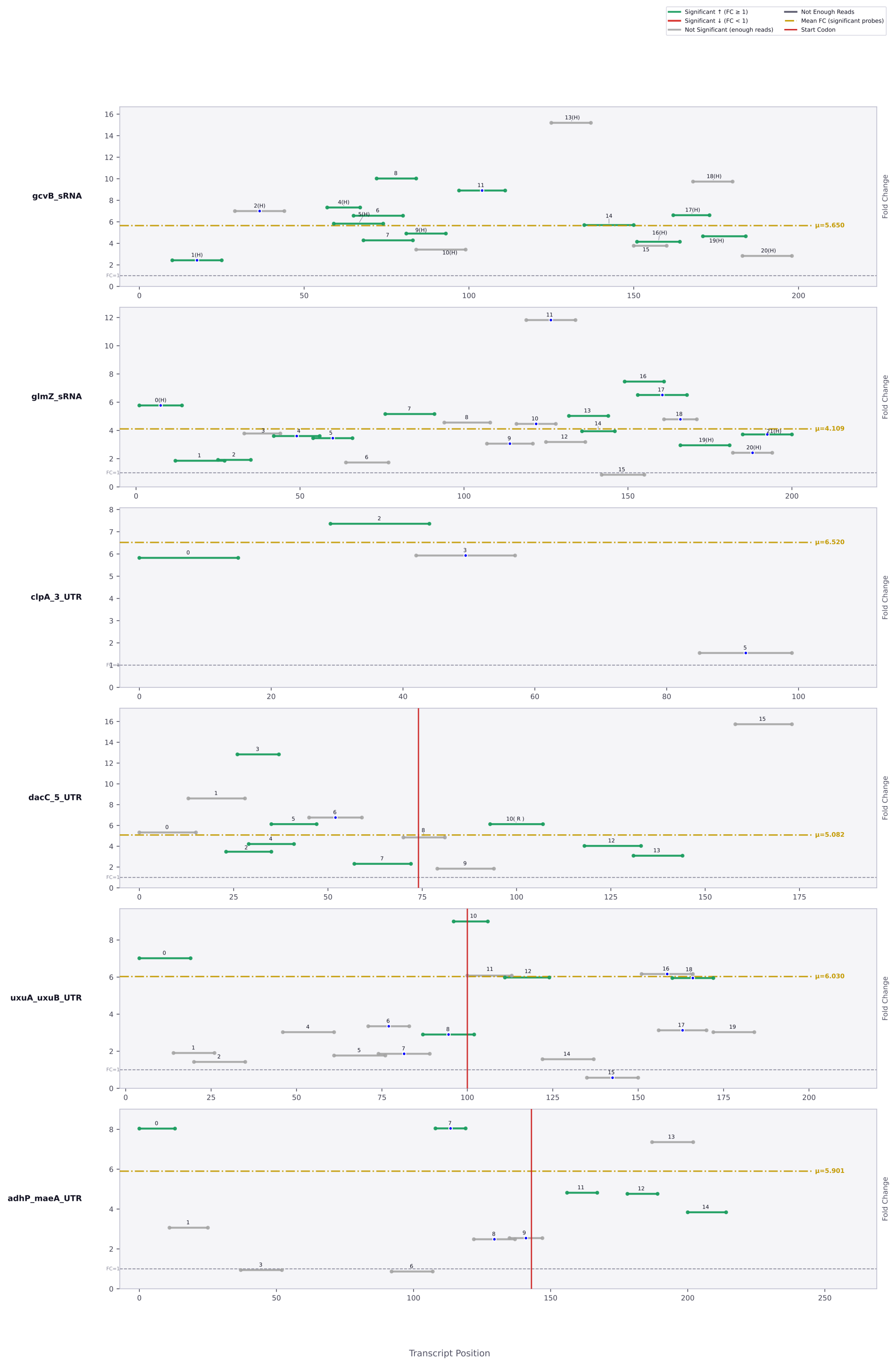

Significant ↑ (FC ≥ 1)
Significant ↓ (FC < 1)
Not Significant (enough reads)
Not Enough Reads
Mean FC (significant probes)
Start Codon
gcvB_sRNA
μ=5.650
glmZ_sRNA
μ=4.109
clpA_3_UTR
μ=6.520
dacC_5_UTR
μ=5.082
uxuA_uxuB_UTR
μ=6.030
adhP_maeA_UTR
μ=5.901
Fold Change
Transcript Position

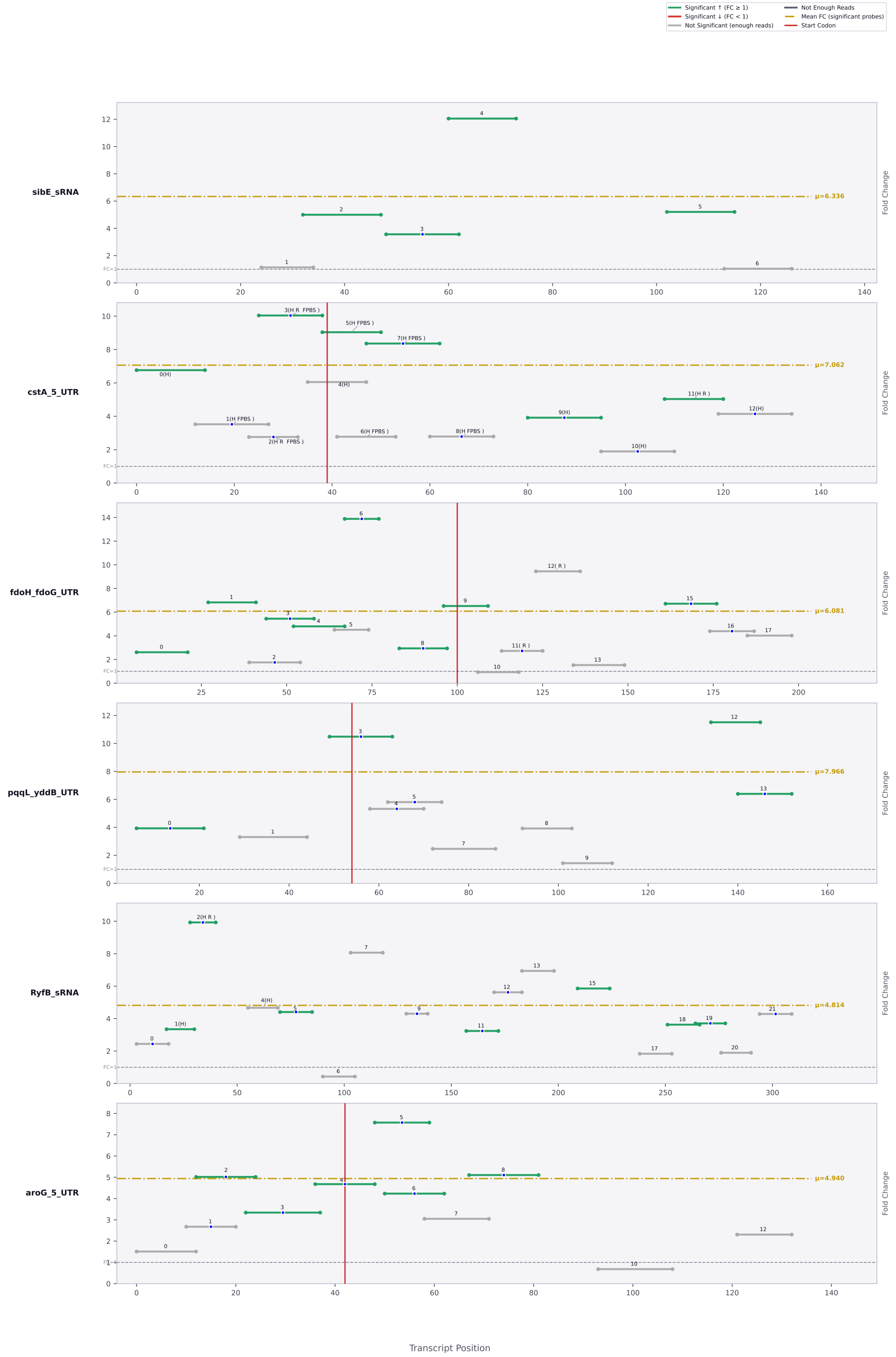

Significant ↑ (FC ≥ 1)
Significant ↓ (FC < 1)
Not Significant (enough reads)
Not Enough Reads
Mean FC (significant probes)
Start Codon
sibE_sRNA
μ=6.336
cstA_5_UTR
μ=7.062
fdoH_fdoG_UTR
μ=6.081
pqqL_yddB_UTR
μ=7.966
RyfB_sRNA
μ=4.814
aroG_5_UTR
μ=4.940
Fold Change
FC=1
Transcript Position

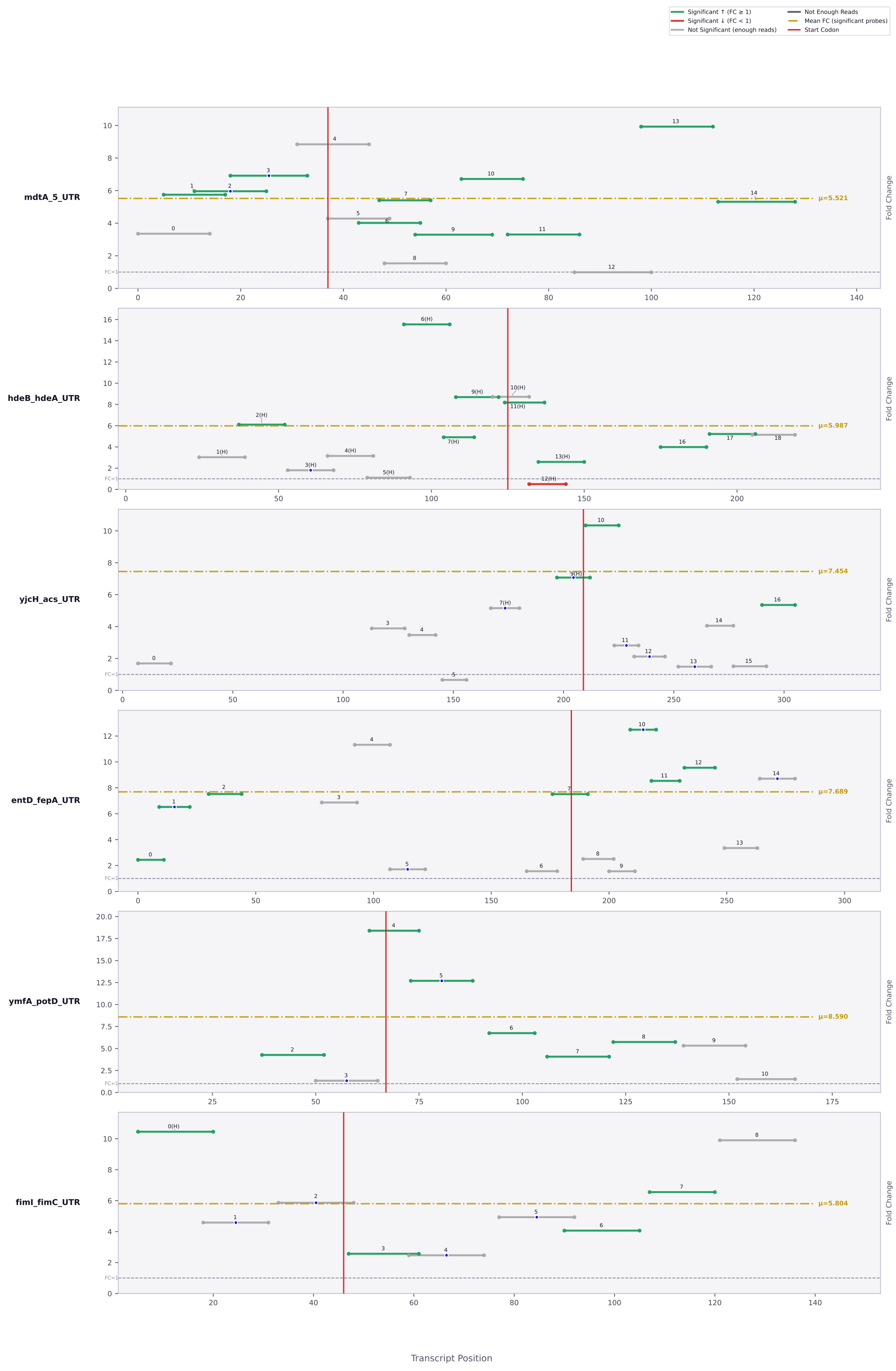

Significant ↑ (FC ≥ 1)
Significant ↓ (FC < 1)
Not Significant (enough reads)
Not Enough Reads
Mean FC (significant probes)
Start Codon
mdtA_5_UTR
μ=5.521
hdeB_hdeA_UTR
μ=5.987
yjcH_acs_UTR
μ=7.454
entD_fepA_UTR
μ=7.689
ymfA_potD_UTR
μ=8.590
fimI_fimC_UTR
μ=5.804
Fold Change
FC=1
Transcript Position

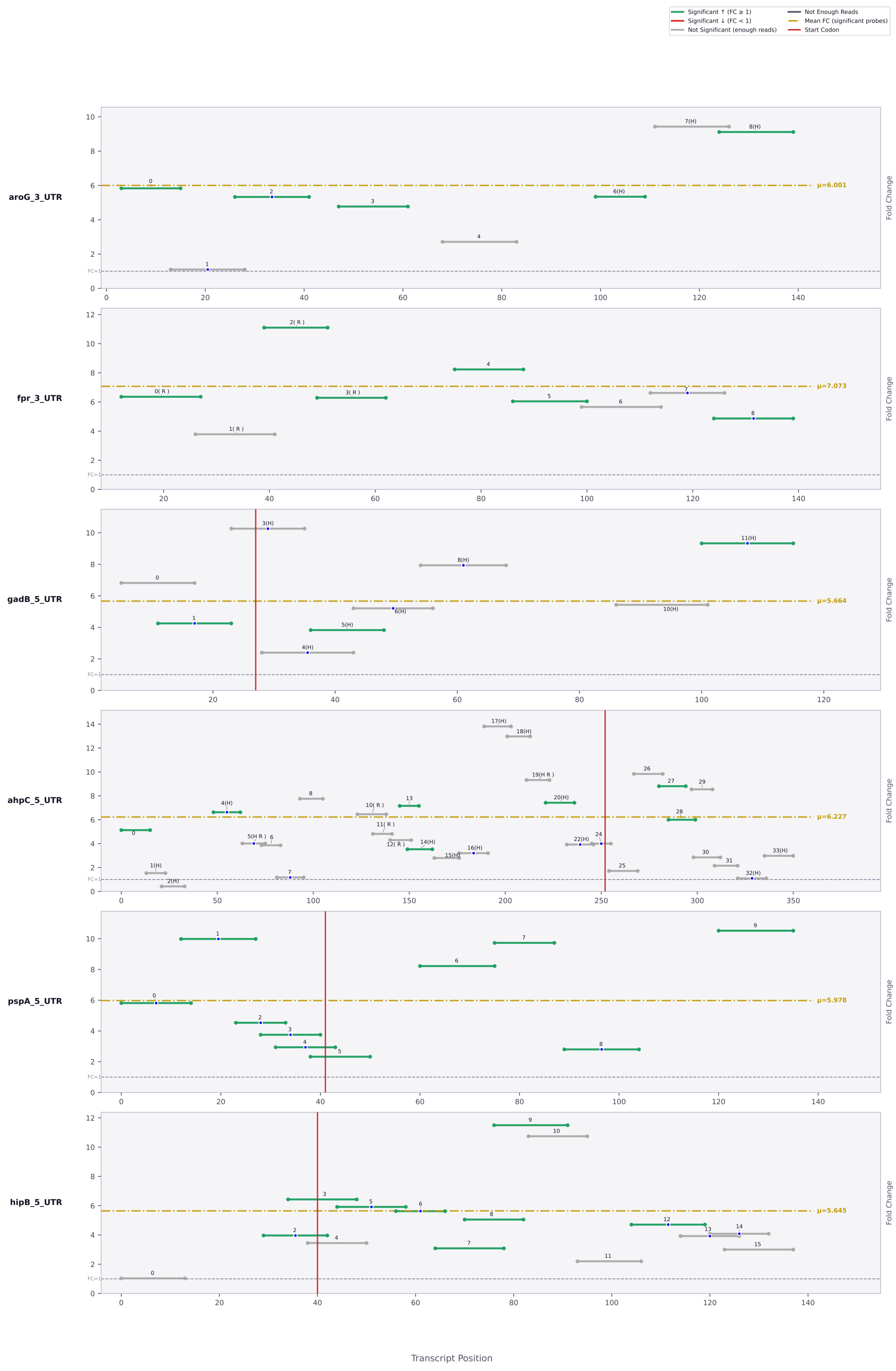

Significant ↑ (FC ≥ 1)
Significant ↓ (FC < 1)
Not Significant (enough reads)
Not Enough Reads
Mean FC (significant probes)
Start Codon
aroG_3_UTR
μ=6.001
fpr_3_UTR
μ=7.073
gadB_5_UTR
μ=5.664
ahpC_5_UTR
μ=6.227
pspA_5_UTR
μ=5.978
hipB_5_UTR
μ=5.645
Fold Change
Transcript Position

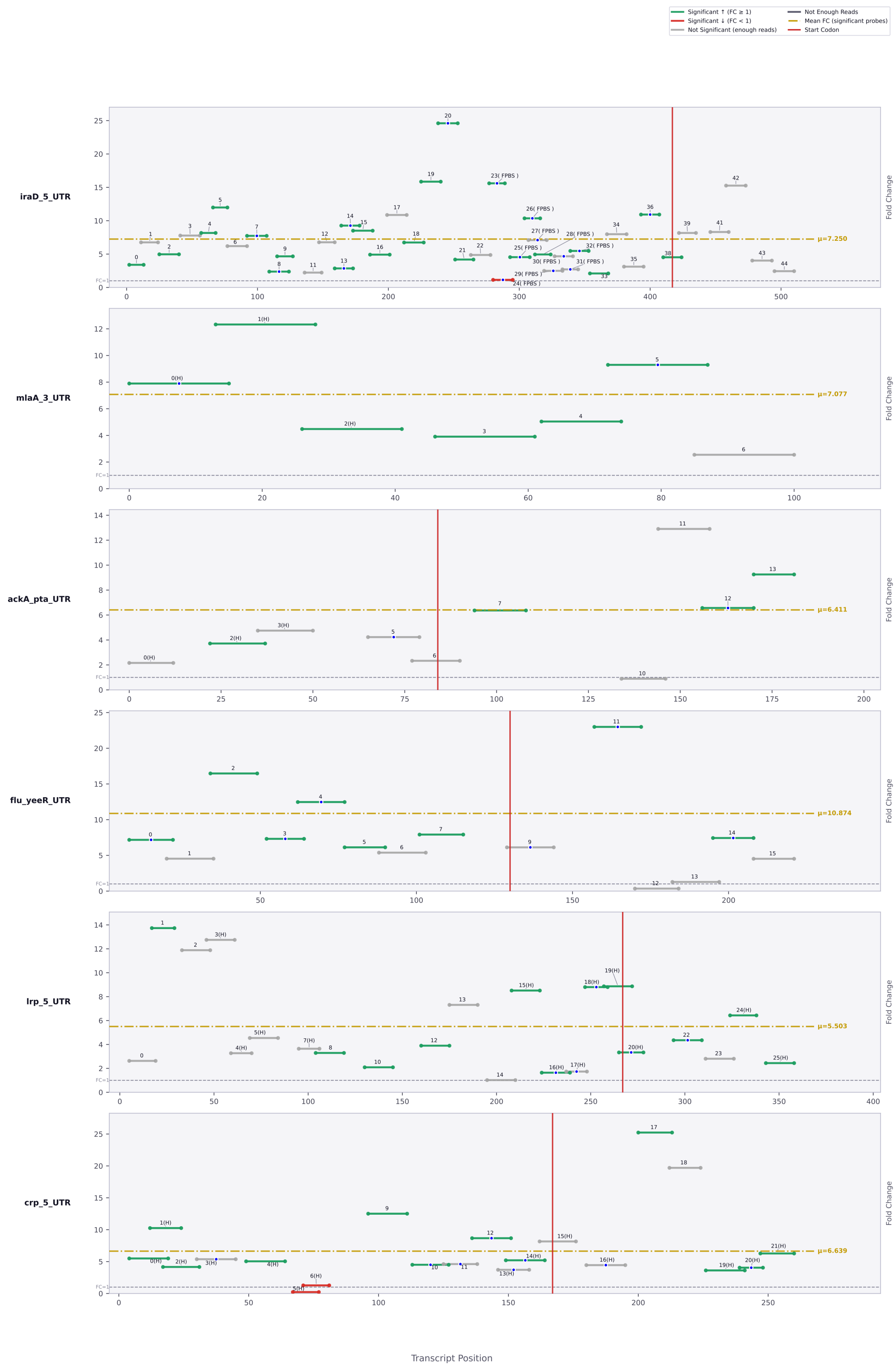

Significant ↑ (FC ≥ 1)
Significant ↓ (FC < 1)
Not Significant (enough reads)
Not Enough Reads
Mean FC (significant probes)
Start Codon
iraD_5_UTR
μ=7.250
mlaA_3_UTR
μ=7.077
ackA_pta_UTR
μ=6.411
flu_yeeR_UTR
μ=10.874
lrp_5_UTR
μ=5.503
crp_5_UTR
μ=6.639
Fold Change
Transcript Position

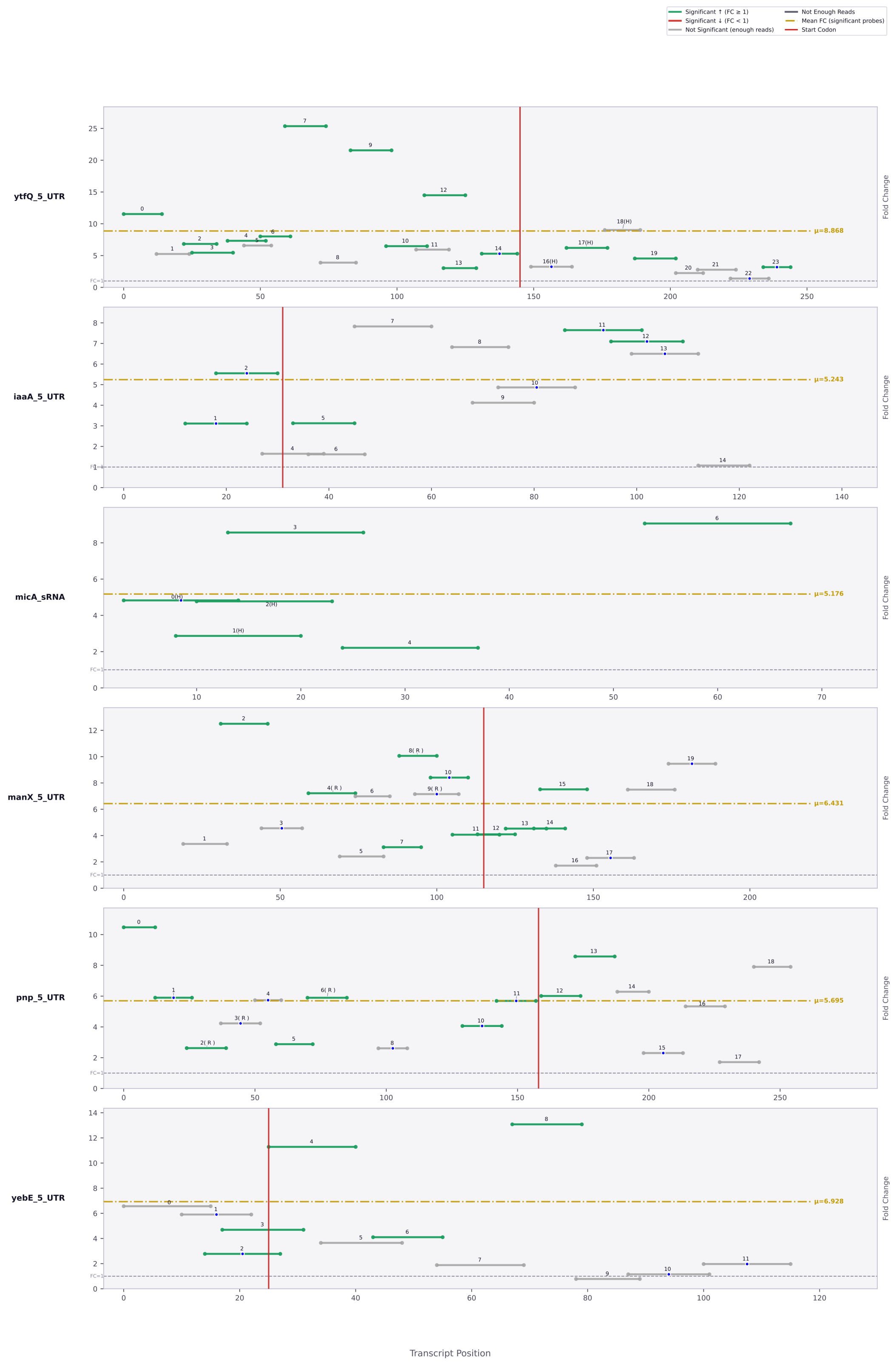

Significant ↑ (FC ≥ 1)
Significant ↓ (FC < 1)
Not Significant (enough reads)
Not Enough Reads
Mean FC (significant probes)
Start Codon
ytfQ_5_UTR
μ=8.868
iaaA_5_UTR
μ=5.243
micA_sRNA
μ=5.176
manX_5_UTR
μ=6.431
pnp_5_UTR
μ=5.695
yebE_5_UTR
μ=6.928
FC=1
Fold Change
Transcript Position

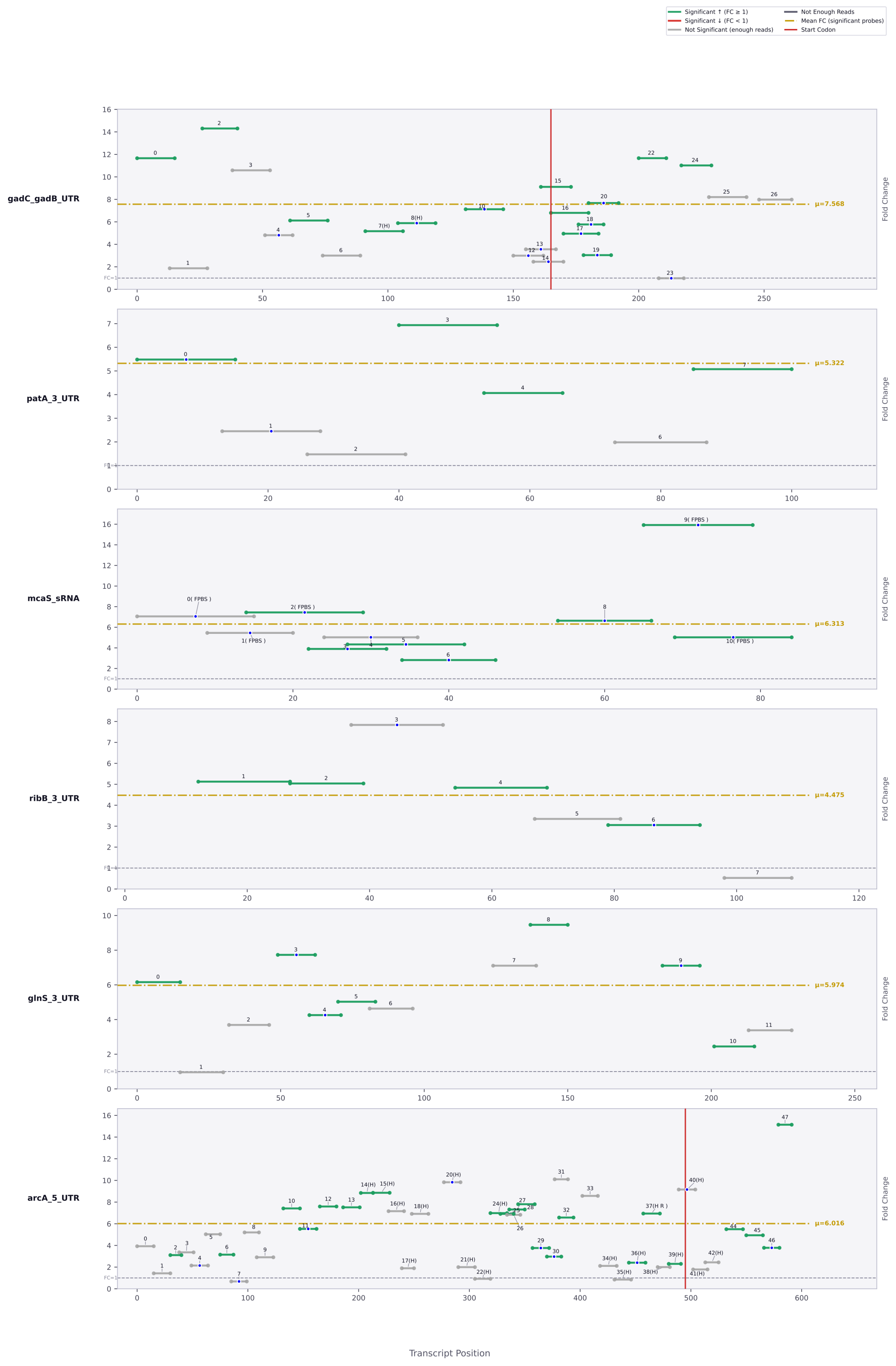

Significant ↑ (FC ≥ 1)
Significant ↓ (FC < 1)
Not Significant (enough reads)
Not Enough Reads
Mean FC (significant probes)
Start Codon
gadC_gadB_UTR
μ=7.568
patA_3_UTR
μ=5.322
mcaS_sRNA
μ=6.313
ribB_3_UTR
μ=4.475
glnS_3_UTR
μ=5.974
arcA_5_UTR
μ=6.016
Fold Change
FC=1
Transcript Position

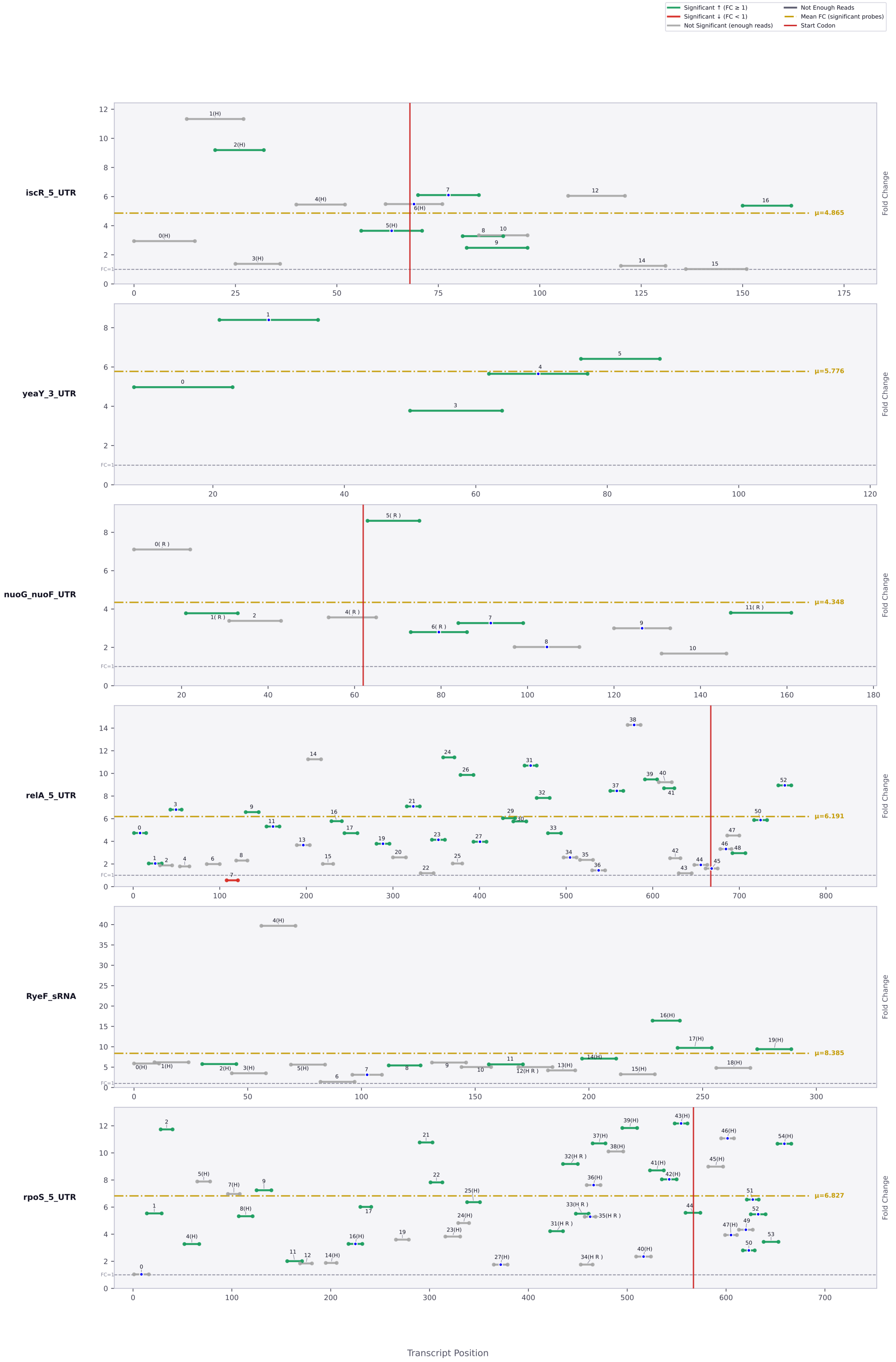

Significant ↑ (FC ≥ 1)
Significant ↓ (FC < 1)
Not Significant (enough reads)
Not Enough Reads
Mean FC (significant probes)
Start Codon
iscR_5_UTR
μ=4.865
yeaY_3_UTR
μ=5.776
nuoG_nuoF_UTR
μ=4.348
relA_5_UTR
μ=6.191
RyeF_sRNA
μ=8.385
rpoS_5_UTR
μ=6.827
Fold Change
FC=1
Transcript Position

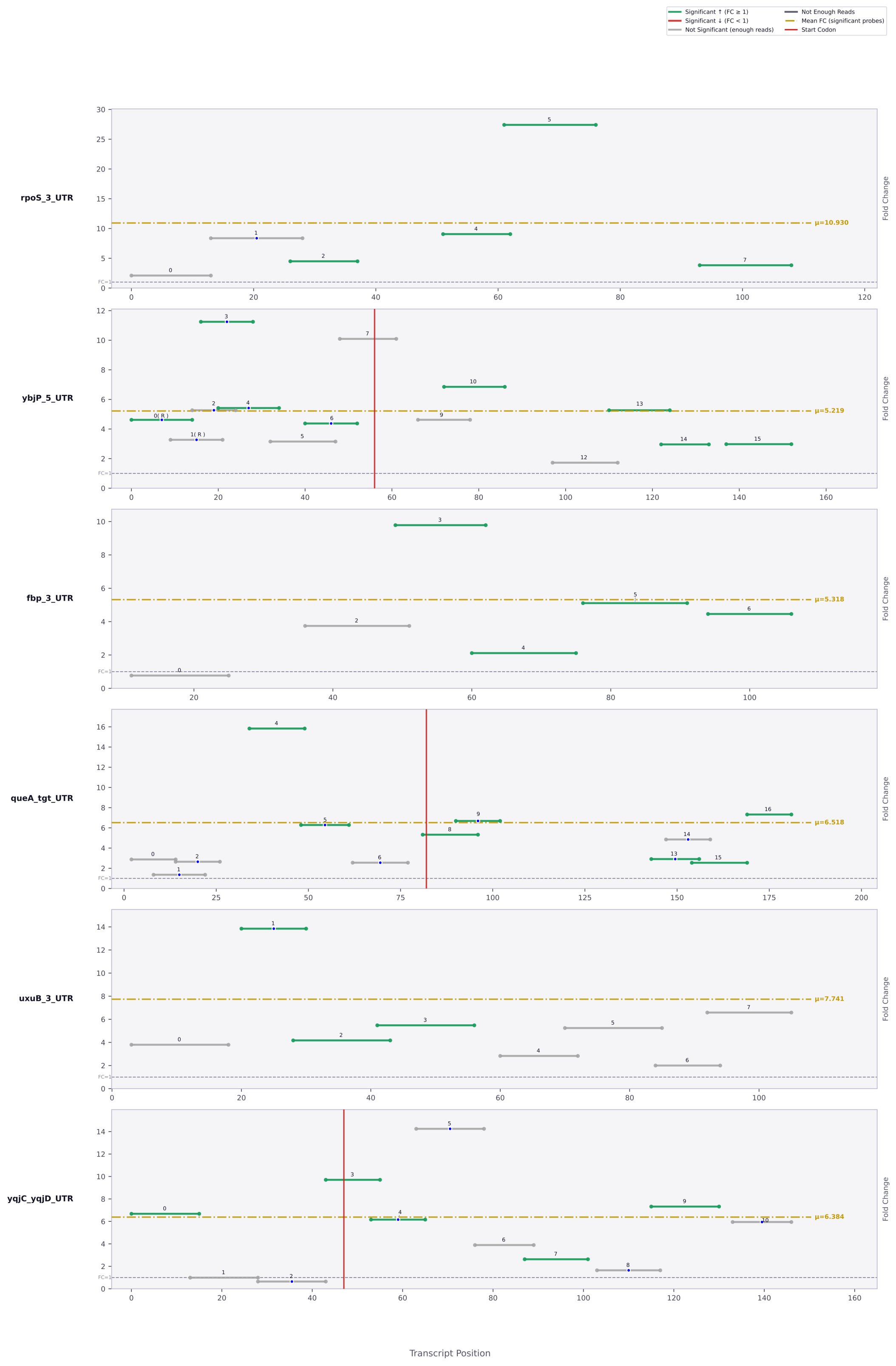

Significant ↑ (FC ≥ 1)
Significant ↓ (FC < 1)
Not Significant (enough reads)
Not Enough Reads
Mean FC (significant probes)
Start Codon
rpoS_3_UTR
μ=10.930
ybjP_5_UTR
μ=5.219
fbp_3_UTR
μ=5.318
queA_tgt_UTR
μ=6.518
uxuB_3_UTR
μ=7.741
yqjC_yqjD_UTR
μ=6.384
Fold Change
Transcript Position

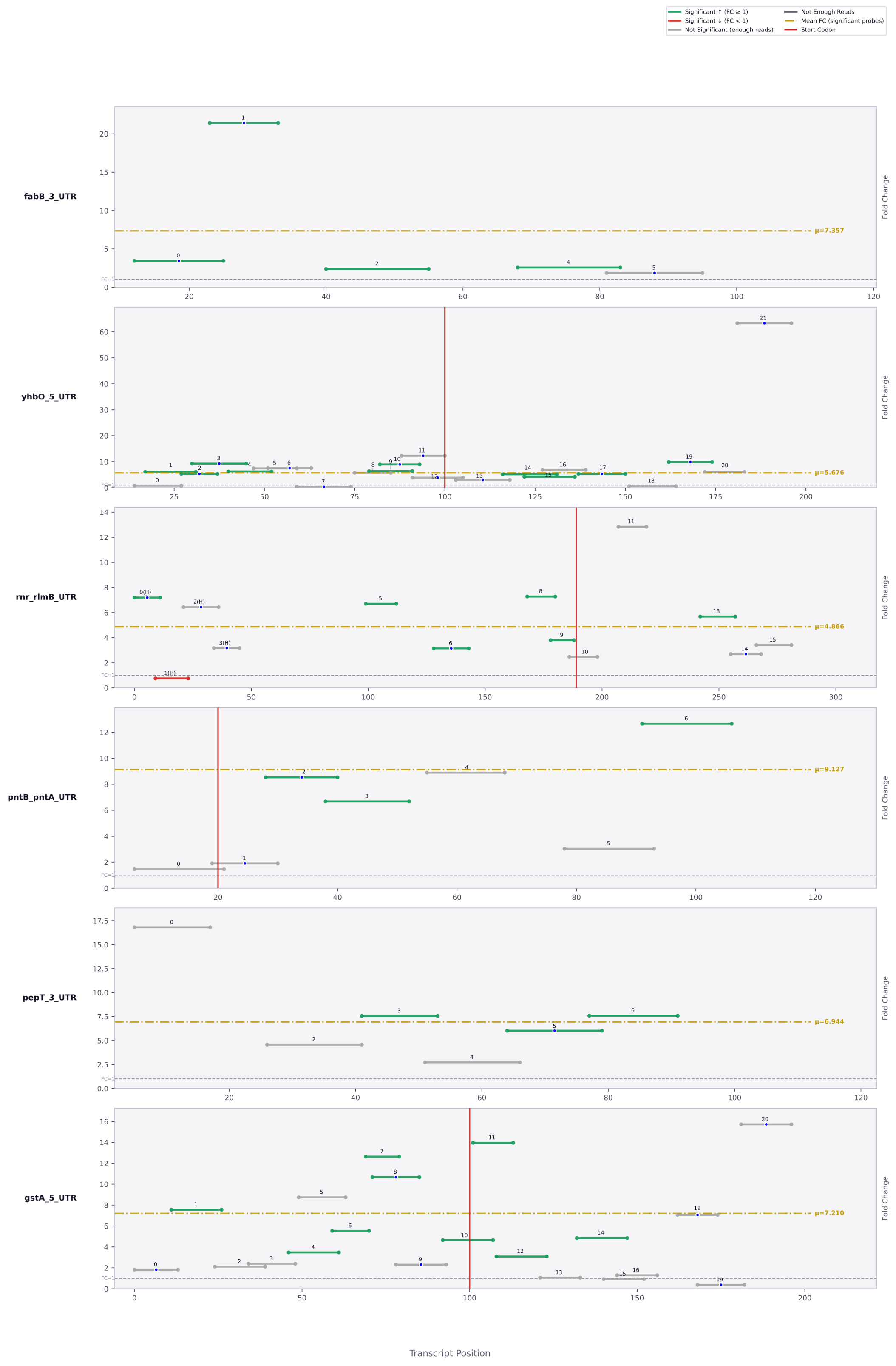

Significant ↑ (FC ≥ 1)
Significant ↓ (FC < 1)
Not Significant (enough reads)
Not Enough Reads
Mean FC (significant probes)
Start Codon
fabB_3_UTR
μ=7.357
yhbO_5_UTR
μ=5.676
rnr_rlmB_UTR
μ=4.866
pntB_pntA_UTR
μ=9.127
pepT_3_UTR
μ=6.944
gstA_5_UTR
μ=7.210
Fold Change
FC=1
Transcript Position

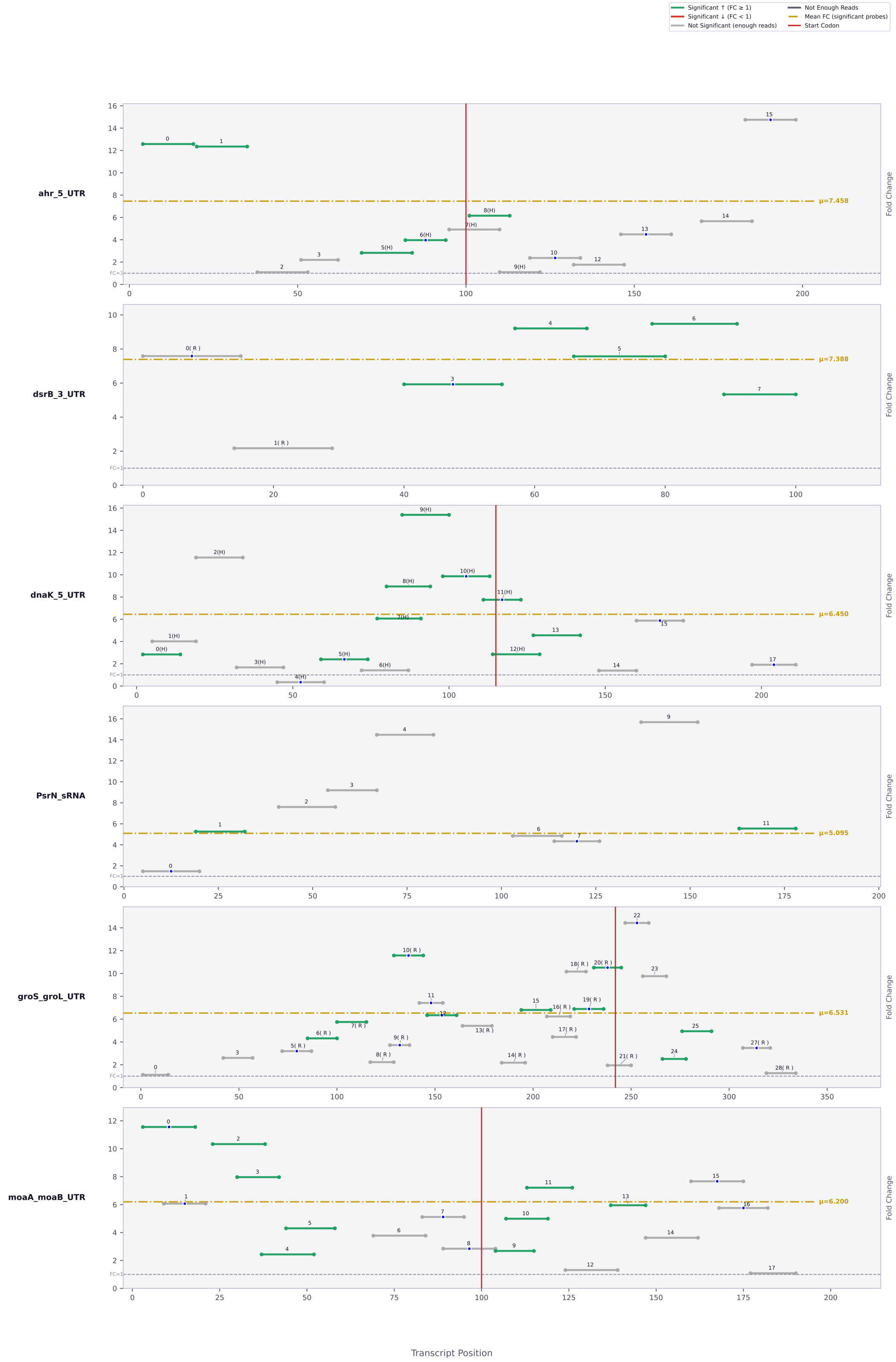

Significant ↑ (FC ≥ 1)
Significant ↓ (FC < 1)
Not Significant (enough reads)
Not Enough Reads
Mean FC (significant probes)
Start Codon
ahr_5_UTR
μ=7.458
dsrB_3_UTR
μ=7.388
dnaK_5_UTR
μ=6.450
PsrN_sRNA
μ=5.095
groS_groL_UTR
μ=6.531
moaA_moaB_UTR
μ=6.200
Fold Change
FC=1
Transcript Position

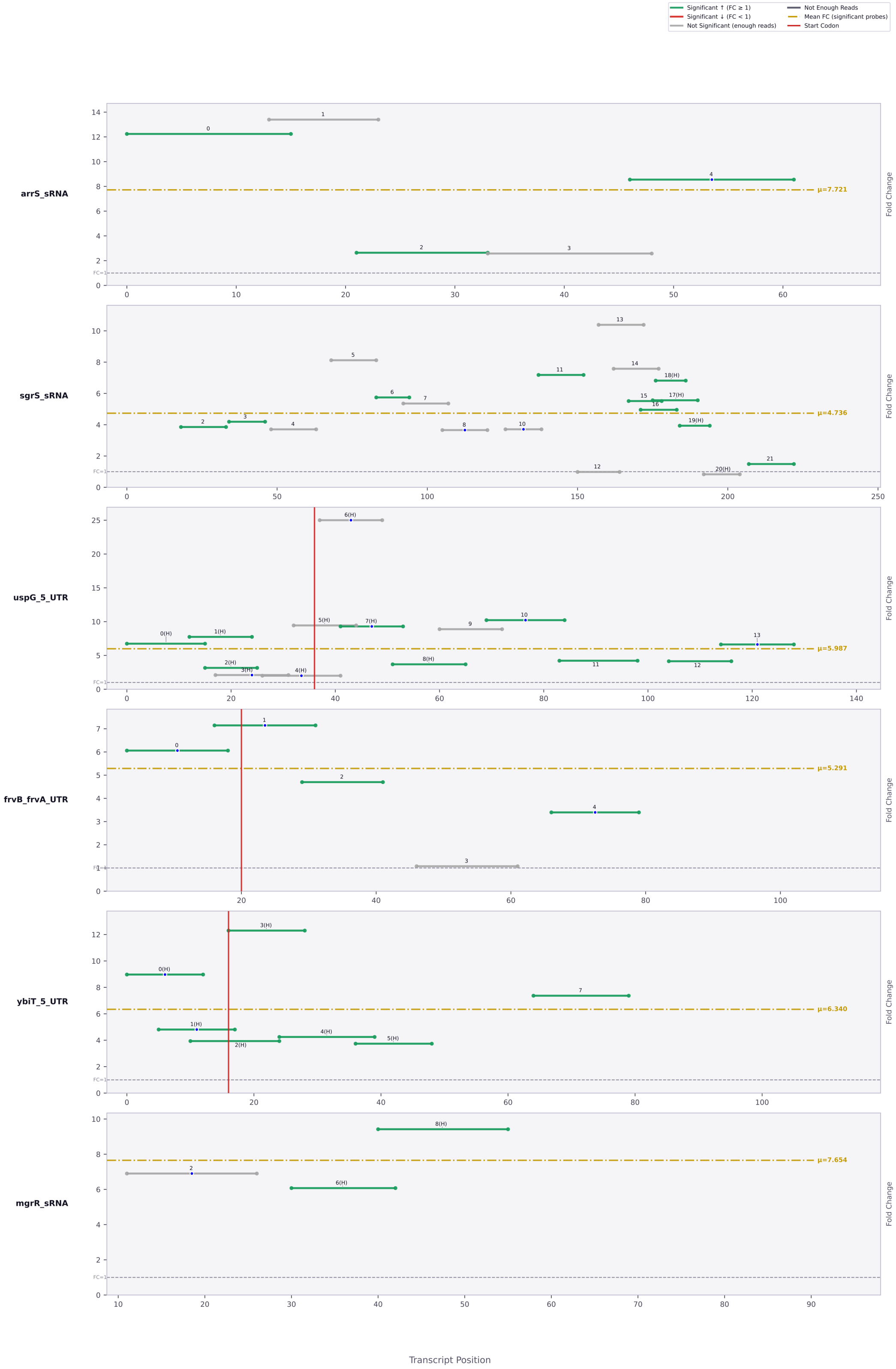

Significant ↑ (FC ≥ 1)
Significant ↓ (FC < 1)
Not Significant (enough reads)
Not Enough Reads
Mean FC (significant probes)
Start Codon
arrS_sRNA
μ=7.721
sgrS_sRNA
μ=4.736
uspG_5_UTR
μ=5.987
frvB_frvA_UTR
μ=5.291
ybiT_5_UTR
μ=6.340
mgrR_sRNA
μ=7.654
Fold Change
Transcript Position

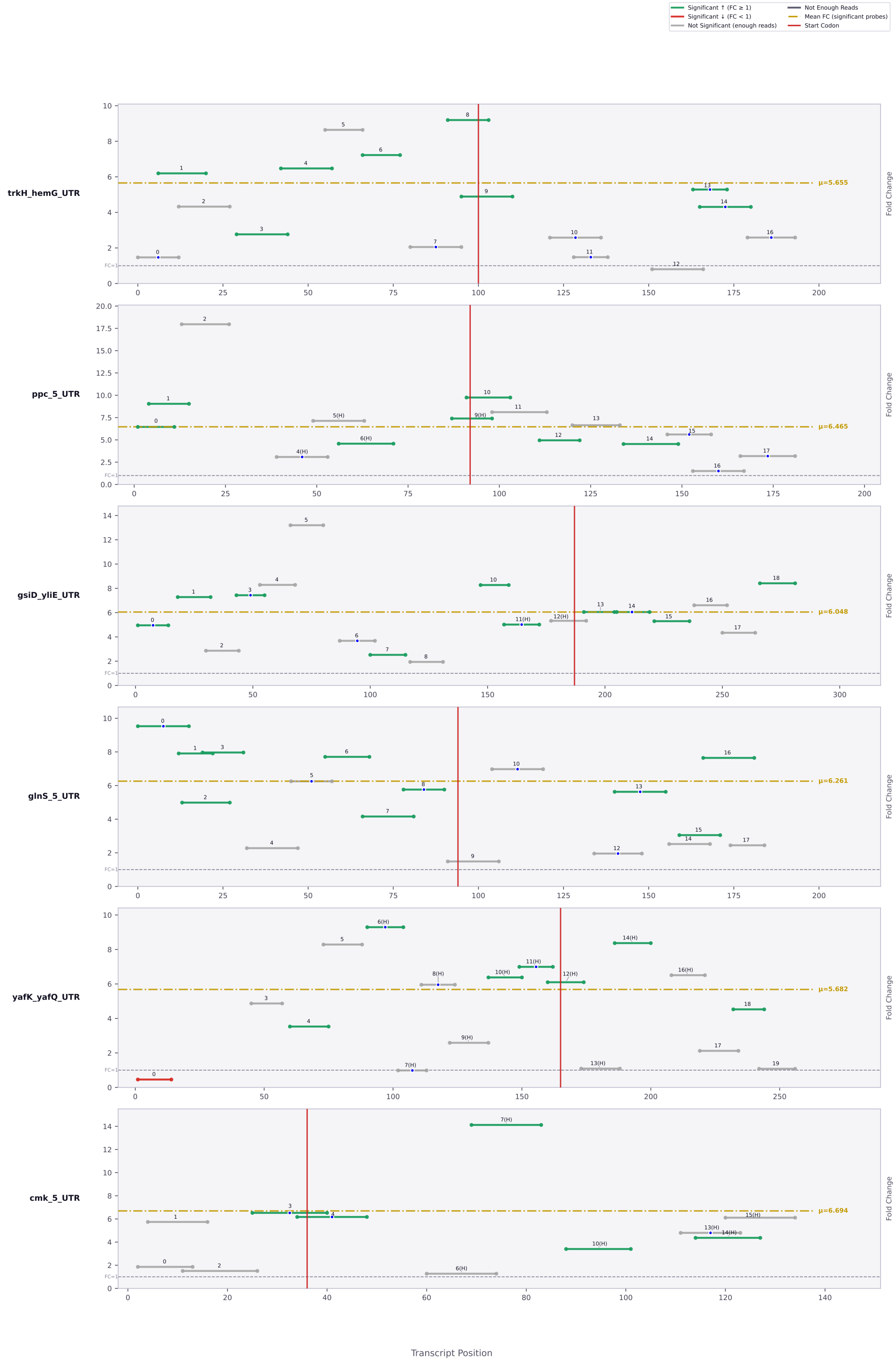

Significant ↑ (FC ≥ 1)
Significant ↓ (FC < 1)
Not Significant (enough reads)
Not Enough Reads
Mean FC (significant probes)
Start Codon
trkH_hemG_UTR
μ=5.655
ppc_5_UTR
μ=6.465
gsiD_yliE_UTR
μ=6.048
glnS_5_UTR
μ=6.261
yafK_yafQ_UTR
μ=5.682
cmk_5_UTR
μ=6.694
Fold Change
FC=1
Transcript Position

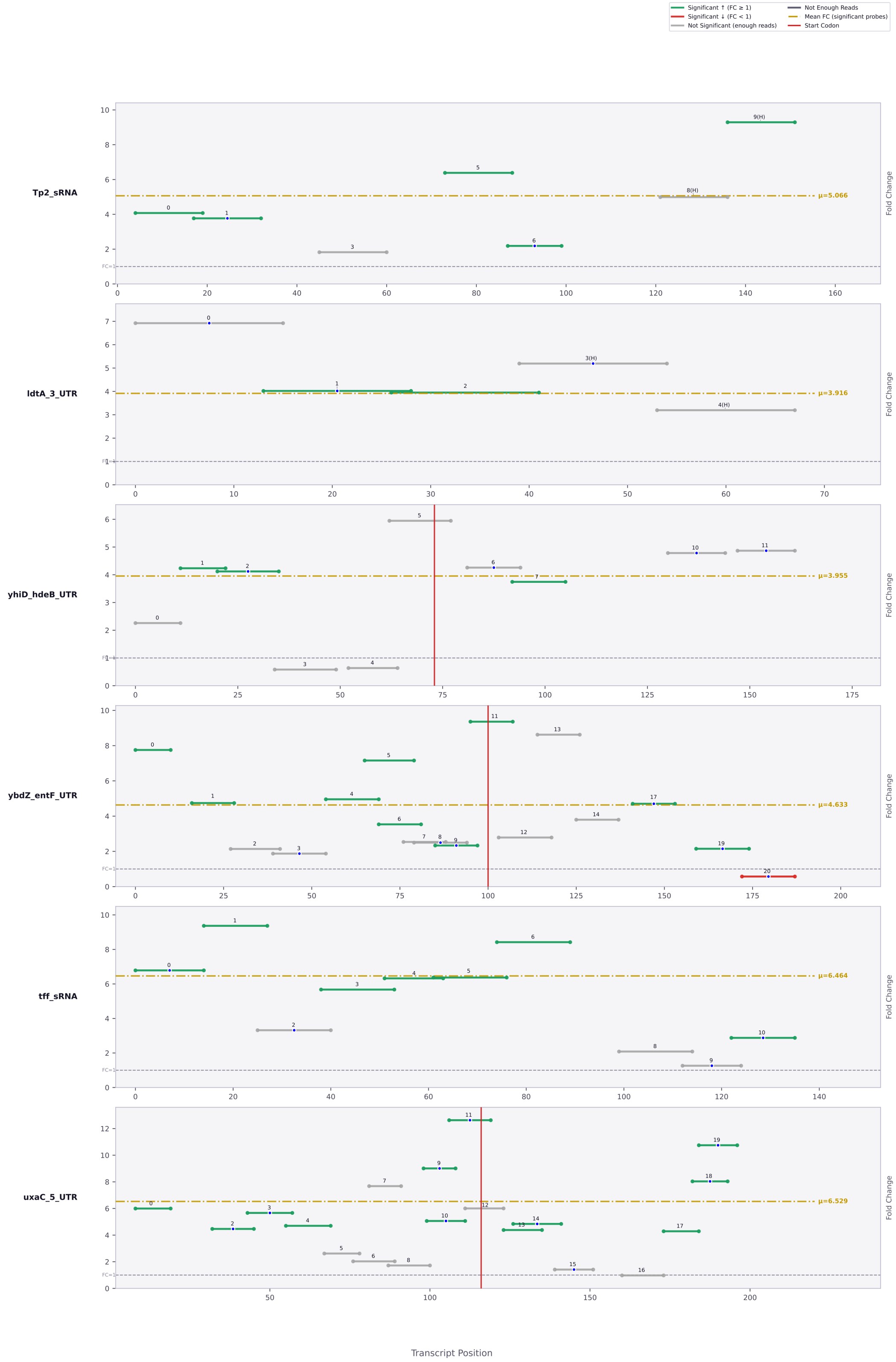

Significant ↑ (FC ≥ 1)
Significant ↓ (FC < 1)
Not Significant (enough reads)
Not Enough Reads
Mean FC (significant probes)
Start Codon
Tp2_sRNA
μ=5.066
ldtA_3_UTR
μ=3.916
yhiD_hdeB_UTR
μ=3.955
ybdZ_entF_UTR
μ=4.633
tff_sRNA
μ=6.464
uxaC_5_UTR
μ=6.529
Fold Change
FC=1
Transcript Position

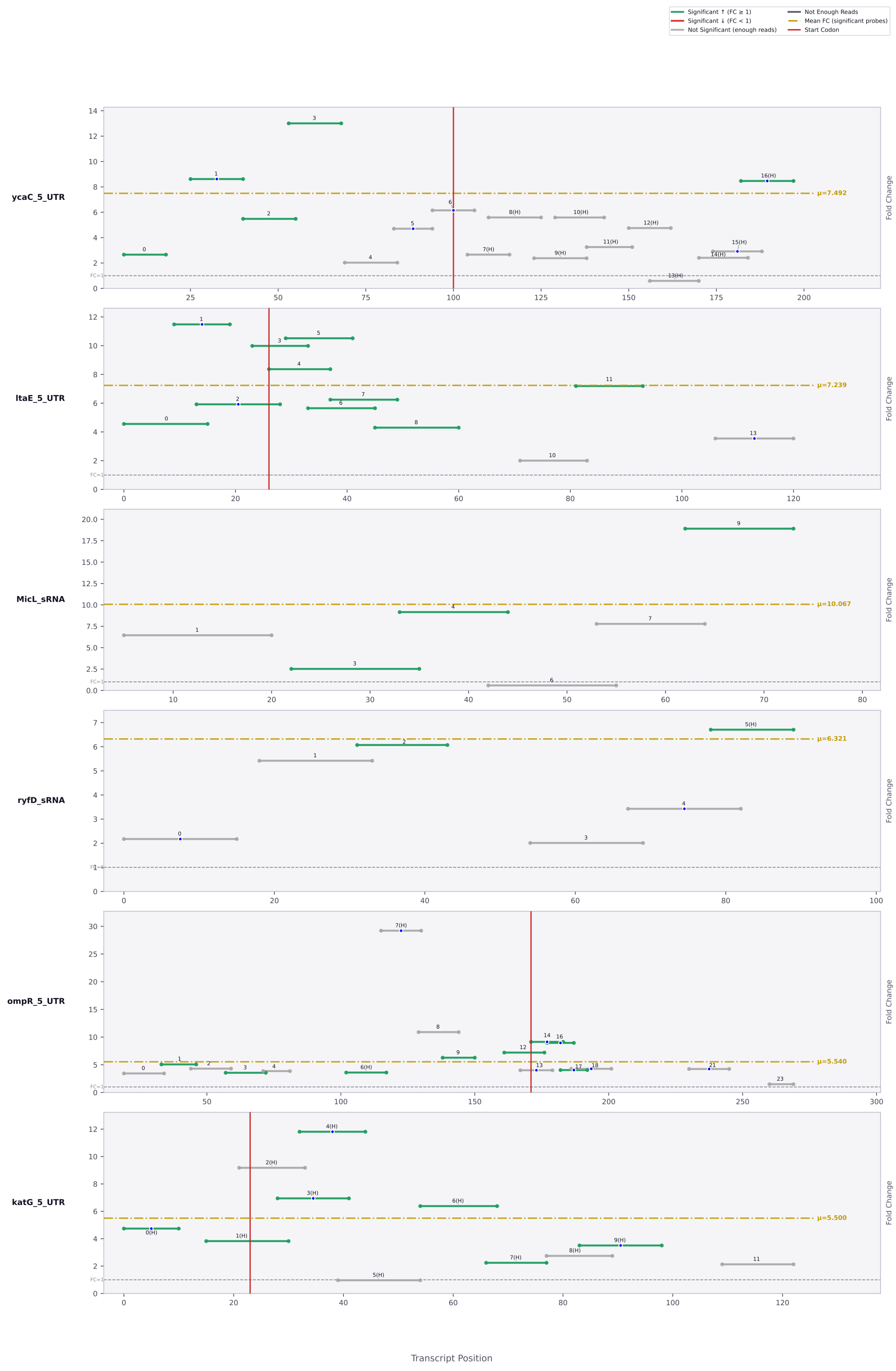

Significant ↑ (FC ≥ 1)
Significant ↓ (FC < 1)
Not Significant (enough reads)
Not Enough Reads
Mean FC (significant probes)
Start Codon
ycaC_5_UTR
μ=7.492
ltaE_5_UTR
μ=7.239
MicL_sRNA
μ=10.067
ryfD_sRNA
μ=6.321
ompR_5_UTR
μ=5.540
katG_5_UTR
μ=5.500
Fold Change
FC=1
Transcript Position

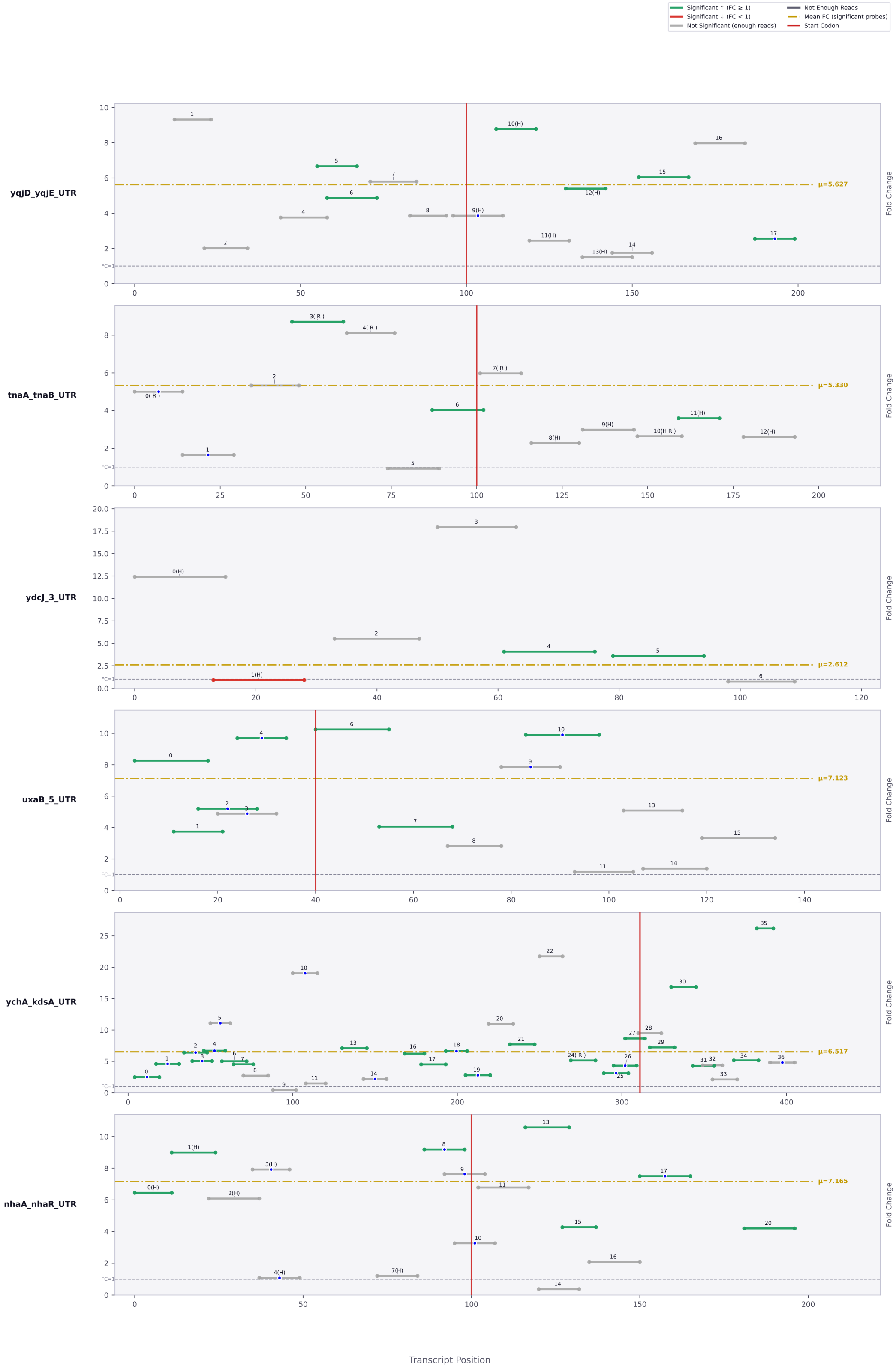

Significant ↑ (FC ≥ 1)
Significant ↓ (FC < 1)
Not Significant (enough reads)
Not Enough Reads
Mean FC (significant probes)
Start Codon
yqjD_yqjE_UTR
μ=5.627
tnaA_tnaB_UTR
μ=5.330
ydcJ_3_UTR
μ=2.612
uxaB_5_UTR
μ=7.123
ychA_kdsA_UTR
μ=6.517
nhaA_nhaR_UTR
μ=7.165
Fold Change
FC=1
Transcript Position

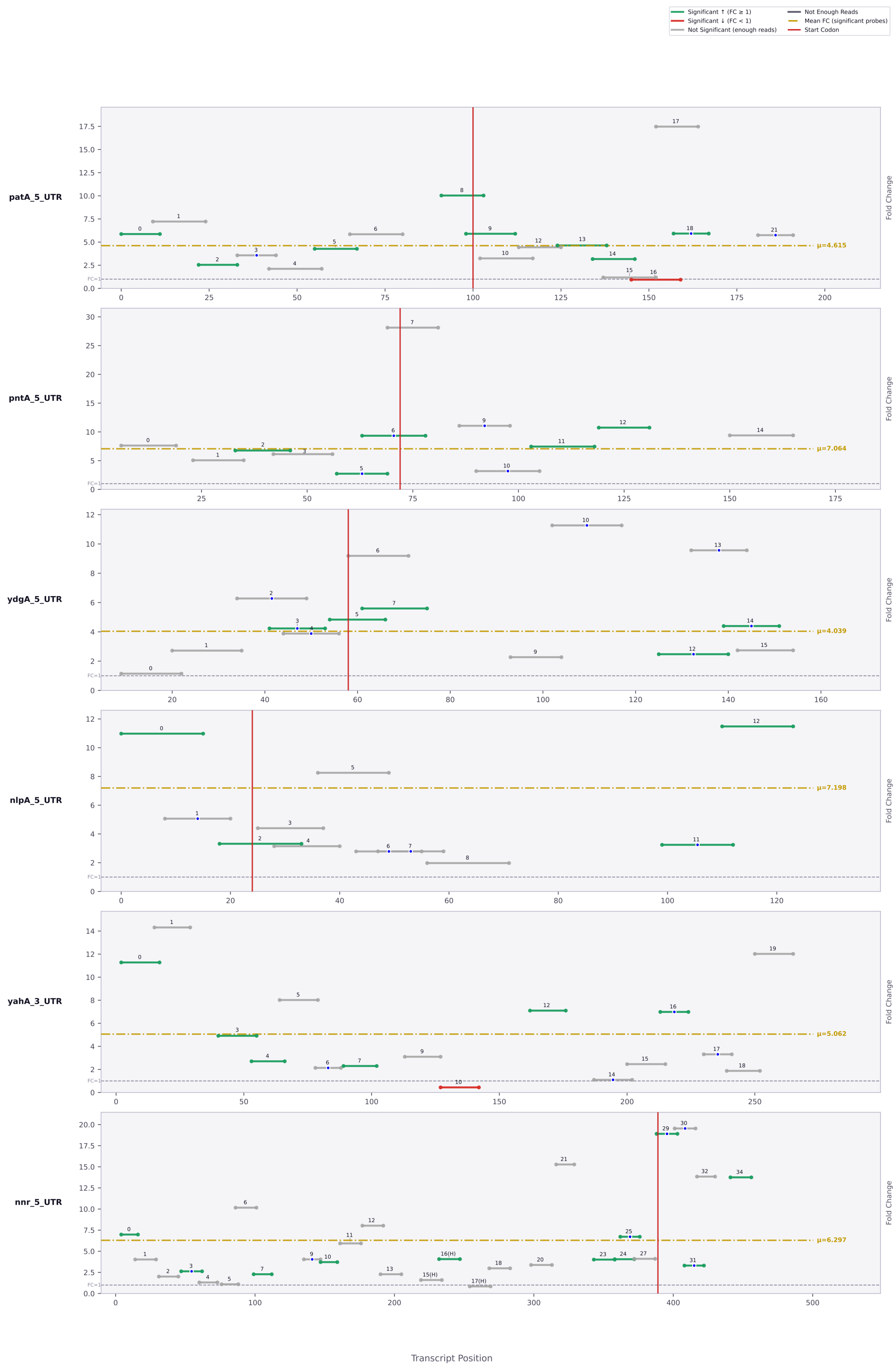

Significant ↑ (FC ≥ 1)
Significant ↓ (FC < 1)
Not Significant (enough reads)
Not Enough Reads
Mean FC (significant probes)
Start Codon
patA_5_UTR
μ=4.615
pntA_5_UTR
μ=7.064
ydgA_5_UTR
μ=4.039
nlpA_5_UTR
μ=7.198
yahA_3_UTR
μ=5.062
nnr_5_UTR
μ=6.297
Fold Change
Transcript Position

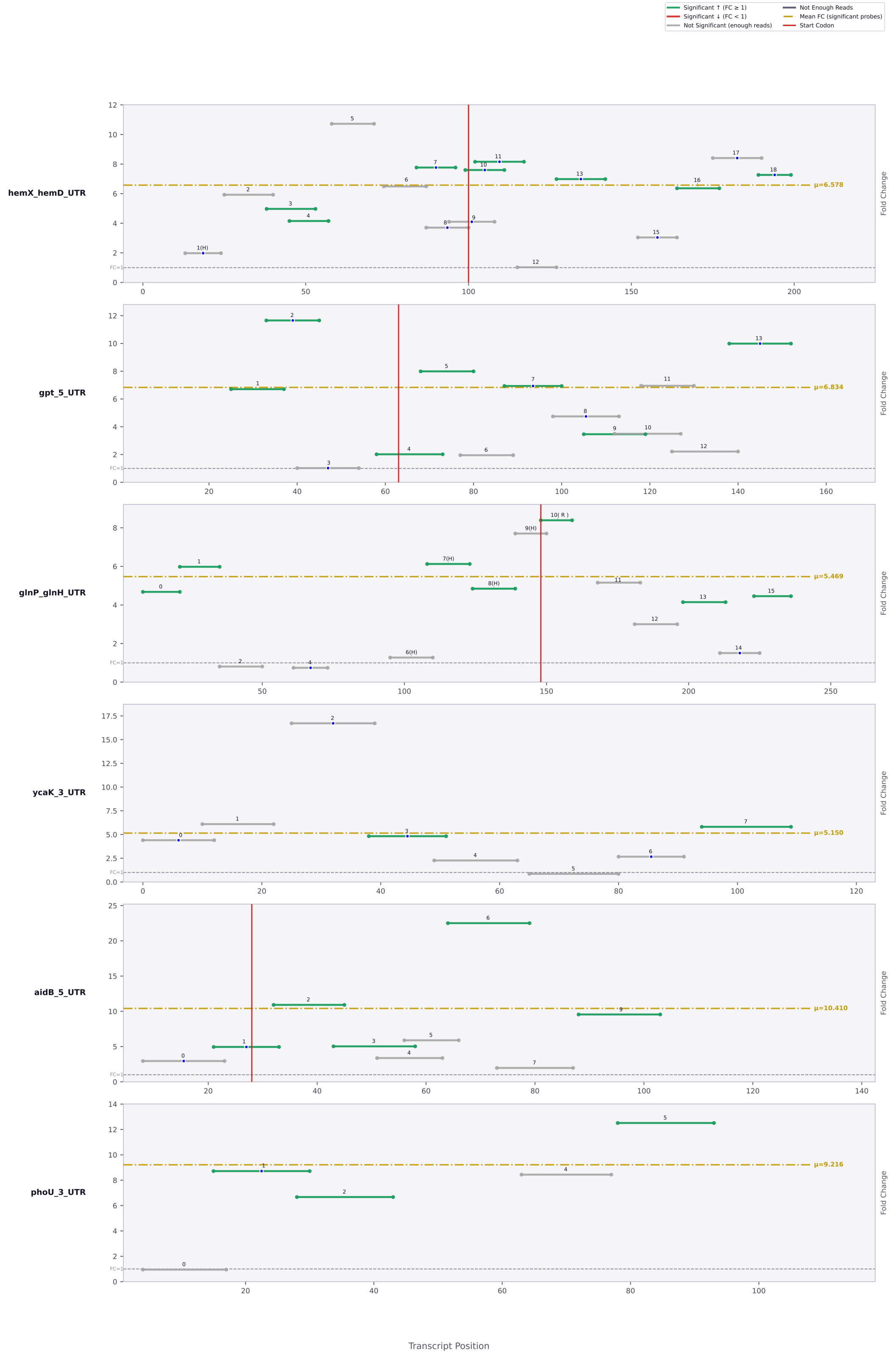

Significant ↑ (FC ≥ 1)
Significant ↓ (FC < 1)
Not Significant (enough reads)
Not Enough Reads
Mean FC (significant probes)
Start Codon
hemX_hemD_UTR
μ=6.578
gpt_5_UTR
μ=6.834
glnP_glnH_UTR
μ=5.469
ycaK_3_UTR
μ=5.150
aidB_5_UTR
μ=10.410
phoU_3_UTR
μ=9.216
FC=1
Fold Change
Transcript Position

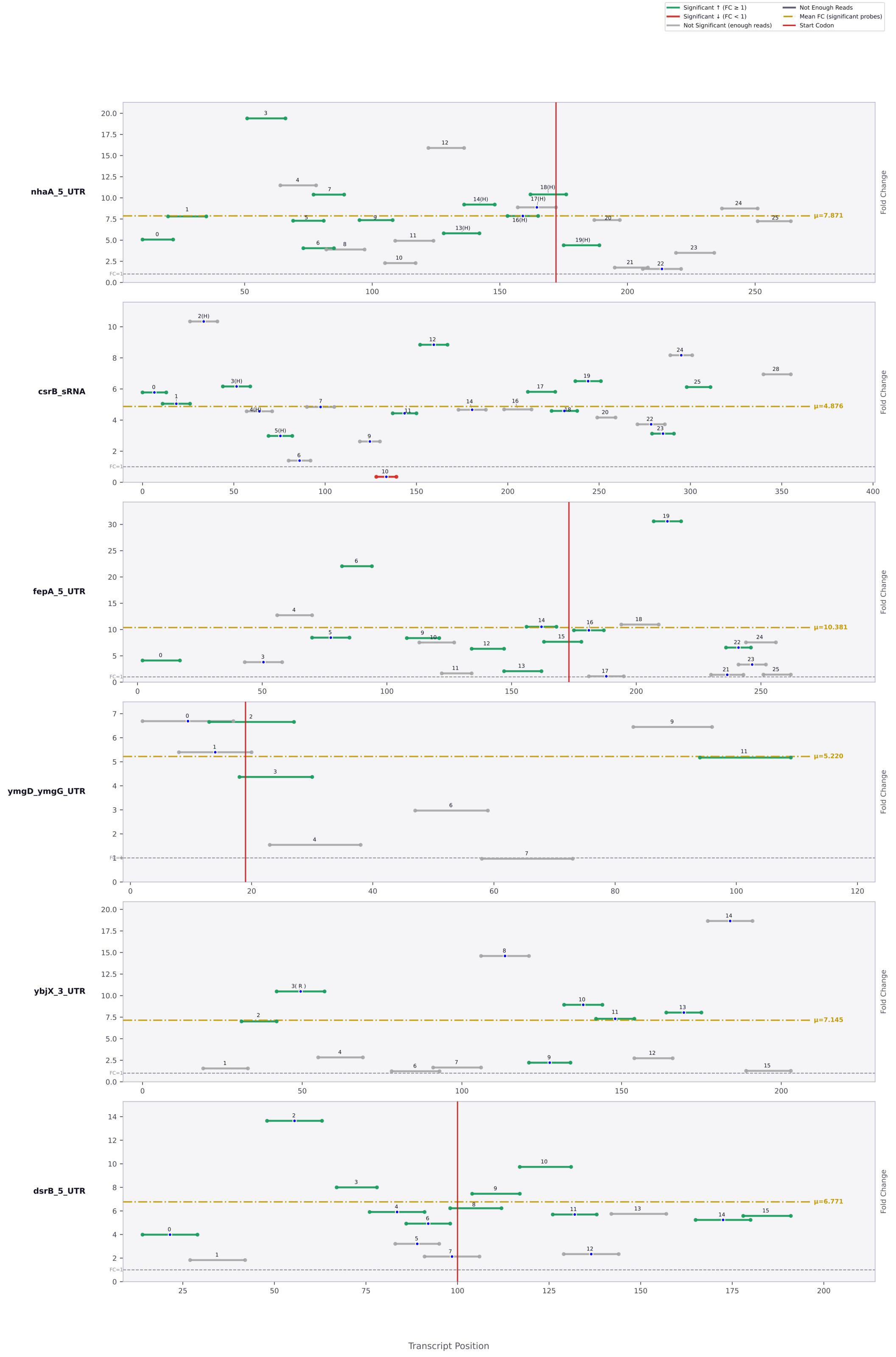

Significant ↑ (FC ≥ 1)
Significant ↓ (FC < 1)
Not Significant (enough reads)
Not Enough Reads
Mean FC (significant probes)
Start Codon
nhaA_5_UTR
μ=7.871
csrB_sRNA
μ=4.876
fepA_5_UTR
μ=10.381
ymgD_ymgG_UTR
μ=5.220
ybjX_3_UTR
μ=7.145
dsrB_5_UTR
μ=6.771
Fold Change
Transcript Position

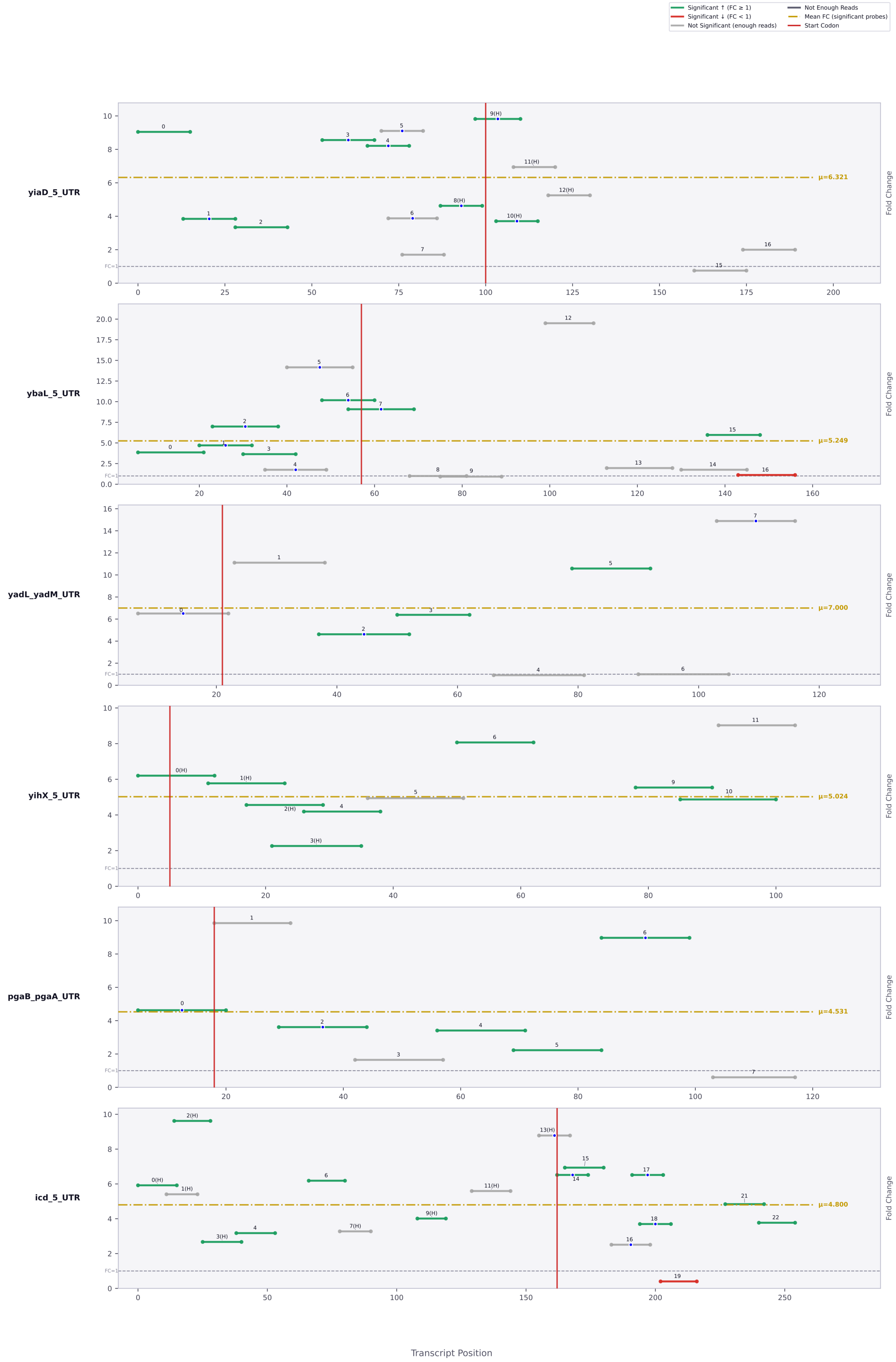

Significant ↑ (FC ≥ 1)
Significant ↓ (FC < 1)
Not Significant (enough reads)
Not Enough Reads
Mean FC (significant probes)
Start Codon
yiaD_5_UTR
μ=6.321
ybaL_5_UTR
μ=5.249
yadL_yadM_UTR
μ=7.000
yihX_5_UTR
μ=5.024
pgaB_pgaA_UTR
μ=4.531
icd_5_UTR
μ=4.800
Fold Change
FC=1
Transcript Position

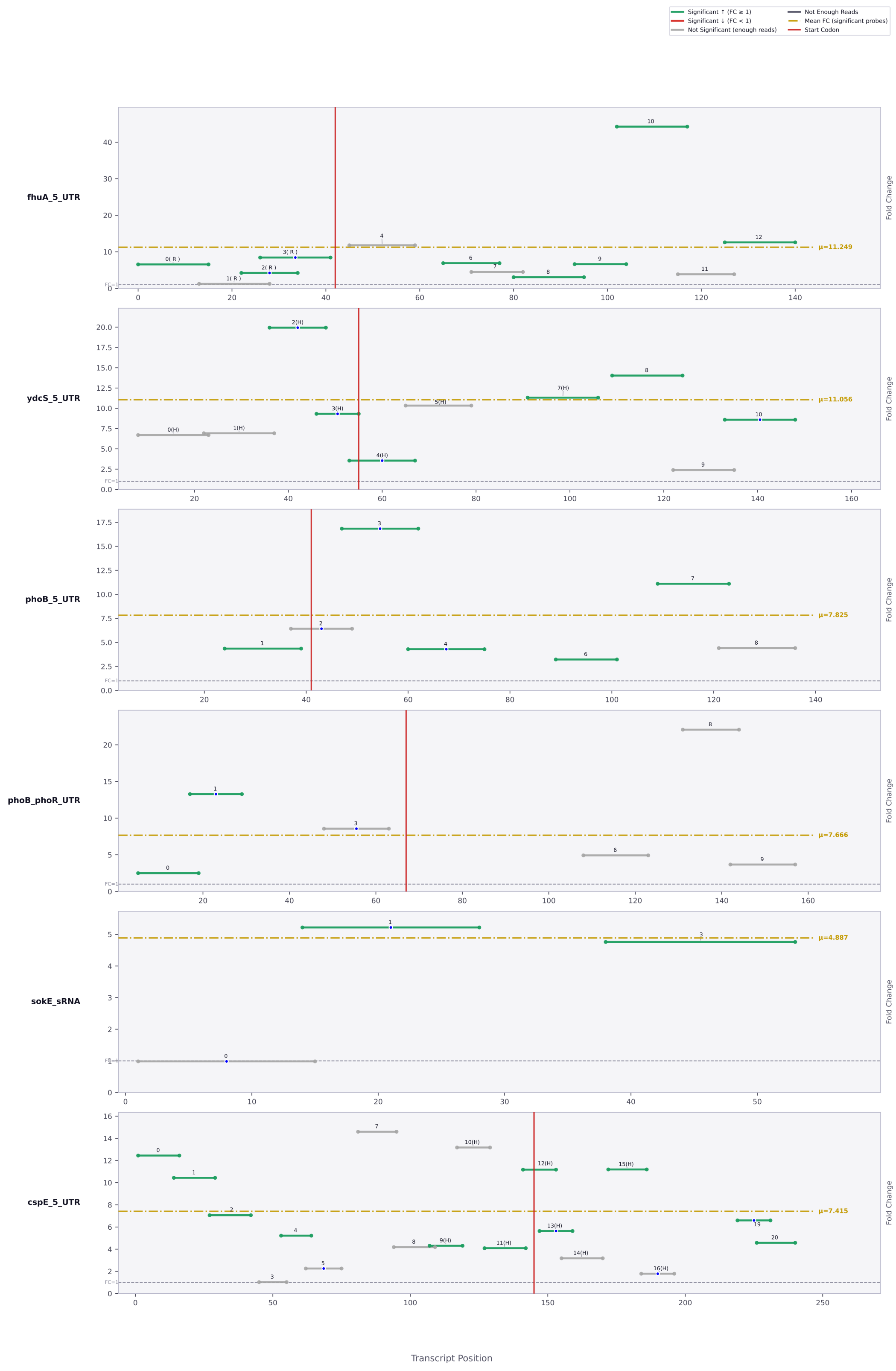

Significant ↑ (FC ≥ 1)
Significant ↓ (FC < 1)
Not Significant (enough reads)
Not Enough Reads
Mean FC (significant probes)
Start Codon
fhuA_5_UTR
μ=11.249
ydcS_5_UTR
μ=11.056
phoB_5_UTR
μ=7.825
phoB_phoR_UTR
μ=7.666
sokE_sRNA
μ=4.887
cspE_5_UTR
μ=7.415
Fold Change
Transcript Position

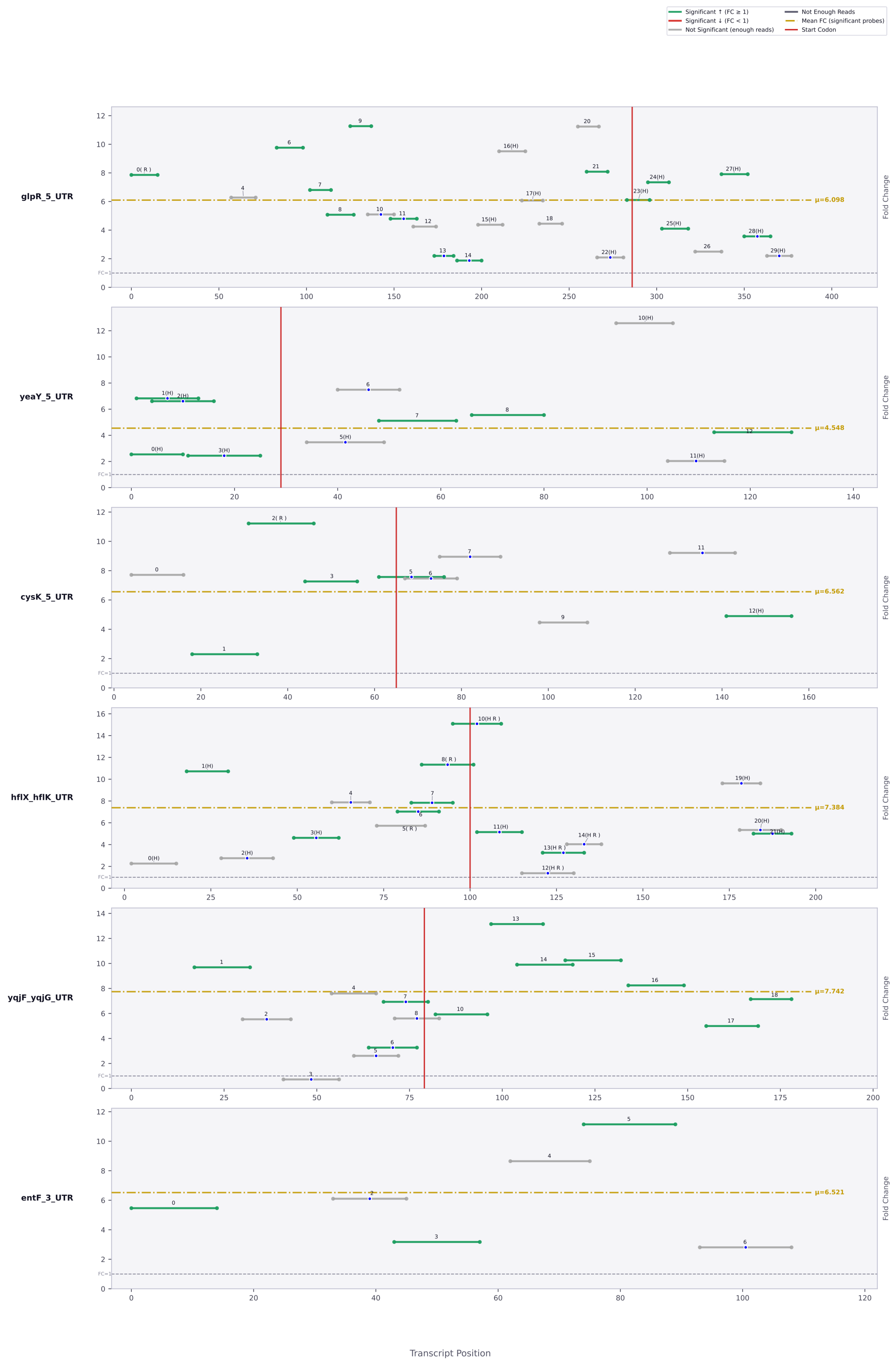

Significant ↑ (FC ≥ 1)
Significant ↓ (FC < 1)
Not Significant (enough reads)
Not Enough Reads
Mean FC (significant probes)
Start Codon
glpR_5_UTR
μ=6.098
yeaY_5_UTR
μ=4.548
cysK_5_UTR
μ=6.562
hflX_hflK_UTR
μ=7.384
yqjF_yqjG_UTR
μ=7.742
entF_3_UTR
μ=6.521
Fold Change
Transcript Position

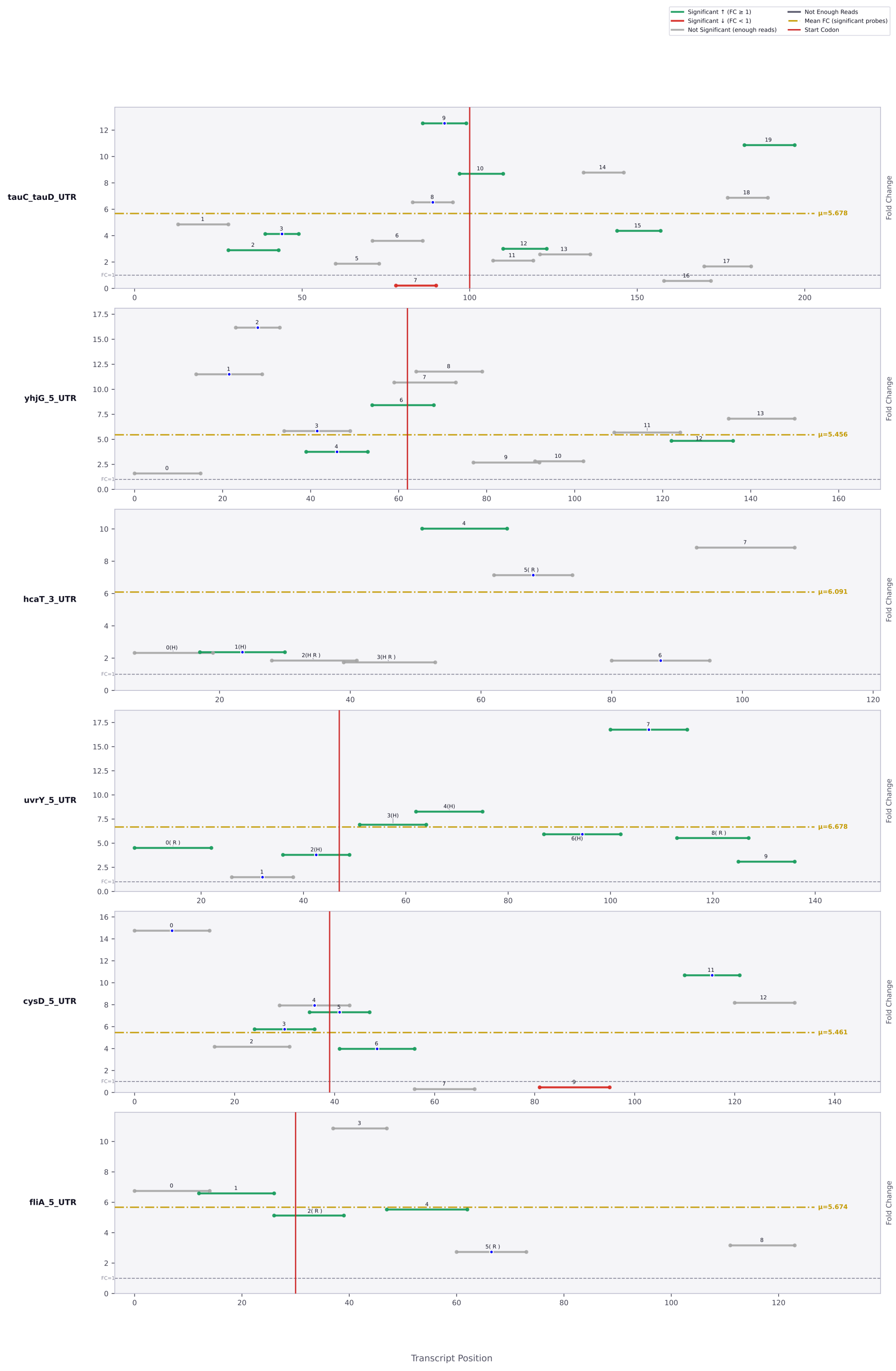

Significant ↑ (FC ≥ 1)
Significant ↓ (FC < 1)
Not Significant (enough reads)
Not Enough Reads
Mean FC (significant probes)
Start Codon
tauC_tauD_UTR
μ=5.678
yhjG_5_UTR
μ=5.456
hcaT_3_UTR
μ=6.091
uvrY_5_UTR
μ=6.678
cysD_5_UTR
μ=5.461
fliA_5_UTR
μ=5.674
FC=1
Fold Change
Transcript Position

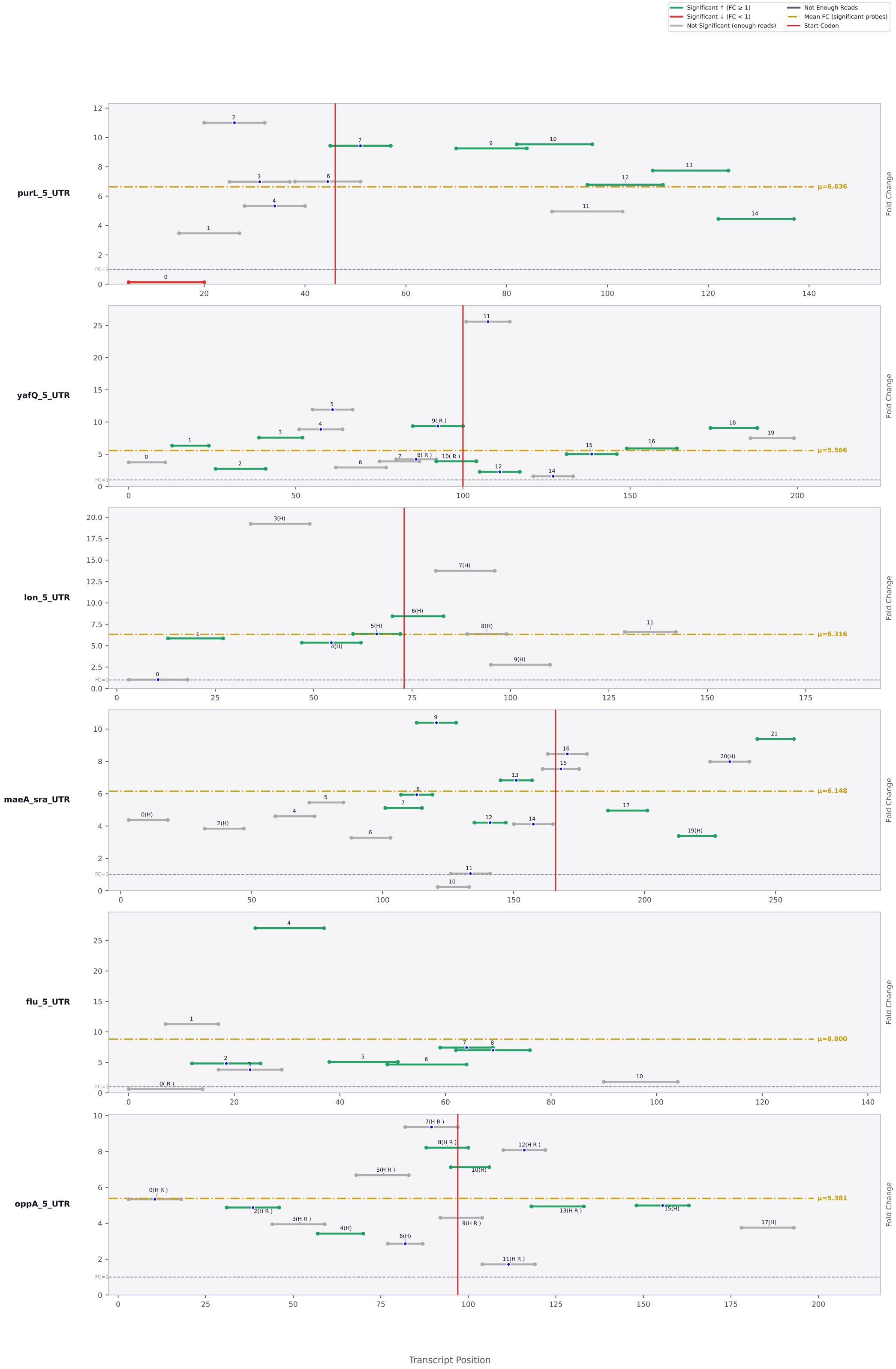

Significant ↑ (FC ≥ 1)
Significant ↓ (FC < 1)
Not Significant (enough reads)
Not Enough Reads
Mean FC (significant probes)
Start Codon
purL_5_UTR
μ=6.636
yafQ_5_UTR
μ=5.566
lon_5_UTR
μ=6.316
maeA_sra_UTR
μ=6.148
flu_5_UTR
μ=8.800
oppA_5_UTR
μ=5.381
Fold Change
FC=1
Transcript Position

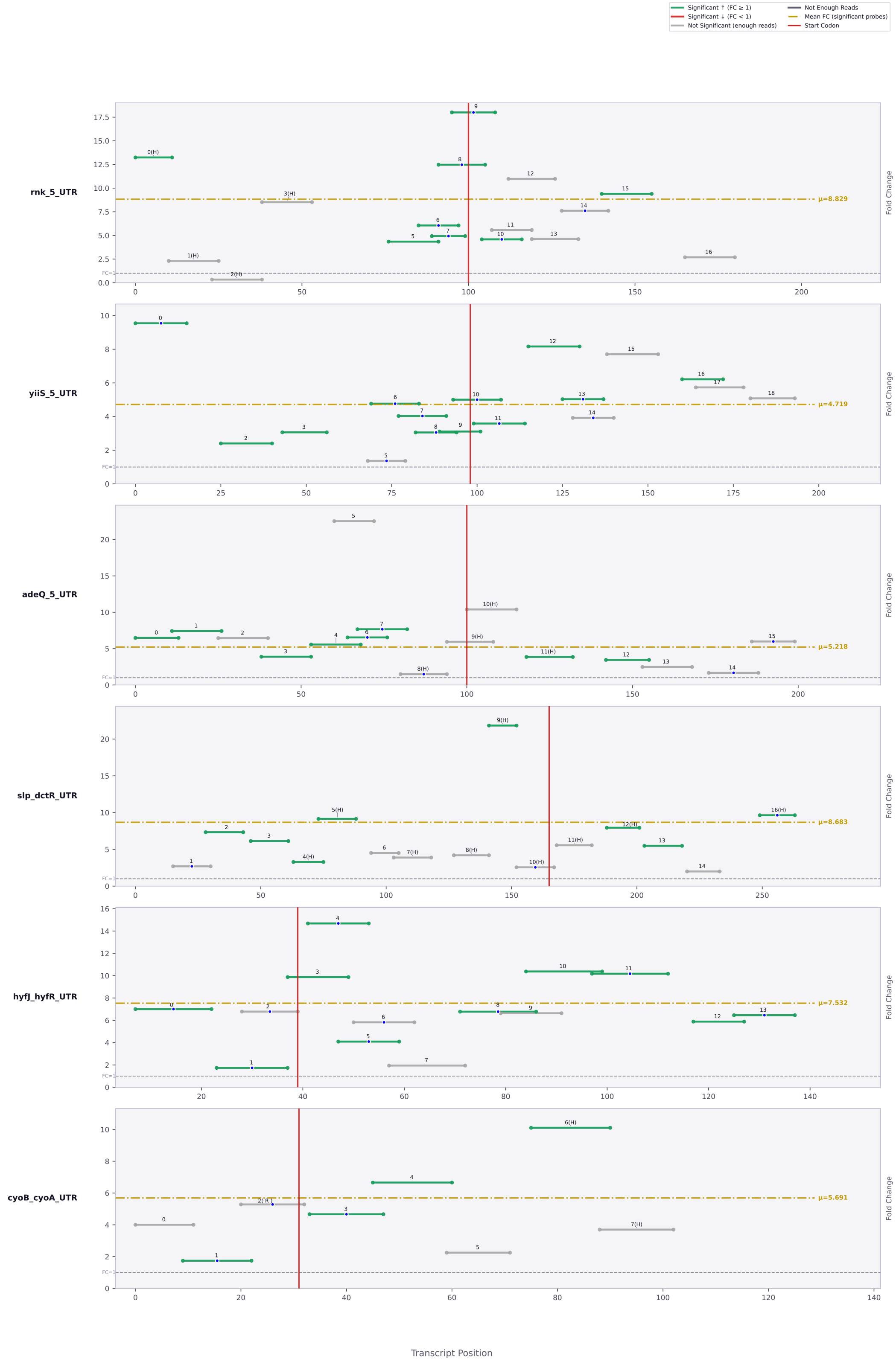

Significant ↑ (FC ≥ 1)
Significant ↓ (FC < 1)
Not Significant (enough reads)
Not Enough Reads
Mean FC (significant probes)
Start Codon
rnk_5_UTR
μ=8.829
yiiS_5_UTR
μ=4.719
adeQ_5_UTR
μ=5.218
slp_dctR_UTR
μ=8.683
hyfJ_hyfR_UTR
μ=7.532
cyoB_cyoA_UTR
μ=5.691
Fold Change
Transcript Position

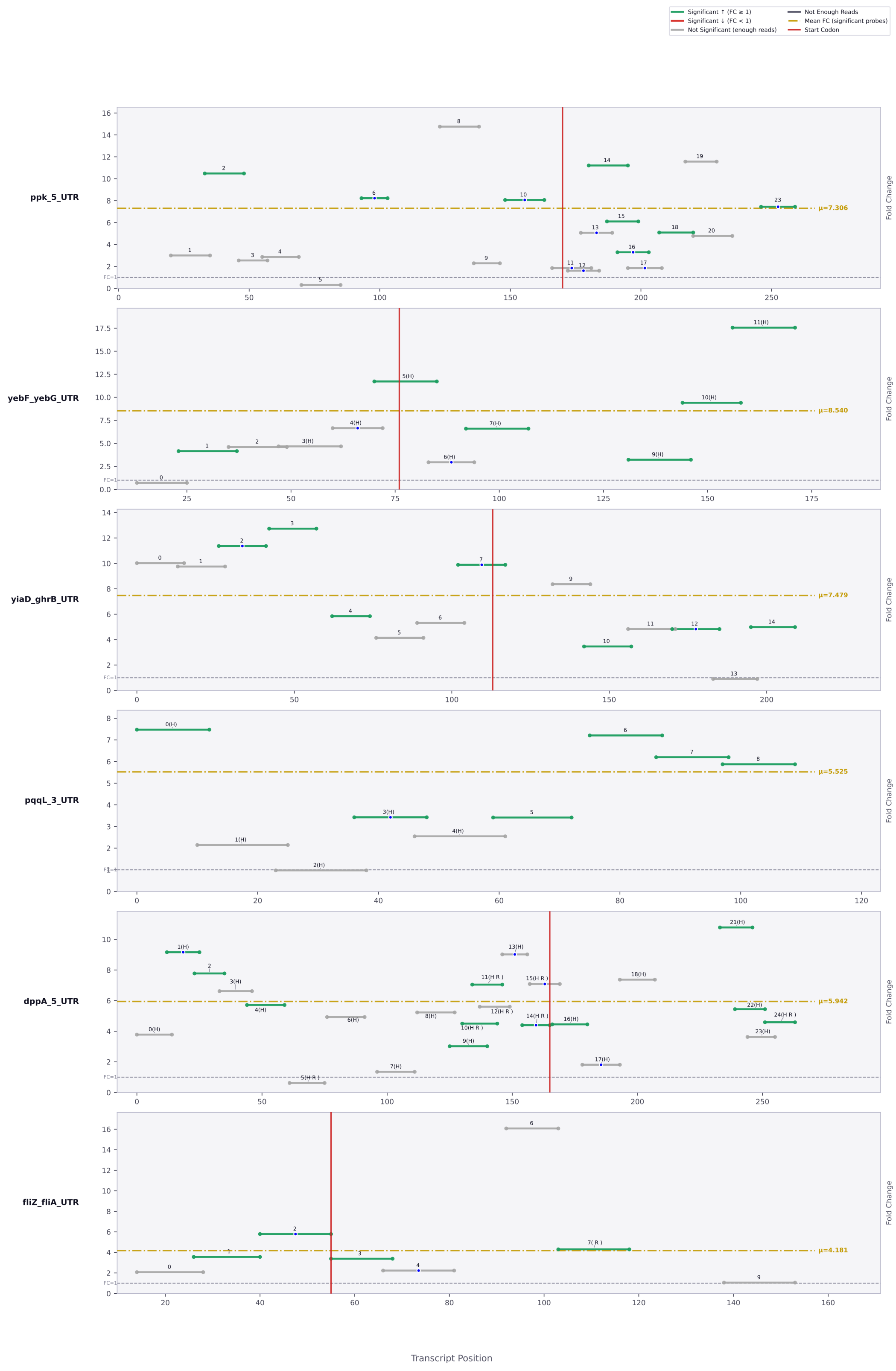

Significant ↑ (FC ≥ 1)
Significant ↓ (FC < 1)
Not Significant (enough reads)
Not Enough Reads
Mean FC (significant probes)
Start Codon
ppk_5_UTR
μ=7.306
yebF_yebG_UTR
μ=8.540
yiaD_ghrB_UTR
μ=7.479
pqqL_3_UTR
μ=5.525
dppA_5_UTR
μ=5.942
fliZ_fliA_UTR
μ=4.181
Fold Change
Transcript Position

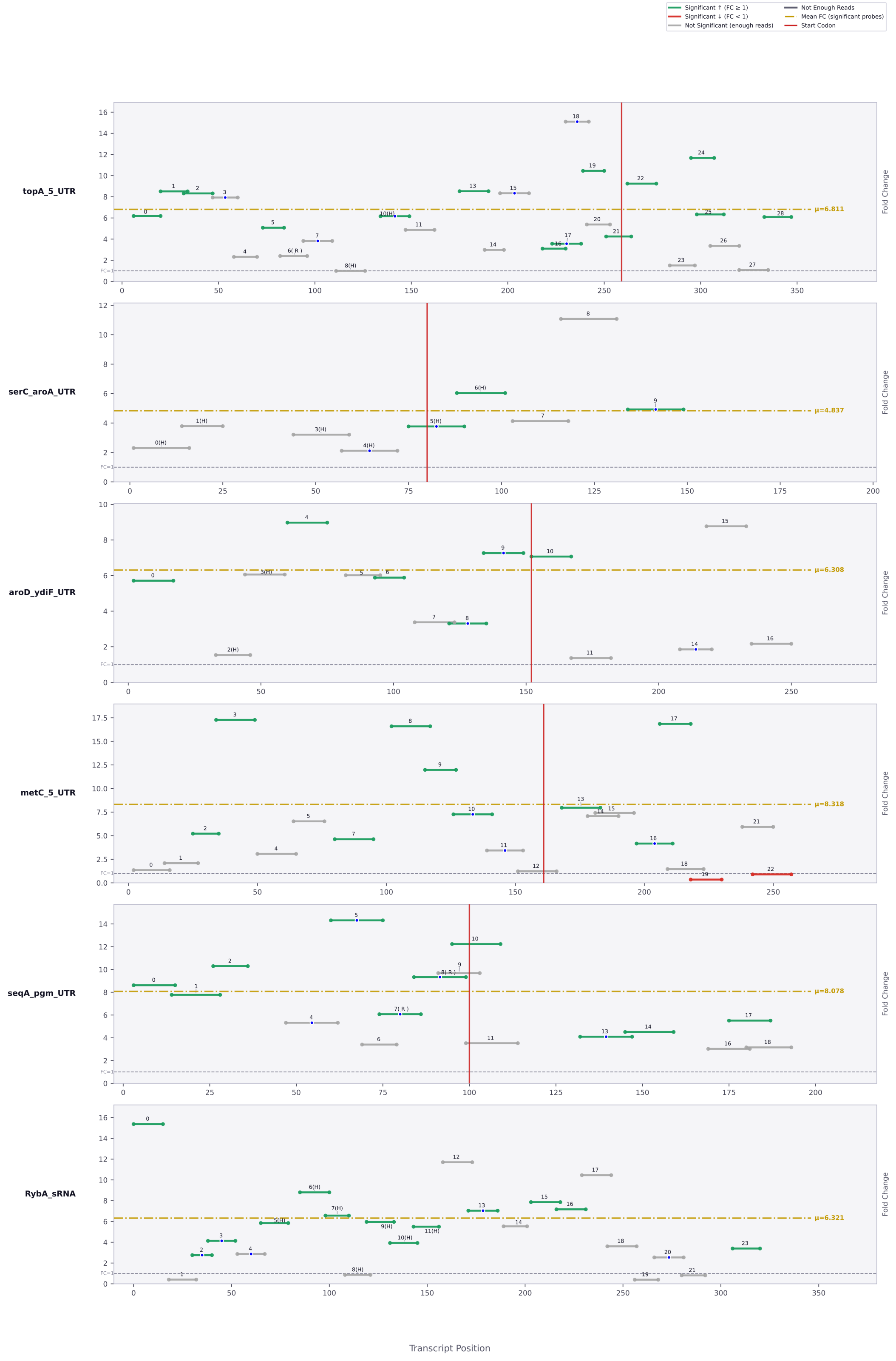

Significant ↑ (FC ≥ 1)
Significant ↓ (FC < 1)
Not Significant (enough reads)
Not Enough Reads
Mean FC (significant probes)
Start Codon
topA_5_UTR
μ=6.811
serC_aroA_UTR
μ=4.837
aroD_ydiF_UTR
μ=6.308
metC_5_UTR
μ=8.318
seqA_pgm_UTR
μ=8.078
RybA_sRNA
μ=6.321
Fold Change
FC=1
Transcript Position

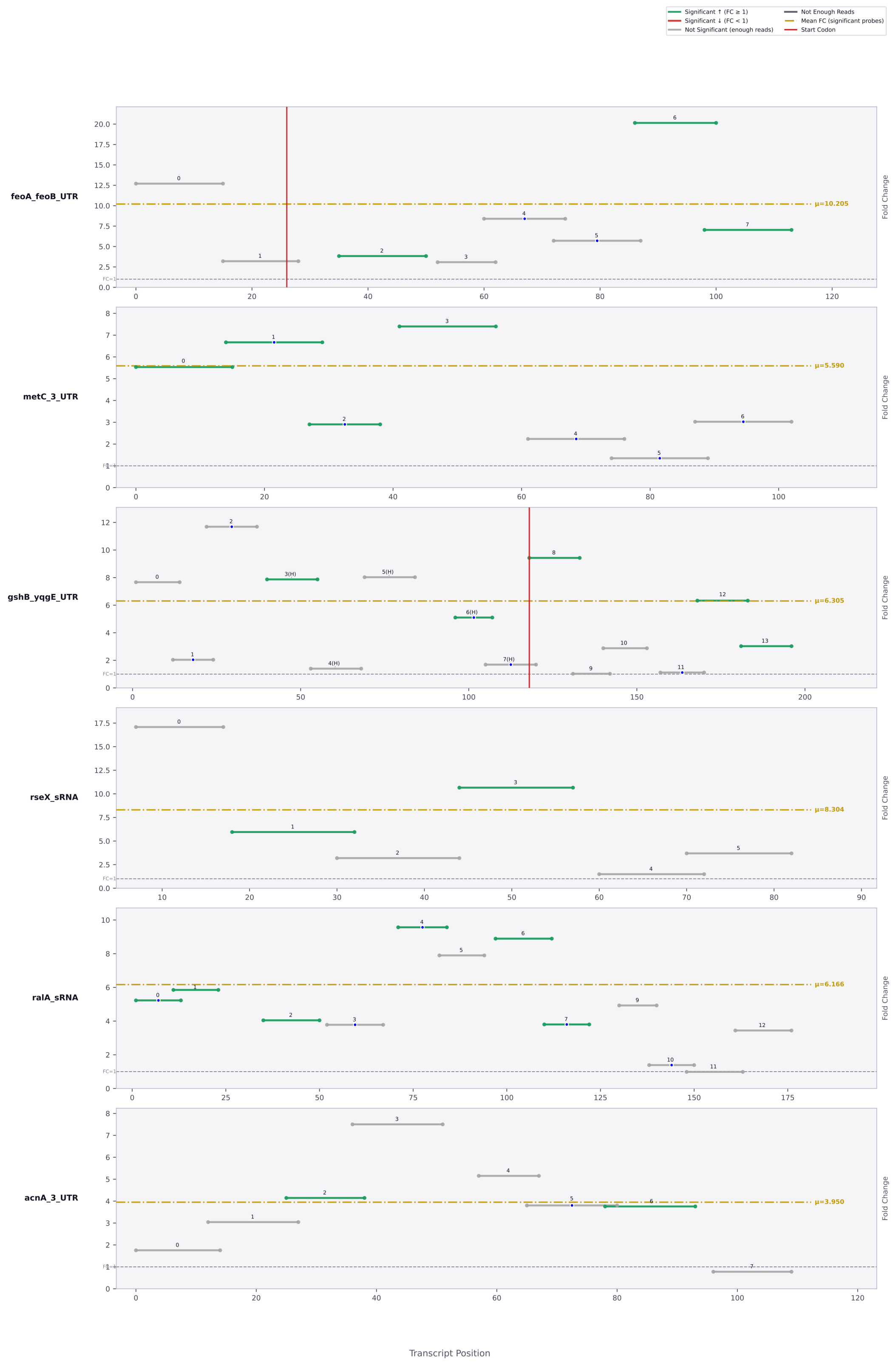

Significant ↑ (FC ≥ 1)
Significant ↓ (FC < 1)
Not Significant (enough reads)
Not Enough Reads
Mean FC (significant probes)
Start Codon
feoA_feoB_UTR
μ=10.205
metC_3_UTR
μ=5.590
gshB_yqgE_UTR
μ=6.305
rseX_sRNA
μ=8.304
ralA_sRNA
μ=6.166
acnA_3_UTR
μ=3.950
Fold Change
FC=1
Transcript Position

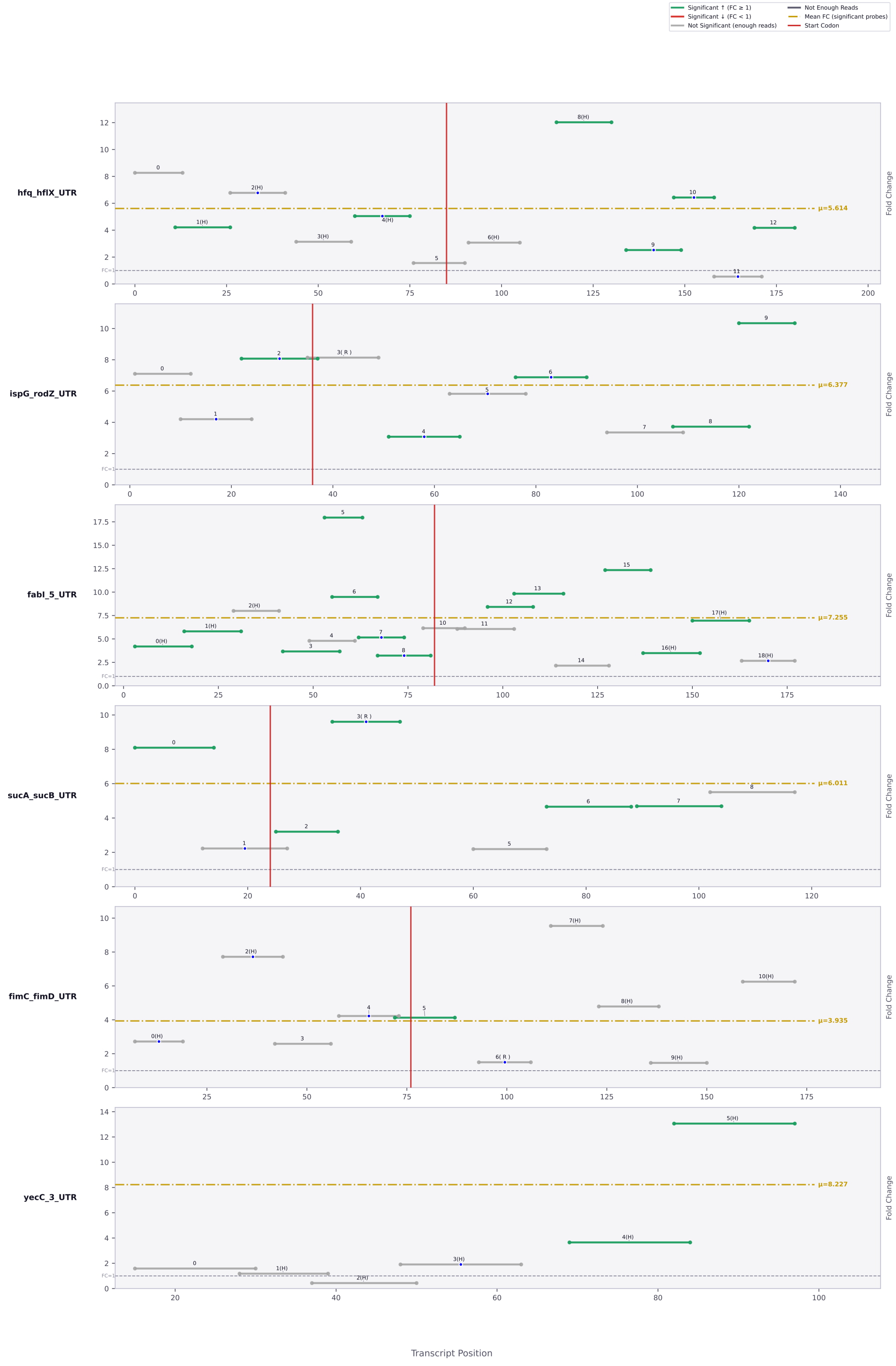

Significant ↑ (FC ≥ 1)
Significant ↓ (FC < 1)
Not Significant (enough reads)
Not Enough Reads
Mean FC (significant probes)
Start Codon
hfq_hflX_UTR
μ=5.614
ispG_rodZ_UTR
μ=6.377
fabI_5_UTR
μ=7.255
sucA_sucB_UTR
μ=6.011
fimC_fimD_UTR
μ=3.935
yecC_3_UTR
μ=8.227
Fold Change
Transcript Position

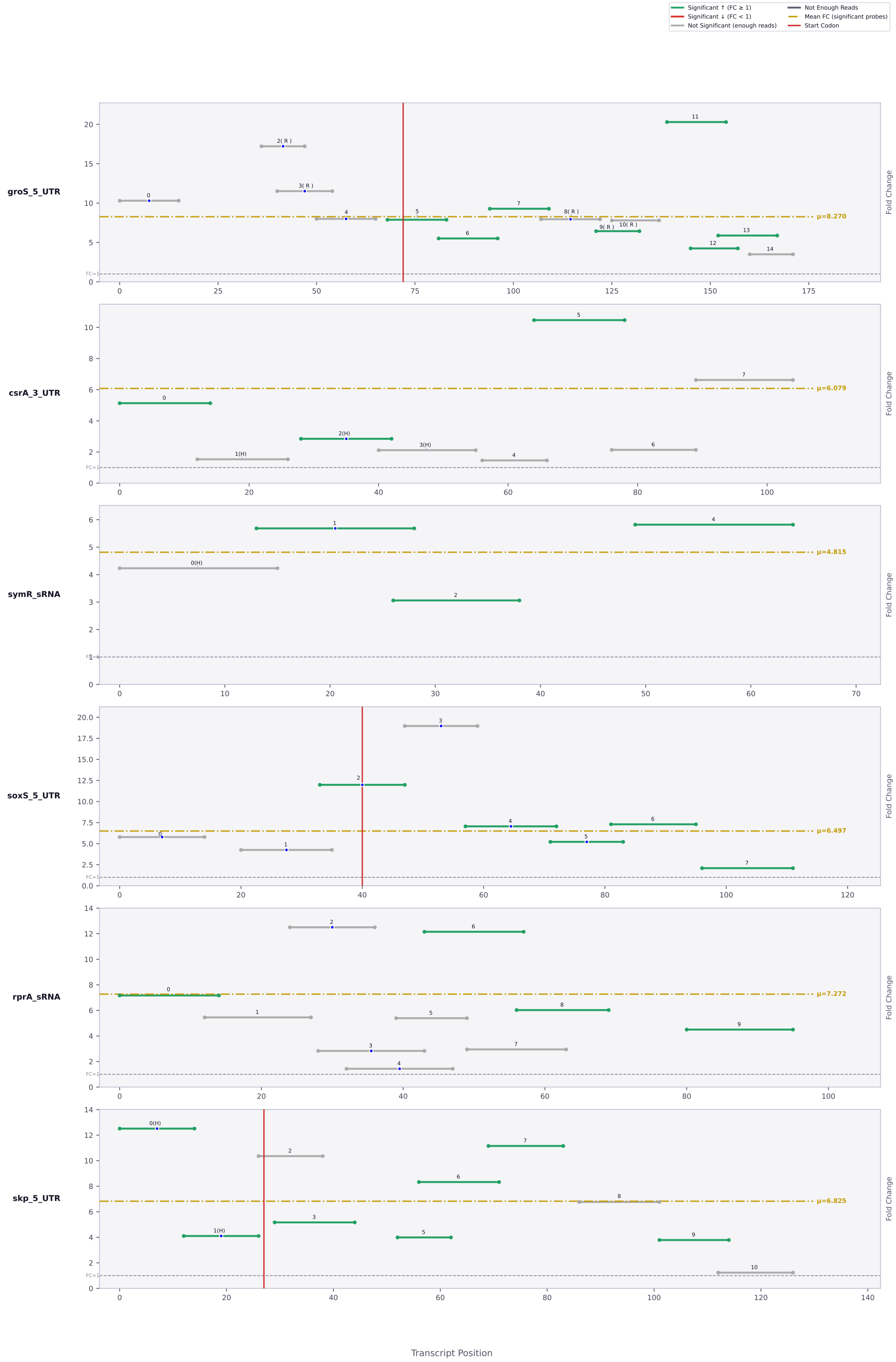

Significant ↑ (FC ≥ 1)
Significant ↓ (FC < 1)
Not Significant (enough reads)
Not Enough Reads
Mean FC (significant probes)
Start Codon
groS_5_UTR
μ=8.270
csrA_3_UTR
μ=6.079
symR_sRNA
μ=4.815
soxS_5_UTR
μ=6.497
rprA_sRNA
μ=7.272
skp_5_UTR
μ=6.825
Fold Change
Transcript Position

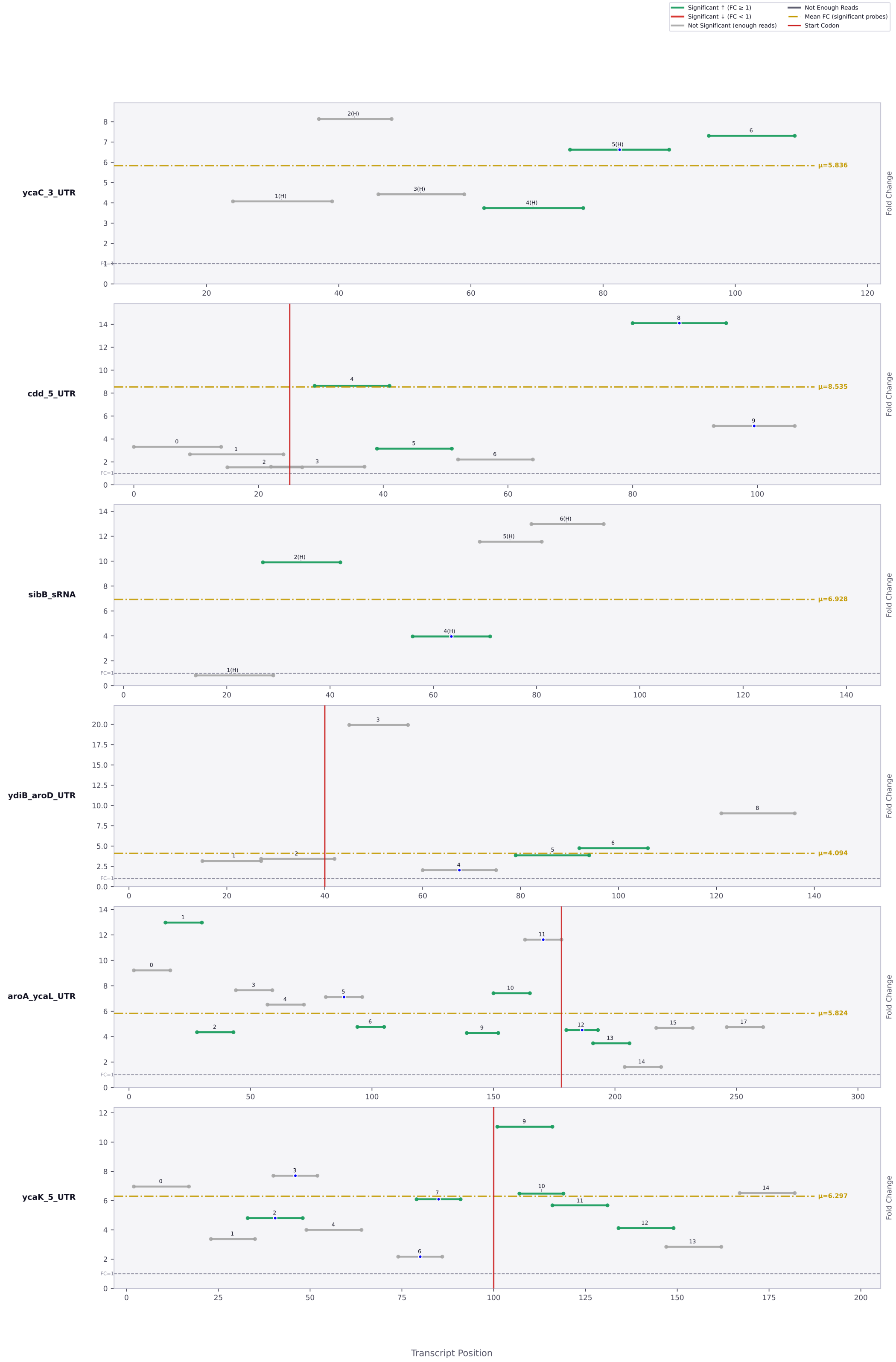

Significant ↑ (FC ≥ 1)
Significant ↓ (FC < 1)
Not Significant (enough reads)
Not Enough Reads
Mean FC (significant probes)
Start Codon
ycaC_3_UTR
μ=5.836
cdd_5_UTR
μ=8.535
sibB_sRNA
μ=6.928
ydiB_aroD_UTR
μ=4.094
aroA_ycaL_UTR
μ=5.824
ycaK_5_UTR
μ=6.297
Fold Change
Transcript Position

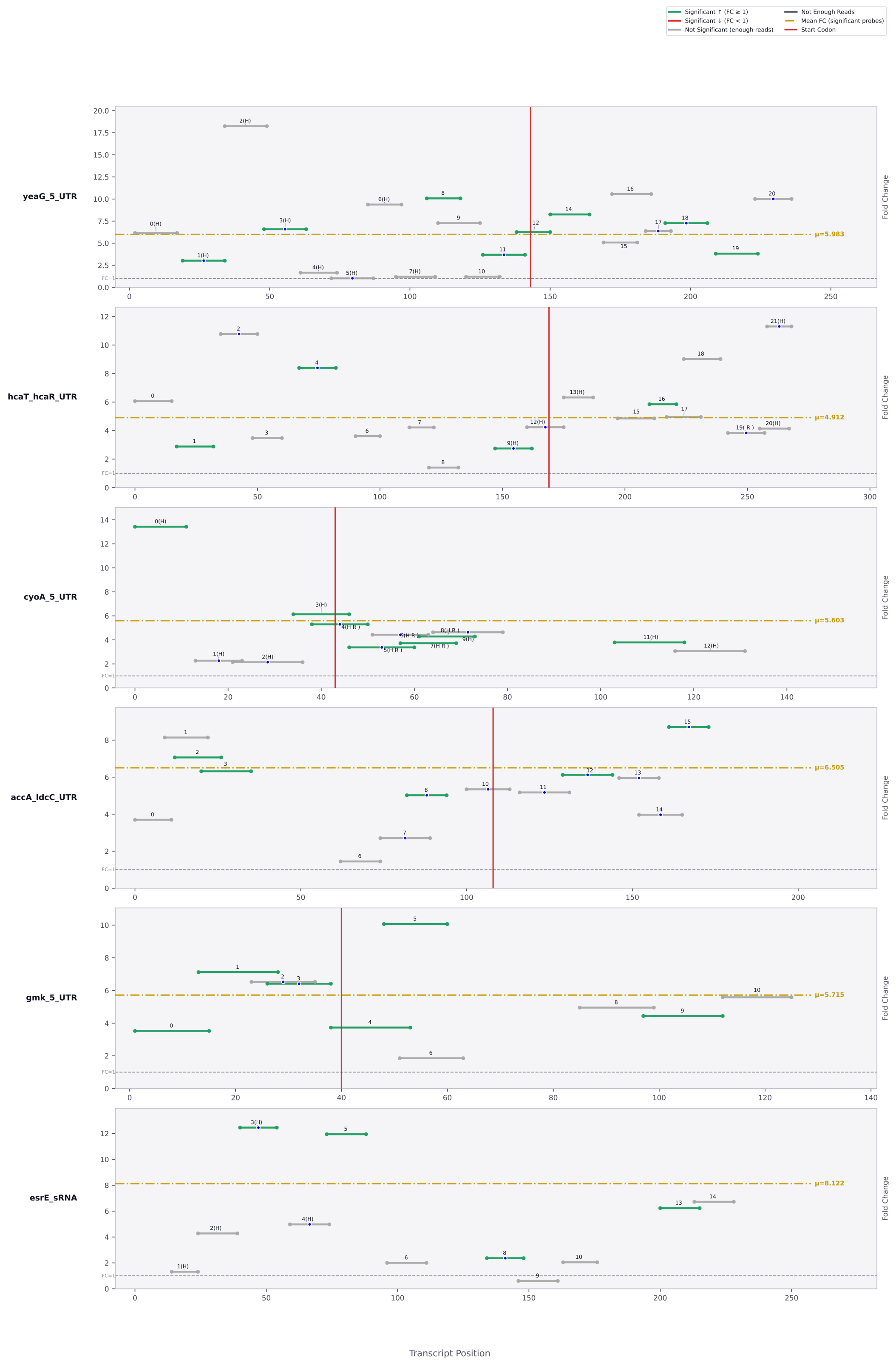

Significant ↑ (FC ≥ 1)
Significant ↓ (FC < 1)
Not Significant (enough reads)
Not Enough Reads
Mean FC (significant probes)
Start Codon
yeaG_5_UTR
μ=5.983
hcaT_hcaR_UTR
μ=4.912
cyoA_5_UTR
μ=5.603
accA_ldcC_UTR
μ=6.505
gmk_5_UTR
μ=5.715
esrE_sRNA
μ=8.122
Fold Change
FC=1
Transcript Position

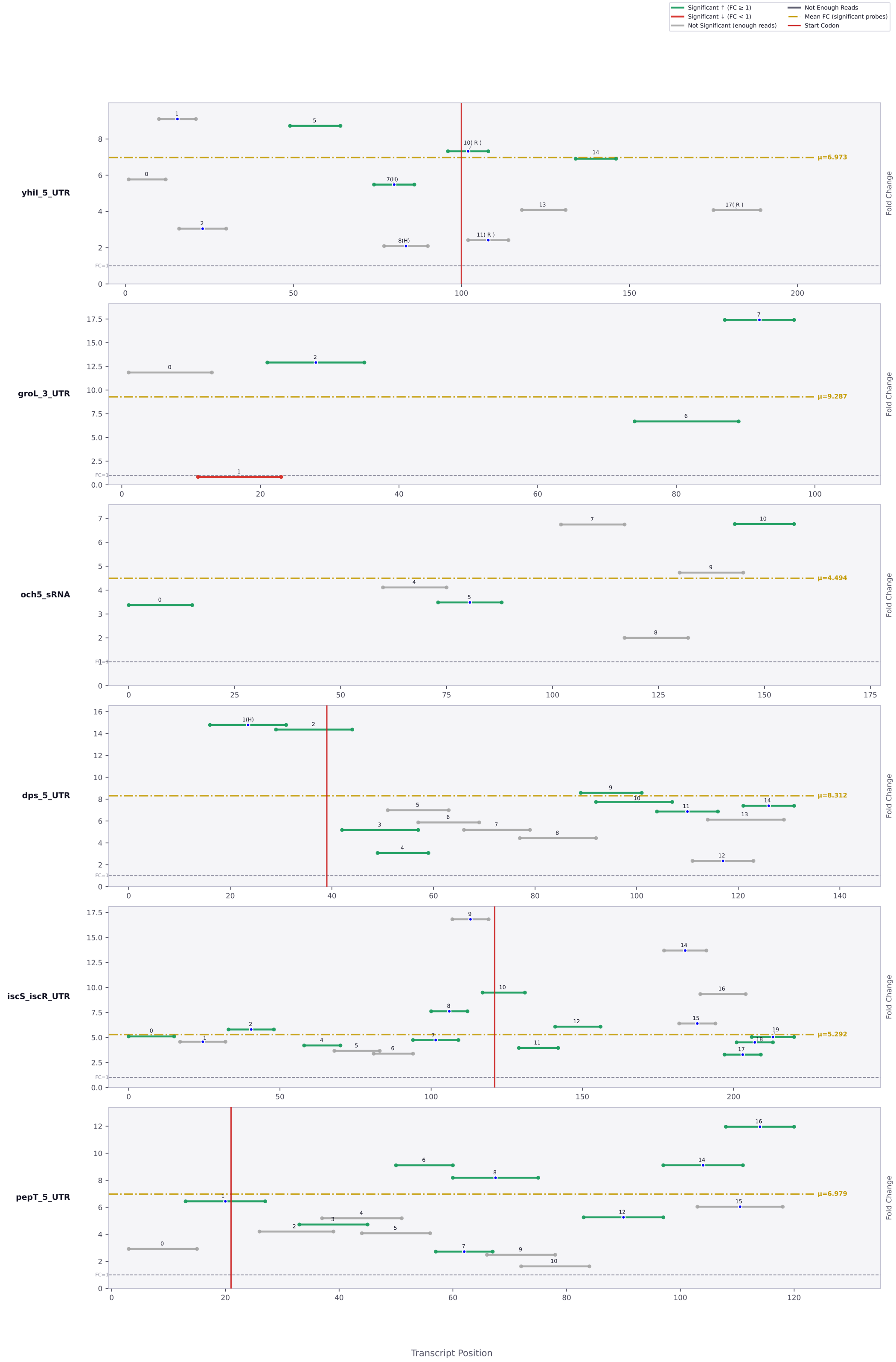

Significant ↑ (FC ≥ 1)
Significant ↓ (FC < 1)
Not Significant (enough reads)
Not Enough Reads
Mean FC (significant probes)
Start Codon
yhiI_5_UTR
μ=6.973
groL_3_UTR
μ=9.287
och5_sRNA
μ=4.494
dps_5_UTR
μ=8.312
iscS_iscR_UTR
μ=5.292
pepT_5_UTR
μ=6.979
Fold Change
Transcript Position

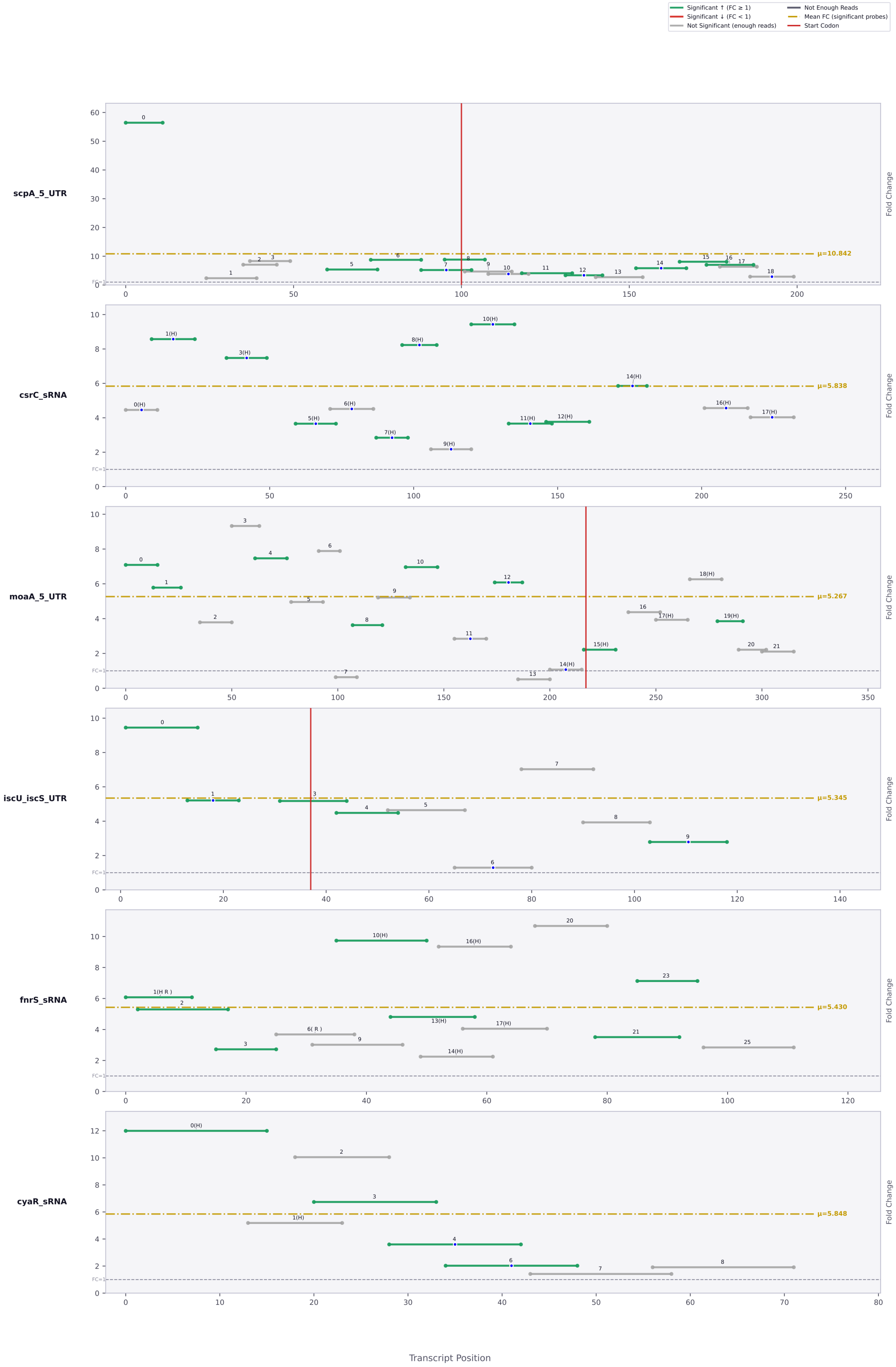

Significant ↑ (FC ≥ 1)
Significant ↓ (FC < 1)
Not Significant (enough reads)
Not Enough Reads
Mean FC (significant probes)
Start Codon
scpA_5_UTR
μ=10.842
csrC_sRNA
μ=5.838
moaA_5_UTR
μ=5.267
iscU_iscS_UTR
μ=5.345
fnrS_sRNA
μ=5.430
cyaR_sRNA
μ=5.848
Fold Change
Transcript Position

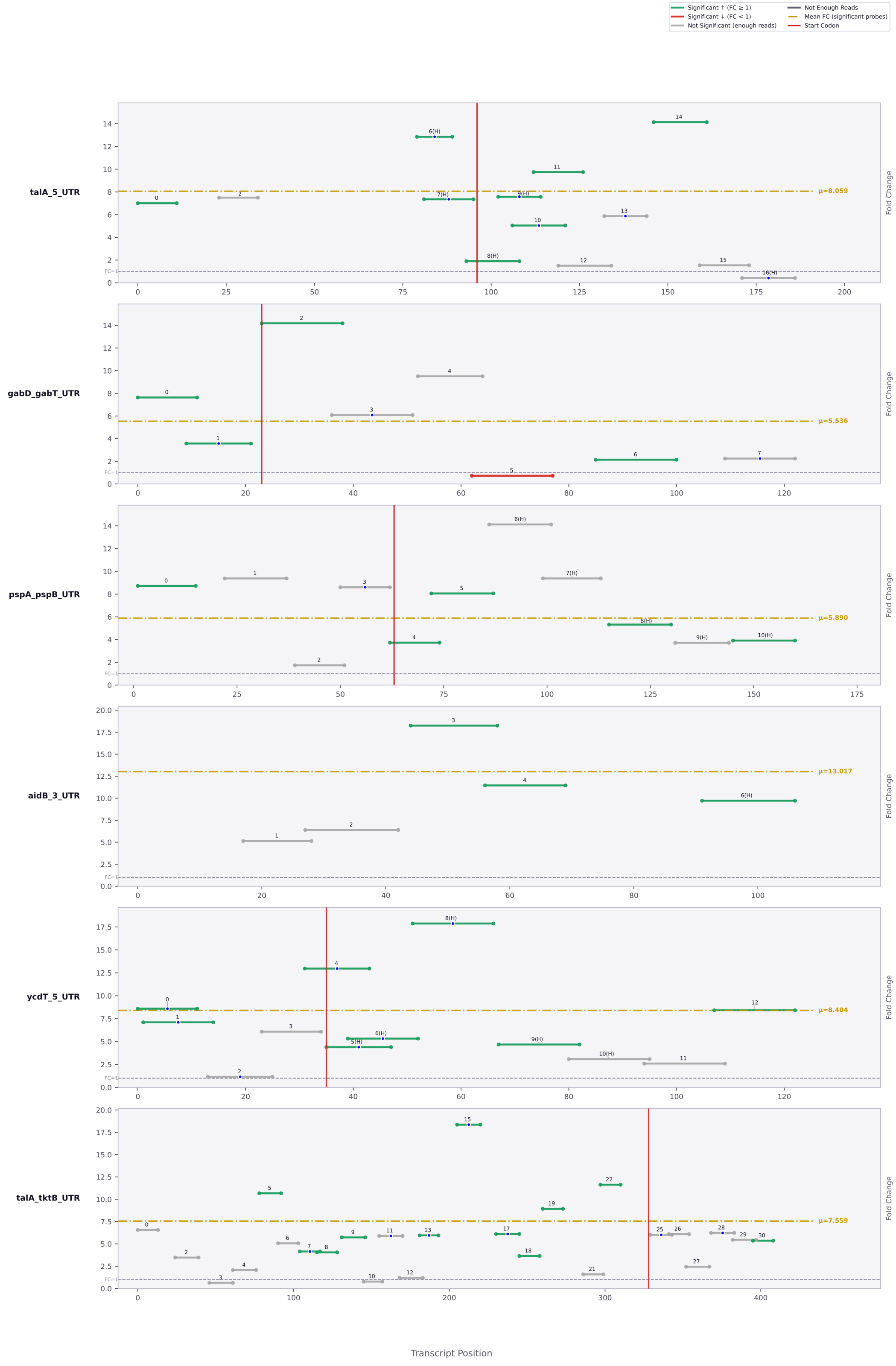

Significant ↑ (FC ≥ 1)
Significant ↓ (FC < 1)
Not Significant (enough reads)
Not Enough Reads
Mean FC (significant probes)
Start Codon
talA_5_UTR
μ=8.059
gabD_gabT_UTR
μ=5.536
pspA_pspB_UTR
μ=5.890
aidB_3_UTR
μ=13.017
ycdT_5_UTR
μ=8.404
talA_tktB_UTR
μ=7.559
Fold Change
Transcript Position

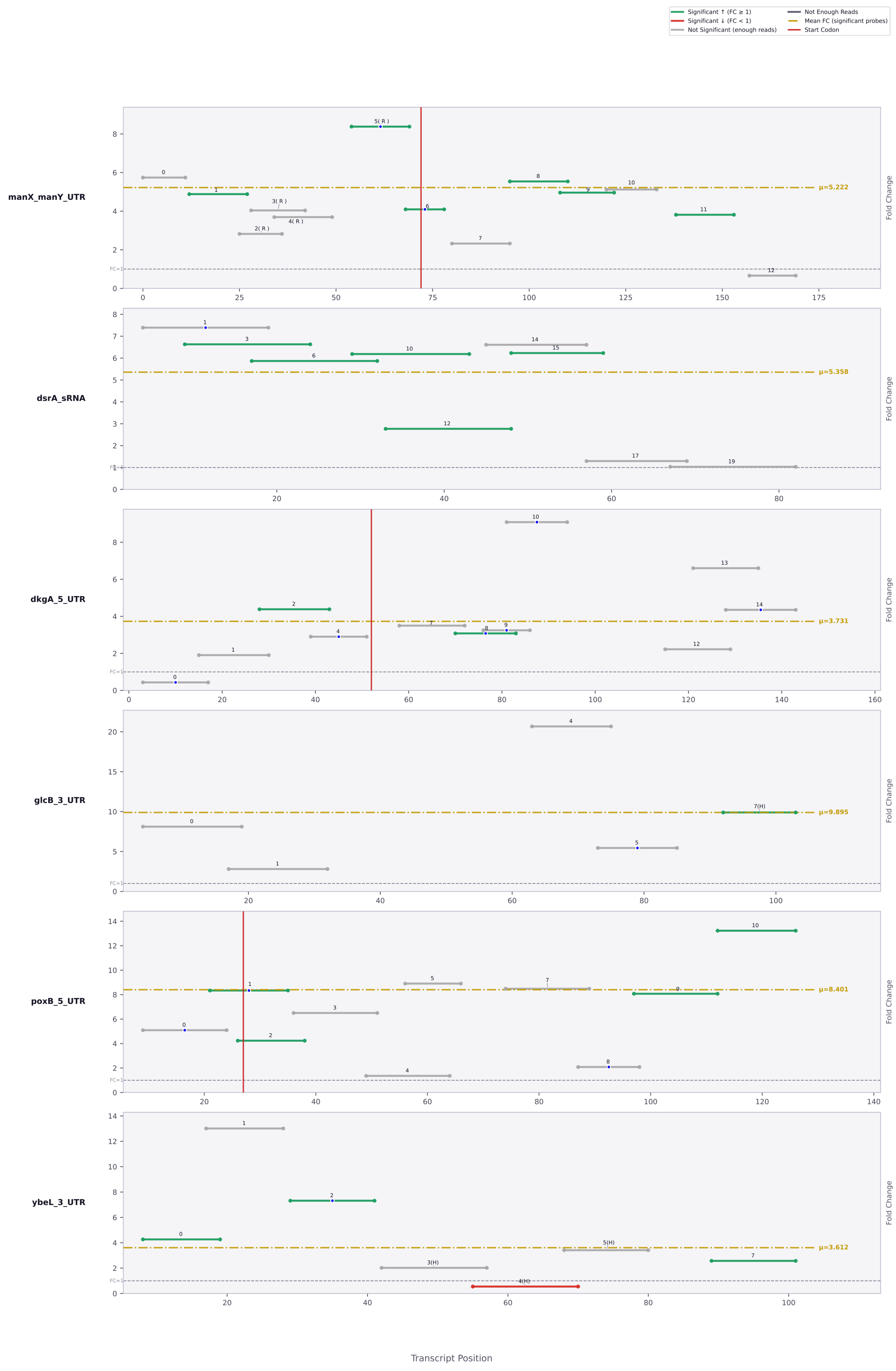

Significant ↑ (FC ≥ 1)
Significant ↓ (FC < 1)
Not Significant (enough reads)
Not Enough Reads
Mean FC (significant probes)
Start Codon
manX_manY_UTR
μ=5.222
dsrA_sRNA
μ=5.358
dkgA_5_UTR
μ=3.731
glcB_3_UTR
μ=9.895
poxB_5_UTR
μ=8.401
ybeL_3_UTR
μ=3.612
Fold Change
Transcript Position

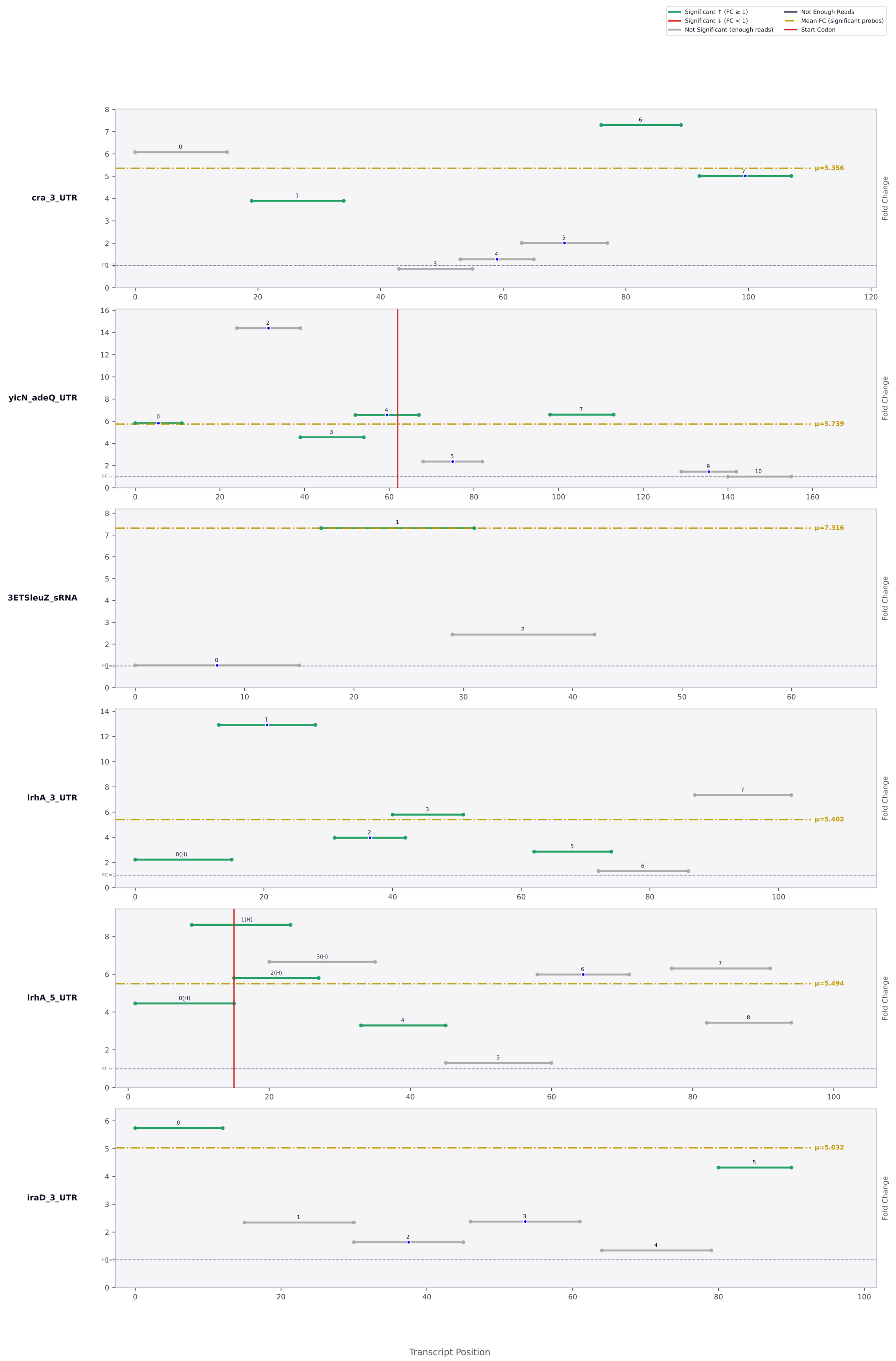

Significant ↑ (FC ≥ 1)
Significant ↓ (FC < 1)
Not Significant (enough reads)
Not Enough Reads
Mean FC (significant probes)
Start Codon
cra_3_UTR
μ=5.356
yicN_adeQ_UTR
μ=5.739
3ETSleuZ_sRNA
μ=7.316
lrhA_3_UTR
μ=5.402
lrhA_5_UTR
μ=5.494
iraD_3_UTR
μ=5.032
Fold Change
FC=1
Transcript Position

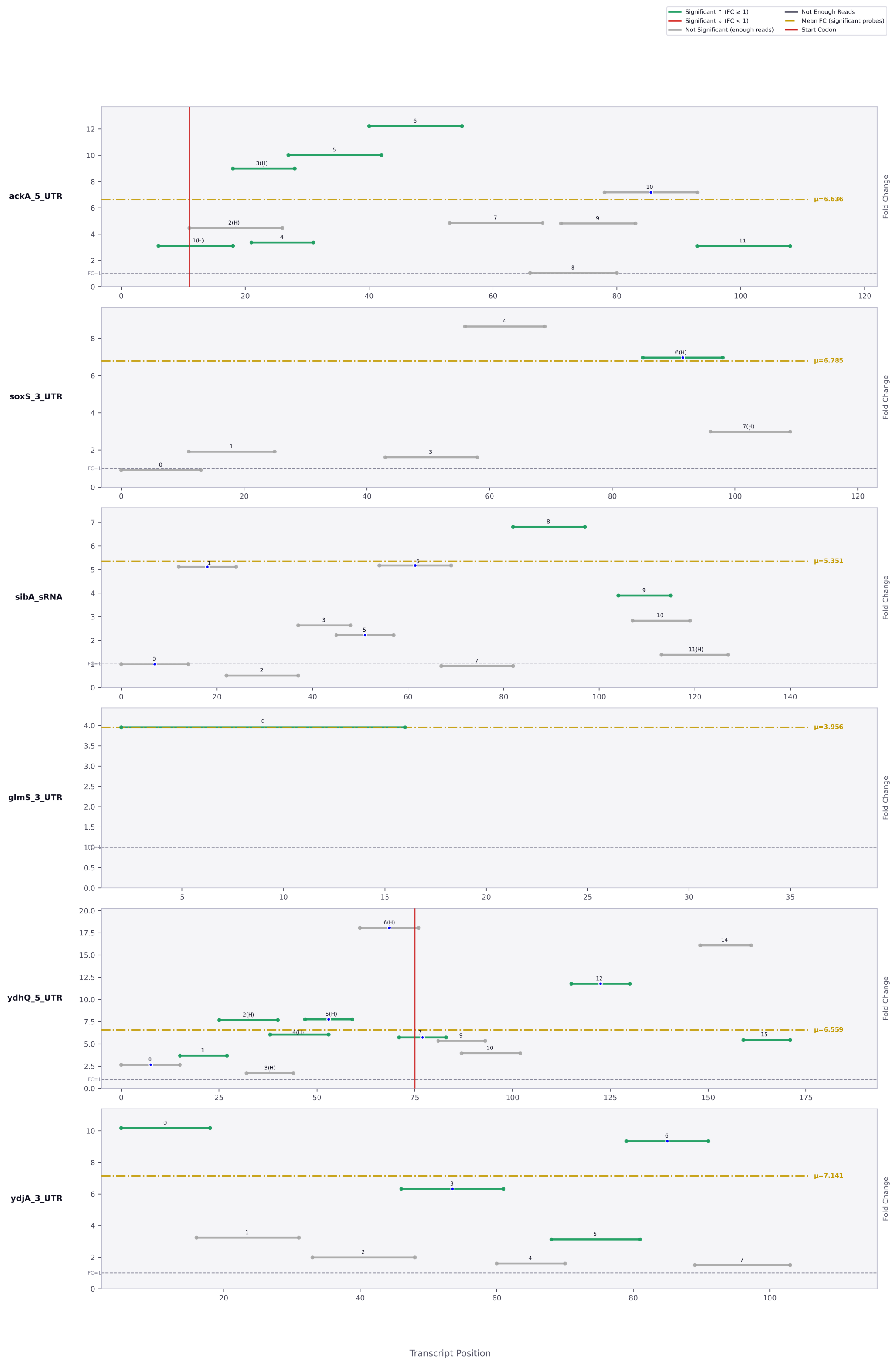

Significant ↑ (FC ≥ 1)
Significant ↓ (FC < 1)
Not Significant (enough reads)
Not Enough Reads
Mean FC (significant probes)
Start Codon
ackA_5_UTR
μ=6.636
soxS_3_UTR
μ=6.785
sibA_sRNA
μ=5.351
glmS_3_UTR
μ=3.956
ydhQ_5_UTR
μ=6.559
ydjA_3_UTR
μ=7.141
Fold Change
Transcript Position

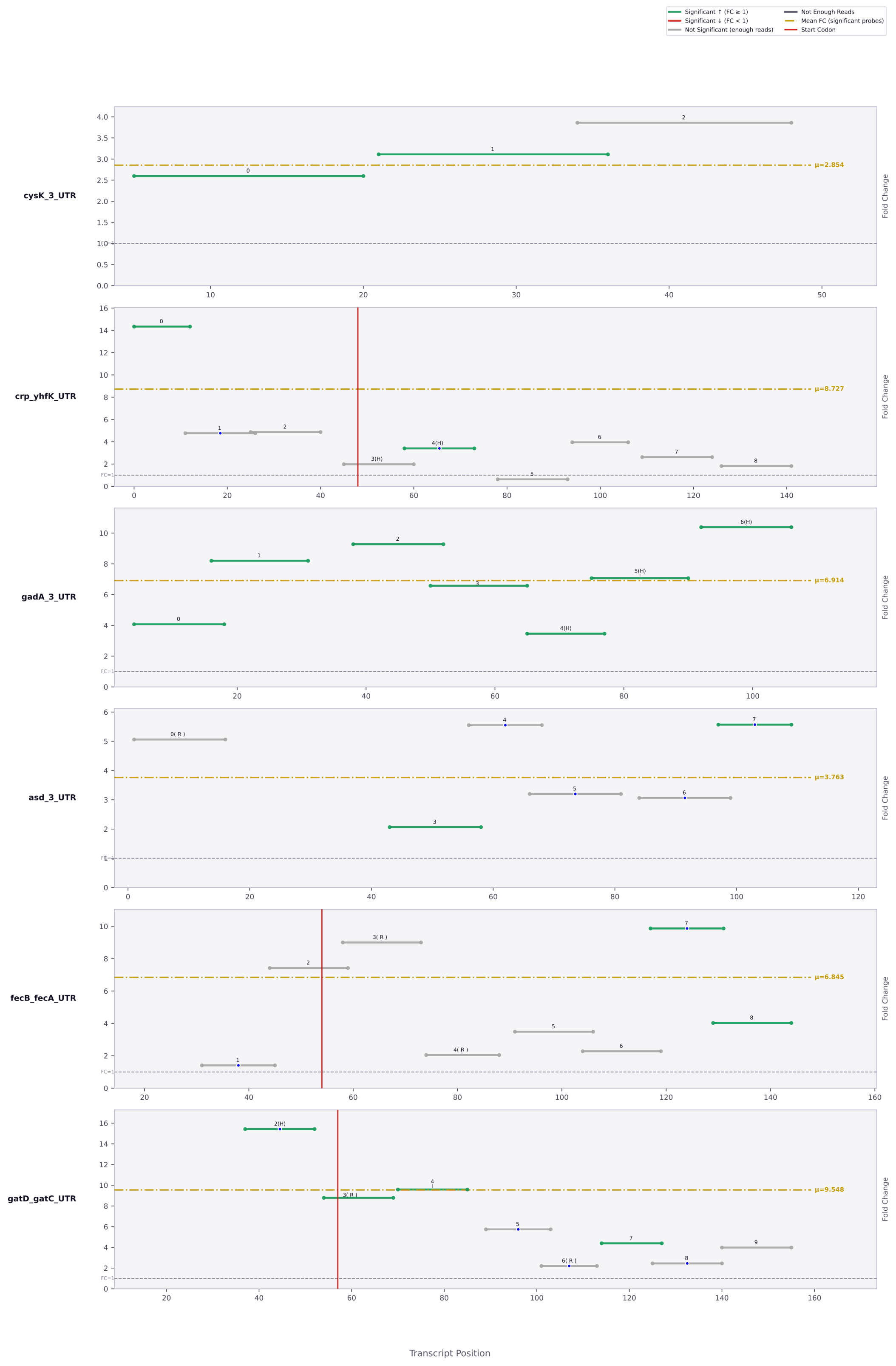

Significant ↑ (FC ≥ 1)
Significant ↓ (FC < 1)
Not Significant (enough reads)
Not Enough Reads
Mean FC (significant probes)
Start Codon
cysK_3_UTR
μ=2.854
crp_yhfK_UTR
μ=8.727
gadA_3_UTR
μ=6.914
asd_3_UTR
μ=3.763
fecB_fecA_UTR
μ=6.845
gatD_gatC_UTR
μ=9.548
Fold Change
Transcript Position

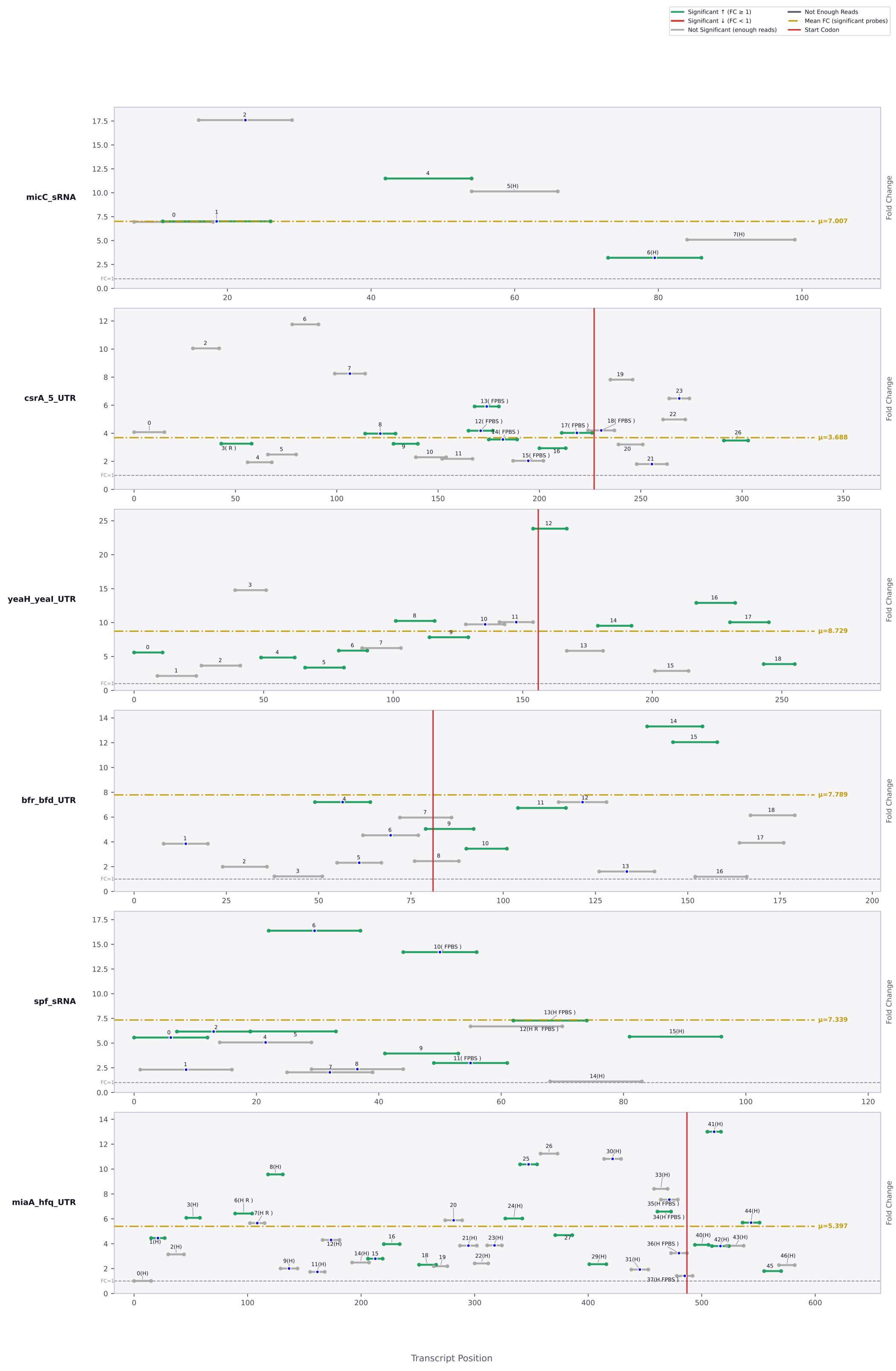

Significant ↑ (FC ≥ 1)
Significant ↓ (FC < 1)
Not Significant (enough reads)
Not Enough Reads
Mean FC (significant probes)
Start Codon
micC_sRNA
μ=7.007
csrA_5_UTR
μ=3.688
yeaH_yeaI_UTR
μ=8.729
bfr_bfd_UTR
μ=7.789
spf_sRNA
μ=7.339
miaA_hfq_UTR
μ=5.397
Fold Change
Transcript Position

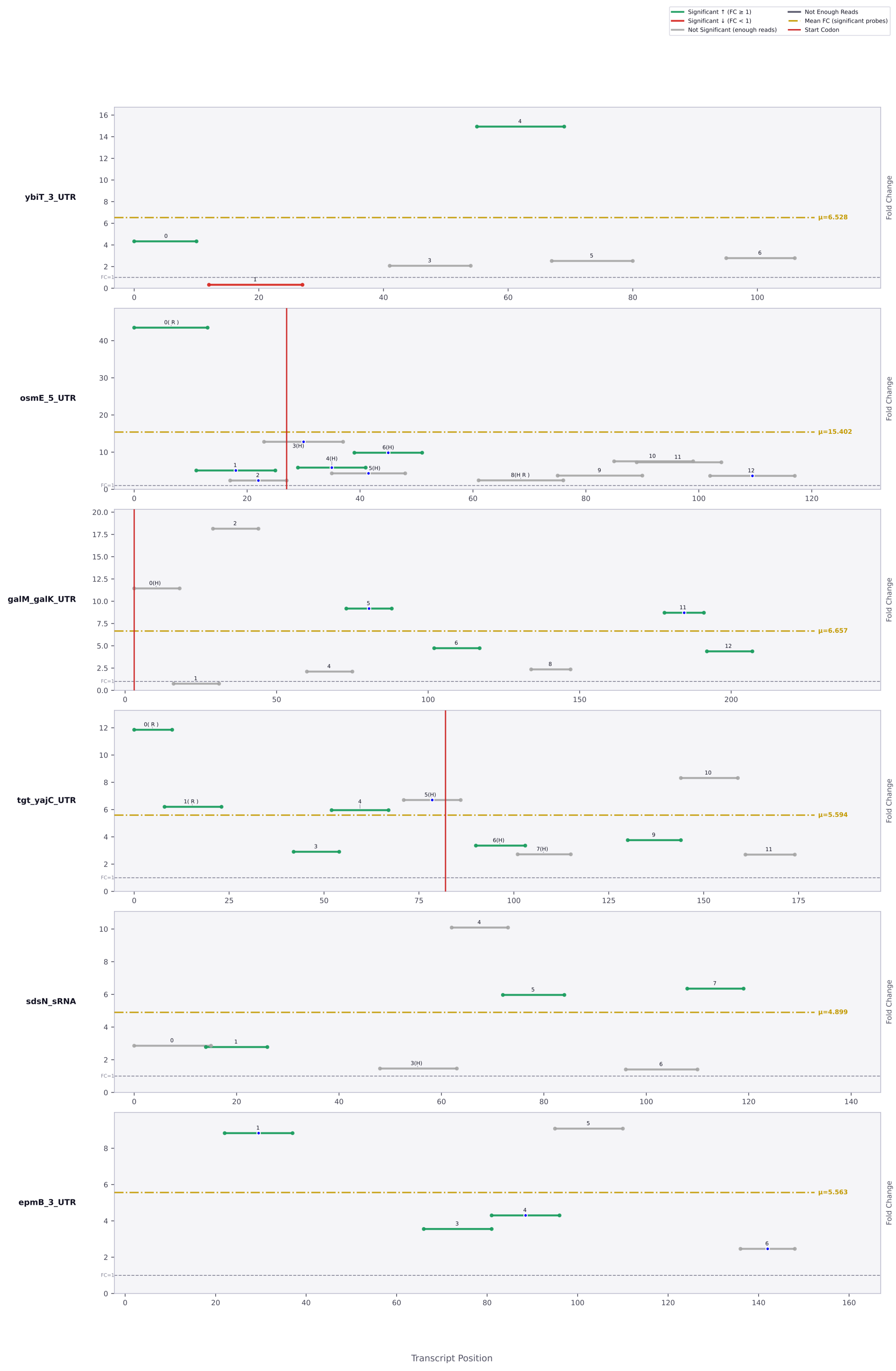

Significant ↑ (FC ≥ 1)
Significant ↓ (FC < 1)
Not Significant (enough reads)
Not Enough Reads
Mean FC (significant probes)
Start Codon
ybiT_3_UTR
μ=6.528
osmE_5_UTR
μ=15.402
galM_galK_UTR
μ=6.657
tgt_yajC_UTR
μ=5.594
sdsN_sRNA
μ=4.899
epmB_3_UTR
μ=5.563
Fold Change
FC=1
Transcript Position

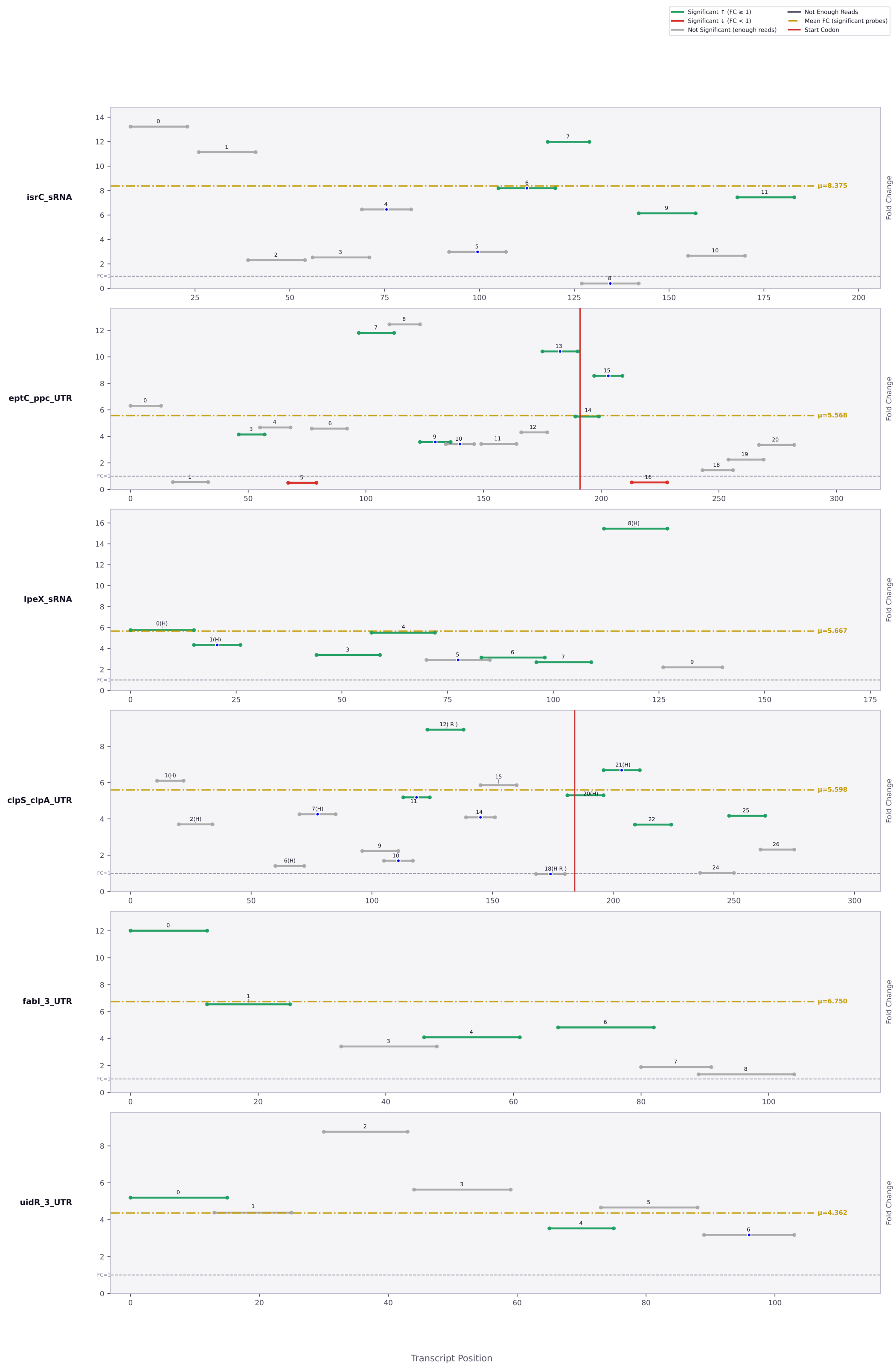

Significant ↑ (FC ≥ 1)
Significant ↓ (FC < 1)
Not Significant (enough reads)
Not Enough Reads
Mean FC (significant probes)
Start Codon
isrC_sRNA
μ=8.375
eptC_ppc_UTR
μ=5.568
lpeX_sRNA
μ=5.667
clpS_clpA_UTR
μ=5.598
fabI_3_UTR
μ=6.750
uidR_3_UTR
μ=4.362
Fold Change
FC=1
Transcript Position

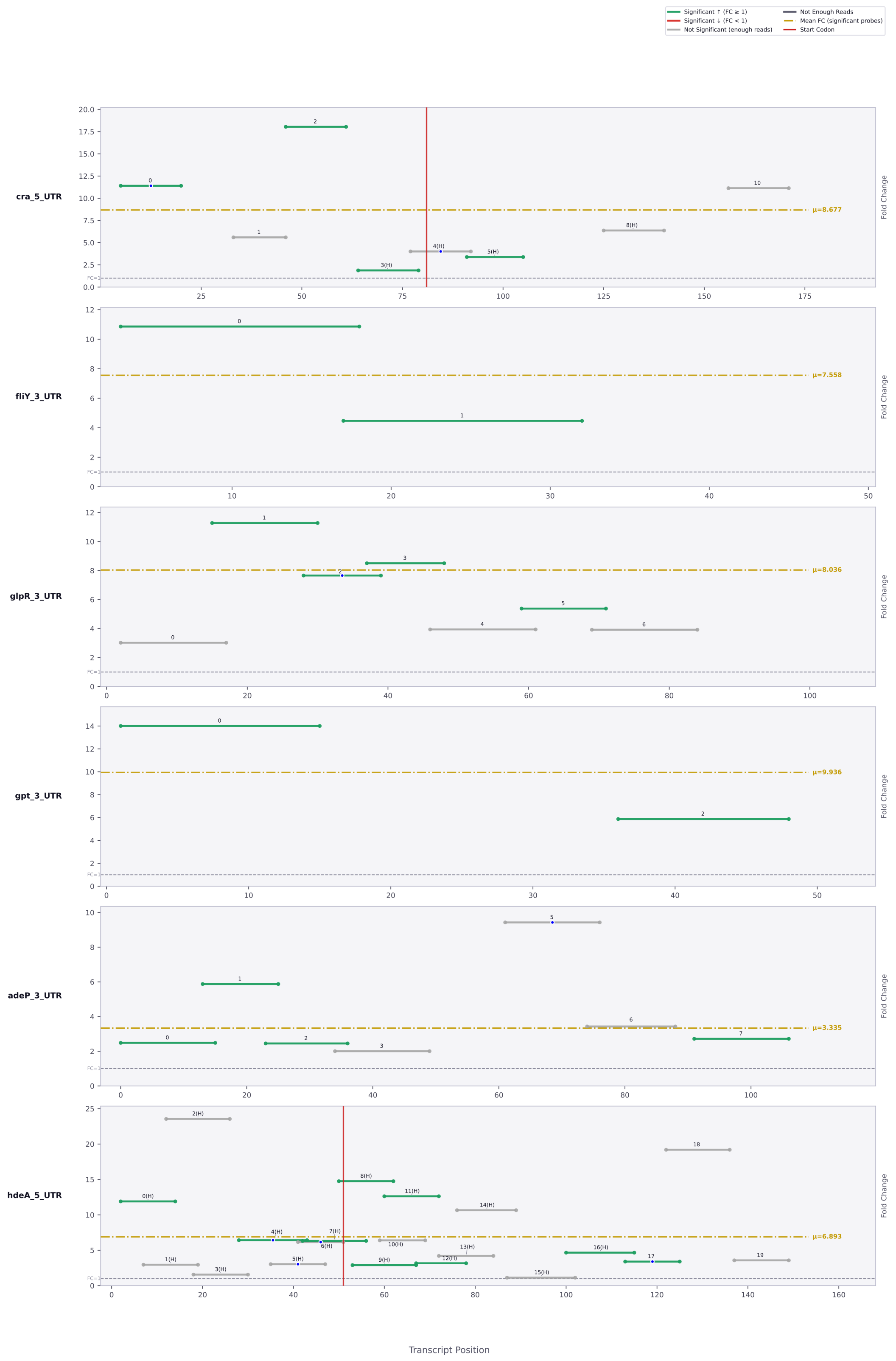
Significant ↑ (FC ≥ 1)
Significant ↓ (FC < 1)
Not Significant (enough reads)
Not Enough Reads
Mean FC (significant probes)
Start Codon
cra_5_UTR
μ=8.677
fliY_3_UTR
μ=7.558
glpR_3_UTR
μ=8.036
gpt_3_UTR
μ=9.936
adeP_3_UTR
μ=3.335
hdeA_5_UTR
μ=6.893
Fold Change
FC=1
Transcript Position

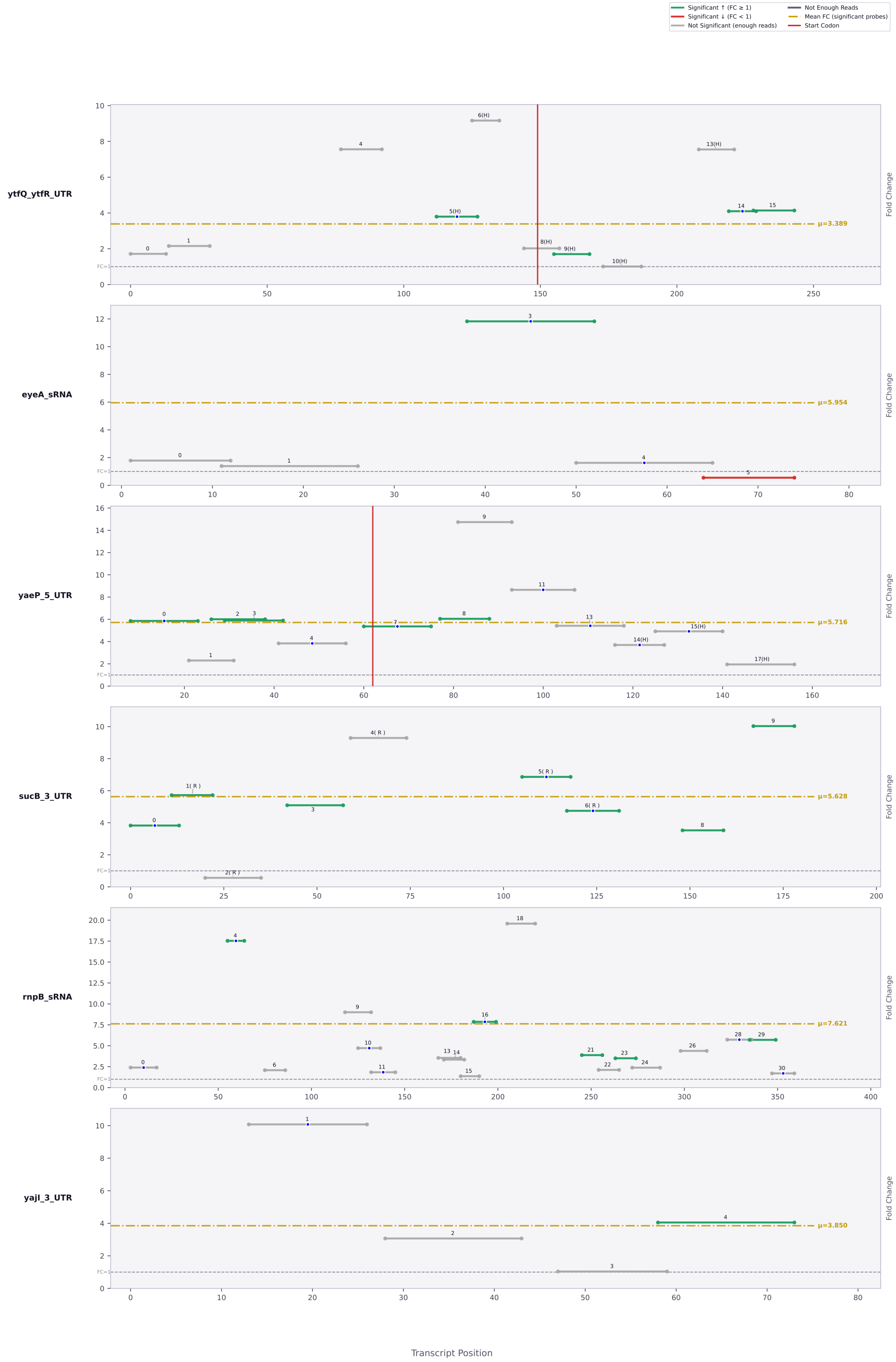

Significant ↑ (FC ≥ 1)
Significant ↓ (FC < 1)
Not Significant (enough reads)
Not Enough Reads
Mean FC (significant probes)
Start Codon
ytfQ_ytfR_UTR
μ=3.389
eyeA_sRNA
μ=5.954
yaeP_5_UTR
μ=5.716
sucB_3_UTR
μ=5.628
rnpB_sRNA
μ=7.621
yajI_3_UTR
μ=3.850
Fold Change
Transcript Position

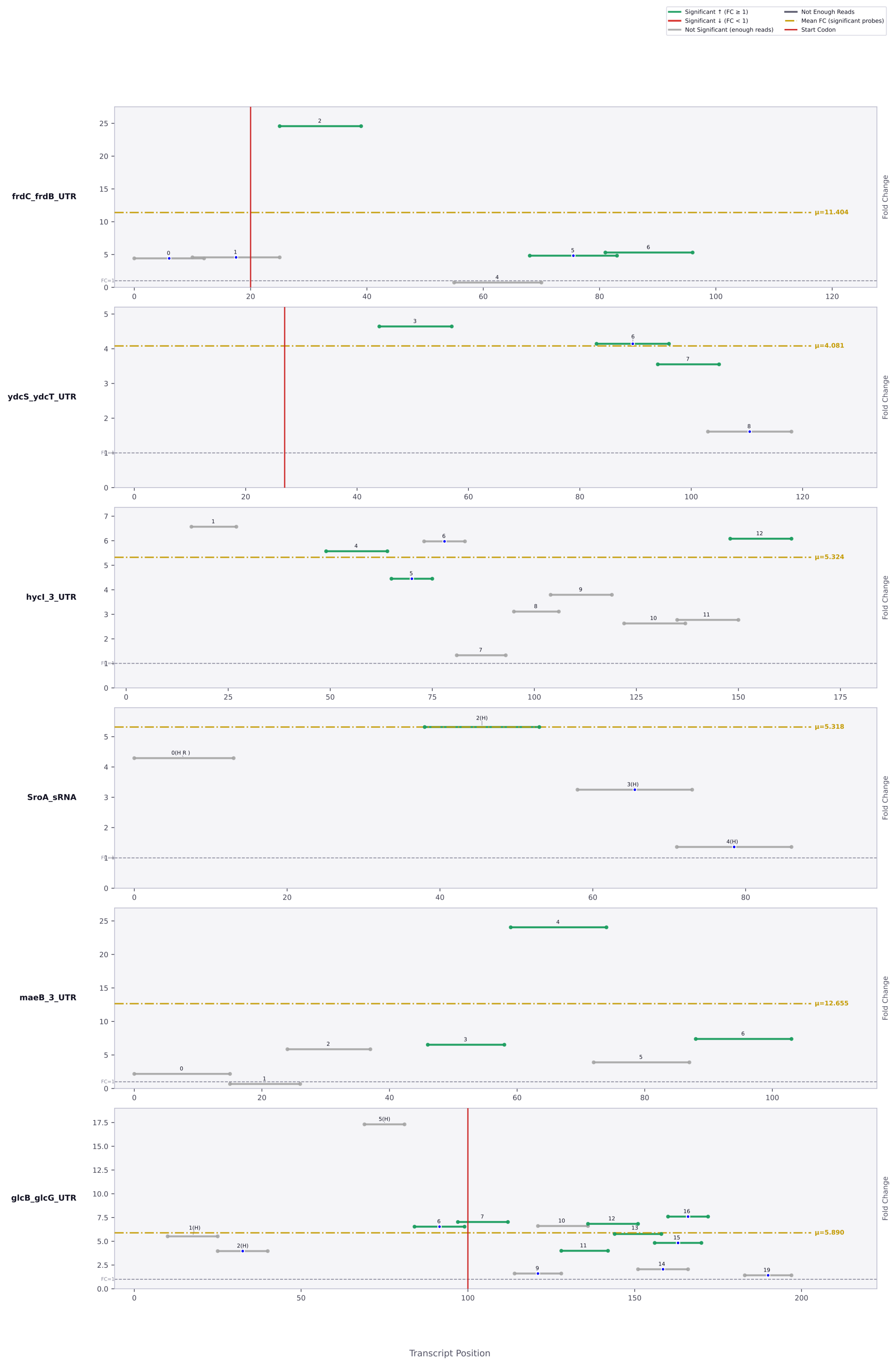

Significant ↑ (FC ≥ 1)
Significant ↓ (FC < 1)
Not Significant (enough reads)
Not Enough Reads
Mean FC (significant probes)
Start Codon
frdC_frdB_UTR
μ=11.404
ydcS_ydcT_UTR
μ=4.081
hycI_3_UTR
μ=5.324
SroA_sRNA
μ=5.318
maeB_3_UTR
μ=12.655
glcB_glcG_UTR
μ=5.890
Fold Change
Transcript Position

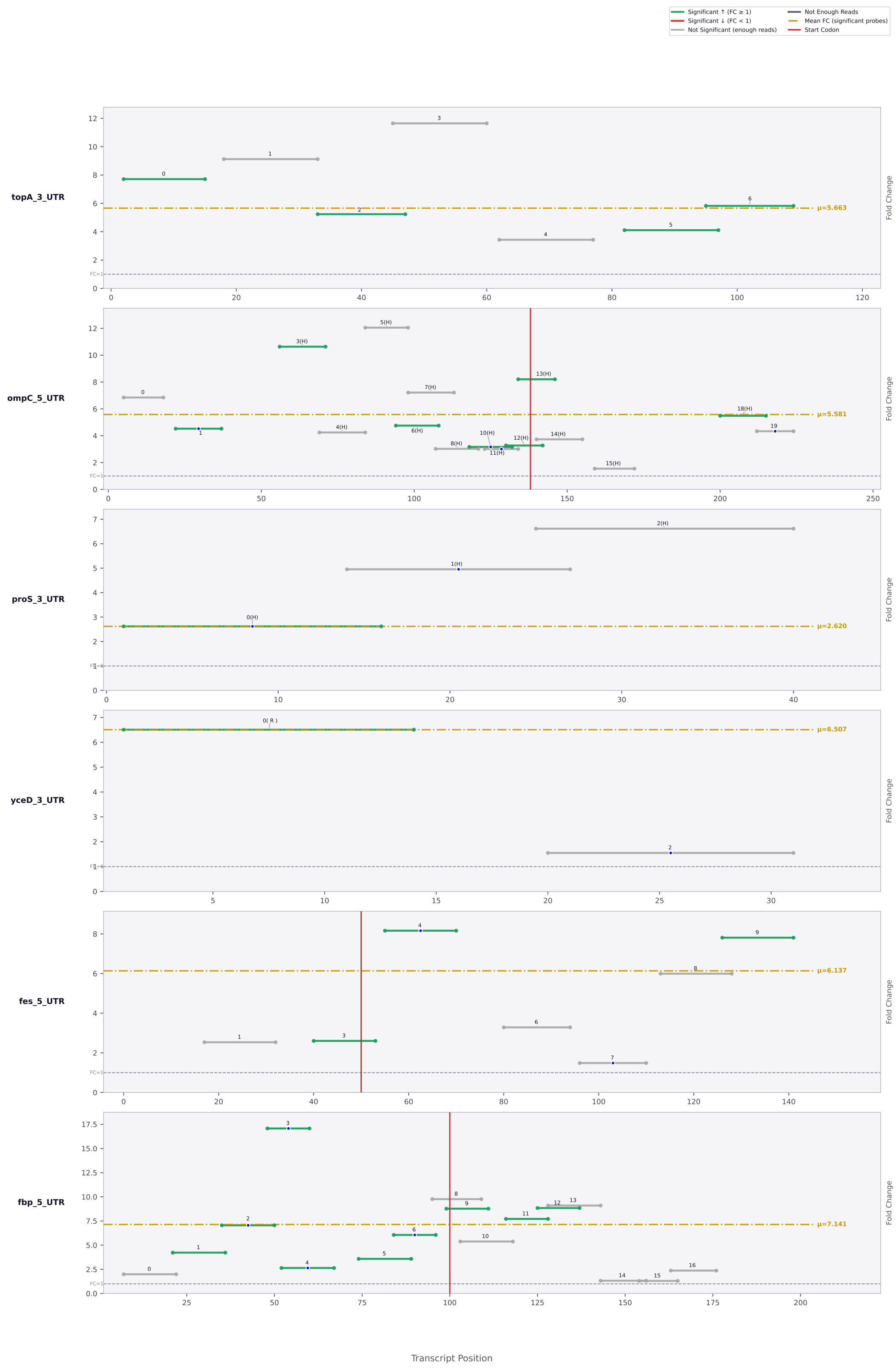

Significant ↑ (FC ≥ 1)
Significant ↓ (FC < 1)
Not Significant (enough reads)
Not Enough Reads
Mean FC (significant probes)
Start Codon
topA_3_UTR
μ=5.663
ompC_5_UTR
μ=5.581
proS_3_UTR
μ=2.620
yceD_3_UTR
μ=6.507
fes_5_UTR
μ=6.137
fbp_5_UTR
μ=7.141
Fold Change
FC=1
Transcript Position

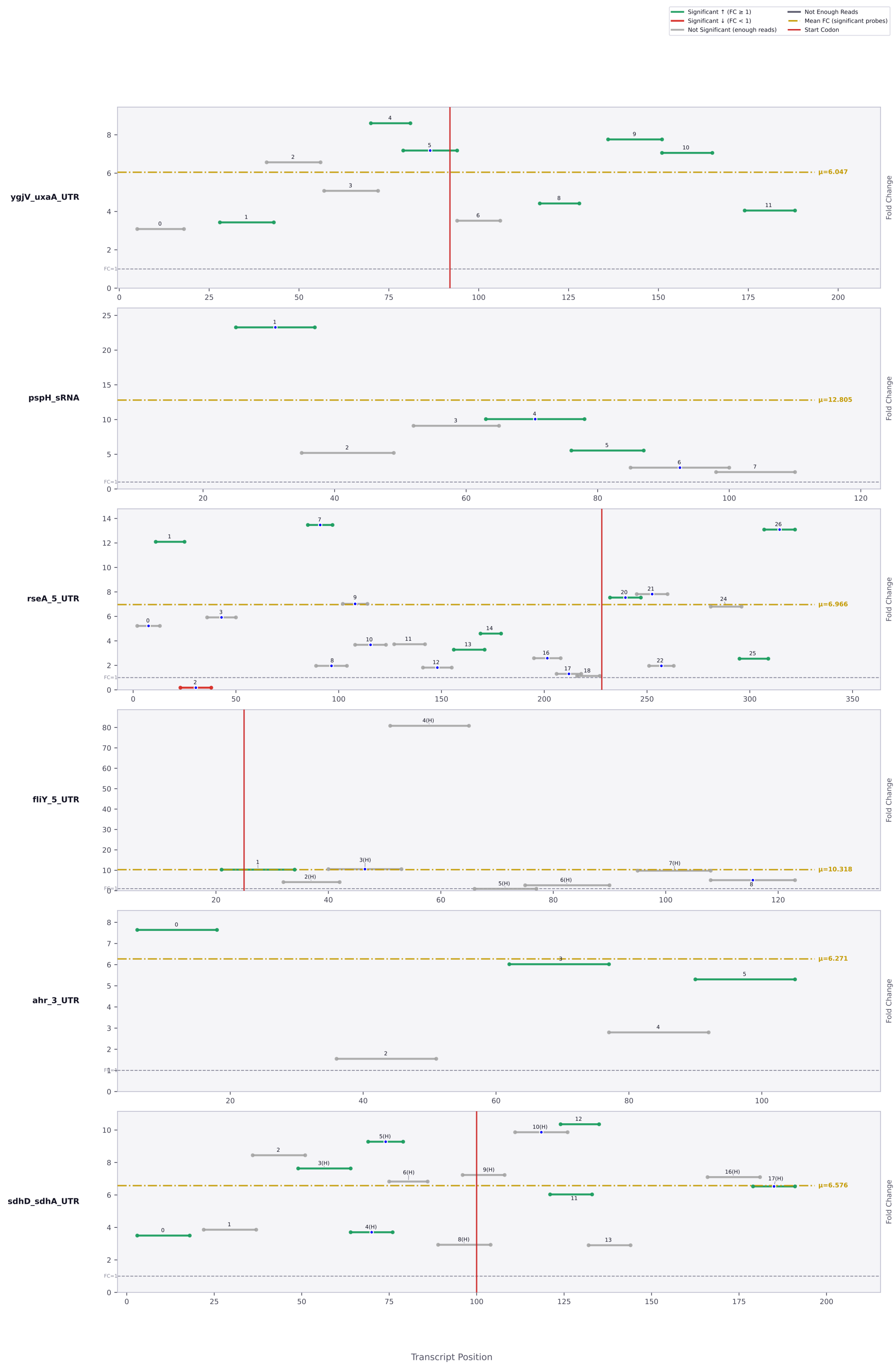

Significant ↑ (FC ≥ 1)
Significant ↓ (FC < 1)
Not Significant (enough reads)
Not Enough Reads
Mean FC (significant probes)
Start Codon
ygjV_uxaA_UTR
μ=6.047
pspH_sRNA
μ=12.805
rseA_5_UTR
μ=6.966
fliY_5_UTR
μ=10.318
ahr_3_UTR
μ=6.271
sdhD_sdhA_UTR
μ=6.576
Fold Change
Transcript Position

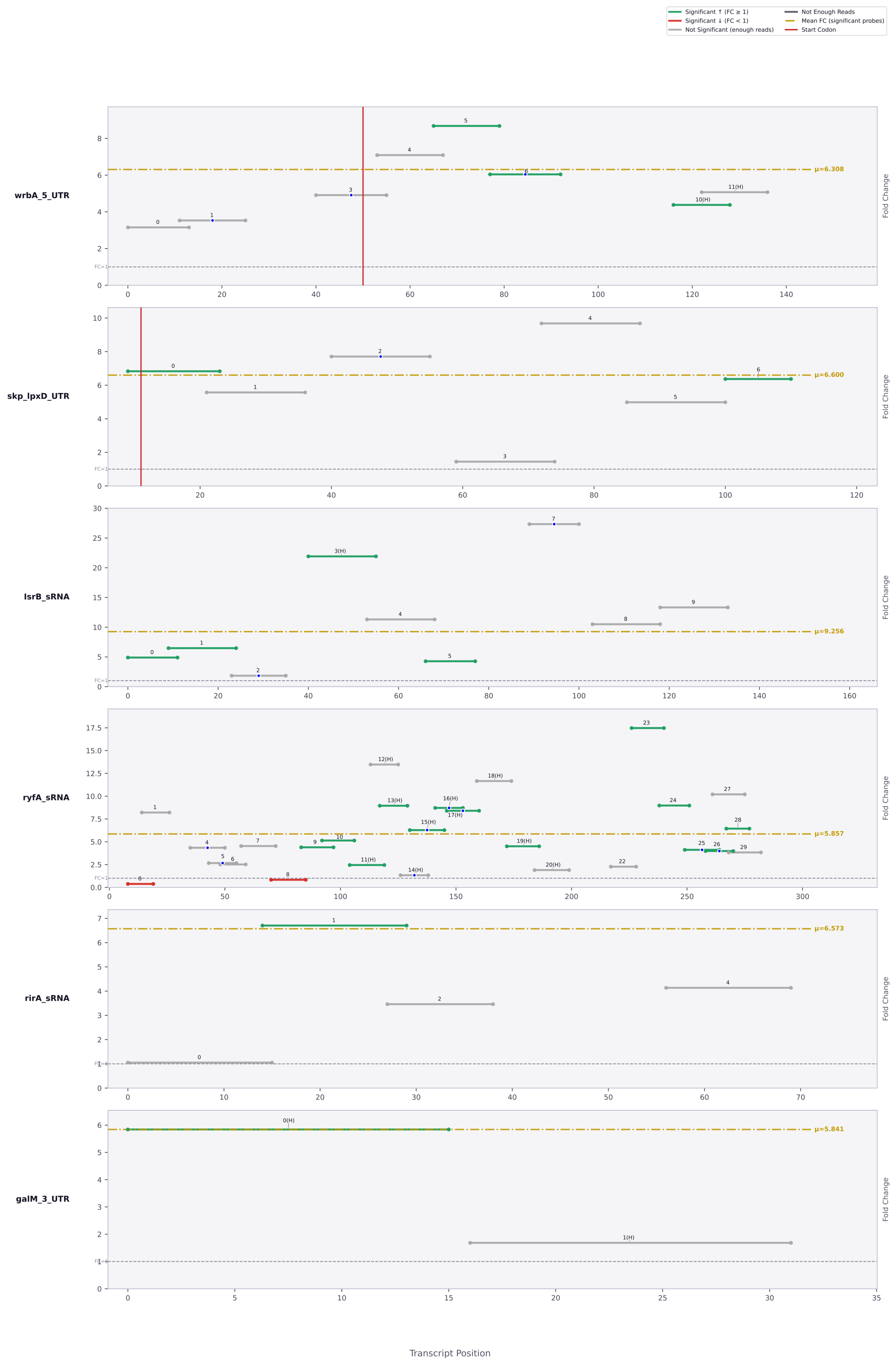
Significant ↑ (FC ≥ 1)
Significant ↓ (FC < 1)
Not Significant (enough reads)
Not Enough Reads
Mean FC (significant probes)
Start Codon
wrbA_5_UTR
μ=6.308
skp_lpxD_UTR
μ=6.600
lsrB_sRNA
μ=9.256
ryfA_sRNA
μ=5.857
rirA_sRNA
μ=6.573
galM_3_UTR
μ=5.841
Fold Change
FC=1
Transcript Position

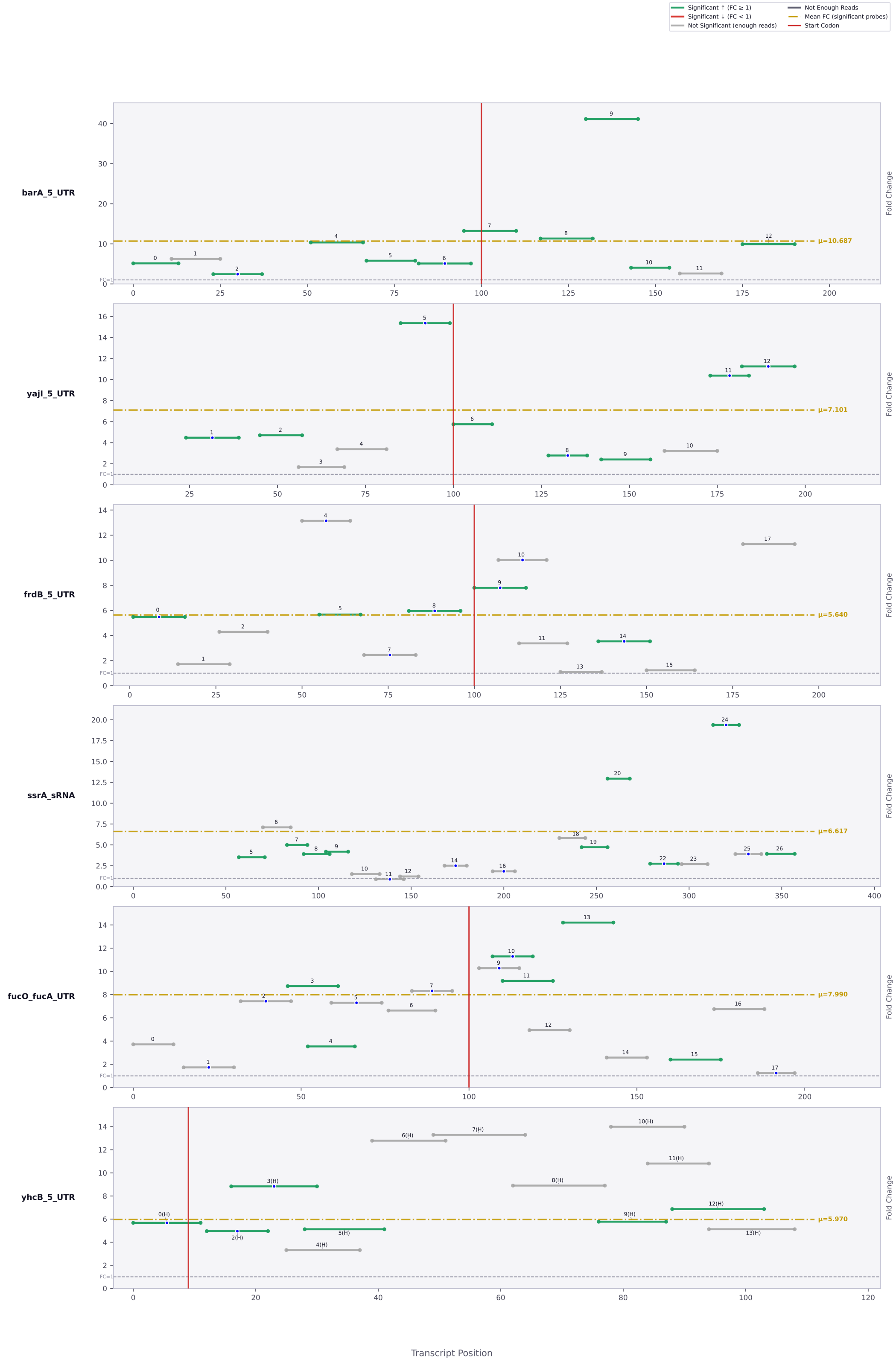

Significant ↑ (FC ≥ 1)
Significant ↓ (FC < 1)
Not Significant (enough reads)
Not Enough Reads
Mean FC (significant probes)
Start Codon
barA_5_UTR
μ=10.687
yajI_5_UTR
μ=7.101
frdB_5_UTR
μ=5.640
ssrA_sRNA
μ=6.617
fucO_fucA_UTR
μ=7.990
yhcB_5_UTR
μ=5.970
Fold Change
FC=1
Transcript Position

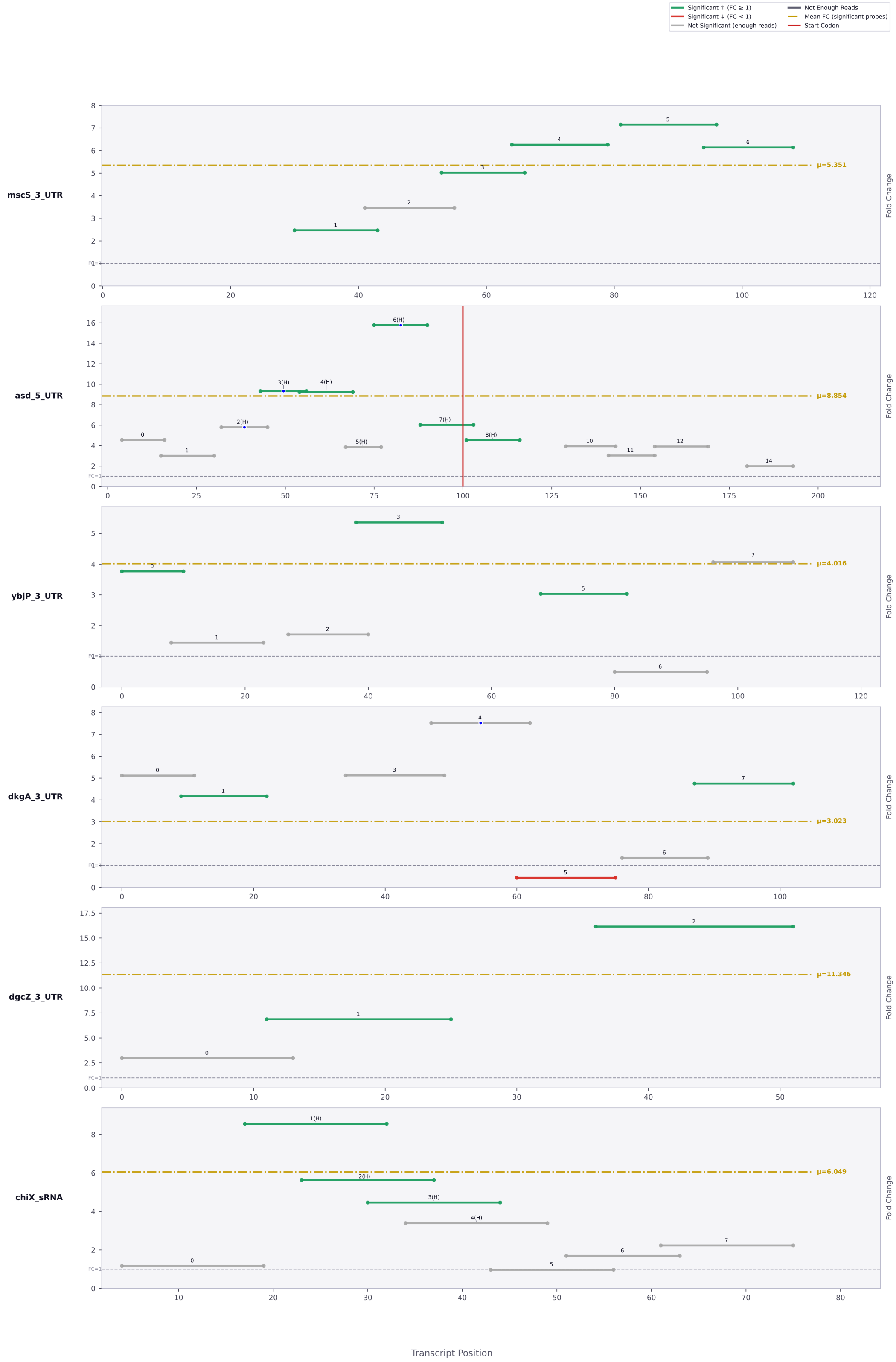

Significant ↑ (FC ≥ 1)
Significant ↓ (FC < 1)
Not Significant (enough reads)
Not Enough Reads
Mean FC (significant probes)
Start Codon
mscS_3_UTR
μ=5.351
asd_5_UTR
μ=8.854
ybjP_3_UTR
μ=4.016
dkgA_3_UTR
μ=3.023
dgcZ_3_UTR
μ=11.346
chiX_sRNA
μ=6.049
Fold Change
Transcript Position

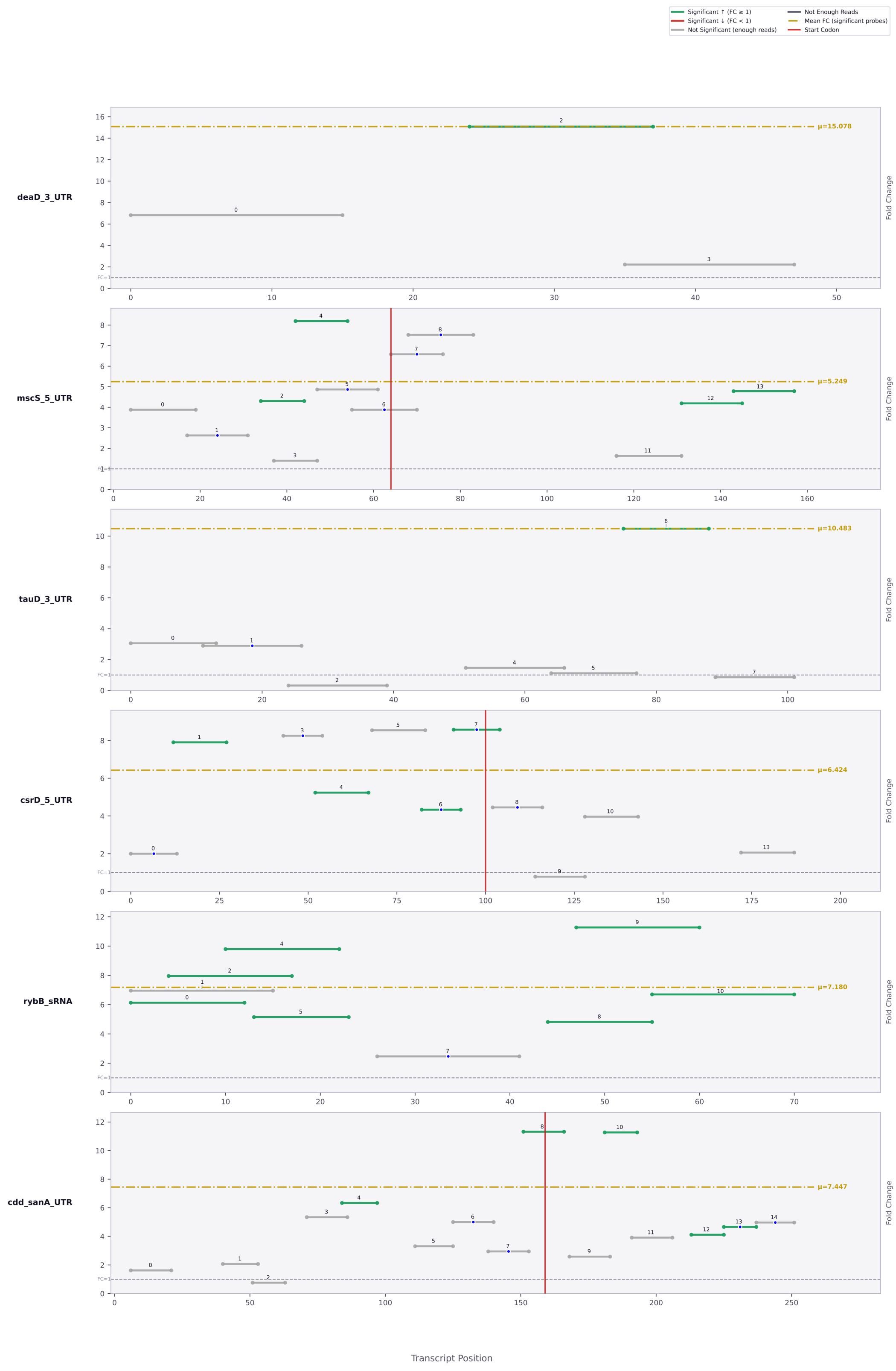

Significant ↑ (FC ≥ 1)
Significant ↓ (FC < 1)
Not Significant (enough reads)
Not Enough Reads
Mean FC (significant probes)
Start Codon
deaD_3_UTR
μ=15.078
mscS_5_UTR
μ=5.249
tauD_3_UTR
μ=10.483
csrD_5_UTR
μ=6.424
rybB_sRNA
μ=7.180
cdd_sanA_UTR
μ=7.447
Fold Change
Transcript Position

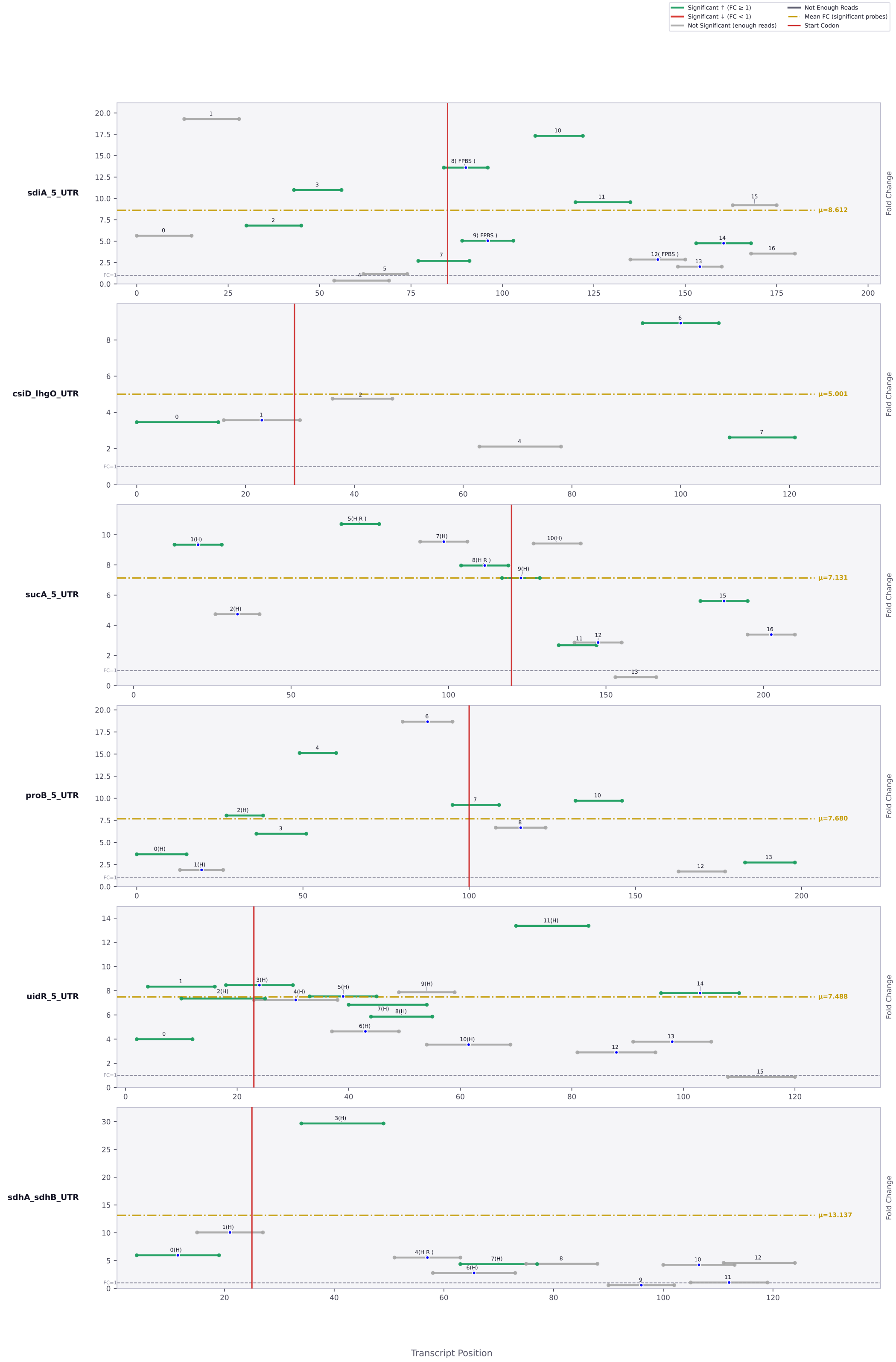

Significant ↑ (FC ≥ 1)
Significant ↓ (FC < 1)
Not Significant (enough reads)
Not Enough Reads
Mean FC (significant probes)
Start Codon
sdiA_5_UTR
μ=8.612
csiD_lhgO_UTR
μ=5.001
sucA_5_UTR
μ=7.131
proB_5_UTR
μ=7.680
uidR_5_UTR
μ=7.488
sdhA_sdhB_UTR
μ=13.137
Fold Change
FC=1
Transcript Position

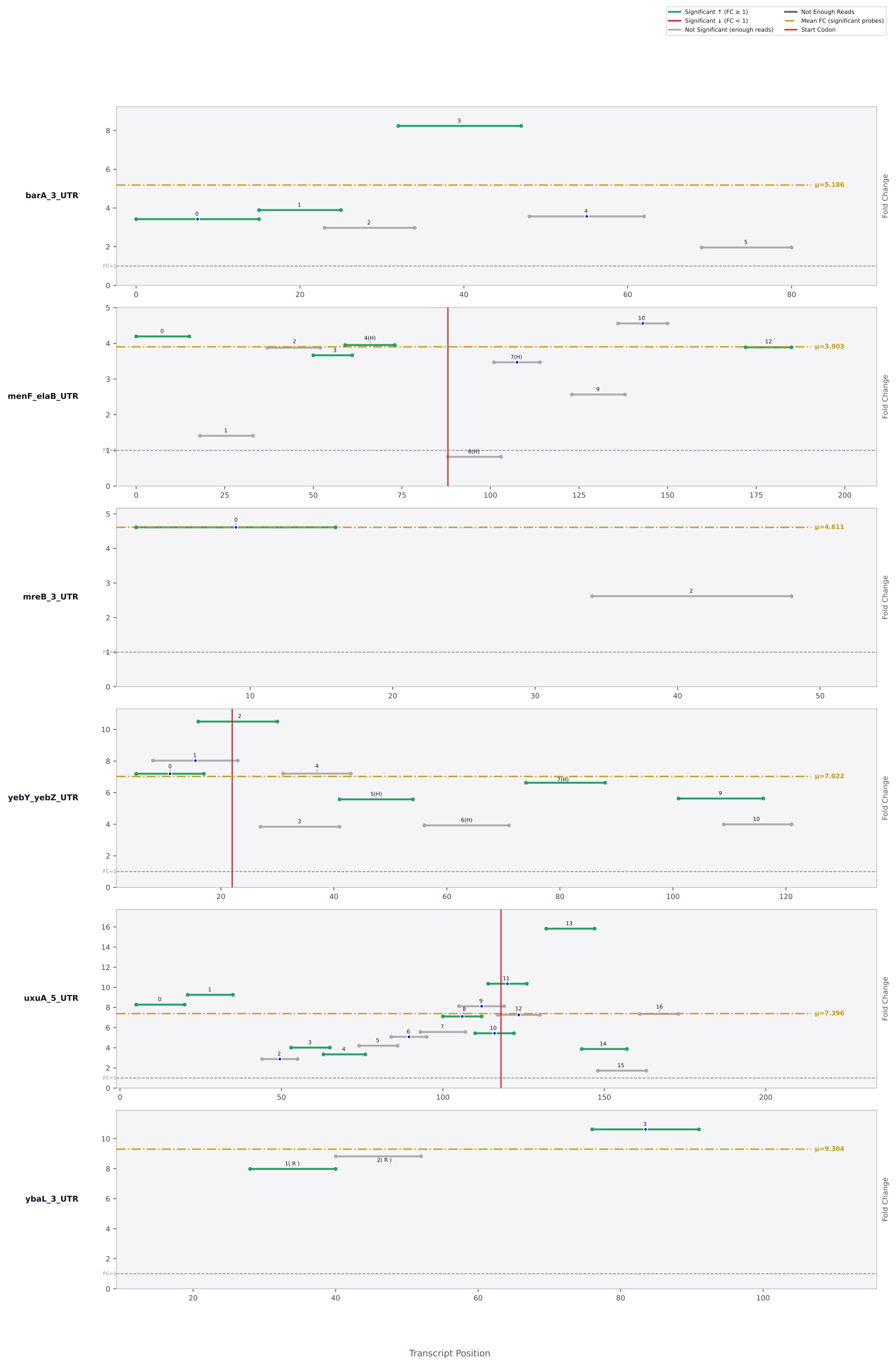

Significant ↑ (FC ≥ 1)
Significant ↓ (FC < 1)
Not Significant (enough reads)
Not Enough Reads
Mean FC (significant probes)
Start Codon
barA_3_UTR
μ=5.186
menF_elaB_UTR
μ=3.903
mreB_3_UTR
μ=4.611
yebY_yebZ_UTR
μ=7.022
uxuA_5_UTR
μ=7.396
ybaL_3_UTR
μ=9.304
Fold Change
Transcript Position

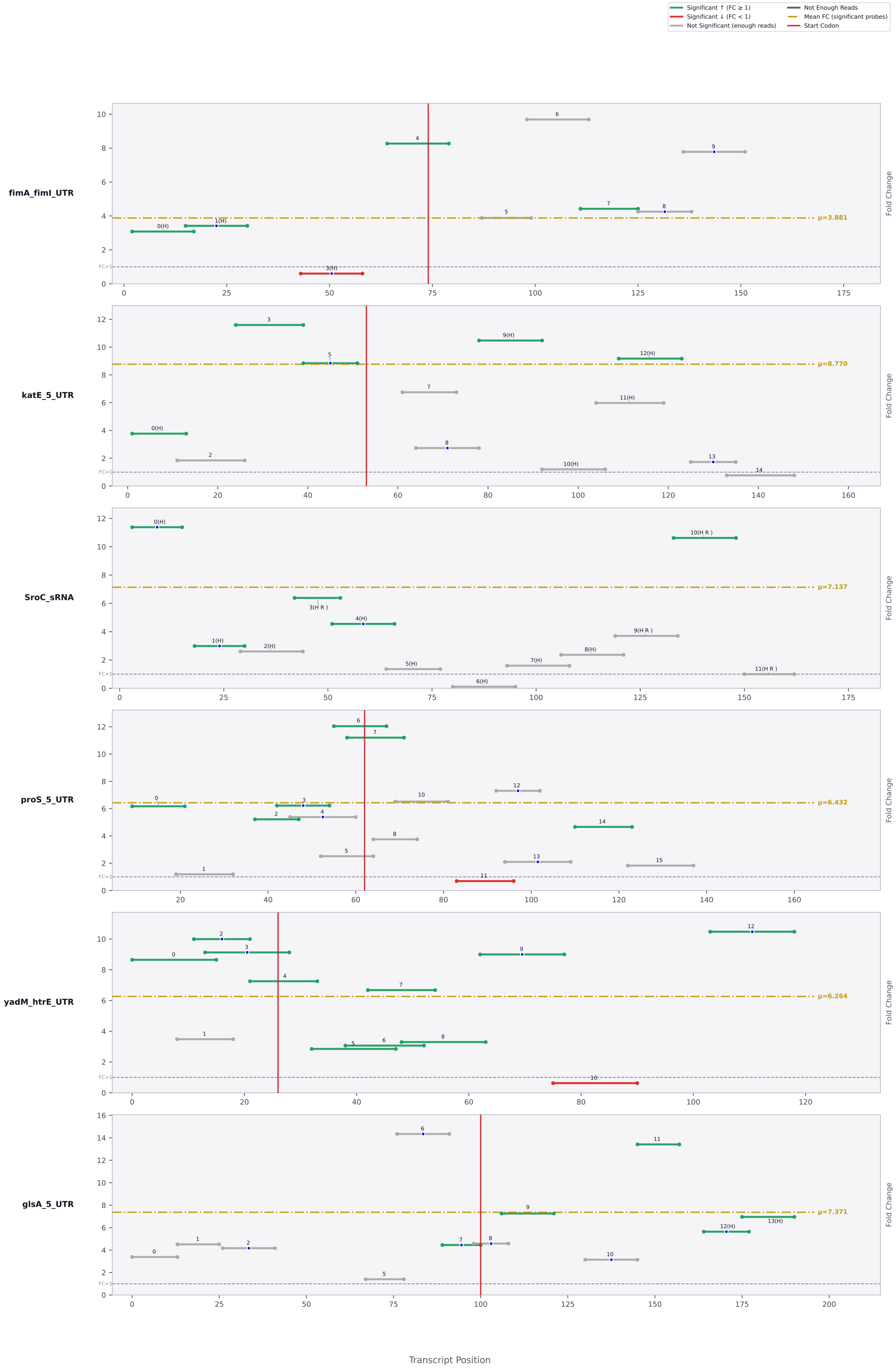

Significant ↑ (FC ≥ 1)
Significant ↓ (FC < 1)
Not Significant (enough reads)
Not Enough Reads
Mean FC (significant probes)
Start Codon
fimA_fimI_UTR
μ=3.881
katE_5_UTR
μ=8.770
SroC_sRNA
μ=7.137
proS_5_UTR
μ=6.432
yadM_htrE_UTR
μ=6.264
glsA_5_UTR
μ=7.371
Fold Change
FC=1
Transcript Position

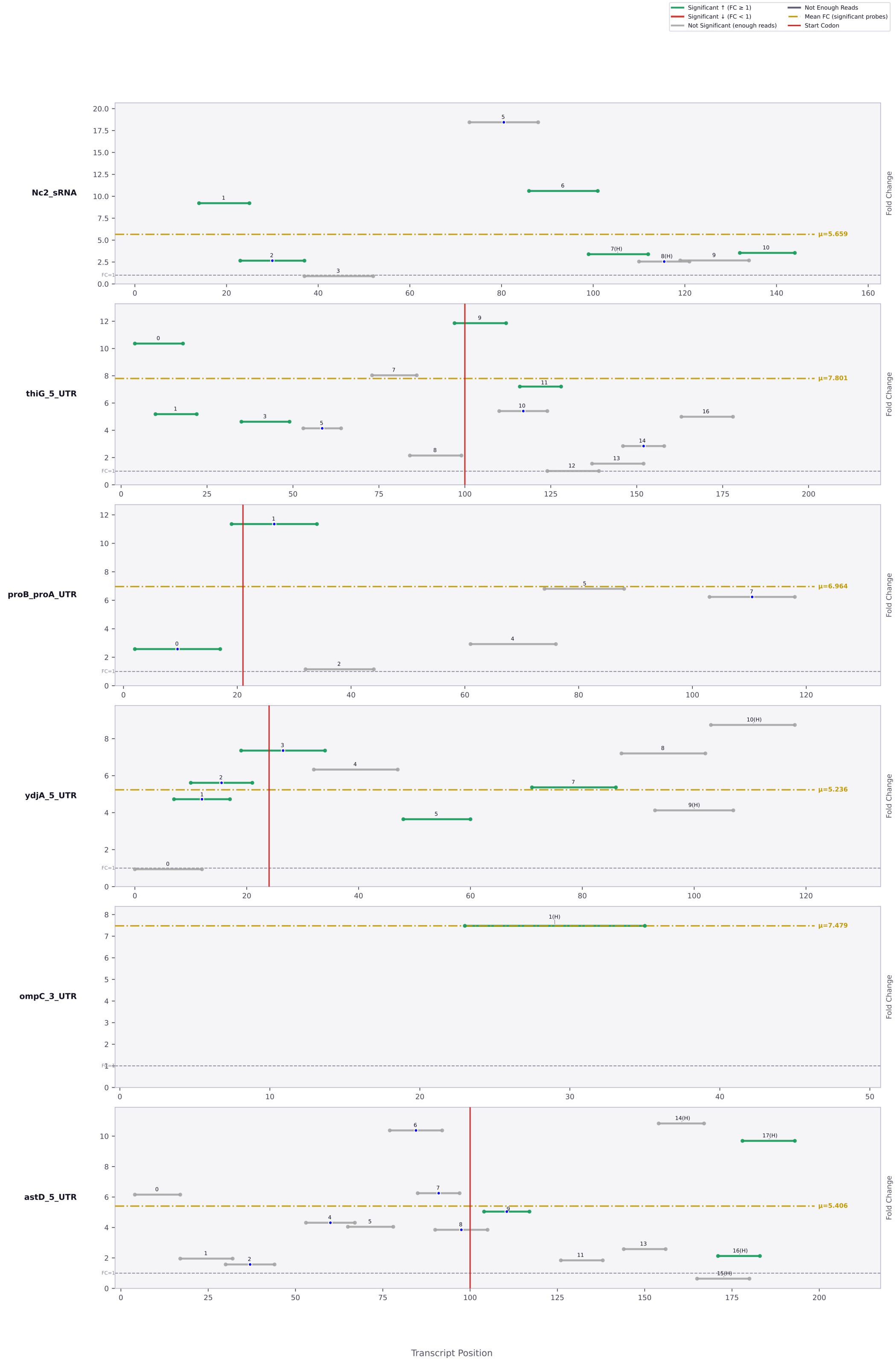

Significant ↑ (FC ≥ 1)
Significant ↓ (FC < 1)
Not Significant (enough reads)
Not Enough Reads
Mean FC (significant probes)
Start Codon
Nc2_sRNA
μ=5.659
thiG_5_UTR
μ=7.801
proB_proA_UTR
μ=6.964
ydjA_5_UTR
μ=5.236
ompC_3_UTR
μ=7.479
astD_5_UTR
μ=5.406
Fold Change
FC=1
Transcript Position

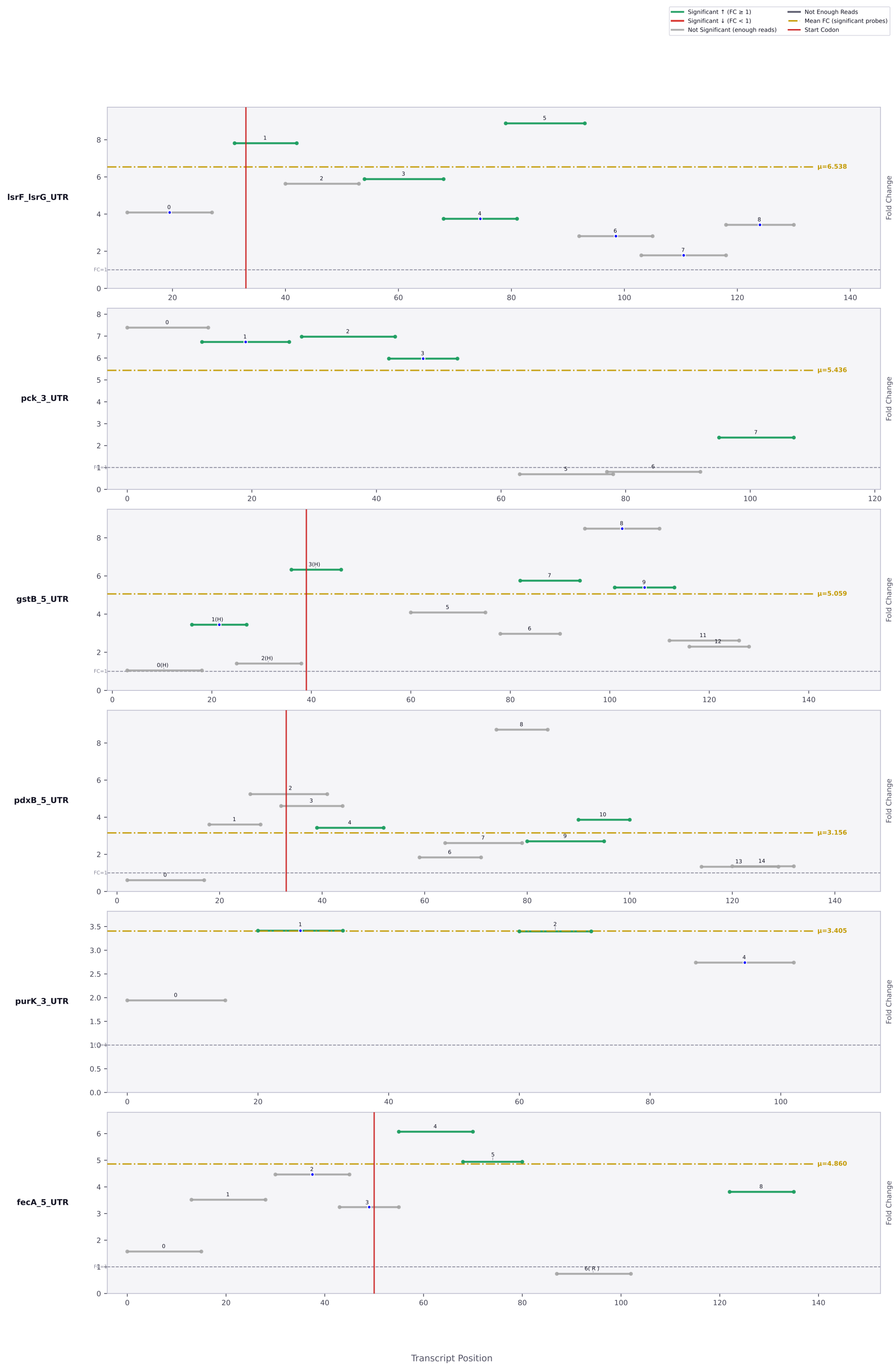

Significant ↑ (FC ≥ 1)
Significant ↓ (FC < 1)
Not Significant (enough reads)
Not Enough Reads
Mean FC (significant probes)
Start Codon
lsrF_lsrG_UTR
μ=6.538
pck_3_UTR
μ=5.436
gstB_5_UTR
μ=5.059
pdxB_5_UTR
μ=3.156
purK_3_UTR
μ=3.405
fecA_5_UTR
μ=4.860
Fold Change
Transcript Position

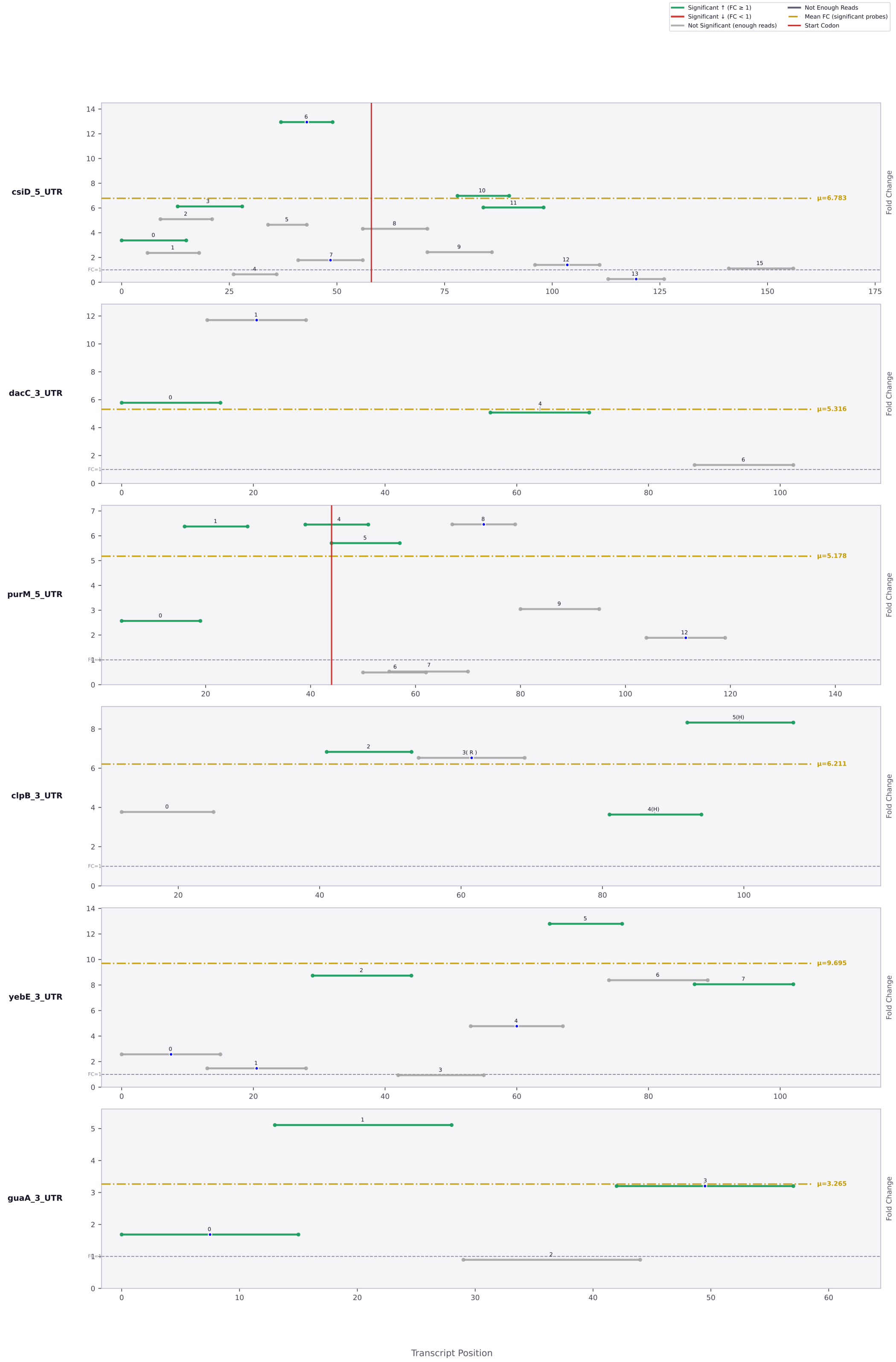

Significant ↑ (FC ≥ 1)
Significant ↓ (FC < 1)
Not Significant (enough reads)
Not Enough Reads
Mean FC (significant probes)
Start Codon
csiD_5_UTR
μ=6.783
dacC_3_UTR
μ=5.316
purM_5_UTR
μ=5.178
clpB_3_UTR
μ=6.211
yebE_3_UTR
μ=9.695
guaA_3_UTR
μ=3.265
Fold Change
FC=1
Transcript Position

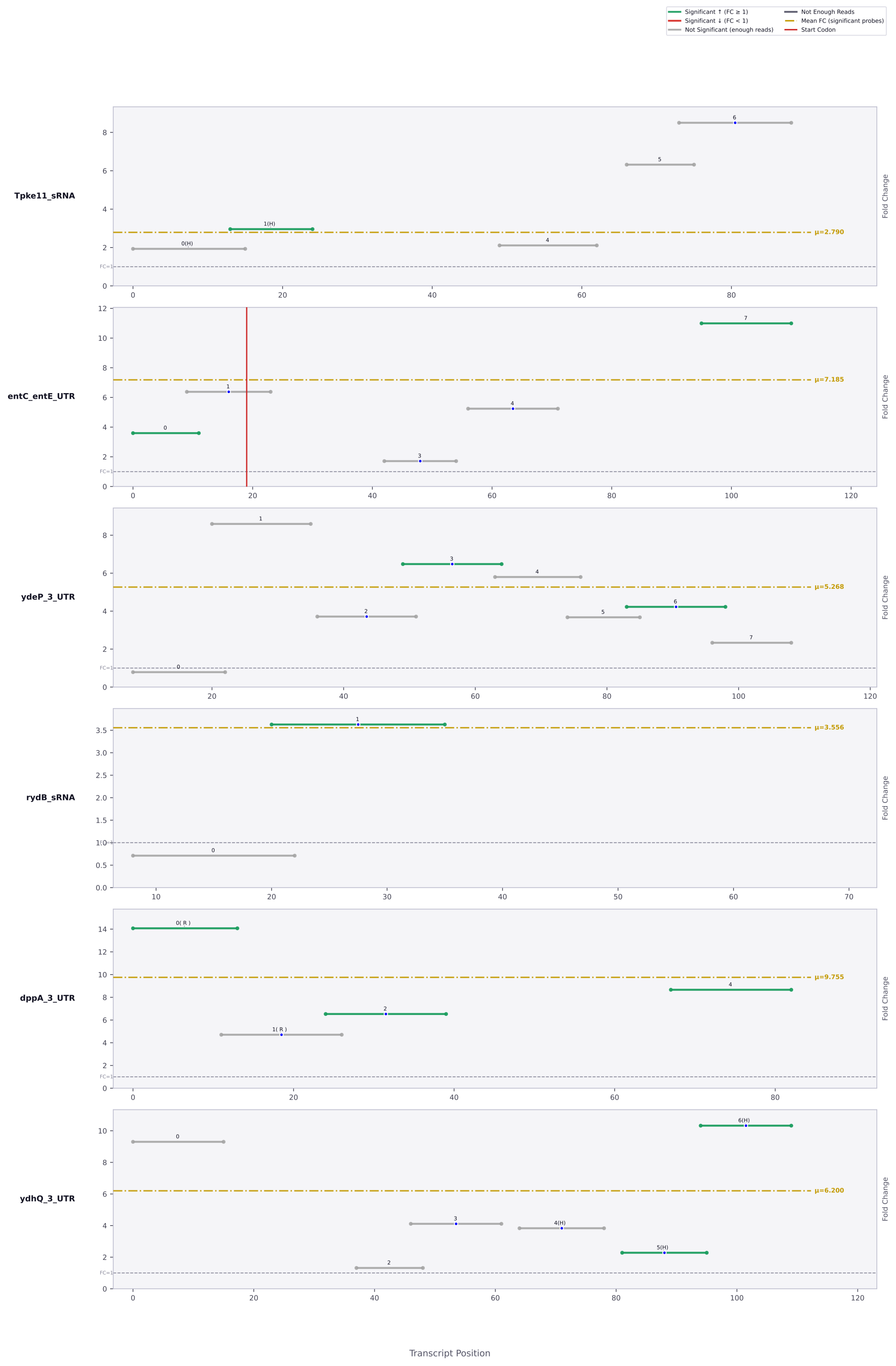

Significant ↑ (FC ≥ 1)
Significant ↓ (FC < 1)
Not Significant (enough reads)
Not Enough Reads
Mean FC (significant probes)
Start Codon
Tpke11_sRNA
μ=2.790
entC_entE_UTR
μ=7.185
ydeP_3_UTR
μ=5.268
rydB_sRNA
μ=3.556
dppA_3_UTR
μ=9.755
ydhQ_3_UTR
μ=6.200
Fold Change
Transcript Position

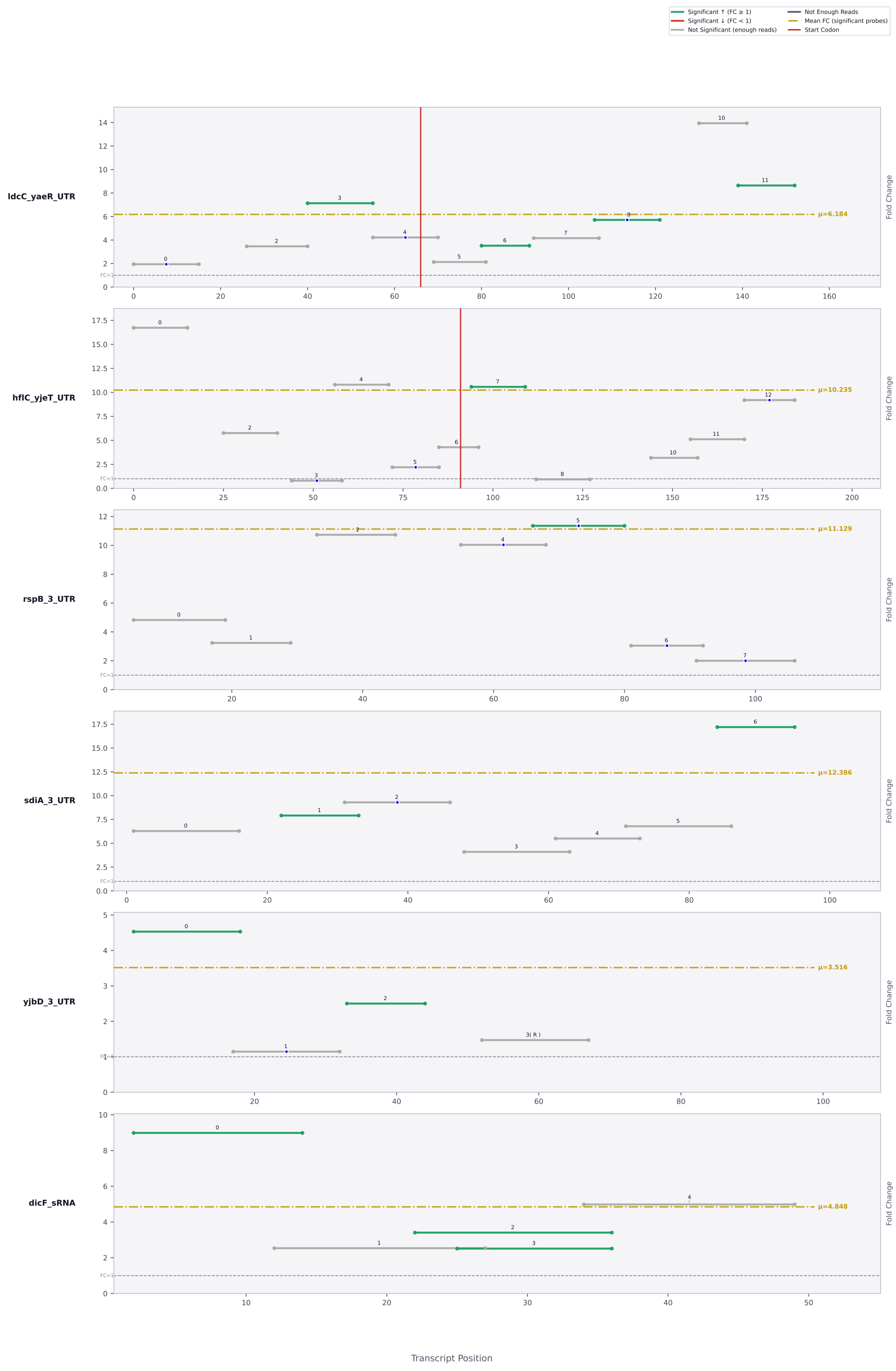
Significant ↑ (FC ≥ 1)
Significant ↓ (FC < 1)
Not Significant (enough reads)
Not Enough Reads
Mean FC (significant probes)
Start Codon
ldcC_yaeR_UTR
μ=6.184
hflC_yjeT_UTR
μ=10.235
rspB_3_UTR
μ=11.129
sdiA_3_UTR
μ=12.386
yjbD_3_UTR
μ=3.516
3( R )
dicF_sRNA
μ=4.848
FC=1
Fold Change
Transcript Position

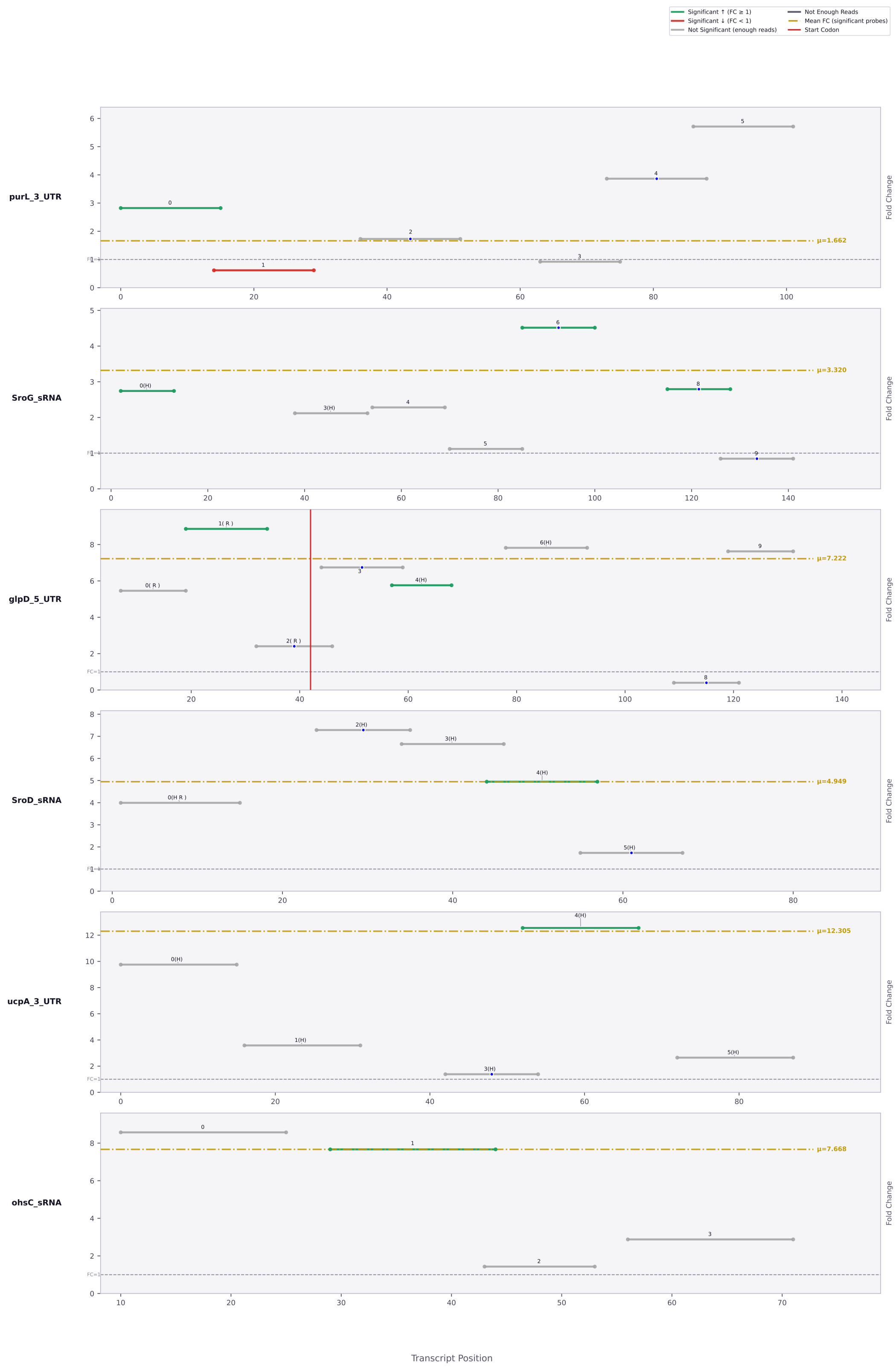

Significant ↑ (FC ≥ 1)
Significant ↓ (FC < 1)
Not Significant (enough reads)
Not Enough Reads
Mean FC (significant probes)
Start Codon
purL_3_UTR
μ=1.662
SroG_sRNA
μ=3.320
glpD_5_UTR
μ=7.222
SroD_sRNA
μ=4.949
ucpA_3_UTR
μ=12.305
ohsC_sRNA
μ=7.668
Fold Change
Transcript Position

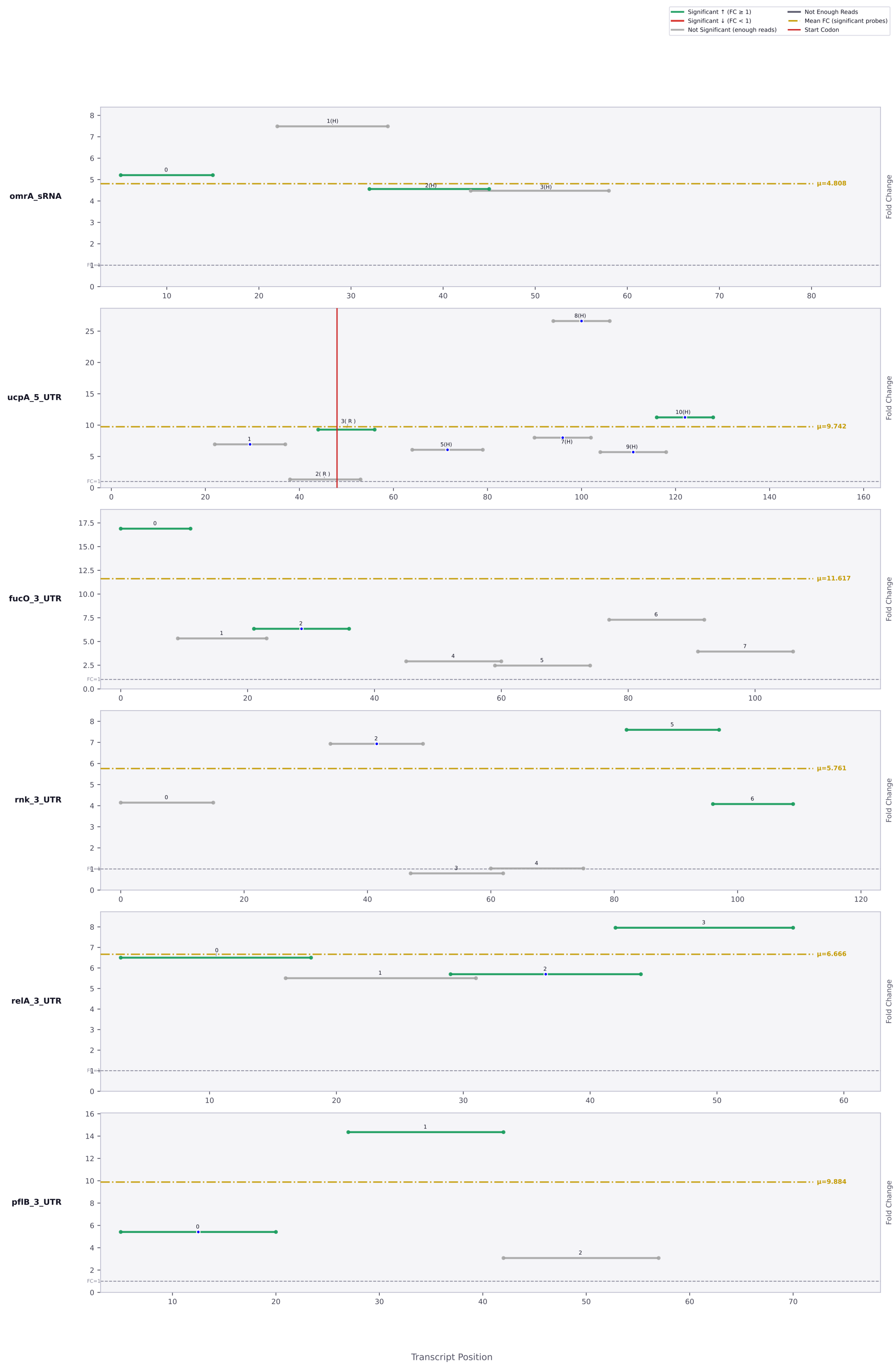

Significant ↑ (FC ≥ 1)
Significant ↓ (FC < 1)
Not Significant (enough reads)
Not Enough Reads
Mean FC (significant probes)
Start Codon
omrA_sRNA
μ=4.808
ucpA_5_UTR
μ=9.742
fucO_3_UTR
μ=11.617
rnk_3_UTR
μ=5.761
relA_3_UTR
μ=6.666
pflB_3_UTR
μ=9.884
Fold Change
Transcript Position

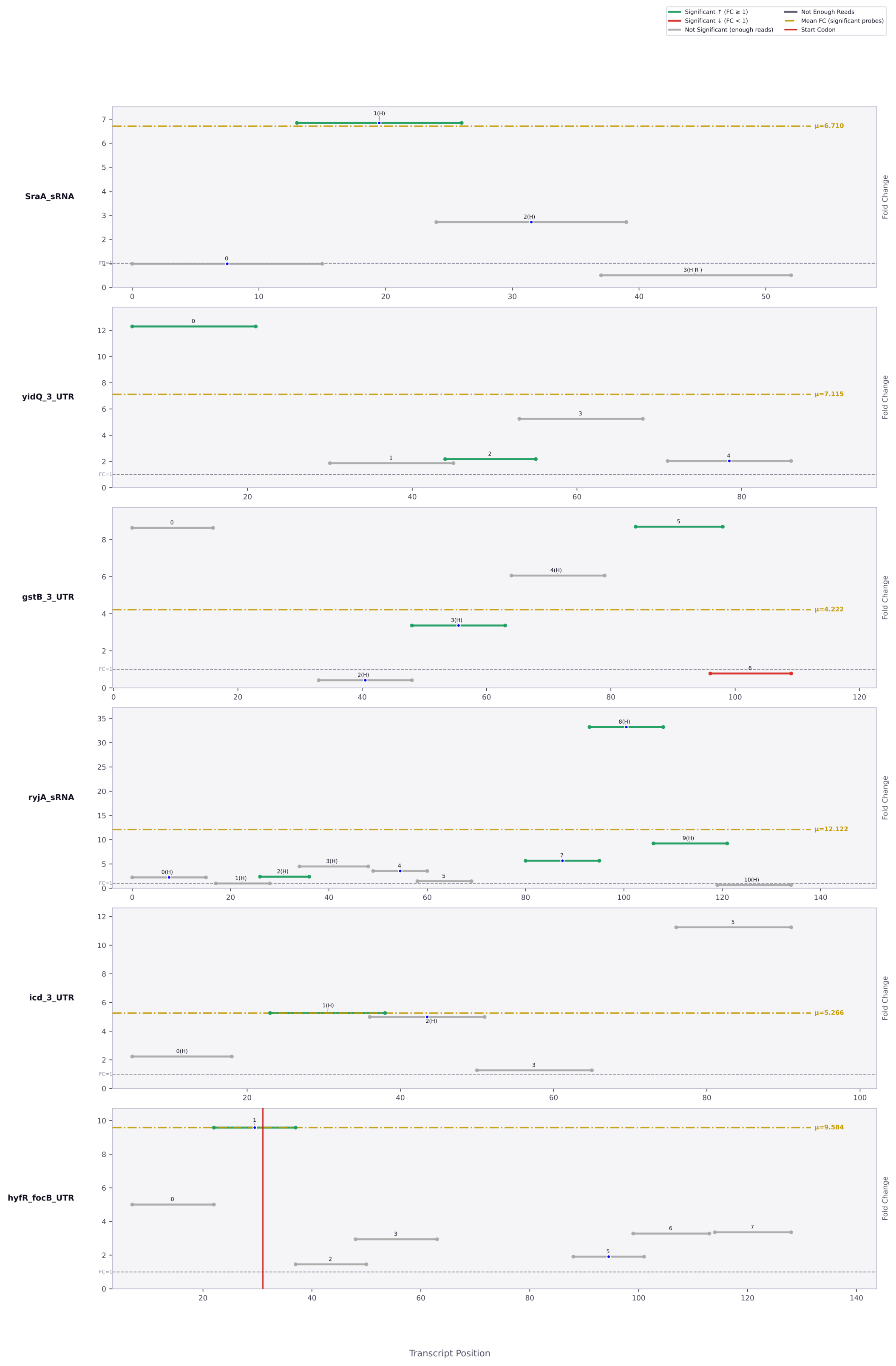

Significant ↑ (FC ≥ 1)
Significant ↓ (FC < 1)
Not Significant (enough reads)
Not Enough Reads
Mean FC (significant probes)
Start Codon
SraA_sRNA
μ=6.710
yidQ_3_UTR
μ=7.115
gstB_3_UTR
μ=4.222
ryjA_sRNA
μ=12.122
icd_3_UTR
μ=5.266
hyfR_focB_UTR
μ=9.584
Fold Change
Transcript Position

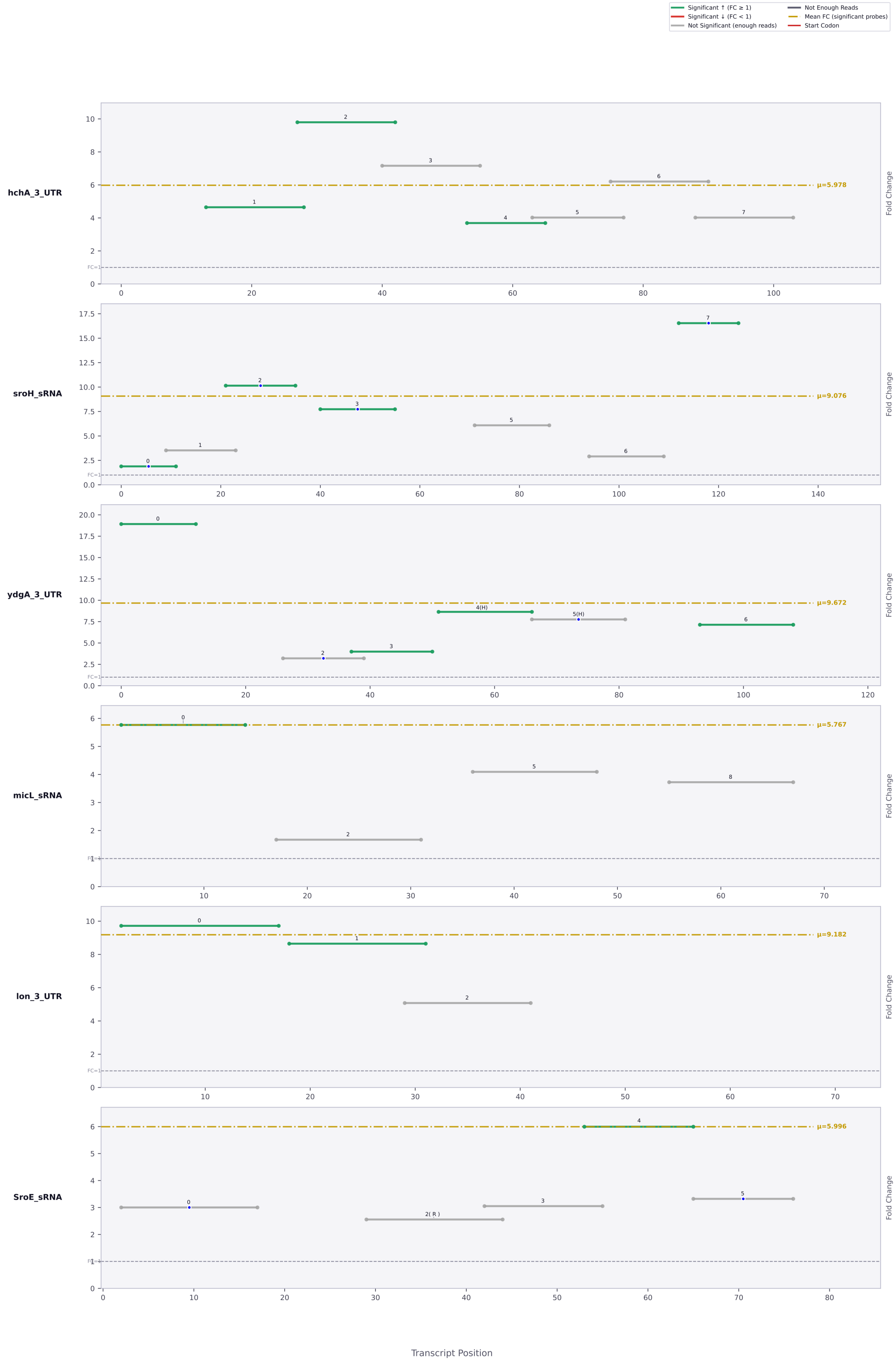

Significant ↑ (FC ≥ 1)
Significant ↓ (FC < 1)
Not Significant (enough reads)
Not Enough Reads
Mean FC (significant probes)
Start Codon
hchA_3_UTR
μ=5.978
sroH_sRNA
μ=9.076
ydgA_3_UTR
μ=9.672
micL_sRNA
μ=5.767
lon_3_UTR
μ=9.182
SroE_sRNA
μ=5.996
Fold Change
FC=1
Transcript Position

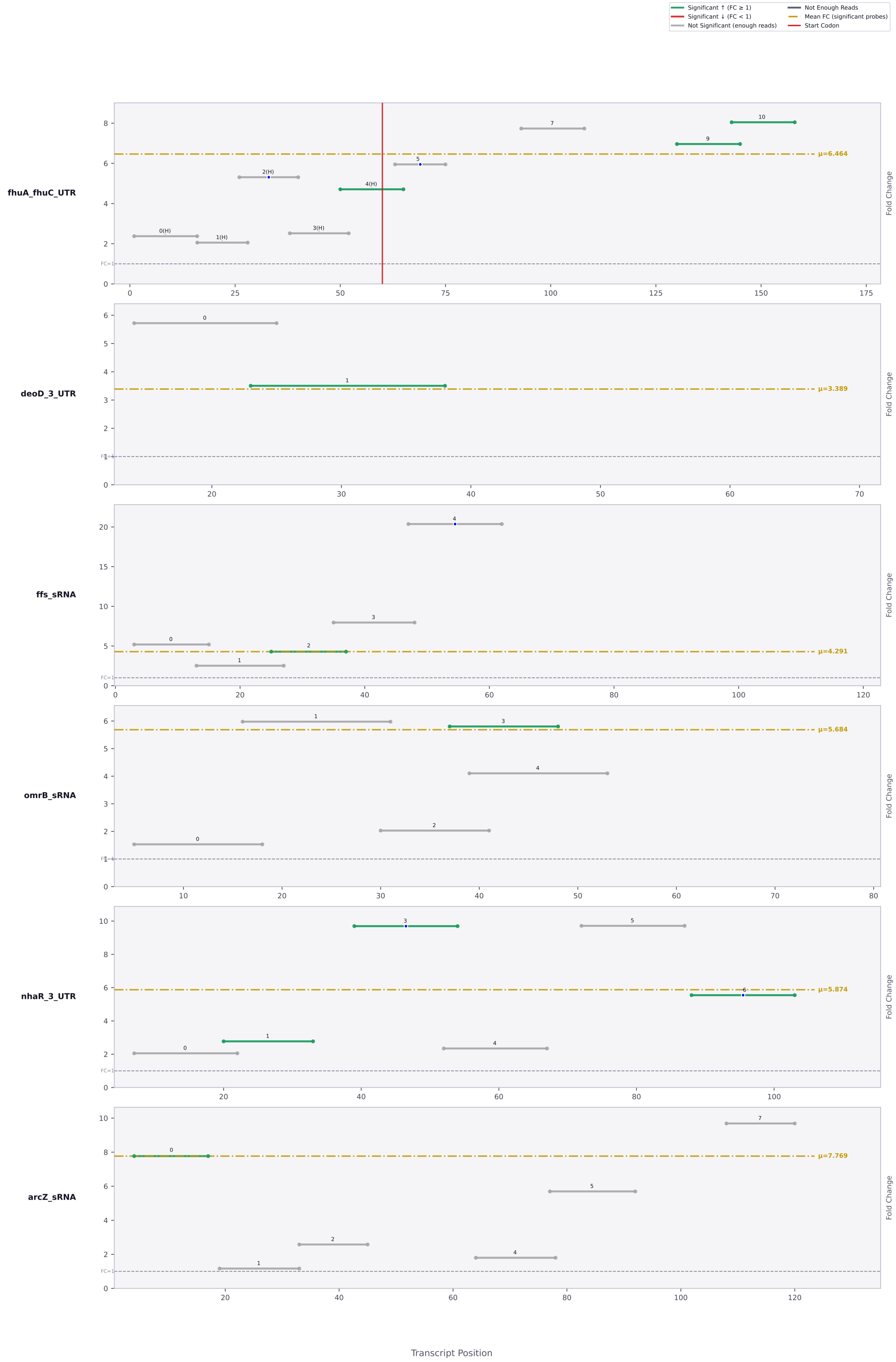

Significant ↑ (FC ≥ 1)
Significant ↓ (FC < 1)
Not Significant (enough reads)
Not Enough Reads
Mean FC (significant probes)
Start Codon
fhuA_fhuC_UTR
μ=6.464
deoD_3_UTR
μ=3.389
ffs_sRNA
μ=4.291
omrB_sRNA
μ=5.684
nhaR_3_UTR
μ=5.874
arcZ_sRNA
μ=7.769
FC=1
Fold Change
Transcript Position

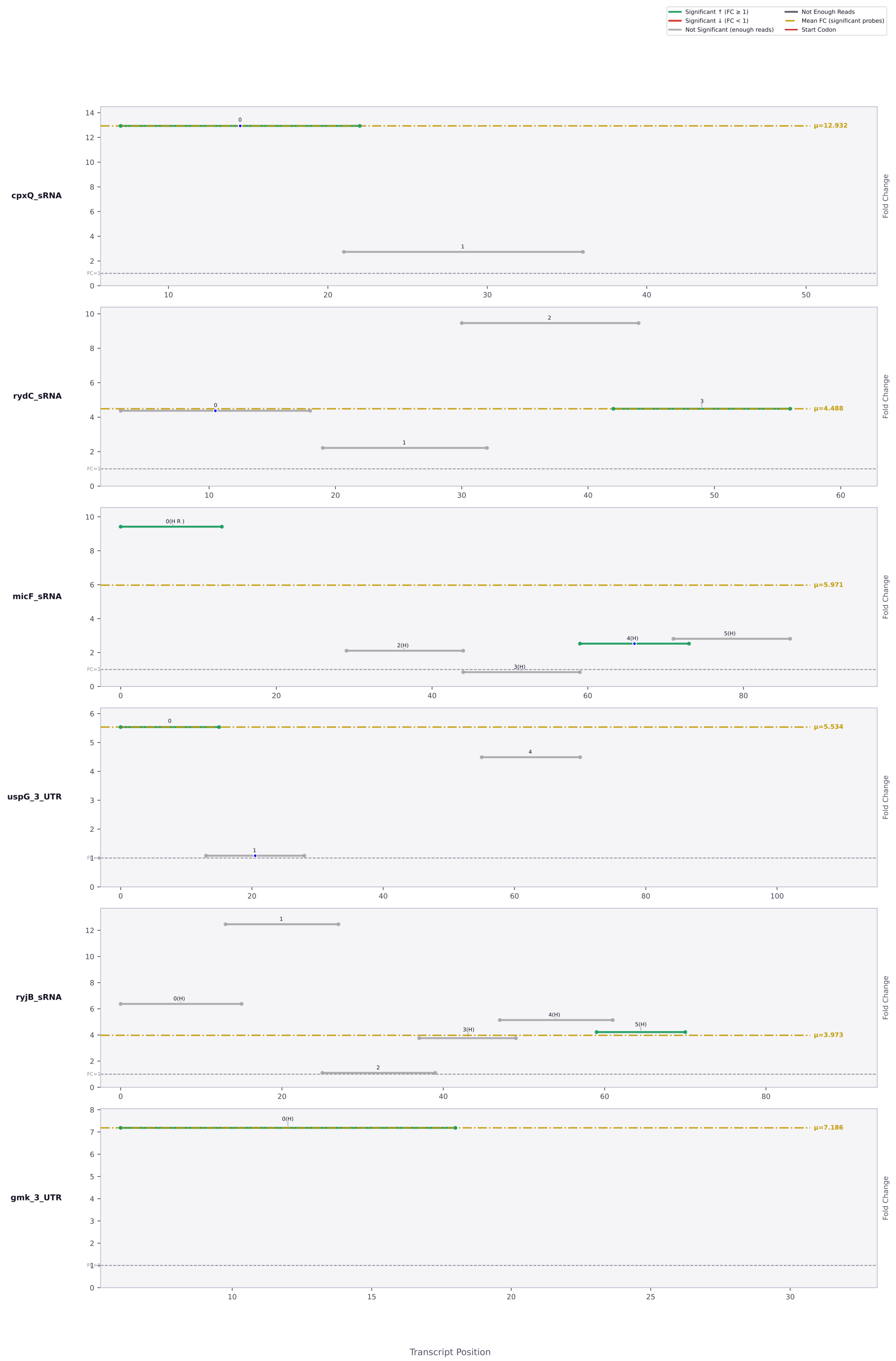
Significant ↑ (FC ≥ 1)
Significant ↓ (FC < 1)
Not Significant (enough reads)
Not Enough Reads
Mean FC (significant probes)
Start Codon
cpxQ_sRNA
μ=12.932
rydC_sRNA
μ=4.488
micF_sRNA
μ=5.971
uspG_3_UTR
μ=5.534
ryjB_sRNA
μ=3.973
gmk_3_UTR
μ=7.186
Fold Change
FC=1
Transcript Position

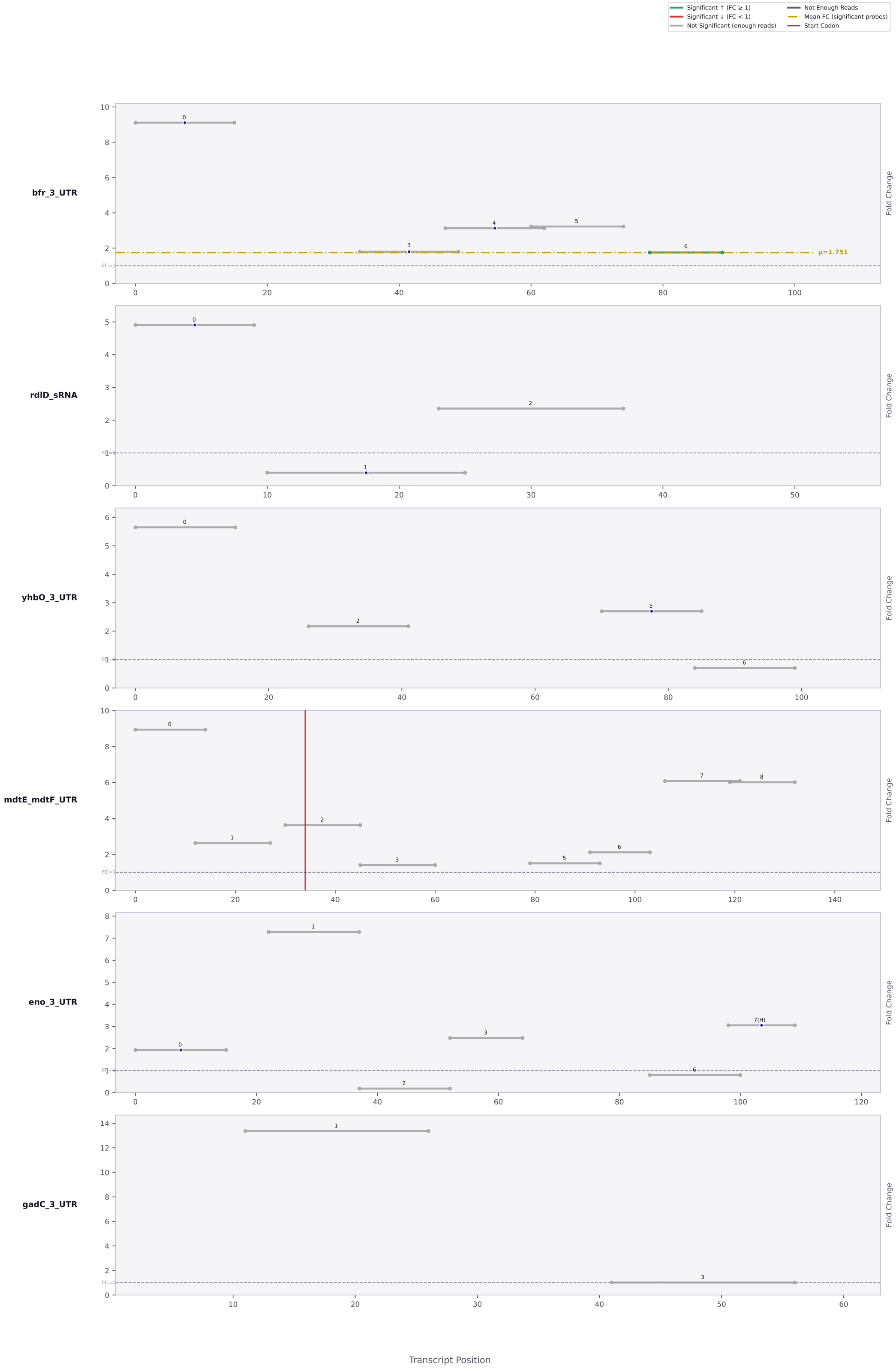
Significant ↑ (FC ≥ 1)
Significant ↓ (FC < 1)
Not Significant (enough reads)
Not Enough Reads
Mean FC (significant probes)
Start Codon
bfr_3_UTR
μ=1.751
rdlD_sRNA
yhbO_3_UTR
mdtE_mdtF_UTR
eno_3_UTR
gadC_3_UTR
Fold Change
FC=1
Transcript Position

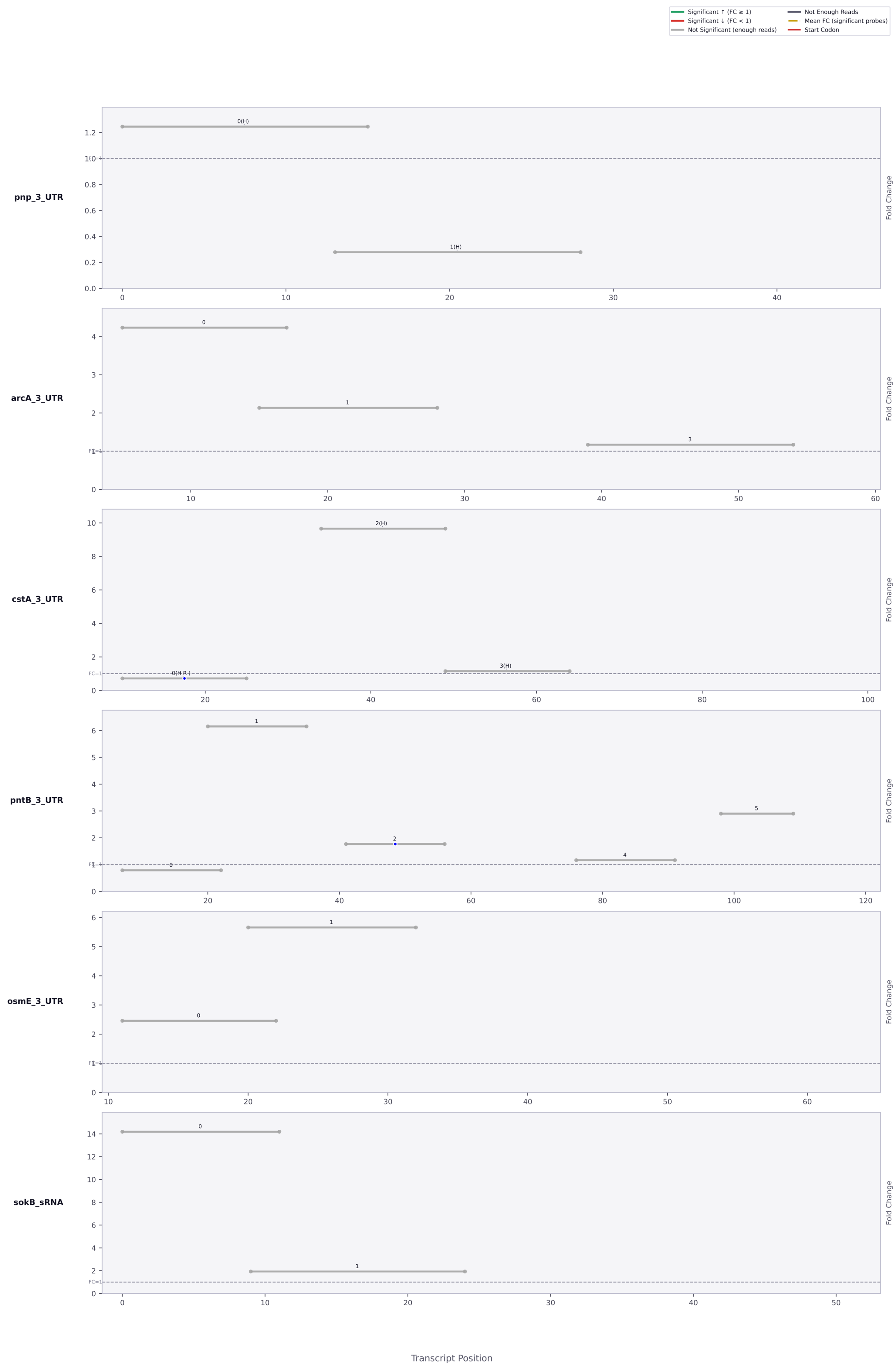

Significant ↑ (FC ≥ 1)
Significant ↓ (FC < 1)
Not Significant (enough reads)
Not Enough Reads
Mean FC (significant probes)
Start Codon
pnp_3_UTR
arcA_3_UTR
cstA_3_UTR
pntB_3_UTR
osmE_3_UTR
sokB_sRNA
Fold Change
Transcript Position

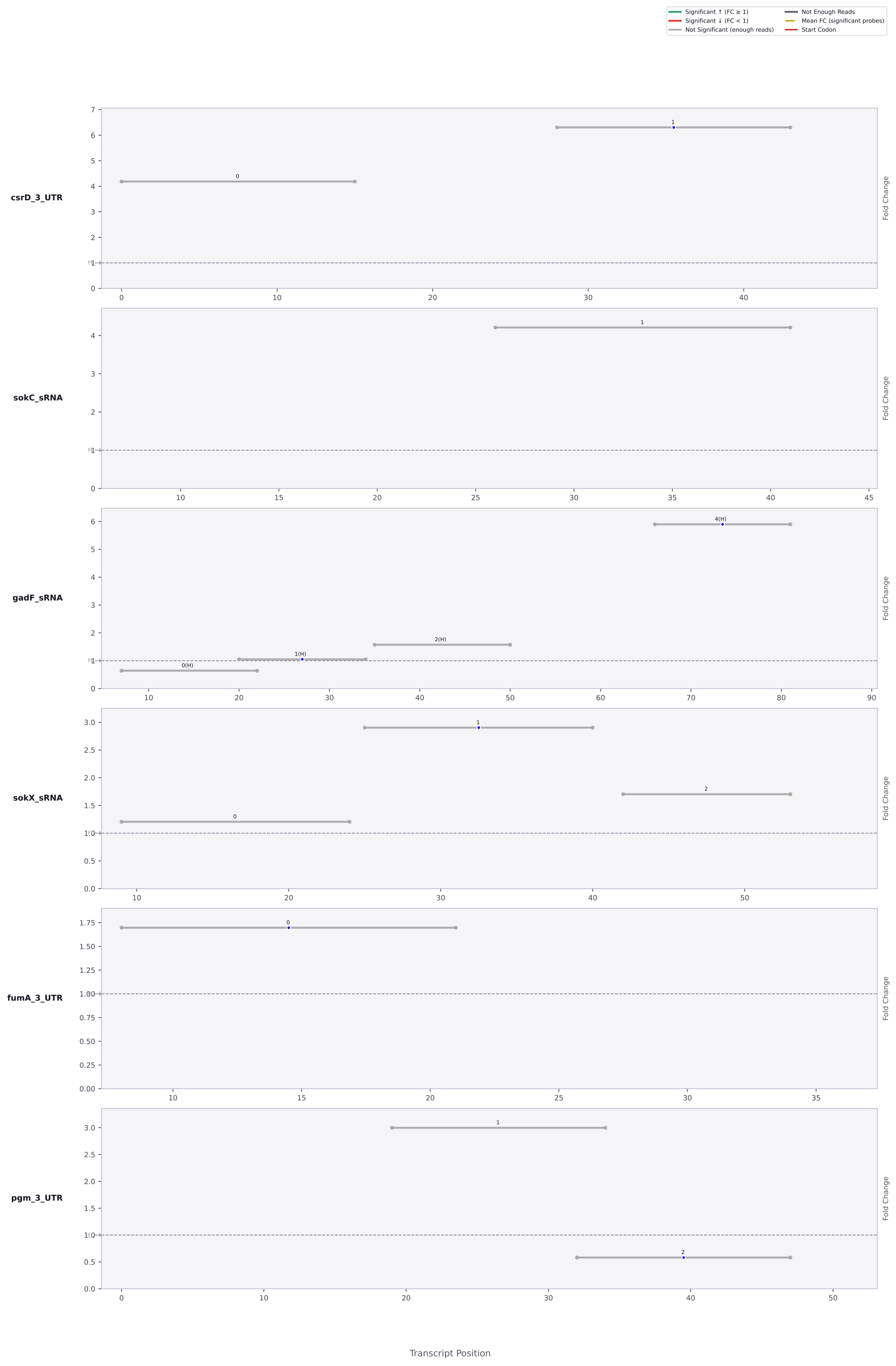

Significant ↑ (FC ≥ 1)
Significant ↓ (FC < 1)
Not Significant (enough reads)
Not Enough Reads
Mean FC (significant probes)
Start Codon
csrD_3_UTR
sokC_sRNA
gadF_sRNA
sokX_sRNA
fumA_3_UTR
pgm_3_UTR
Fold Change
Transcript Position

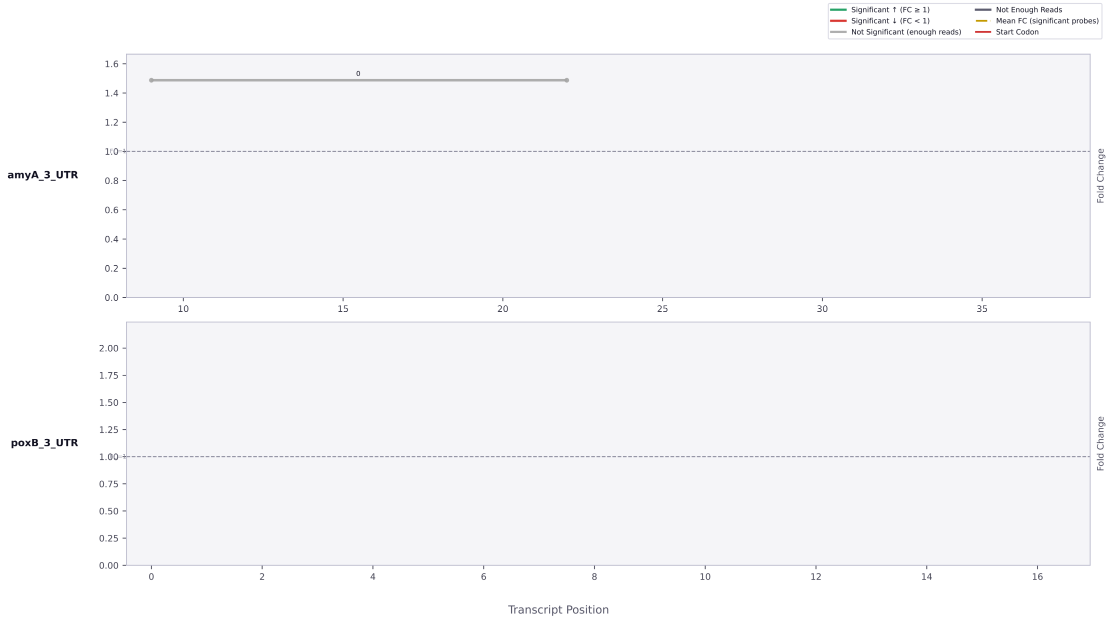

Significant ↑ (FC ≥ 1)
Significant ↓ (FC < 1)
Not Significant (enough reads)
Not Enough Reads
Mean FC (significant probes)
Start Codon
amyA_3_UTR
Fold Change
poxB_3_UTR
Fold Change
Transcript Position

***Figure S10.*** Accessibility map of all tested RNAs in this study. Each horizontal bar in the map of the transcript represents a single asRNA probe. The position on the x-axis signifies the location within the transcript, and the position on the y-axis represents the magnitude of the fold change between the *csrA:kan* and WT strains. Probes containing a blue dot are those that are antisense to a region containing an AGA or GGA motif, signifying a potential CsrA binding site. The color of each bar represents the statistical significance of the probe between the two strains, with green/red bars being statistically significant, and gray bars not significant. The vertical red bar signifies the start codon. The horizontal yellow bar indicates the mean accessibility change of the transcript.
